# A Covalent PFKL Activator Suppresses Tumor Growth

**DOI:** 10.1101/2025.10.02.680014

**Authors:** Xiaoding Jiang, Eric M. Lynch, Congcong Lyu, Crystal Wilson, Lauren E. Salay, Scott Lyons, Mu-Jie Lu, Shuangyu Luo, Gibae Kim, Hsin-Ru Chan, Wes Wolfe, Yi-Chih Lin, Justin M. Kollman, Xiaolu A. Cambronne, Ku-Lung Hsu

## Abstract

Glycolysis fuels vital cellular functions and its dysregulation is implicated in cancer, neurodegeneration, antibiotic resistance and diabetes. The glycolytic dependency of cancer, known as the Warburg effect, presents a key vulnerability for developing targeted anticancer agents but remains challenging due to metabolic heterogeneity and resistance. Here, we developed a first-in-class covalent phosphofructokinase-1 liver type (PFKL) activator that induces metabolic imbalance coupled to delivery of a cytotoxic payload to cancer cells *in vitro* and *in vivo*. The electrophile-drug conjugate (EDC) site-specifically and proteome-wide selectively modifies K677 in the allosteric effector site to stabilize the R-state tetramer of PFKL and destabilize cell metabolism. We introduce EDCs as a new delivery mechanism analogous to antibody-drug conjugates but differentiated by selective covalent targeting of intracellular proteins.

## INTRODUCTION

Cell metabolism is essential for life by supplying fuel, signaling molecules and building blocks required for growth, survival and adaptation to environmental fluctuations (*1*). Dysregulation of metabolic pathways has emerged as a hallmark of various diseases, including cancer (*2*), diabetes (*3*), neurodegenerative disorders (*4*) and metabolic syndromes (*5*). Notably, many cancer cells adopt a metabolic phenotype with increased dependence on aerobic glycolysis despite availability of oxygen. This metabolic reprogramming, also known as the Warburg effect (*6*), drives increased glucose uptake and lactate production, supporting rapid cell proliferation and adaptation to nutrient-deprived or hypoxic environments (*7*).

While glycolytic rewiring facilitates tumor growth and survival, it concurrently creates vulnerabilities that can be exploited therapeutically (*8*). Beyond catabolism, altered glucose metabolism generates key metabolic intermediates that serve as signaling molecules, orchestrating diverse cellular processes including gene expression, epigenetic modifications, and redox homeostasis (*9–12*). Excessive accumulation of glycolytic intermediates in cancer cells has been reported to induce imbalances in bioenergetics and impair tumor growth (*13, 14*). Alterations in tumor microenvironment (e.g. pH) or intracellular enzymes (e.g., GAPDH) differentially regulated during the Warburg effect have been targeted as anticancer strategies (*15, 16*). Nevertheless, inhibiting the upregulation of glycolysis in cancer cells remains a challenge due to their inherent metabolic plasticity, including resistance through utilization of alternative carbon sources such as amino acids to support growth (*17*).

Glycolytic flux is regulated principally by phosphofructokinase-1 (PFK1) activity. PFK1 catalyzes the ATP-dependent phosphorylation of fructose 6-phosphate (F6P) to fructose 1,6-bisphosphate (FBP) (*18*). PFK1 is the rate-limiting enzyme in glycolysis and tightly regulated by:

1. cellular energy status, where a low ATP/ADP ratio activates PFK1 and a high ratio is inhibitory, post-translational modifications (PTMs) such as phosphorylation (*19*), acetylation (*20*), and glycosylation (*21*), (iii) allosteric modulators that stabilize its tetrameric conformational states (*22*). In mammals, PFK1 is expressed as three isoforms (PFKL, PFKM, PFKP) with high sequence homology but distinct allosteric regulation (*23*) and moonlighting functions (*24, 25*). Genetic mutations resulting in increased or decreased activity of individual PFK1 isoforms has been implicated in cancer cells (*18, 26*). Collectively, PFK1 activity could be exploited for anticancer applications but requires selective targeting modalities that can address the metabolic plasticity of cancer cells.

Herein, we developed a covalent, allosteric PFKL activator (**XJ-4-85**) loaded with a cytotoxic leaving group as an anticancer modality. **XJ-4-85** site-specifically bound K677 and its leaving group interacted with the nucleotide effector site to stabilize the active R-state tetramer of PFKL as determined by a cryo-electron microscopy structure (3.2 Å resolution). Quantitative chemoproteomics revealed excellent PFK1 isoform and proteome-wide selectivity of **XJ-4-85** in cells. **XJ-4-85** treatments resulted in glycolytic reprogramming and reduced tumor burden *in vivo* compared with the free leaving group, presenting a rationale for electrophile-drug conjugates (EDCs) as a payload delivery strategy targeting intracellular metabolic proteins and vulnerabilities.

## RESULTS

### Discovery of a site-specific, proteome-wide selective covalent PFKL activator

Sulfur-triazole exchange (SuTEx) chemistry has emerged as a versatile platform for developing covalent inhibitors that target functional tyrosine and lysine residues on proteins (*27–29*). In a prior study, we developed the alkynyl probe **TH211** by embedding a kinase binding fragment RF001 into a reactive SuTEx electrophile, allowing broad chemoproteomic profiling of kinases in cells (*30*). Whether RF001-based scaffolds can be optimized into selective ligands for targeting kinases remains largely unexplored. To address this question, we synthesized a library of **TH211** analogues bearing various modifications as starting points for optimizing targeted covalent ligands of kinases (fig. S1). To identify hits capable of engaging target protein(s) in dynamic cellular environments, we conducted gel-based competitive activity-based protein profiling (ABPP) *in situ*. HEK293T cells were treated with SuTEx ligand (1 μM, 1 h) followed by **TH211** probe labeling (25 μM, 2 h) and reduced fluorescent protein labeling from probe competition indicated cellular target engagement.

Notably, SuTEx ligands featuring electron-donating groups, such as methyl or methoxy, on the *para* position of the phenyl sulfone moiety effectively blocked labeling of a ∼85 kDa protein (Fig. S2A). Next, we deployed a competitive tandem mass tag (TMT)-ABPP assay (*31*) to deduce the molecular identity of the ∼85 kDa cellular target (fig. S3). Probe-enriched sites detected by LC-MS/MS and exhibiting significant competition with ligand pretreatment compared to DMSO were designated as liganded sites (competition ratio or CR < 0.5, *P* < 0.05; fig. S2B). By cross referencing these liganded sites with the molecular weights of corresponding proteins, the glycolytic enzyme PFKL was identified as a prominent liganded target exhibiting a competition profile congruent with the gel-based analysis (fig. S2, A and B). Among the active hits, **TH220** demonstrated superior potency (IC_50_ = 305 nM, fig. S2, C and D) for blockade of PFKL probe labeling that was site specific (K677) and isoform selective as evidenced by negligible activity against the homologous lysine site on PFKP (K688, fig. S2, C to E, fig. S4, and Table S1).

To investigate the impact of **TH220** on PFKL biochemical activity, we employed a reported substrate assay using purified His-tagged PFKL and monitored conversion of F6P to FBP (*32*) (fig. S5A). K677 resides in the nucleotide effector site of PFKL and covalent modification at this residue with electrophilic natural products was previously shown to inactivate PFKL (*33*). Unexpectedly, the K677-directed ligand **TH220** activated PFKL in a dose-dependent manner with a moderate half maximal effective concentration (EC_50_) of 56 µM (fig. S5B). Structure–activity relationship (SAR) exploration revealed (i) molecular flexibility and length of the linker, and (ii) introduction of additional fluorine atoms as hydrogen bond acceptors as key for activity (Fig. 1A and fig. S5, C and D). These efforts produced the optimal ligand **XJ-4-85**, which exhibited dose- and time-dependent activation of PFKL (Fig. 1B). Notably, the leaving group alone (**XJ-4-119)** or analogs with a different recognition motif (**XJ-4-97)** did not activate PFKL, identifying the latter as a suitable negative control (Fig. 1, A and B, Table S2).

**Fig. 1.**
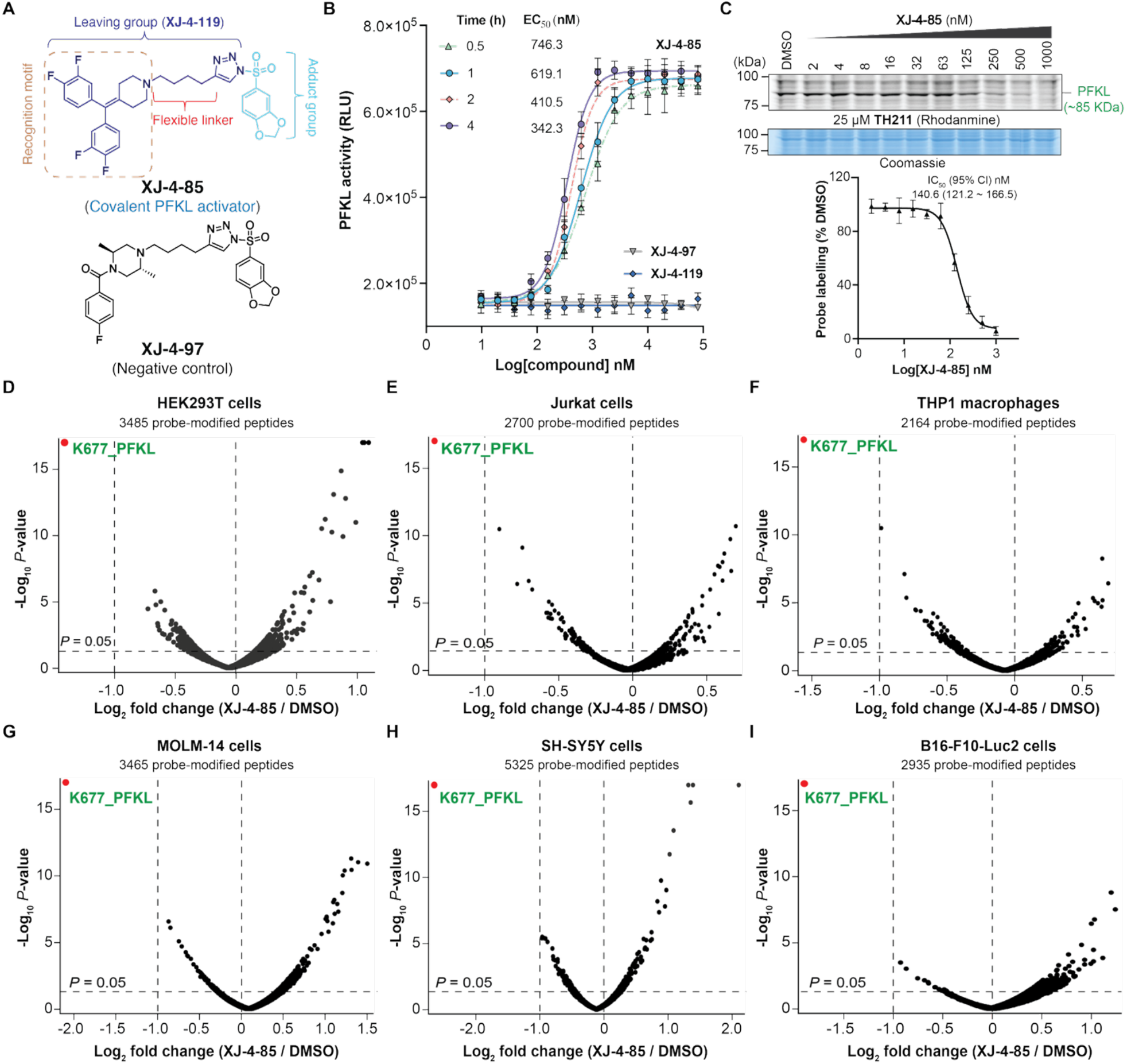
Discovery of a site-specific, proteome wide-selective covalent PFKL activator. (A) Chemical structures of **XJ-4-85, XJ-4-97** (negative control) and **XJ-4-119** (Leaving group). (B) Dose-and time-dependent biochemical activation of PFKL (WT) by **XJ-4-85** compared to **XJ-4-97** and **XJ-4-119** using a substrate assay. (C) Gel-based competitive ABPP evaluation of **XJ-4-85** binding activity in HEK293T cells. Representative data from n = 4 replicates are shown. (D-I) Competitive TMT-ABPP analysis of proteomes from **XJ-4-85**-treated cells (5 µM, 1 h) to demonstrate PFKL site specificity (K677) and proteome-wide selectivity. Data shown are representative of n = 3 biological replicates. See Table S1 for detailed proteomic data.

We performed gel-based competitive ABPP and determined **XJ-4-85** potently engaged PFKL in HEK293T cells (IC_50_ = 141 nM, Fig. 1C). Next, we performed competitive TMT-ABPP to assess PFKL selectivity of **XJ-4-85** in treated cells. Across ∼3,000 quantified probe-modified peptides, PFKL K677 was the only significantly liganded site detected with **XJ-4-85** treatment (CR < 0.5, *P* < 0.05, Fig. 1D). This exquisite selectivity was not cell line or species dependent as evidenced by nearly identical findings in human (Jurkat, THP1, MOLM-14, SH-SY5Y) and mouse (B16-F10-Luc2) cell lines (Fig. 1, E to I). These competition events were not due to PFKL expression changes (fig. S6). Collectively, our findings establish **XJ-4-85** as a site specific (K677), proteome-wide selective covalent activator that effectively engages PFKL but no other detectable PFK1 isoforms across a panel of diverse cell lines.

### Structural insights into PFKL activation by XJ-4-85

To decipher the activation mechanism by **XJ-4-85**, we solved the cryo-electron microscopy (cryo-EM) structure of PFKL in complex with **XJ-4-85** and substrates ATP and F6P (PDB ID: 9P0J; Fig. 2A and fig. S7 and S8). We performed 3D classification to identify the highest quality PFKL tetramers, and subsequently performed D2 symmetry expansion, 3D classification, and local refinement to identify the highest quality monomers in the dataset. The monomer classes that reached the highest resolution (3.2-3.3Å) exhibited clear density for the **XJ-4-85** leaving group in the allosteric activating site between the PFKL catalytic and regulatory domains, adjacent to residue K677 (Fig. 2B). The covalent moiety was less well-resolved, with additional density next to K677 only apparent in a subset of the monomer classes, suggesting flexibility in the covalently modified residue (Fig. 2B). The **XJ-4-85** binding pose overlapped with those of native activators (AMP and ADP) and the reversible PFKL activator NA-11 (*34*) (Fig. 2C). The linear alkyl chain was not resolved, presumably due to conformational flexibility, and therefore was not included in the atomic model (Fig. 2).

**Fig. 2.**
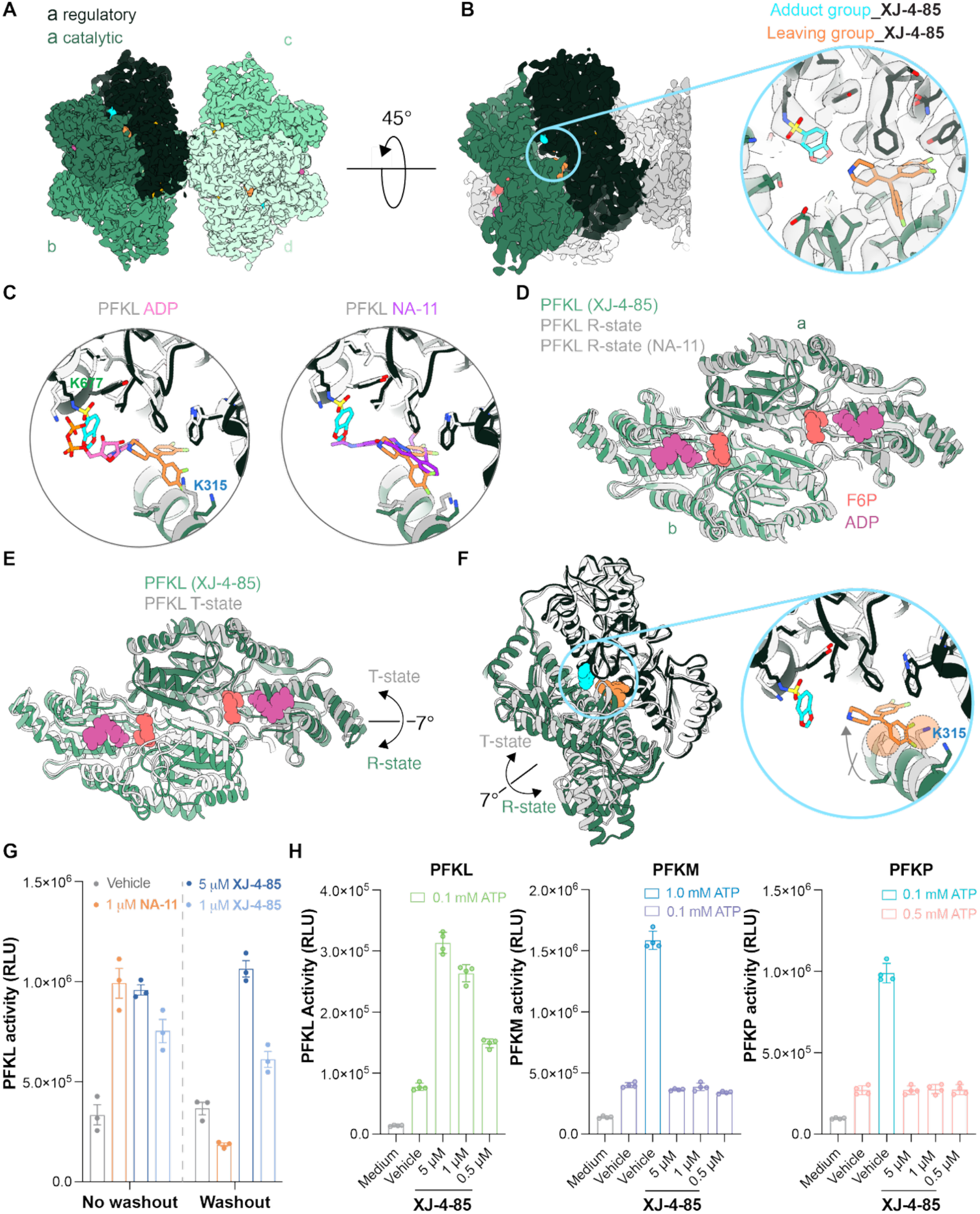
Cryo-EM structure of XJ-4-85 bound to PFKL. (A) Cryo-EM structure of a PFKL tetramer bound to **XJ-4-85**, colored by monomer. Monomer-a is colored by domain. (B) Cryo-EM structure of a PFKL monomer bound to **XJ-4-85** (PDB: 9P0J). Inset shows an expanded view of the **XJ-4-85** adduct and leaving group binding sites. (C) Overlay of **XJ-4-85** with ADP (PDB: 8W2G) and NA-11 (PDB: 7LW1) in the allosteric activating site. (D-E) Catalytic domain dimer from the **XJ-4-85**-bound PFKL structure compared with PFKL in the R-state (PDB: 8W2G, PDB: 7LW1) and T-state (PDB: 8W2H). Structures are aligned on subunit-a. F6P and ADP are shown in the active sites of the **XJ-4-85**-bound structure. PFKL bound to **XJ-4-85** is in the R-state conformation, which involves a 7° rotation between catalytic domains relative to the T-state conformation. (F) Monomers of **XJ-4-85**-bound and T-state PFKL aligned on their regulatory domains. Transition to the T-state involves a 7° rotation between the catalytic and regulatory domains, which is sterically inhibited by the **XJ-4-85** leaving group. Orange circles in the expanded view highlight residues that would clash with the **XJ-4-85** leaving group in the T-state conformation. (G) Washout experiment demonstrated the irreversible binding of **XJ-4-85**. (H) PFK1 isoform specificity of **XJ-4-85** using a substrate assay. See Table S2 for details.

To understand the effect of **XJ-4-85** on the overall conformation of PFKL, we performed a local refinement using particles from the best monomer class, expanding the refinement mask to encompass a full tetramer (Fig. 2A). The resulting structure showed **XJ-4-85**-bound PFKL was in the active R-state conformation, with F6P and ADP bound at active sites, aligning with the activating function of **XJ-4-85** (Fig. 2D and fig. S8). As previously described for R-state PFKL, the allosteric sugar-binding sites were also occupied and we modelled these ligands as FBP, likely produced by PFKL activity during sample preparation (fig. S8). The active R-state conformation contrasts with the inactive T-state conformation, where the catalytic domains rotate to disrupt the enzyme active sites (Fig. 2E). The R-to T-state transition also involved a rotation between the catalytic and regulatory domains within monomers, which compresses the allosteric activating site in the T-state conformation (Fig. 2F). Akin to AMP, ADP, and NA-11, **XJ-4-85** appeared to activate PFKL by sterically preventing compression of the allosteric activating site, thereby resisting transition to the T-state conformation. Rotation to the T-state would result in steric clashes between the **XJ-4-85** leaving group and incoming catalytic domain (Fig. 2F).

We conducted a washout experiment after preincubation with compounds and showed that PFKL activation was retained with **XJ-4-85** but not the reversible ligand NA-11 (Fig. 2G). These data combined with the lack of activity for the free leaving group, support covalency as a requirement for **XJ-4-85**-mediated PFKL activation (Fig. 1B and 2G). We tested and found that **XJ-4-85** failed to activate FPKP or PFKM despite these isoforms containing a homologous lysine residue (K688 and K678, respectively) for covalent binding (Fig. 2H and fig. S9, A and B). To further probe PFK1 isoform-selective ligand recognition, we mutated the non-conserved K315 located in proximity to the bound leaving group, which abolished **XJ-4-85**-mediated PFKL activation (fig. S9, A and C).

Collectively, our studies reveal **XJ-4-85** functions as an allosteric activator of PFKL that achieves isoform selectivity through covalent (K677) and non-covalent (K315) binding distinct from the reported reversible PFKL activator (*34*).

### XJ-4-85 reprograms glycolysis across multiple cell types

Detection of intracellular FBP is widely utilized as an indicator of glycolytic pathway activity, as its intracellular concentration closely reflects the rate of glycolysis as the product of PFK1. Here we investigated whether **XJ-4-85** can activate endogenous PFKL in cells using the genetically encoded single-fluorescent protein HYlight biosensor (*35*) to monitor levels of FBP, the direct product of PFKL (Fig. 3A).

**Fig. 3.**
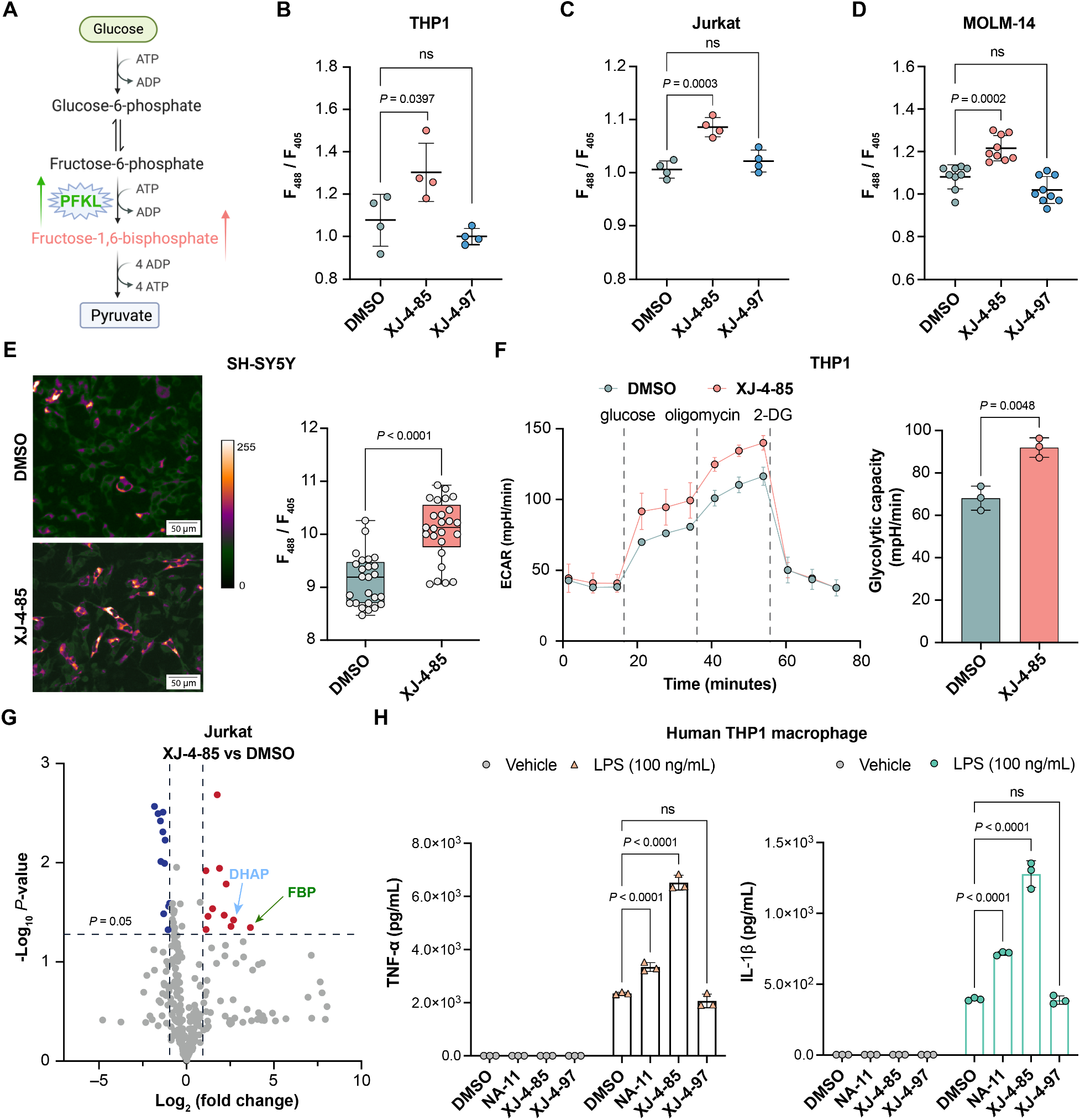
XJ-4-85 treatment induces rapid alterations in glycolytic metabolism and signaling. (A) Schematic of glycolytic pathway. Image was created with BioRender. Quantification of the HYlight sensor ratio for THP1 (B), Jurkat (C) and MOLM-14 (D) cells treated with DMSO, **XJ-4-85** or **XJ-4-97**. (E) Left - a representative image in SH-SY5Y illustrates the sensor’s response to fructose-1,6-bisphosphate (FBP) detection following 2 hours of treatment with either DMSO vehicle or **XJ-4-85**. Right - the fluorescence ratios (488/405 nm) were statistically analyzed across the same fields. Data are presented as mean ± SEM, n = 24. (F) Upregulation of glycolysis in THP1 cells by Seahorse assay. (G) Volcano plot showing metabolomic changes in Jurkat cells treated with **XJ-4-85** compared to DMSO vehicle (2 h). Down- or up-regulated metabolites are shown in blue (log_2_(fold change) < –1) or red (log_2_(fold change) > 1), respectively. Significance was calculated using a two-tailed Student’s *t*-test (n = 4 biologically independent replicates). (H) Enhanced expression of proinflammatory cytokine TNF-α and IL-1β in THP1 M0 cells pretreated with DMSO vehicle, NA-11 or **XJ-4-85** (5 µM, 2 h) followed by 4 h stimulation with LPS (n = 3 biologically independent samples). See Table S3 for details.

An acute treatment (2 h) with **XJ-4-85** resulted in markedly elevated intracellular FBP levels in THP1, Jurkat, MOLM-14 and SH-SY5Y cells relative to vehicle control. No significant changes were observed with the PFKL-inactive **XJ-4-97** control or with a T152E control sensor that cannot bind FBP (Fig. 3, B to D and Table S3). In agreement with flow cytometry measurements, we used live cell imaging and the sensor to observe increased levels of intracellular FBP in **XJ-4-85**-treated SH-SY5Y cells (Fig. 3E).

To determine whether the elevated FBP levels resulted in changes to glycolytic activity, we used the Seahorse extracellular acidification assay and observed increased extracellular acidification rates (ECAR) and a higher glycolytic capacity in **XJ-4-85**- but not **XJ-4-97**-treated cells (Fig. 3F and fig. S10). We performed untargeted metabolomics in cells to corroborate that acute **XJ-4-85** treatment resulted in a significant accumulation of glycolytic intermediates (see Table S4 for details). Specifically, we detected a prominent increase in FBP and dihydroxyacetone phosphate (DHAP) compared to vehicle control (Fig. 3G).

We also investigated the effects of **XJ-4-85** treatment in lipopolysaccharide (LPS) stimulated THP1 macrophage cytokine release, a glycolysis-dependent process (*36*). Following LPS stimulation, **XJ-4-85** significantly potentiated expression of the pro-inflammatory cytokines TNF-α and IL-1β compared to DMSO vehicle **XJ-4-97** or NA-11 treatments (Fig. 3H). This increase in inflammatory cytokine secretion supports that **XJ-4-85**-mediated glycolytic activation can functionally impact signaling responses.

### The covalent PFKL activator XJ-4-85 is cytotoxic to cancer cells

Next, a panel of cancer cell lines was treated with **XJ-4-85** to assess the impact of covalent PFKL activation on tumor cell proliferation. The panel included THP1 (monocytic leukemia), Jurkat (T-cell leukemia), HepG2 (hepatocellular carcinoma), A549 (lung adenocarcinoma), B16-F10-Luc2 (melanoma), MDA-MB-231 (triple-negative breast cancer), MOLM-14 (acute myeloid leukemia), and SH-SY5Y (neuroblastoma) cells. Unexpectedly, we found that treatment with the PFKL activator was broadly cytotoxic across the cancer cell lines tested. The observed cytotoxicity appeared somewhat selective for cancer cells since non-cancerous HEK293T cells appeared less sensitive to **XJ-4-85** treatment (Fig. 4A and Table S5).

**Fig. 4.**
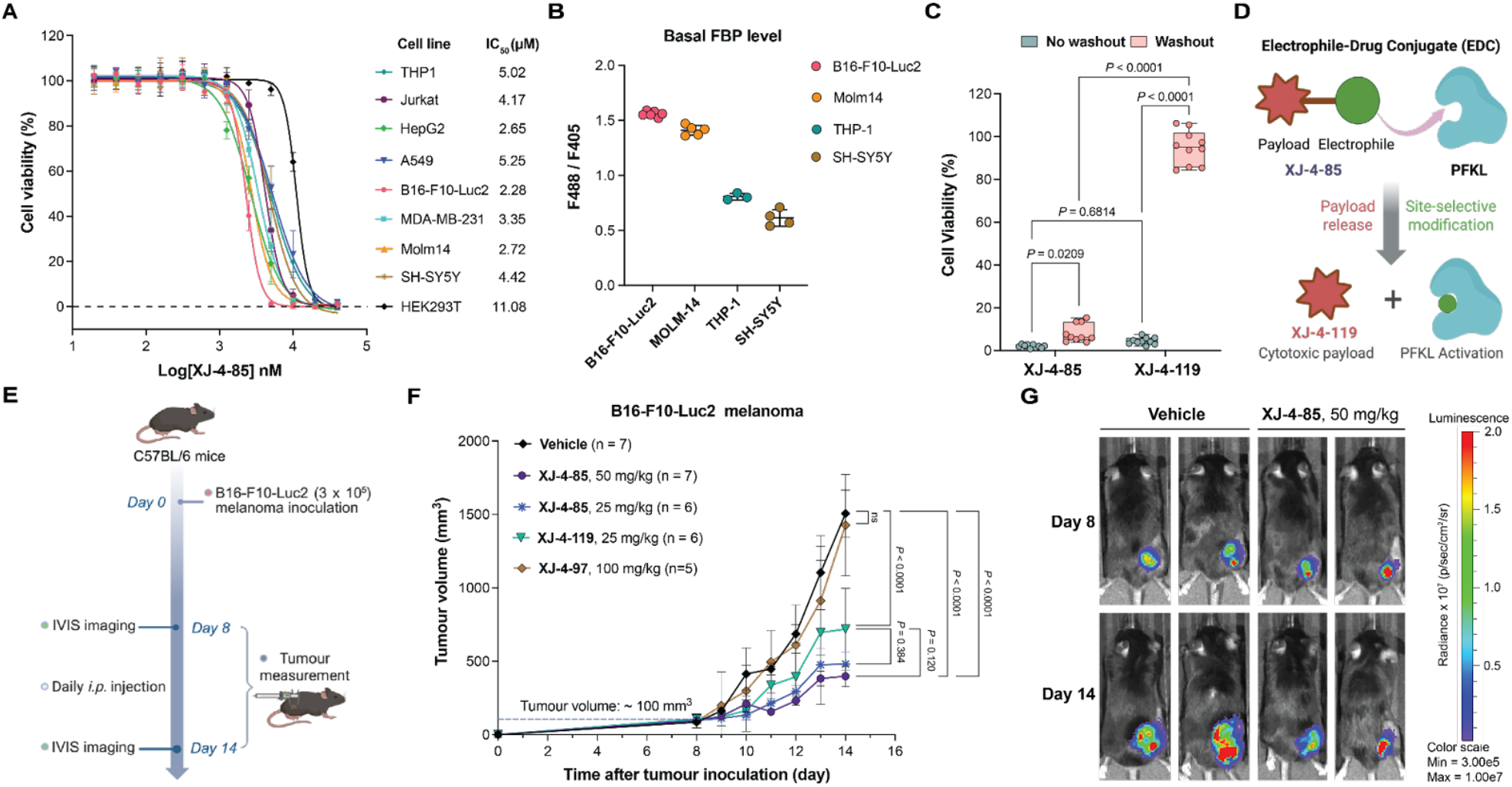
XJ-4-85 blocks tumor growth *in vivo*. (A) Dose-response curves for a panel of cancer cells treated with increasing concentrations of **XJ-4-85** (48 h). The noncancerous HEK293T cell line was included for comparison. Cell viability was performed as described in Supplementary Methods. Data points are normalized relative to vehicle-treated controls for each respective treatment and shown as mean ± SD (n = 4 replicates). (B) Basal FBP levels in B16-F10-Luc2, MOLM-14, THP1, SH-SY5Y cells. (C) Cell viability after 48 h in B10-F10-Luc2 cells pretreated with 10 μM of **XJ-4-85** or **XJ-4-119** for 2 h (n = 10 biological replicates). (D) Schematic of electrophile-drug conjugate (EDC) concept. (E) Treatment paradigm for evaluating compound treatments on B16-F10-Luc2 tumor outgrowth. Scheme was created with BioRender. (F) C57BL/6 mice bearing B16-F10-Luc2 tumors were treated daily with **XJ-4-85, XJ-4-97, XJ-4-119** or vehicle (intraperitoneal injections) at the indicated dose. Tumor measurements were performed daily after tumors became palpable (indicated numbers of mice per group are shown). Data are shown as mean ± SEM. (G) Representative *in vivo* bioluminescence imaging at day 8 and day 14 in B16-F10-Luc2-implanted male mice treated with 50 mg/kg of **XJ-4-85** or vehicle. See Table S5 for details.

We compared basal FBP levels across the cell lines and observed that those with relatively higher basal FBP levels, including B16-F10-Luc2 and MOLM-14, were among the most sensitive to prolonged **XJ-4-85** treatment (IC_50_ values of ∼2.0 µM, Fig. 4, A and B). Washout experiments supported that the effect of extended **XJ-4-85** treatment was mediated through irreversible PFKL binding, consistent with its covalent mode of action. The effects with **XJ-4-119**, which lacks the covalent warhead, were reversible after washout (Fig. 4C).

### XJ-4-85 limited melanoma tumor outgrowth *in vivo*

The unexpected cytotoxic activity of **XJ-4-85** in cancer cell lines prompted further investigations into the long-term effects of metabolic imbalance on tumor outgrowth *in vivo*. The observed selectivity of **XJ-4-85** for PFKL and the cytotoxic activity of its leaving group present an opportunity for its delivery to FBP-high tumor cells. We tested this electrophile-drug conjugate (EDC) concept by directly comparing activity of **XJ-4-85** and **XJ-4-119** in a tumor model *in vivo* (Fig. 4D).

We selected B16-F10-Luc2 melanoma cells for proof-of-concept studies *in vivo* because of enhanced basal FBP levels and sensitivity to **XJ-4-85** (Fig. 4, A and B). We performed a dose-escalation pilot study to identify 10 – 50 mg/kg as the tolerated range for **XJ-4-85** treatments in mice (fig. S11A). B16-F10-Luc2 melanoma cells (3 × 10^5^ cells) were subcutaneously injected into the right flanks of 6- to 8-week-old C57BL/6 mice. Once tumors reached approximately 100 mm^3^, mice received daily intraperitoneal injections of **XJ-4-85** (25 and 50 mg/kg), **XJ-4-119** (leaving group, 25 mg/kg), **XJ-4-97** (negative control, 100 mg/kg) or vehicle control (Fig. 4E).

After two weeks, tumor expansion was evident in the vehicle-treated mice and dramatically and significantly reduced in **XJ-4-85** treated mice (Fig. 4F and Table S5). Even at lower doses (25 mg/kg), **XJ-4-85** treatment significantly controlled B16-F10-Luc2 tumor outgrowth. Mice treated with a high dose of **XJ-4-97** (100 mg/kg) presented with tumor sizes comparable to vehicle-treated cohorts. Notably, administration of an equivalent amount of **XJ-4-119**, representing a maximal exposure of the payload *in vivo*, resulted in reduced efficacy compared to **XJ-4-85** in support of an EDC mechanism. **XJ-4-119** showed tumor-suppressive effects but was less effective at reducing tumor burden than **XJ-4-85** (Fig. 4F). Importantly, none of the treatments at these doses resulted in overt toxicity or weight loss in mice during the 2-week treatment paradigm (fig. S11B). We performed *in vivo* luminescence imaging to further confirm that **XJ-4-85** was efficacious at reducing tumor burden compared to the vehicle group (Fig. 4G and fig. S12 and S13).

Collectively, the covalent PFKL activator **XJ-4-85** blocked tumor growth in animal models with no apparent toxicity between 10 and 50 mg/kg and with evidence for an EDC mechanism.

### XJ-4-85 reprograms cancer cell glycolysis and signaling

To gain insights into **XJ-4-85** mode of action, we performed untargeted LC-MS/MS metabolomic profiling of **XJ-4-85**-treated B16-F10-Luc2 cells (5 μM, 2 h). These studies revealed significant alterations across multiple metabolic pathways. KEGG pathway enrichment analysis of significantly altered metabolites indicated disruptions in starch and sucrose metabolism, pyrimidine metabolism, the pentose phosphate pathway, glutathione metabolism, and glycolysis/gluconeogenesis (Fig. 5A and Table S4).

**Fig. 5.**
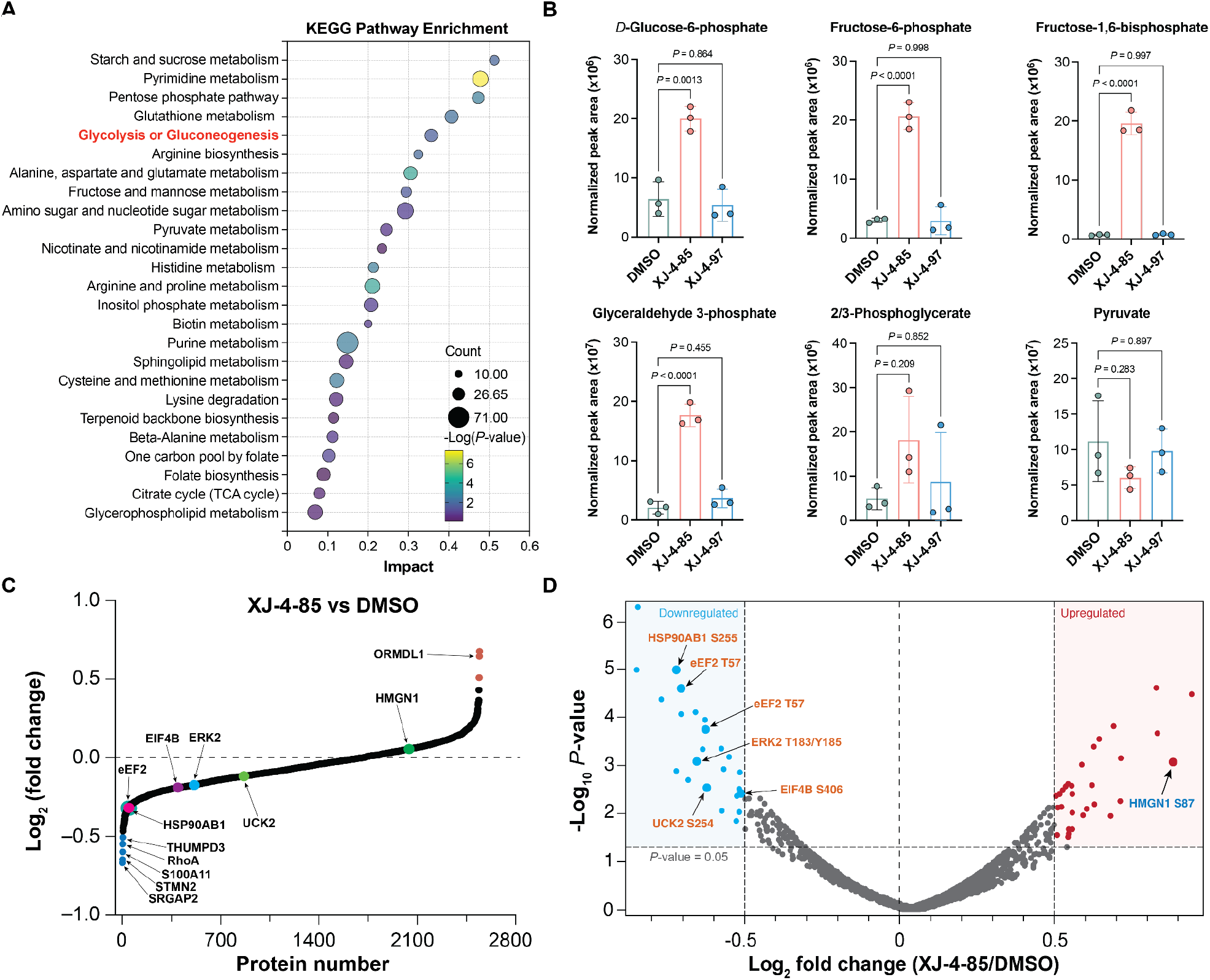
XJ-4-85 destabilizes cancer cell glycolysis and signaling. (A) KEGG pathway analysis of untargeted LC-MS/MS metabolomics profiling showing metabolic pathways significantly enriched following the treatment of **XJ-4-85** (5 µM, 2h) in B16-F10-Luc2 cells. Size of circles denotes the number of metabolites in each respective category. Coloring of circles shows the degree of significance in changes from **XJ-4-85** treatments (enrichment *P* values). (B) Alterations in glycolysis-related metabolites following **XJ-4-85** treatment in B16-F10-Luc2 cells. Data are shown as mean ± SEM, *n* = 3 biological replicates. (C) Waterfall plot showing protein abundance changes in B16-F1-Luc2 cells treated with **XJ-4-85** (5 µM, 2 h) relative to DMSO vehicle, n = 3 biologically independent replicates. (D) Volcano plot showing phosphorylated sites that were significantly up- or down-regulated in B16-F10-Luc2 cells treated with **XJ-4-85** (5 µM, 2 h) relative to DMSO vehicle, n = 3 biologically independent replicates. Phosphorylation sites with a *P* value of ≤ 0.05 and |Log_2_(fold change)| ≥ 0.5 relative to DMSO vehicle treatment are highlighted in color. Associated datasets are provided in Table S4.

Deeper analysis uncovered a distinct glycolytic metabolic profile. **XJ-4-85** treatment led to accumulation of key glycolytic metabolites, including glucose-6-phosphate, fructose-6-phosphate, FBP and glyceraldehyde-3-phosphate (G3P) while pyruvate levels remained relatively unchanged (Fig. 5B). This metabolic profile suggested that PFKL activation by **XJ-4-85** may induce bottlenecks at specific steps of glycolysis and impair glucose utilization.

Next, we deployed TMT-based proteomics and phosphoproteomics to quantify changes in the proteome and phosphoproteome of B16-F10-Luc2 in response to **XJ-4-85** treatments (5 µM). Cells were treated acutely (2 h) to capture earlier changes in cell biology. Comparative analysis revealed that **XJ-4-85** increased expression of ORMDL1, a protein associated with tumor suppression (*37*). Whereas it significantly reduced the abundance of several proteins including SRGAP2, STMN2, S100A11, RhoA and THUMPD3, previously reported to suppress cancer cell proliferation when down-regulated (*38–43*) (Fig. 5, C and D).

Additionally, **XJ-4-85** treatments led to the loss of site-specific phosphorylation for multiple oncogenic signaling proteins including HSP90AB1 (S255), eEF2 (T57), ERK2 (T183/Y185), EIF4B (S406) and UCK2 (S254) (*44–46*) (Fig. 5D). For example, inhibition of ERK2 phosphorylation at T183/Y185 demonstrated anticancer effects against melanoma (*47*). **XJ-4-85** treatment increased phosphorylation of HMGN1 at S87, which has been shown to disrupt DNA repair and chromatin regulation in cancer cells (*48*). Matching proteomics confirmed phosphorylation changes were not due to alterations in protein expression (Fig. 5C and Table S4). Collectively, these data suggest that **XJ-4-85** exerts broader cellular effects beyond metabolic reprogramming, impacting key signaling pathways that regulate cell survival, growth, and stress responses.

## DISCUSSION

Antibody-drug conjugates (ADCs) are an important therapeutic modality in cancer with demonstrated efficacy across a wide range of malignancies (*49*). The rapidly growing interest in ADCs in the clinic stems from the ability to selectively deliver cytotoxic payloads to cancer cells, increasing the therapeutic index (*50, 51*). ADC development, however, faces several significant challenges including limited access to cell surface proteins, high manufacturing costs and complexity, and the potential for immunogenic responses (*49, 52*) . A fully small molecule counterpart to ADCs could overcome many of these limitations including the ability to access intracellular targets. Here, we disclose an electrophile-drug conjugate (EDC) strategy that targets glycolytically enhanced tumor cells using a selective covalent activator of PFKL to induce metabolic stress coupled to delivery of a cytotoxic leaving group.

The development of covalent activators in contrast with inhibitors is less common due to challenges with targeting allosteric regions on proteins (*53*). An initial clue to the distinct activating mechanism of **XJ-4-85** included the striking PFK1 isoform selectivity despite high conservation of the modified lysine site on PFKL. The cryo-EM structure of PFKL bound to **XJ-4-85** revealed covalent and non-covalent interactions in the allosteric activating region engaged by ADP and NA-11 (*34*). These structural studies present two models for explaining the observed selectivity: (i) non-covalent binding of **XJ-4-85** positions the SuTEx electrophile in proximity for K677 modification and/or (ii) covalent adduction at K677 facilitates binding of the departed leaving group in the allosteric pocket (Fig. 2). A direct outcome from the distinct binding mode of **XJ-4-85** is the exquisite proteome-wide selectivity observed in treated cells (Fig. 1). Covalent PFKL activators in contrast with reversible ligands (*34*) provide prolonged activation (Fig. 3H), which can be particularly beneficial in therapeutic contexts where boosting glycolysis is desired (*54–56*).

The high PFK1 isoform- and proteome-wide selectivity of **XJ-4-85** supported its use as a chemical probe to elucidate metabolic and signaling impact of PFKL activation in cancer cells. Single-cell, real-time analyses revealed treatment with **XJ-4-85** confers rapid production of FBP and a concomitant increase in both glycolytic flux and glycolytic metabolites. Importantly, **XJ-4-85** was found to mediate metabolic reprogramming and alter cytokine production and global phosphorylation, revealing a broader role for PFKL in tumor cell biology. Notably, the prominent accumulation of key glycolytic metabolites with **XJ-4-85** treatments indicated metabolic stress that is supported by broad perturbations to the phosphoproteome (Fig. 3 and 5).

Our findings identify **XJ-4-85** as an antitumor agent that exerts its effects through a coupled mechanism involving: (i) metabolic imbalance through PFKL activation, and (ii) release of a leaving group that itself exhibits cytotoxic activity (Fig. 4 and 5). The enhanced antitumor activity of **XJ-4-85** compared with **XJ-4-119** at the same dose provides *in vivo* evidence in support of an EDC mechanism (Fig. 4). Antitumor activity using a PFKL activator, while initially surprising given the Warburg effect, is supported by reports that excessive accumulation of intracellular FBP can disrupt glycolytic flux, induce energy stress, and thus inhibit tumor growth (*13*). Additionally, FBP analogs have been shown to inhibit phosphoglycerate mutase 1 (PGAM1) activation, thereby suppressing cancer cell proliferation (*57*). PFKL is also reported to regulate lipid metabolism and targeting this moonlighting function could affect tumor cell proliferation (*25*).

Future studies include a deeper understanding of how **XJ-4-85** and potentially more potent analogs disrupt the R- to T-state conformation transition and if this allosteric targeting mechanism is more generalizable. PFKL assembles into higher order structures in cells (*22, 58*) and the effects of **XJ-4-85** on these large protein assemblies is not known. The molecular target(s) of **XJ-4-119** mediating the observed cytotoxicity remain to be elucidated. Finally, although we focus on cancer in this report, a PFKL activator has broader applications including reversing the glycolytically deficient state of neurons in neurodegeneration and resensitizing bacteria to antibiotic treatments.

Collectively, our study discloses, to the best of our knowledge, the first covalent, allosteric activator of PFKL that achieves PFK1 isoform and global selectivity. **XJ-4-85** serves as a template for developing EDCs for accessing the cancer intracellular proteome for targeted payload delivery. More broadly, covalent PFKL activation by **XJ-4-85** represents a distinct approach for targeting metabolic vulnerabilities in cancer cells.

## Supporting information

PDB validation report

## ACKNOWLEDGMENTS

We thank H. Hansen and B.A. Webb (West Virginia University) for providing recombinant protein and helpful discussions. We thank T. Huang for help with initial synthesis of compounds. We thank all members of the Hsu and Cambronne labs for helpful discussions and review of the manuscript. We thank R. Goodman (Yale University School of Medicine) for his scientific insights and discussions. We thank T. Mathews and L. Zacharias for metabolomics analysis (Children’s Research Institute Metabolomics Facility), which is supported by the Cancer Prevention Research Institute of Texas (CPRIT Core Facilities Support Award RP240494). This work was supported by the National Institutes of Health grant nos. GM152218 (X.A.C), CA272490 (X.A.C), GM144472 (K.-L.H.), DA043571 (K.-L.H.), AI169412 (K.-L.H.), GM149542 (J.M.K.), GM154453 (L.E.S.), the University of Washington Beckman Cryo-EM Center (S10OD023476 to J.M.K.), the Robbins Family MRA Young Investigator Award from the Melanoma Research Alliance (https://doi.org/10.48050/pc.gr.80540 to K.-L.H.), the Mark Foundation for Cancer Research (Emerging Leader Award to K.-L.H.), a Research Grant Award from The Welch Foundation (F-2143-20230405 to K.-L.H.).and a Recruitment of Rising Stars Award from CPRIT (RR220063 to K.-L.H.).

## Author contributions

K.-L. H., X. A. C. and X.J. conceived and directed the project and wrote the manuscript. X.J., E.M.L., C.L., C.W., L.E.S., S.L. and M.-J.L. performed experimental work and analyzed the data. E.M.L., L.E.S. and J.M.K performed cryo-electron microscopy structural studies. X.J. and C.W. performed animal studies. X.J. performed all proteomic studies. C.L., S.L. and M.-J.L. performed sensor, seahorse and imaging studies. S.L., G.K., H.-R.C., W.W., and Y.-C.L. helped with experiments. The manuscript was edited through contributions of all authors. All authors have given approval to the final version of the manuscript.

## Supplementary Materials

Materials and Methods

Figs. S1 to S16

Tables S1 to S5

References (*59–68*)

**For Table of Contents (TOC) Graphic only**

---

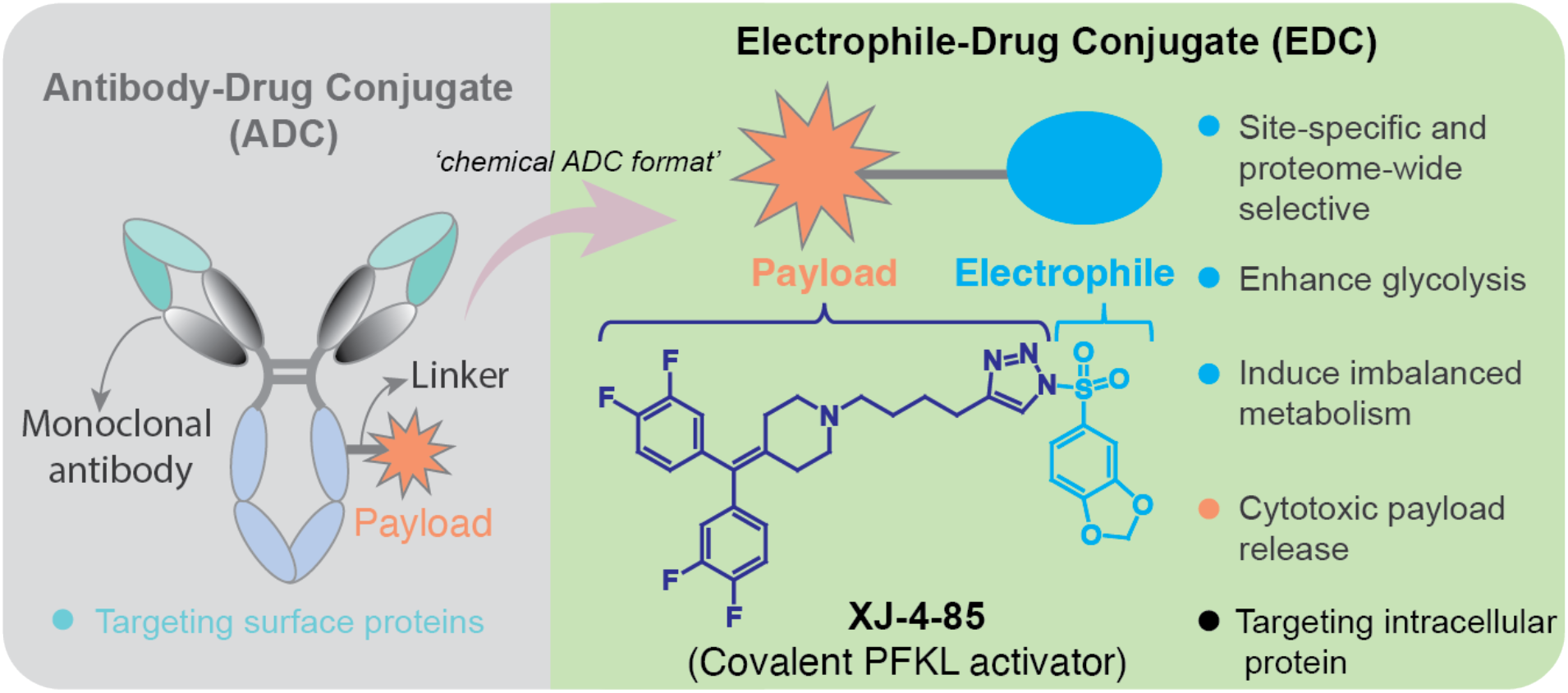

