## Supplementary Information for "A Covalent PFKL Activator Suppresses Tumor Growth"

**CONTENTS**:

1. Supplementary Figures
2. Materials and Methods
3. Chemical Synthesis
4. NMR and HRMS Spectra
5. References

### 1. SUPPLEMENTARY FIGURES

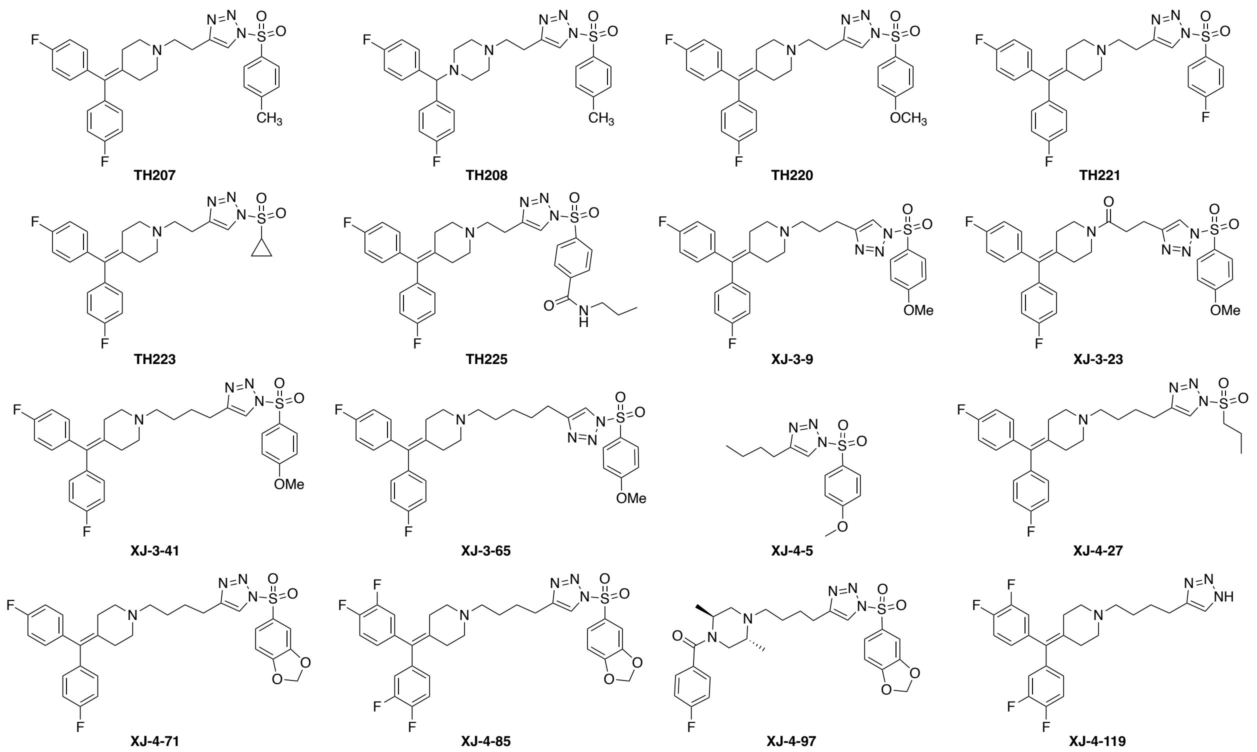


**Fig. S1**. The chemical structures and designated names of SuTEx ligands screened for PFKL activation in this report.

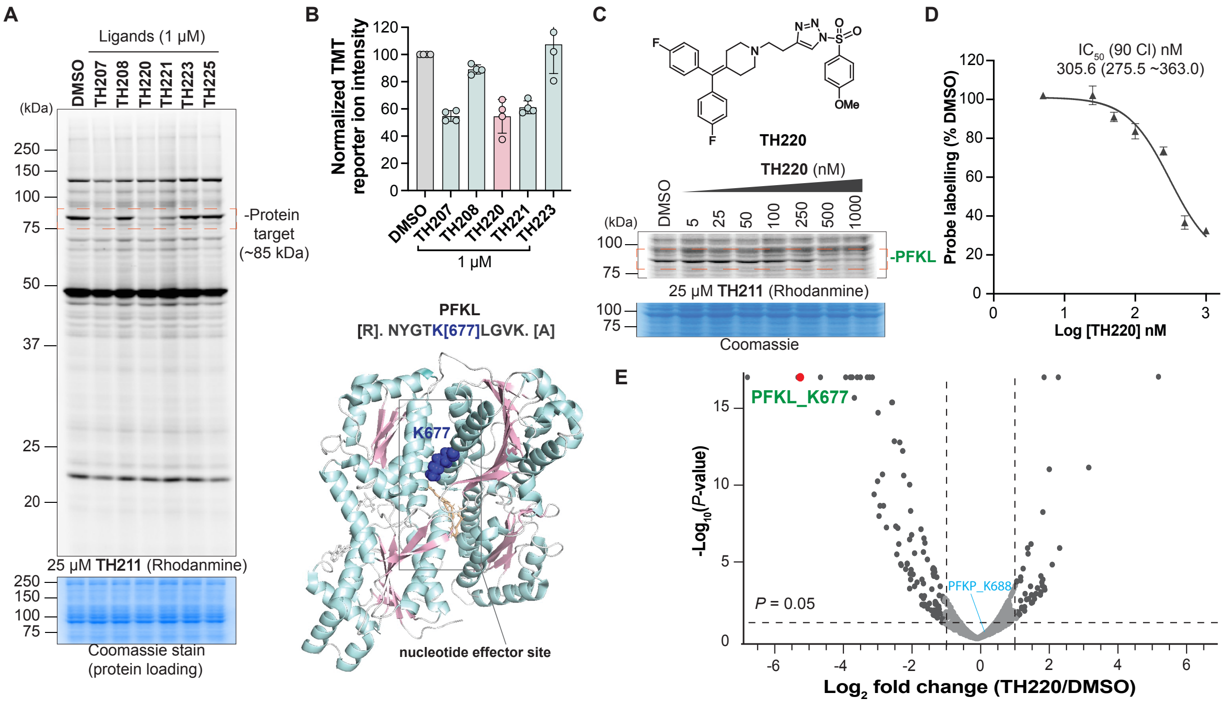


**Fig. S2**. Identification of sulfonyl-triazole ligands that covalently bind a conserved lysine on PFKL in live cells. (A) Gel-based competitive ABPP screening in HEK293T cells using **TH211** probe. HEK293T cells were treated with DMSO vehicle or SuTEx ligands (1 µM) for 1 h followed by treatment with **TH211** probe (25 µM, 2 h). (B) Competitive TMT-ABPP analysis. The normalized TMT reporter ion intensities are shown in bar plot (top), n = 4 biological replicates. K677 is highlighted (blue) in the protein structure of PFKL (bottom). (C) The chemical structure of **TH220** (top) and its dose-dependent binding activity in cells as determined by gel-based competitive ABPP analysis (bottom). HEK293T cells were treated with DMSO vehicle or varying concentrations of **TH220** for 1 h followed by treatment with **TH211** probe (25 µM, 2 h). (D) The integrated band intensities from gel-based studies were quantified in Image Lab. Data shown are mean ± SD, n = 4 biological replicates. (E) Selectivity of **TH220** in HEK293T cells as determined by competitive TMT-ABPP analysis. See Table S1 for detailed proteomics data.

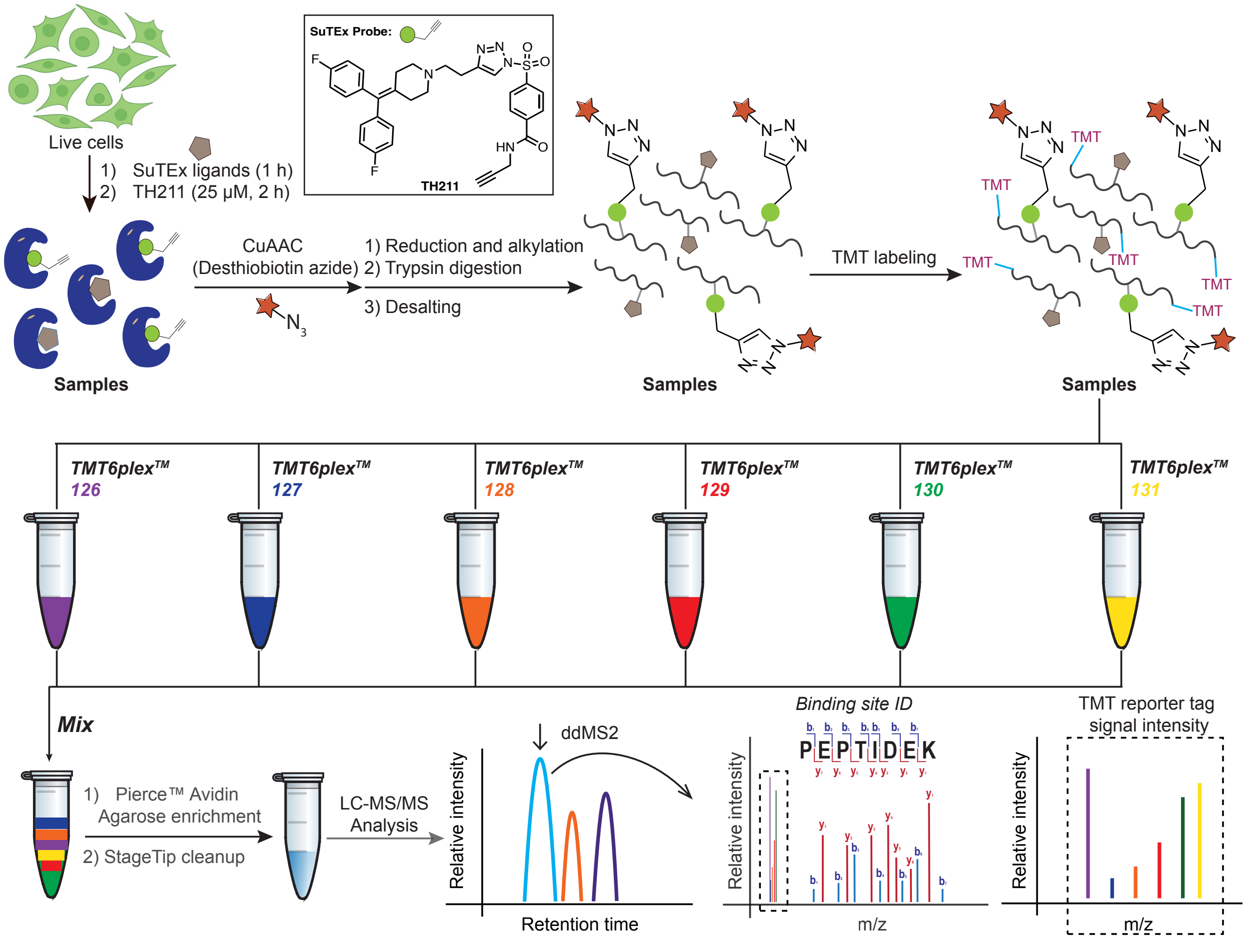


**Fig. S3**. Workflow for competitive TMT-ABPP analysis. Chemical structure of the SuTEx probe TH211 used for chemical proteomics is shown.

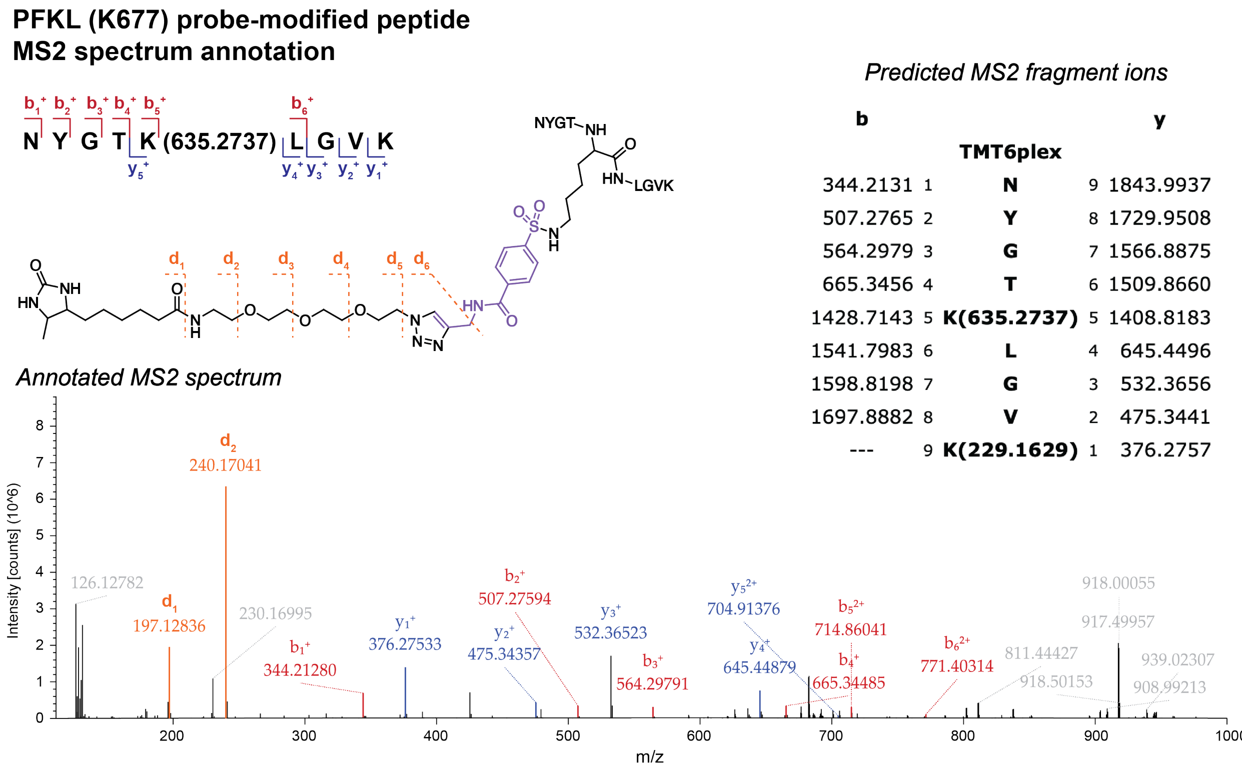


*Predicted MS2 fragment ions*

| b | # | TMT6plex | # | y |
|---|---|---|---|---|
| 344.2131 | 1 | **N** | 9 | 1843.9937 |
| 507.2765 | 2 | **Y** | 8 | 1729.9508 |
| 564.2979 | 3 | **G** | 7 | 1566.8875 |
| 665.3456 | 4 | **T** | 6 | 1509.8660 |
| 1428.7143 | 5 | **K(635.2737)** | 5 | 1408.8183 |
| 1541.7983 | 6 | **L** | 4 | 645.4496 |
| 1598.8198 | 7 | **G** | 3 | 532.3656 |
| 1697.8882 | 8 | **V** | 2 | 475.3441 |
| --- | 9 | **K(229.1629)** | 1 | 376.2757 |

**Fig. S4**. MS2 annotation of the PFKL (K677) TH211-modified tryptic peptide. Left panel: Probe-modified peptide sequence and MS2 fragment ion annotation of NYGTK*LGVK (residues 673-681) peptide from PFKL. Covalent reaction of TH211 with K677 results in a modified lysine (K*) with the addition of +635.2737 Da. Fragmentation of the desthiobiotin-containing tag is also shown. Right panel: predicted MS2 b- and y-fragment ions from CID as determined using Protein Prospector software. Bottom panel: annotation of the MS2 spectrum for the TH211-modified PFKL K677 tryptic peptide including fragment ions containing the probe modified lysine site. Data shown are representative of n = 4 biologically independent experiments.

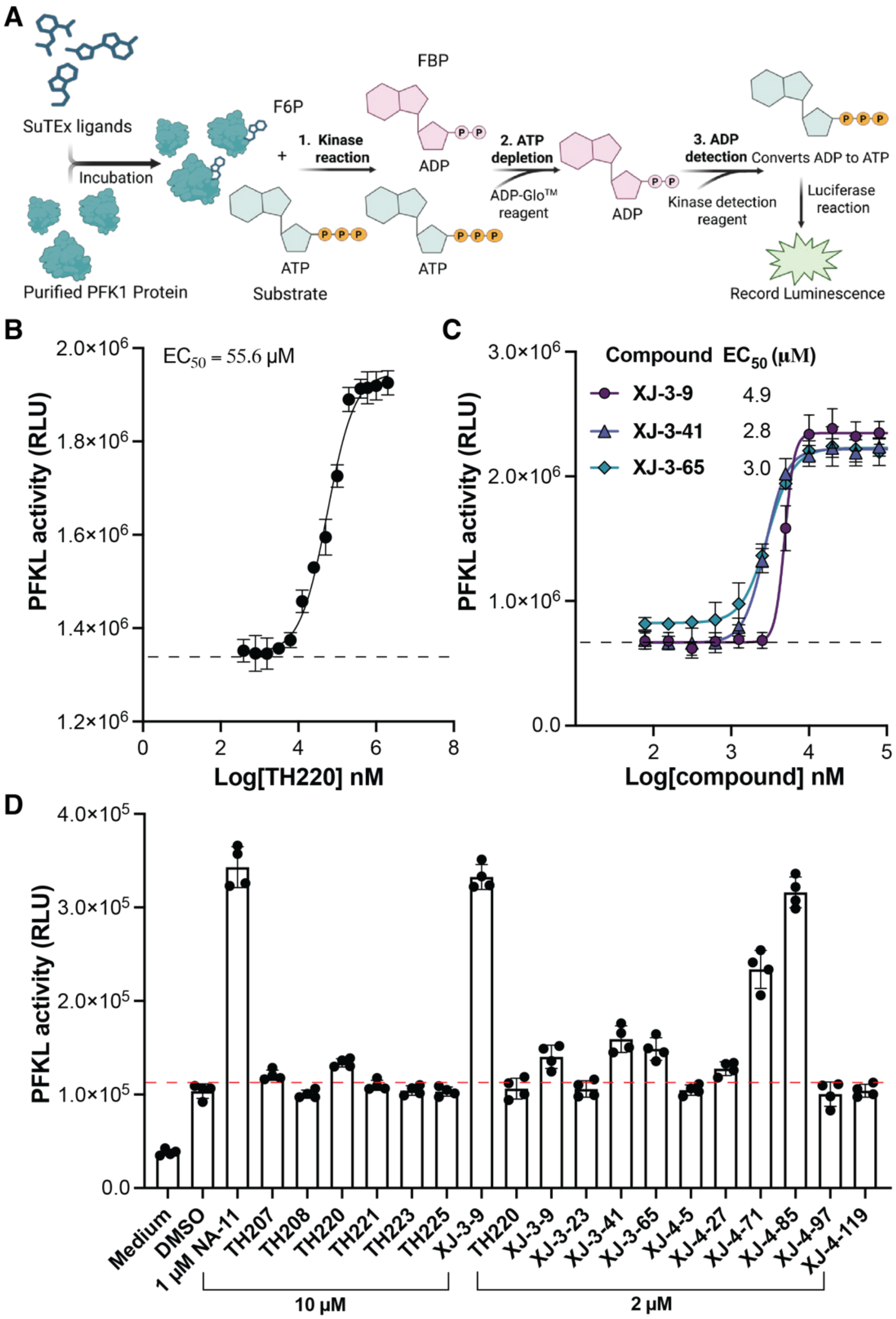


**Fig. S5**. Structure and activity relationship (SAR) study for PFKL activators tested. (A) Schematic of PFKL biochemical assay using ADP-Glo$^{TM}$. (B) Dose-dependent activation of PFKL by **TH220** after incubation for 30 min. (C) Dose-response activation of PFKL by **XJ-3-9**, **XJ-3-41** and **XJ-3-65** after incubation for 30 min. (D) Biochemical screening results for PFKL activation using substrate assay. See Table S2 for details.

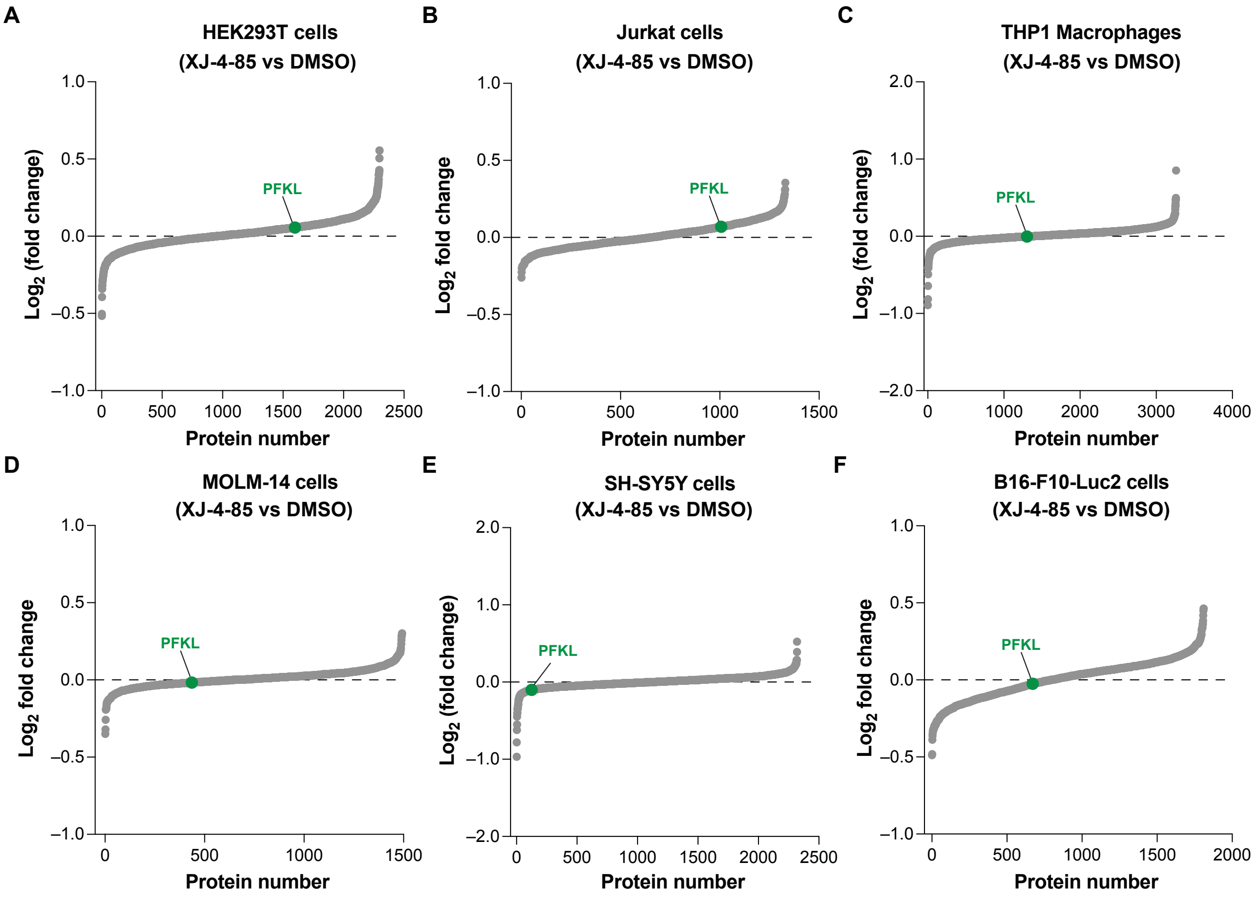


**Fig. S6**. TMT proteomics analysis of protein expression changes in HEK293T, Jurkat, THP1 macrophages, MOLM-14, SH-SY5Y and B16-F10-Luc2 cells treated with DMSO vehicle or **XJ-4-85** (5 µM, 1 h) followed by treatment with **TH211** probe (5 µM, 2 h). PFKL is highlighted in green. See Table S1 for details.

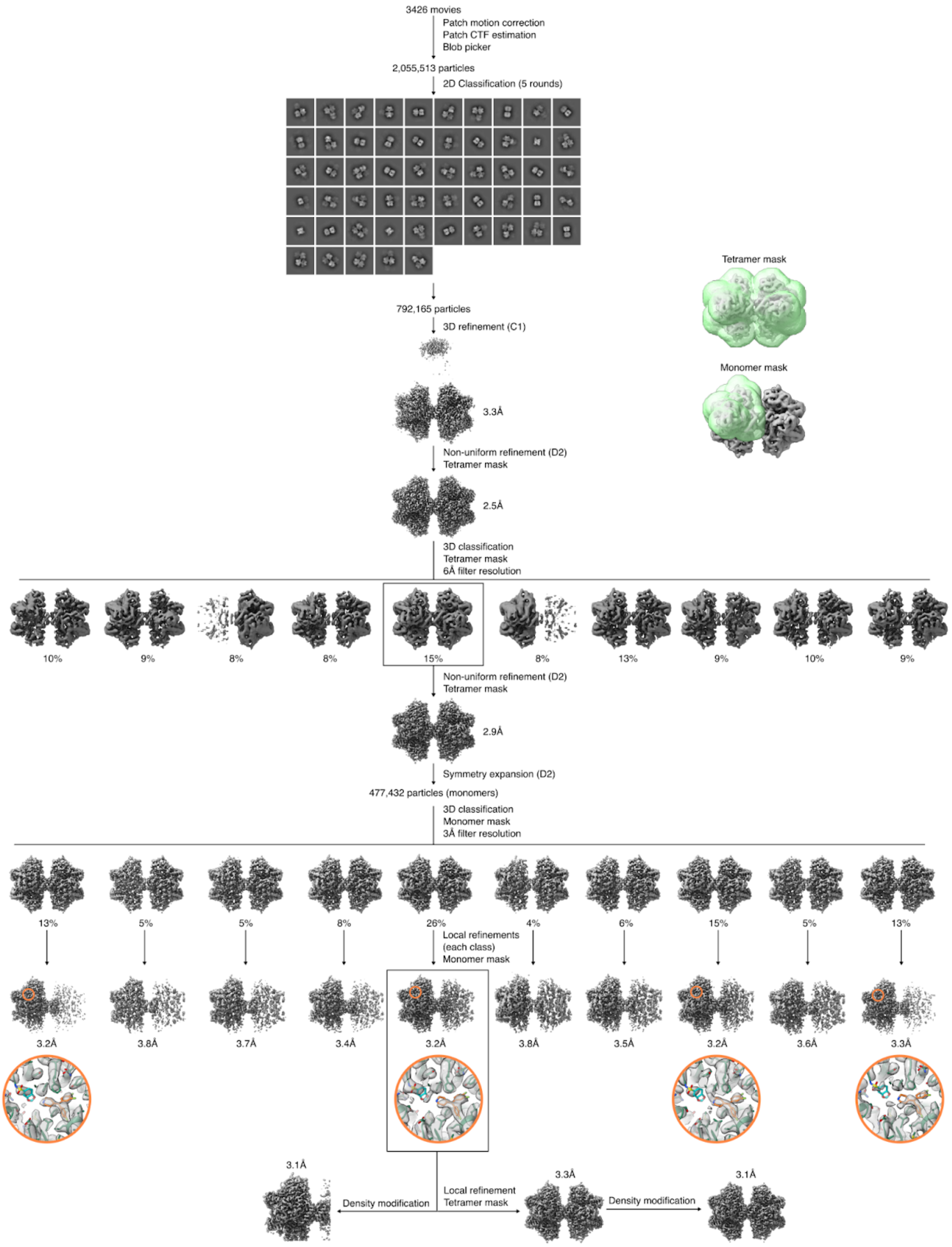


**Fig. S7**. Cryo-EM data processing workflow for PFKL bound to **XJ-4-85**.

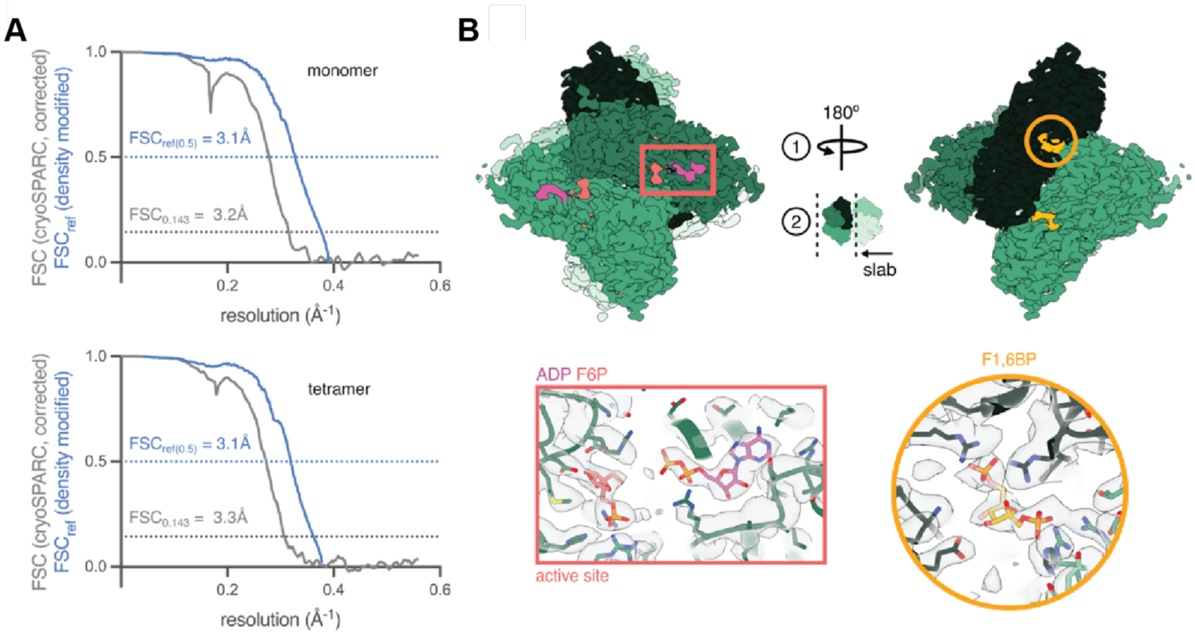


**Fig. S8**. Cryo-EM structure of PFKL bound to **XJ-4-85**. (A) Noise-substituted corrected FSC curves (grey lines) and FSCref curves after density modification (blue lines) and corresponding resolution estimates for the PFKL-**XJ-4-85** monomer and tetramer structures. (B) PFKL-**XJ-4-85** cryo-EM structure, with ADP and F6P highlighted in the active site (pink rectangle) and FBP highlighted in the allosteric sugar-binding site (yellow circle).

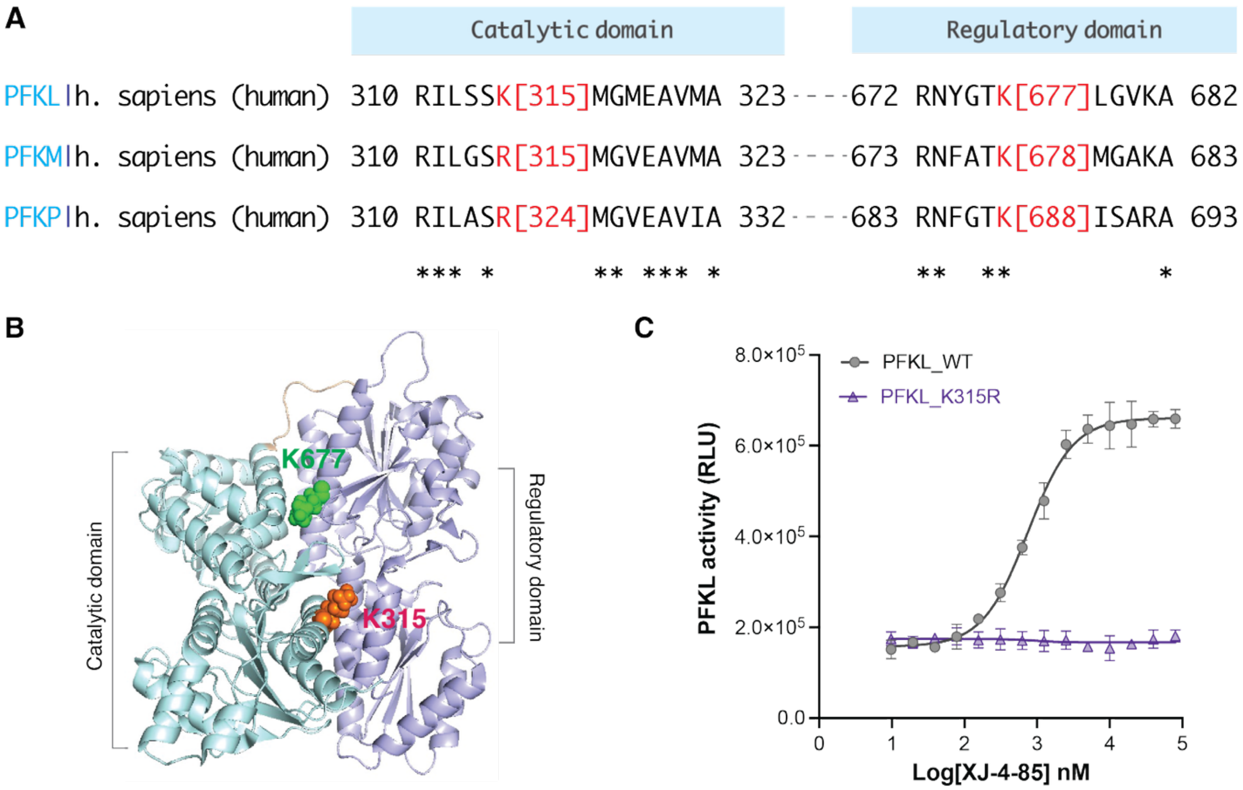


**Fig. S9**. (A) Multiple sequence alignment of PFK1 isoforms. (B) Location of K315 and K677, mapped onto the cryo-EM structure of human PFKL (PDB ID: 7LW1). Catalytic and regulatory domains are colored pale cyan and light purple, respectively. (C) Comparison of biochemical activity of PFKL WT and the K315R mutant after treatment with **XJ-4-85** for 30 min.

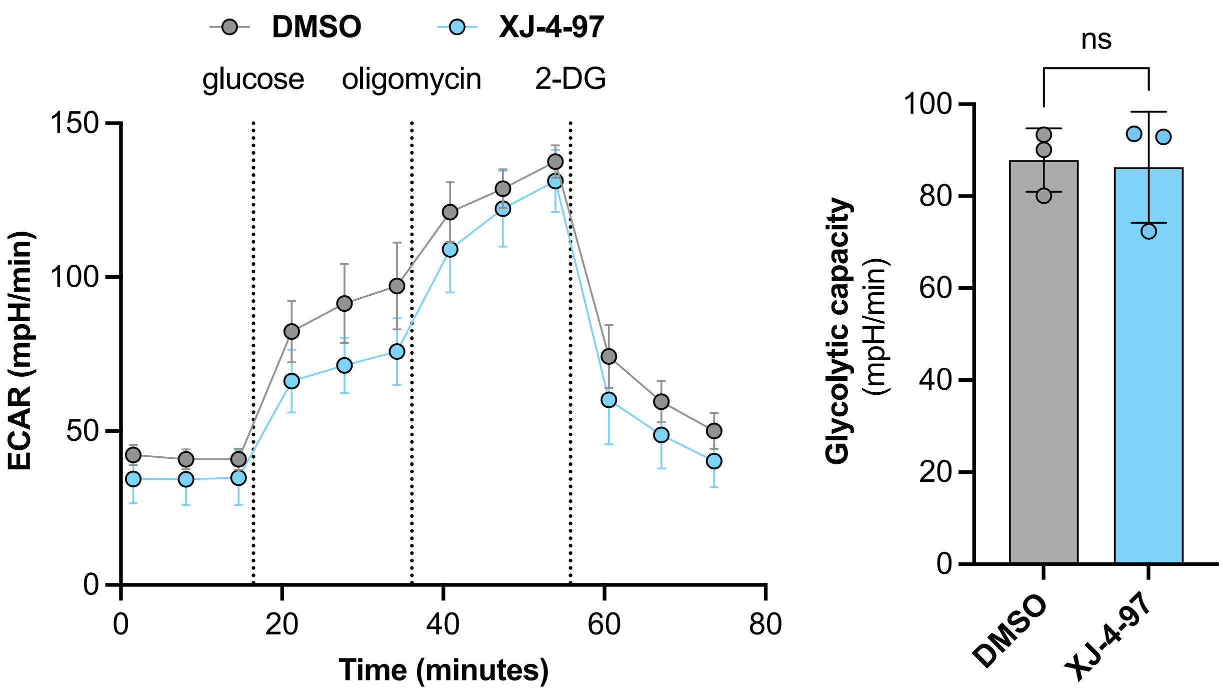


**Fig. S10**. Glycolysis analysis (ECAR) using a Seahorse XF analyzer revealed no difference in glycolytic capacity in THP1 cells treated with **XJ-4-97** (5 μM) compared to DMSO vehicle. See Supplementary Methods for details of assay. Statistical significance was determined by an unpaired Student's *t*-test. Data shown are mean ± SD and representative of n = 3 biologically independent experiments.

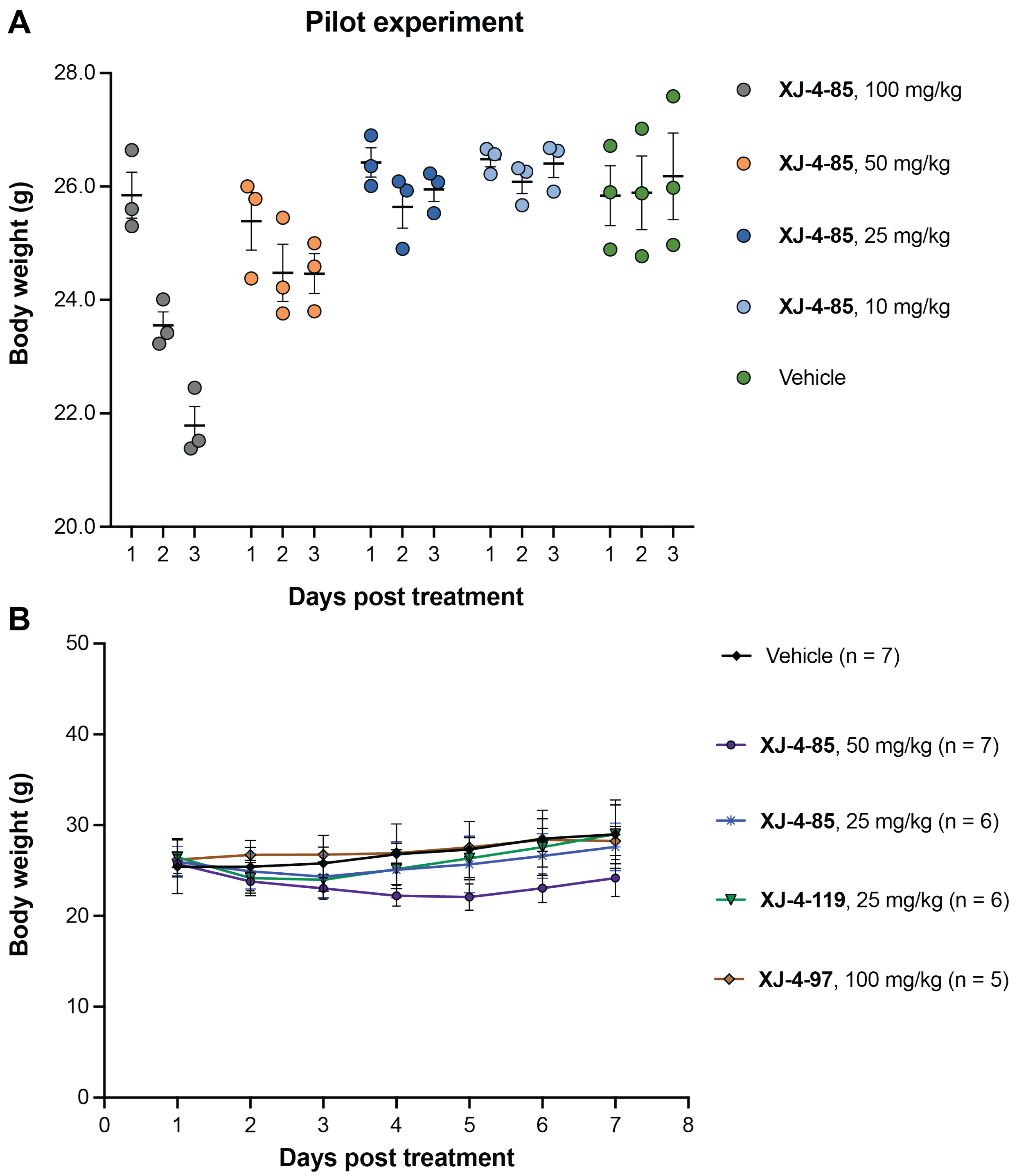


**Fig. S11**. Assessment of body weight changes during *in vivo* compound treatments. (A) Pilot study that identified a tolerated dose range for **XJ-4-85** treatments in mice. (B) Plot showing body weights of mice over time in response to treatment with vehicle, **XJ-4-85**, **XJ-4-119** or **XJ-4-97** (indicated numbers of mice per group are shown). Data are shown as mean ± SEM.

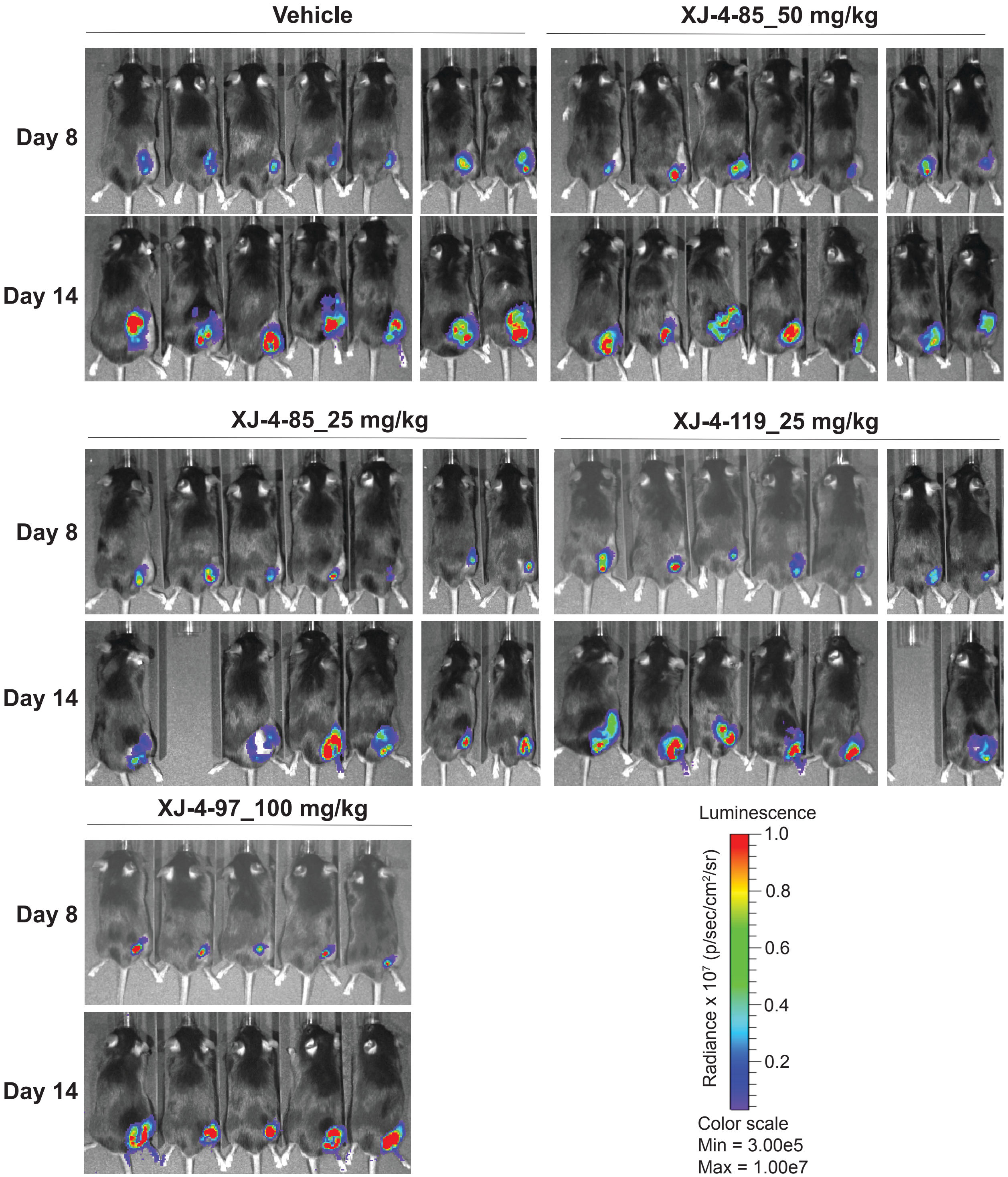


**Fig. S12**. *In vivo* bioluminescence imaging of B16-F10-Luc2-engrafted male mice treated with vehicle, **XJ-4-85**, **XJ-4-119** or **XJ-4-97** (indicated numbers of mice per group are shown).

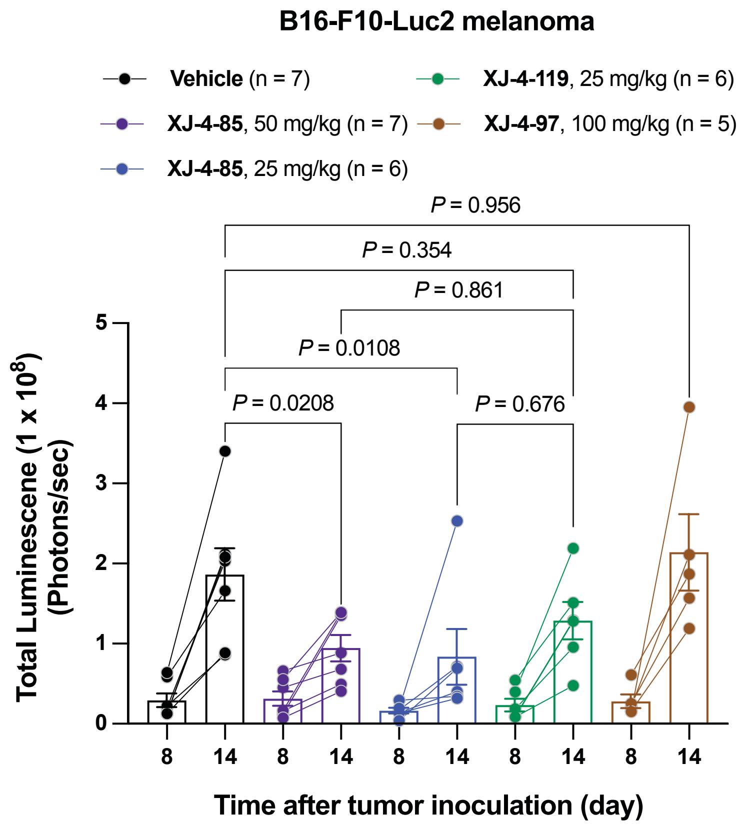


**Fig. S13**. Luminescent intensity of photons emitted from each male mouse in the images from Figure S12 were quantified at day 8 and 14. See Table S5 for details.

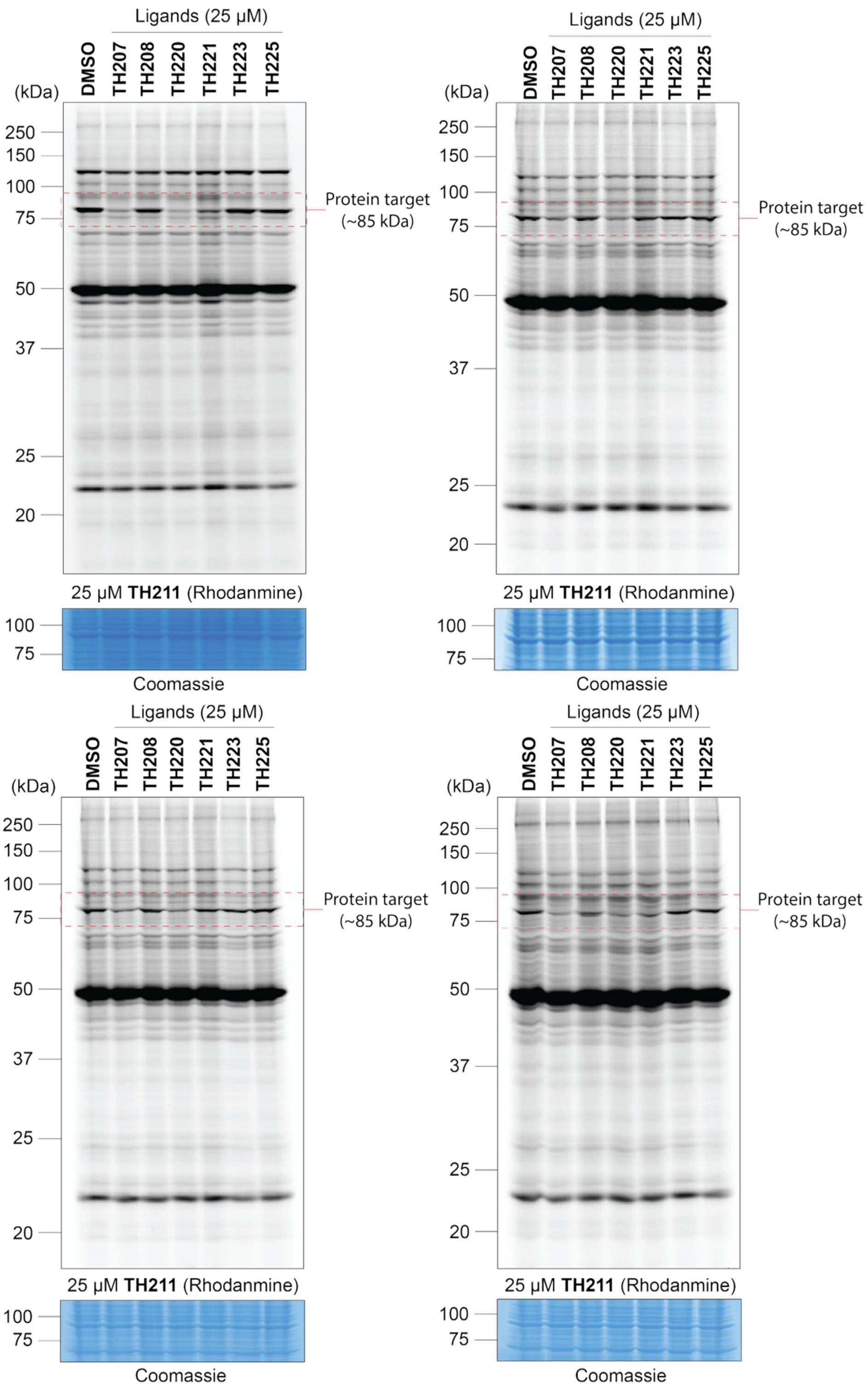


**Fig. S14**. Gel-based competitive ABPP analysis *in situ*. HEK293T cells were treated with DMSO vehicle or the indicated compounds for 1 h, followed by treatment with **TH211** probe (25 μM, 2 h); n = 4 biological replicates. Related to Figure S2A.

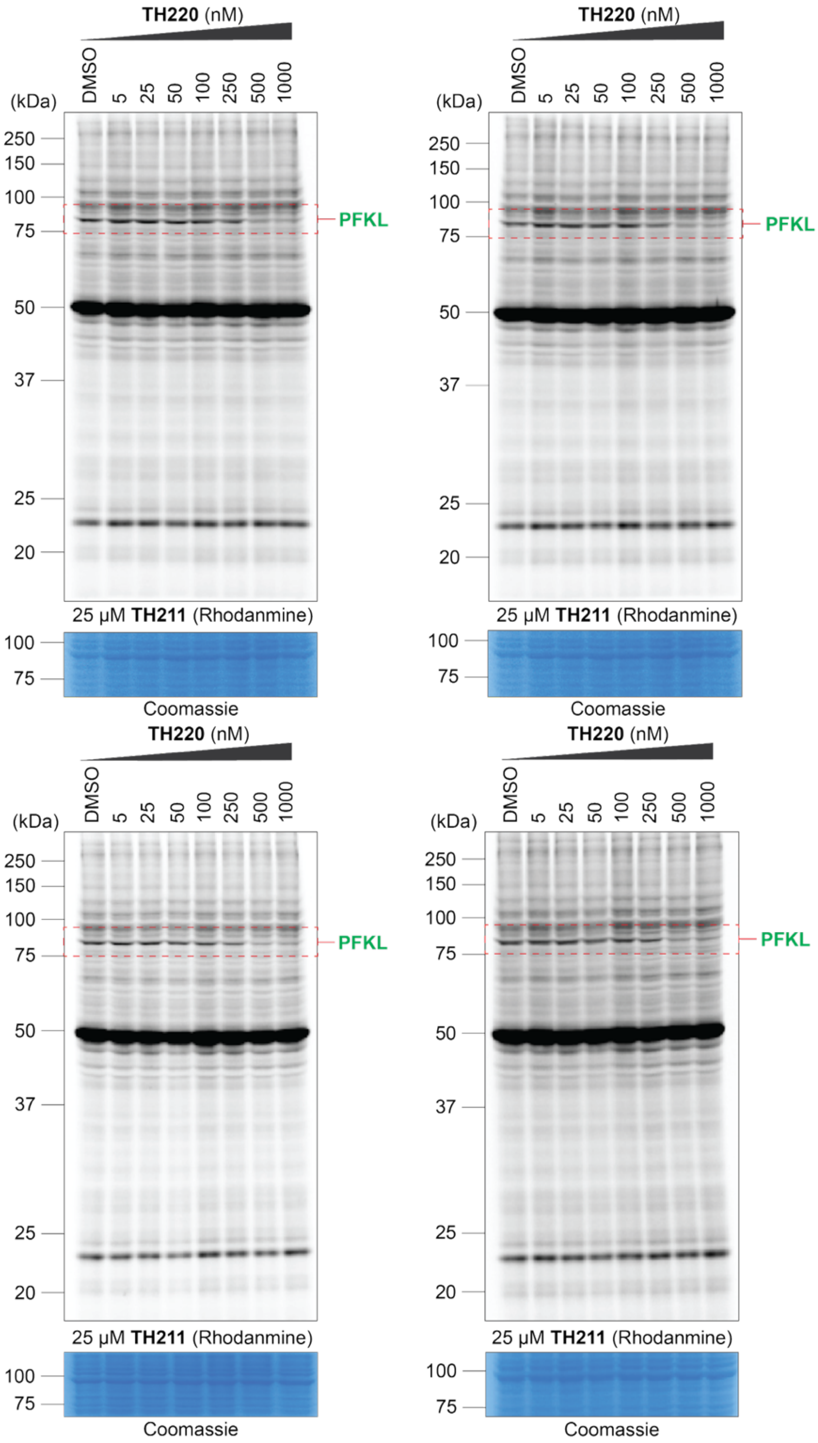


**Fig. S15**. Concentration dependent blockade of probe labeling in HEK293T cells treated with **TH220** as measured by gel-based competitive ABPP. HEK293T cells were treated with DMSO or **TH220** (5-1000 nM, 1 h) followed by treatment with **TH211** probe (25 μM, 2 h); n = 4 biological replicates. Related to Figure S2C and S2D.

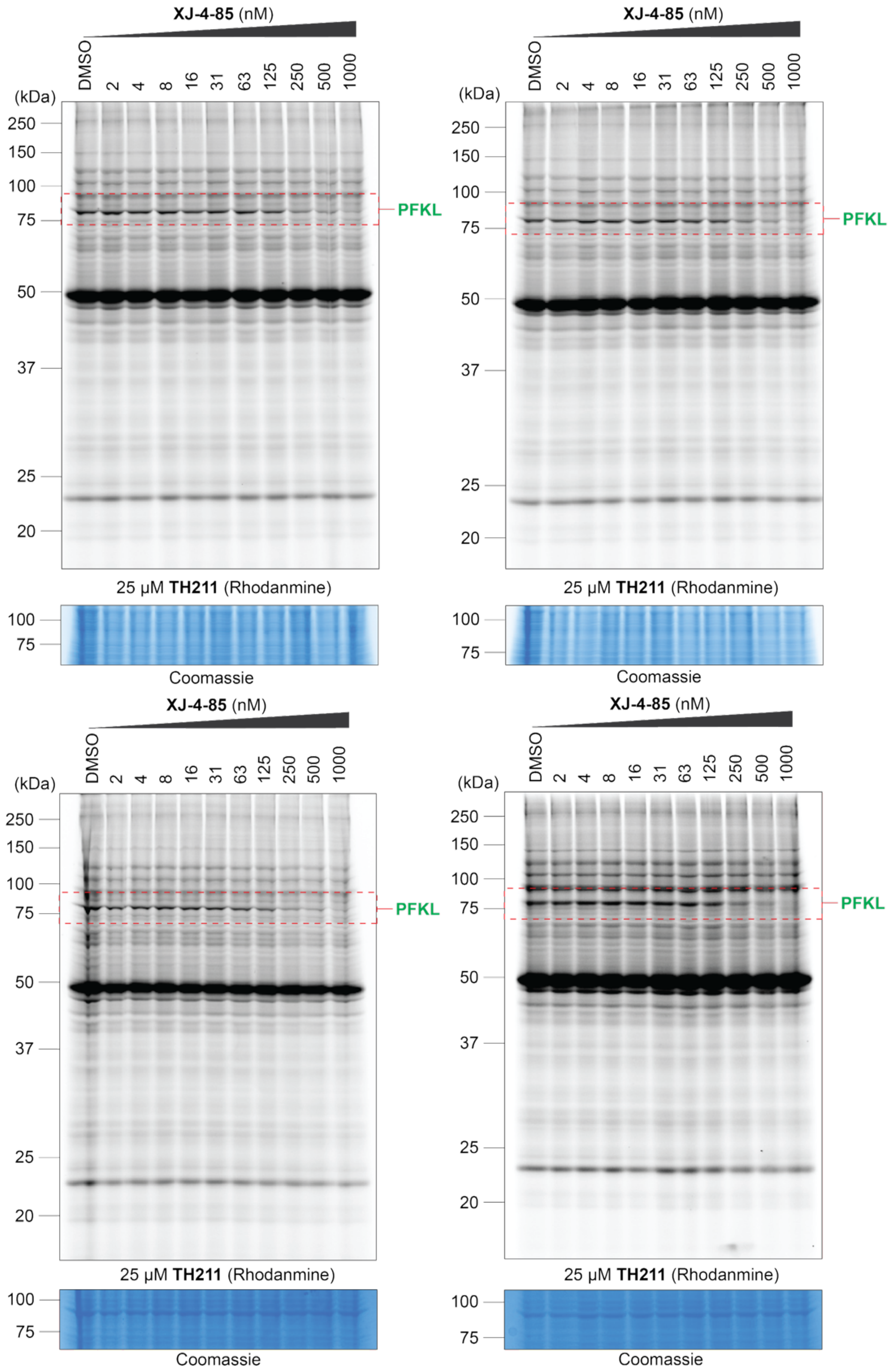


**Fig. S16**. Dose-dependent blockade of probe labeling in **XJ-4-85**-treated HEK293T cells as determined by gel-based competitive ABPP. HEK293T cells were treated with DMSO vehicle or **XJ-4-85** (2-1000 nM, 1 h) followed by treatment with **TH211** probe (25 µM, 2 h); n = 4 biological replicates.  Related to Figure 1C.

#### 2. BIOLOGICAL METHODS

##### Mice

All studies were conducted in 6-12-week-old C57BL/6J from the Jackson Laboratory. Mice were allowed free access to a standard chow diet and water under specific pathogen–free conditions in the animal facility of the Animal Resources Center (ARC) at the University of Texas at Austin. Animal studies were performed in compliance with National Institutes of Health (NIH) guidelines. Studies were approved by the Animal Care and Use Committee (ACUC) of the University of Texas at Austin.

##### Cell culture

All cell lines were cultured at 37 °C in a humidified atmosphere with 5% $CO_2$ and procedures were conducted under aseptic conditions in a biosafety cabinet according to standard operating procedures. HEK293T, HepG2, A549, MDA-MB-231 and B16-F10-Luc2 cells were obtained from the American Type Culture Collection (ATCC) and cultured in complete DMEM media (10% FBS (U.S. Source, Omega Scientific) and 1% L-glutamine (Thermo Fisher Scientific)), penicillin (100 U/mL), streptomycin (100 μg/mL) in 10 $cm^2$ plates. Molm14 cells were cultured in RPMI-1640 medium (Sigma) containing 2 mM L-glutamine, supplemented with 10% fetal bovine serum (Thermo Fisher Scientific) and 1% penicillin-streptomycin (Thermo Fisher Scientific), and 25 mM HEPES. Human neuroblastoma cells SH-SY5Y were maintained in Dulbecco's Modified Eagle Medium/Nutrient Mixture F-12 (DMEM/F12, Sigma) supplemented with 4 mM L-glutamine, 10% fetal bovine serum (FBS, Thermo Fisher Scientific) and 1% penicillin-streptomycin (Thermo Fisher Scientific), and 25 mM HEPES. For starvation experiments, glucose was omitted from the medium to induce a state of nutrient deprivation. Prior to drug treatment, cells were re-supplied with 5 mM glucose (Sigma) to assess cellular responses under re-nourished conditions. The human monocyte cell line THP-1 and Jurkat cells were cultured in RPMI-1640 medium supplemented with 2.05 mM glutamine, 1% antibiotic-antimitotic (penicillin-streptomycin, Gibco), 10% heat-inactivated FBS. To differentiate THP-1 cells into macrophage-like cells, cells were treated with 100 ng/mL PMA in 10% FBS culture medium for 24 h and treated with RPMI-1640 medium containing 5% FBS for another 2 d. To activate M0 macrophages, macrophages were incubated with 5% FBS RPMI-1640 medium containing 100 ng/mL LPS for 4 h. All the cell lines were routinely tested negative for mycoplasma contamination.

***In situ* gel-based competitive activity-based protein profiling**

HEK293T cells were grown to 90% confluence. Cells were carefully washed with Dulbecco's phosphate-buffered saline (DPBS) and replenished with 10 mL of serum-free medium containing the DMSO vehicle or SuTEx ligands at the indicated concentrations, with the final concentration of DMSO at 0.1%. After incubation at 37 °C for 1 h in $CO_2$ incubator, the medium was aspirated and 10 mL of probe TH211 (25 μM) in serum-free medium was added. After incubation at 37 °C for another 2 h, the cells were then harvested in cold DPBS by scraping. After centrifugation at 400 x g for 5 min, the cell pellets were washed with cold DPBS (2 times). Pellets were either directly processed or kept frozen at −80 °C until further use. The cell pellets were resuspended and lysed by sonication (1 sec pulse, 20% amplitude, 3 times) in DPBS in the presence of EDTA-free protease inhibitor cocktail tablet (Pierce). The cell lysates were subject to ultracentrifugation (100,000 x g, 45 min at 4 °C) to yield the cytosolic fraction in the supernatant and the insoluble fraction as a pellet. Protein concentrations were determined by the Bio-Rad DC protein assay. Proteome aliquots (2 mg/mL, 50 μL) were conjugated with fluorophore which was accomplished by copper-catalyzed azide-alkyne cycloaddition (CuAAC) with rhodamine-azide (TAMRA-azide, 1.25 mM, 1 μL, final concentration of 25 μM) in the presence of tris(2- carboxyethyl)phosphine (TCEP, 50 mM fresh in water, 1 μL, final concentration of 1 mM), tris[(1- benzyl-1H-1,2,3-triazol-4-yl)methyl]amine (TBTA, 1.7 mM in 4:1 *t*-butanol/DMSO, 3 μL, final concentration of 100 μM) and $CuSO_4$ (50 mM, 1 μL, final concentration of 1 mM). After incubation at room temperature (RT) for 1 h, reactions were quenched by adding 4X SDS-PAGE loading buffer with beta-mercaptoethanol (17 μL) and samples were resolved by SDS-PAGE and imaged by in-gel fluorescence scanning using a BioRad image system.

**ADP-Glo™ Kinase Assay.**

A 250x stock solution was prepared in 100% DMSO. 2 μL of these dilutions was added to 198 μl assay buffer and mixed well by vortex. A visual inspection of these dilutions was carried out to check for solubility at 1% DMSO. The PFK reaction in the presence of inhibitor was carried out using 10 μL of 1 μg/mL PFKL (final concentration 0.4 μg/ml) which was added to 10 μL of the

1% DMSO compound in a white, non-binding 96-well plate. The protein/compound mixture was incubated for 30 min at RT. The kinase reaction was started by adding a 5 µL of 5x ATP/F6P stock in the assay buffer (final concentrations 0.5 mM F6P and 0.1 mM ATP). Plates were then sealed, centrifuged at 600 x g for 1 min and incubated at RT for 30 min. 5 µL of ADP-Glo™ Reagent was then added to each well, mixed using a multichannel pipette and incubated at RT for 40 min. 10 µL of Kinase Detection Reagent was added and incubated for 50 min at RT. The luminescence was recorded using a CLARIOstar® multi-mode Plate Reader in single-point mode, with an integration time of 750 milliseconds.

**Enzyme-linked immunosorbent assay.**

Supernatants from THP1 macrophages were treated with different reagents for 2 h before stimulation with or without LPS for 4 h (n = 3 biologically independent samples). The supernatants were collected and stored at −80 °C for further measurements. The concentration of cytokines in the extracellular medium was measured using ELISA Duoset kits (R&D Systems) for human TNF-α and IL-1β according to the manufacturer's instructions. Absorbance was measured at 450 nm using a microplate reader. Concentrations were calculated using the corresponding standard curve after accounting for the fourfold dilution of sample in the assay.

**Cell viability assay**

The cell viability assay was performed using the CellTiter-Glo® Luminescent Cell Viability Assay (Promega, Catalog No.: G7572) according to the manufacturer's protocol. Cells were seeded in white-opaque 96-well plates in 100 µL of full growth media at a density of 2,000 ~ 5,000 cells per well and were incubated for 12 h at 37 °C in a humidified 5% $CO_2$ atmosphere. The cells were subsequently treated with the indicated compounds or DMSO (0.1% DMSO final for all wells) in four replicates and incubated at 37 °C in a humidified 5% $CO_2$ atmosphere for 48 h. DMSO without drugs was used as a negative control to normalize the cell viability of drug treated wells. 100 µL of Cell Titer-Glo (Promega) reagent was added to each well. Microplates were incubated with Cell Titer Glo at RT for an additional 10 min to stabilize the luminescence signal. Cell viability was determined by the luminescence measured using the BioTek Synergy H1 microplate reader.

#### Recombinant protein expression and purification

Recombinant His-tagged human PFKL (NP_002617) was expressed and purified as previously described (*59*). Briefly, PFKL cDNA was cloned into the pFastBac HTa vector and baculovirus generated using the Bac-to-Bac Expression system (ThermoFisher Scientific). PFKL was expressed in $2 \times 10^6$ sf9 cells at an MOI of two for 48 h. Cells were pelleted and stored at -80 °C until purification. Cell pellets were lysed with 20 passes of a Dounce homogenizer after resuspension in lysis buffer (20 mM 4-(2-hydroxyethyl)-1-piperazineethanesulfonic acid, pH 7.5; 80 mM potassium phosphate; 1 mM 2-mercapentoethanol; 10% glycerol; 10 mM imidazole) supplemented with EDTA-free Protease Inhibitor Cocktail (Abcam). Cell debris was pelleted by centrifugation and the lysate incubated with HisPur cobalt resin (ThermoFisher Scientific). The resin was washed with 5 bed volumes lysis buffer, 10 bed volumes lysis buffer containing 2 M NaCl, and 5 further bed volumes lysis buffer. PFKL was eluted in lysis buffer containing 100 mM imidazole. Fractions containing protein were pooled, passed over a desalting column equilibrated in freezing buffer (20 mM HEPES pH 7.5, 1 mM DTT, 500 μM ammonium sulfate, 5% glycerol, 1 mM ATP, and 100 μM EDTA) and concentrated using an Amicon Ultracel-30K Centrifugal Filter Unit (MilliporeSigma). Protein was quantified using a Bradford Protein Assay kit (ThermoFisher Scientific) and the integrity and purity of the protein determined by analysis of Coomassie-stained SDS–PAGE gels. Aliquots of protein were frozen in liquid nitrogen and stored at −80 °C. PFKL point mutants N702T and R315K were generated as previously described (*60*,*61*).

#### Cytoplasmic FBP measurement using the FBP biosensor

SH-SY5Y and MOLM-14 cells were transduced with a lentiviral vector expressing a biosensor for fructose-1,6-bisphosphate (FBP). After incubated with 5% $CO_2$ at 37 °C for 48 h, the cells underwent selection with 1 μg/ml puromycin (ThermoFisher Scientific) to enrich populations stably expressing both the sensor and the resistance gene. Ultimately, cells expressing GFP were sorted using a flow cytometer, and populations demonstrating successful sensor expression were considered stable when GFP expression reached 90%.

For SH-SY5Y-sensor cells, seeding occurred at a density of $1\times10^5$ cells per well in a 12-well plate pre-coated with 25 μg/ml poly-L-lysine (Sigma). After 24 h, the medium was switched to glucose-free DMEM to initiate a starvation environment. These cells were then maintained under starvation conditions for an additional 48 h. Subsequently, to evaluate the cellular response to XJ-4-85, these cells were exposed for 2 h to a medium containing 10 µM XJ-4-85, prepared by dissolving in DMSO and further diluting in DMEM supplemented with 5 mM glucose. In the control group, DMSO concentration was kept at 0.5‰ to maintain solvent exposure consistency.

MOLM-14 cells stably expressing the sensor were cultured at a density of $3\times10^5$ cells/mL in T25 flasks under glucose-free RPMI 1640 medium for a 2-d starvation treatment. Post-starvation, $3\times10^5$ cells were transferred to each well of a 12-well plate with a volume of 500 µl per well. An additional 500 µL of medium, containing either XJ-4-85 or XJ-4-97, was added to achieve a final concentration of 10 µM of the compound.

After a 2 h treatment with these compounds, the cells were harvested and analyzed using flow cytometry within 5min on the Novocyte 3000 VYB. Detection was performed using lasers 488-1 (excitation at 488 nm and emission at 530/30 nm) and 405-2 (excitation at 405 nm and emission at 525/50 nm), and gated a stop condition of 10,000 cells with FITC positive. Analysis was done on flowjo generating a derived value (488/405 ratio).

**Confocal imaging of SH-SY5Y cells expressing Hylight sensor**

SH-SY5Y cells, which stably express the FBP sensor, were seeded at a density of $1\times10^4$ cells per well in a 96-well plate and subsequently subjected to 2 d of starvation treatment. Prior to imaging, the cells underwent a 2 h treatment with either 10 µM **XJ-4-85** or 0.5% DMSO. For each group, images were collected from eight distinct wells, and each condition was replicated three different times to ensure biological consistency. Images were captured on an Olympus IXplore SpinSR with a Yokogawa W2 spinning disk and used 100 mW lasers at 405 nm and 488 nm, a 10X Olympus objective, and a Photonics Prime 95B sCMOS camera. Ratiometric measurements were captured between 519 nm excited at 405 nm and 485 nm Method-1 with an excitation wavelength of 495 nm and an emission wavelength of 519 nm, and Method-2 with an excitation wavelength of 405 nm and an emission wavelength of 519 nm. All images were analyzed using ImageJ software, with the average fluorescence intensity of the cells calculated for each frame.

#### Cryo-electron microscopy

##### Protein expression and purification

PFKL was codon optimized for expression in mammalian cells and synthesized commercially (Twist Bioscience). It was subcloned into the pcDNA3.1(+) vector with an N-terminal Twin-Strep tag and TEV cleavage site using Gibson assembly. PFKL was expressed in Expi293F cells (ThermoFisher). Expi293F cells were thawed, maintained, and passaged as described by the manufacturer. To transfect PFKL into cells, Expi293F cells in log phase were seeded in suspension culture at a density of $3 \times 10^6$ live cells/mL. Plasmid DNA (1 mg DNA/mL culture) was diluted in Opti-MEM (ThermoFisher, 35 mL/mL culture). PEI-MAX (Polysciences), at a ratio of 1:3 (w/w) DNA: PEI MAX, was diluted in Opti-MEM (35 mL/mL culture) and mixed with the diluted DNA. The DNA-PEI MAX mixture was incubated for 10 min at RT before it was added dropwise to the cell culture. The cells were maintained in a humid incubator at 37 ºC with 8% $CO_2$ and 120 rpm. After 120 h, the cells were harvested, washed with cold PBS, and stored at -20 ºC.

For purification, the cell pellet was thawed and resuspended in 50 mM HEPES pH 8, 50 mM potassium phosphate, 150 mM KCl, 2 mM DTT, 10% glycerol, 1% triton X and cOmplete EDTA-free protease inhibitor cocktail (Roche). The resuspended cell pellet was incubated on ice for 30 min. The suspension was then clarified by centrifugation at 32500 × g at 4 ºC for 45 min. The clarified supernatant was collected and passed over a pre-equilibrated 1 mL Strep-Tactin XT 4Flow (IBA Life Sciences) gravity column. Once bound, the protein was washed with 5 column volumes (CV) of 50 mM HEPES pH 7.5, 50 mM potassium phosphate, 150 mM KCl, 2 mM DTT, 10% glycerol, 1% triton X. The resin was further washed with 5 CV of 50 mM HEPES pH 8, 50 mM potassium phosphate, 750 mM KCl, 2 mM DTT, 5% glycerol, then 5 CV of 50 mM HEPES pH 8.0, 50 mM potassium phosphate, 150 mM KCl, 2 mM DTT, 5% glycerol. The protein was eluted in 1 mL fractions with 50 mM HEPES pH 8, 50 mM potassium phosphate, 150 mM KCl, 50 mM biotin, 0.5 mM fructose-6-phosphate (F6P), 2 mM DTT, 5% glycerol. The elution fractions were assessed by SDS-PAGE and selected based on purity. Pure fractions were buffer exchanged into 20 mM HEPES pH 8.0, 150 mM KCl, 3% glycerol, 0.5 mM F6P, 2 mM DTT in a 50k MWCO Amicon centrifugal filter (Millipore Sigma). The buffer-exchanged protein was concentrated, filtered, and flash frozen in liquid $N_2$. Protein concentration was determined using a Bradford assay with BSA as the standard. Sample quality was assessed by negative stain electron microscopy and mass photometry.

**Sample preparation, data collection, and image processing**

To prepare samples for cryoEM, 4.2 µM PFKL in 20 mM HEPES pH 8.0, 150 mM KCl, 3% glycerol, 500 µM F6P, and 2 mM DTT was combined with 25 µM XJ-4-85 and incubated for 30 min at RT prior to addition of 2 mM F6P and 1 mM ATP. Sample was applied to glow-discharged ANTcryo M04-Au300-2.0/1.0 grids (SingleParticle) and blotted away four times sequentially, then plunged into liquid ethane using a Vitrobot (Thermofisher). Movies were acquired using a Glacios microscope (Thermofisher) operating at 200kV, equipped with a K3 Direct Detect camera operating in counting mode with a pixel size of 0.885 Å/pixel, 99 frames, and a total dose of 60 electrons/Å$^2$. Data collection was automated with SerielEM. Data processing was performed using cryoSPARC, where movies were aligned, dose-weighted, and summed by patch motion correction, and CTF parameters were estimated using patch CTF. Particles were picked using the blob picker and subjected to 5 rounds of 2D classification. Particles from selected 2D classes were then refined by homogenous 3D refinement (C1 symmetry) followed by 3D classification with a focus mask encompassing a tetramer. Particles selected from high quality 3D classes were then subjected to non-uniform refinement (tetramer mask, D2 symmetry), symmetry expanded (D2), then subjected to 3D classification with a mask encompassing a monomer. Each monomer class was then refined by local refinement (monomer mask). A single monomer class with the best XJ-4-85 ligand density was selected for local refinement with a tetramer mask, to provide a view of the overall PFKL conformation in the XJ-4-85-bound state. Density modification was performed using ResolveCryoEM in Phenix. Atomic models were refined using ISOLDE, with ligands refined in both ISOLDE and Coot. A final round of real space refinement was performed in Phenix, with grid searches, Ramachandran restraints, and rotamer restraints disabled, and with starting model restraints enabled. Custom geometry restraints were used to restrain the sulfone bond of the K677-covalent moiety to approximately tetrahedral geometry. Ligand restraints for the XJ-4-85 leaving group were generated using eLBOW in Phenix. The cryoEM data processing workflow is summarized in Figure S7.

**Seahorse assays**

Seahorse XFp glycolysis stress test kit (Agilent Technologies, 103017-100) was used to measure glycolytic flux (ECAR) according to the manufacture's protocol. Briefly, cells ($2 \times 10^4$ cells per well) were seeded in a Cell-Tak (Corning, 354240) coated XFp miniplates. At 1 h prior to analysis, the medium was replaced with Seahorse XF media (Agilent Technologies, 103681-100) and plates were incubated in a $CO_2$-free incubator at 37 °C. Basal extracellular acidification rate (ECAR) was then analyzed, followed by ECAR measurements after sequential injections of 10 mM glucose, 1 mM oligomycin, and 50 mM 2-deoxyglucose.

**Chemical proteomics**

Cells were grown to 90% confluence for HEK293T, SH-SY5Y and B16-F10-Luc2 or until the cell density reached $2 \times 10^7$ cells/mL for Jurkat, THP1 macrophages and MOLM-14. Cells were treated with SuTEx ligands at the indicated concentrations or DMSO in serum-free media for 1 h, followed by the treatment of 25 µM of TH211 for 2 h (with the final concentration of DMSO maintained at 0.1%). The cells were washed and harvested in cold Dulbecco's phosphate-buffered saline by centrifugation at 400 x g for 5 min. The pellets after DPBS wash for a second time were either directly processed or snap-frozen using liquid nitrogen and stored at −80 °C for further use.

Cell pellets were resuspended and lysed in DPBS with protease inhibitor (10 mL per one tablet ) by sonication at 4 °C and the protein concentrations were determined using the Bio-Rad DC protein assay. Protein concentration was adjusted to 2.3 mg/mL with lysis buffer and probe-modified proteomes (437 µL, 1 mg of proteome) were subjected to CuAAC conjugation to desthiobiotin-PEG3-azide (10 µL of 10 mM stock in DMSO; final concentration of 200 µM) using TCEP (10 µL of fresh 50 mM stock in water, 1 mM final concentration), TBTA ligand (33 µL of a 1.7 mM 4:1 *t*-butanol/DMSO stock, 100 µM final concentration) and $CuSO_4$ (10 µL of 50 mM stock, 1 mM final concentration). Samples were mixed by vortexing and then incubated for 1 h at RT. Excess click reagents were removed by chloroform-methanol extraction as previously described (*63*). Protein pellets were resuspended in 500 µL of 6 M urea:25 mM ammonium bicarbonate and reduced using 10 mM dithiothreitol at 65 °C for 15 min followed by alkylation with 40 mM iodoacetamide at RT for 30 min in the dark. Next, proteins were precipitated with chloroform–methanol, and the protein pellet was resuspended in 500 µL of 25 mM ammonium bicarbonate and then digested overnight with trypsin/Lys-C (7.5 µg in 15 µL of 25 mM ammonium bicarbonate) at 37 °C on a rotator. The resulting peptides were desalted by using Pierce Peptide Desalting Spin Columns. Peptide concentrations were normalized to 5 µg/µL in EPPS (pH 8.5) using the Pierce Quantitative Colorimetric Peptide Assay (Thermo #23275) according to Manufacturer's instruction. 100 µg peptides from each sample were labelled by TMT6plex reagent (Thermo Fisher Scientific) following the SL-TMT protocol (5). 1 µL of each sample were pooled, desalted, and analyzed to check labeling efficiency. After labeling efficiency was verified, the reaction was quenched and acidified with 5% formic acid (pH reduced to ~2 – 3). TMT-labeled peptides were combined, and dried using a SpeedVac. Samples were reconstituted in 550 µL of DPBS and washed streptavidin beads were then added. The SuTEx-modified peptides were enriched for 1 h at RT with rotation and eluted using 150 µL of 50% ACN + 0.1% formic acid (3×). Enriched peptides were dried, desalted using in-house C18 StageTips following standard procedure, and vacuum centrifuged to dryness. The resulting samples were resuspended in 25 µL of 0.1% formic acid and stored at −80 °C until analysis.

Mass spectrometry samples were analyzed on an Exploris 480 or Orbitrap Eclipse Tribrid Mass Spectrometer coupled with a Vanquish Neo UHPLC System (Thermo Fisher Scientific). Peptide separation were achieved on a 75-µm capillary column packed with 20 cm of C18 resin (3 µm, 120 Å; YMC Co., Ltd.) using an 85 min or 120 min gradient of 4 – 50 % mobile phase B with a flow rate of 300 nL/min, where mobile phase A was 0.1% formic acid in water and mobile phase B consisted of 0.1% formic acid in 80% acetonitrile. Eluted peptides were acquired by data-dependent acquisition (DDA) mode. In brief, MS1 spectra were acquired in the scan range of 400 –1600 m/z at an orbitrap resolution of 120,000 with an AGC target set to 1 x $10^6$ and a maximum injection time of 20 ms. The top thirty precursors were then selected for MS/MS analysis. Determined charge states between 2 and 5 were required for sequencing, and an 80 s dynamic exclusion window was used with isotopes excluded. For MS2 scans, HCD collision energy set to 36; normalized AGC target set to 50%; maximum injection time of 75 ms; isolation window of 0.7 Da; resolution set to 15,000 (Turbo); fixed first mass of 110 m/z.

Peptide search was performed in Proteome Discover 3.0 (Thermo Fisher Scientific) against UniProt human protein database or Uniprot mouse protein database with the following parameters: up to two missed cleavages, 10 ppm precursor mass tolerance, 20 ppm fragment mass tolerance and fully tryptic peptides were allowed with a minimum of 6 peptide length. Cysteine residues were searched with a static modification for carboxyamidomethylation (+57.02146 Da). SuTEx probe on tyrosine and lysine (+635.2737), TMT6plex on lysine and peptide N termini (+304.2071 Da) and oxidation of methionine (+15.9949 Da) as variable modifications were included. For the TMT reporter ion quantification, all the identified peptide spectral matches from the MS2 scans were extracted by an in-house program and the reporter ion intensities were adjusted for impurity correction according to the manufacturer's specifications. Peptides used for quantification met the following quality control criteria: PMI-Byonic score ≥ 300, delta ppm err. 5, co-isolation threshold ≤50%, reporter ion S/N threshold ≥ 10. Results were filtered to a peptide false discovery rate of 1%. Volcano plots were generated by grouping PSMs using the peptide isoform node with cutoffs of $Log_2$ fold change ≥ 1.0, and *P*-value ≤ 0.05.

**Phosphoproteomics**

B16-F10-Luc2 cells were seeded in 15 $cm^2$ plates and treated with 5 μM of XJ-4-85 for 2 h upon the confluency reached around 90%. Cells were washed with ice-cold PBS and scraped into 8 M urea lysis buffer (8 M urea, 200 mM EPPS, pH 8.5) with EDTA-free protease/phosphatase inhibitors (Thermo #A32961). Cells were further probe-sonicated on ice. The protein concentrations were determined using the Bio-Rad DC protein assay and adjusted to 2 mg/mL. To 1.0 milligram of proteomes was added 0.5 μL of universal nuclease (Thermo#88700) and incubated at RT for 15 min. The proteomes were then subjected to disulfide bond reduction with 10 mM tris(2-carboxyethyl)phosphine hydrochloride (TCEP) at RT for 30 min followed by alkylation with 20 mM iodoacetamide for 30 min at RT in the dark. Excess iodoacetamide was quenched with dithiothreitol (DTT, 10 mM) at RT for 15 min. Proteins were precipitated by chloroform-methanol precipitation. The resulting protein pellet was resuspended in 500 μL of 25 mM ammonium bicarbonate and then digested with trypsin/Lys-C (7.5 μg in 15 μL of 25 mM ammonium bicarbonate) at 37 °C overnight on a rotator. The resulting peptides were desalted by using Pierce Peptide Desalting Spin Columns. Peptide concentrations were normalized to 5 μg/μL in EPPS (pH 8.5) using the Pierce Quantitative Colorimetric Peptide Assay (Thermo #23275) according to Manufacturer's instruction. 100 μg peptides from each sample were labelled by TMT6plex reagent (Thermo Fisher Scientific) following the SL-TMT protocol (*63*). The labeled peptides were acidified with 10% formic acid (pH reduced to ~2 – 3), pulled together, and dried down in a SpeedVac. TMT-labeled samples were subsequently desalted by using Pierce Peptide Desalting Spin Columns. Ten percent of the resulting peptides were used for protein abundance analysis. The rest were subjected to phosphopeptide enrichment sequentially using the High-Select $TiO_2$ (Thermo Scientific; #A32993) and Fe-NTA (Thermo Scientific; #A32992) Phosphopeptide Enrichment Kits in accordance with the manufacturer's instructions. The resulting phosphopeptides were resuspended with 25 µL 0.1% formic acid before LC-MS/MS analysis.

For the LC-MS/MS analysis, 5 µL of resuspended peptides were separated on an in-house packed 75-µm capillary column with 20 cm of C18 resin (3 µm, 120 Å; YMC Co., Ltd.) using an 140 min gradient of 4 – 45% mobile phase B with a flow rate of 300 nL/min, where mobile phase A was 0.1% formic acid in water and mobile phase B consisted of 0.1% formic acid in 80% ACN. Peptide separations were achieved on an Thermo Scientific™ Orbitrap Eclipse™ Tribrid™ Mass Spectrometer operating in data-dependent acquisition (DDA) mode. For MS1 spectra, resolution for the precursor scan (m/z 400 – 1600) was set to 120,000 with a normalized AGC target set to 250% and a maximum injection time of 50 ms. The 30 most intense ions were selected for MS/MS analysis. Determined charge states between 2 and 5 were required for sequencing, and an 80 s dynamic exclusion window was used with isotopes excluded. For MS2 scans, HCD collision energy set to 36; normalized AGC target set to 50%; maximum injection time of 86 ms; isolation window of 0.7 Da; resolution set to 50,000; fixed first mass of 110 m/z. For data analysis, Thermo Scientific™ Proteome Discoverer™ 3.1 software was used with a precursor mass tolerance of 10 ppm and MS/MS m/z tolerance of 0.02 Da. Carbamidomethylation (+57.021464 Da) on cysteine was used as a fixed modification with methionine oxidation (+15.994915 Da), TMT tags on peptide N terminus/lysine residues (+229.162932 Da) and phosphorylation (+79.966331 Da, T, Y, S) as variable modifications with phosphoRS for site localization. Data was searched against a Uniprot mouse protein database with 1% FDR criteria using Percolator.

**Untargeted Metabolomic analysis**

Metabolites were extracted and measured according to a previously reported protocol (*64*,*65*). Jurkat cells (1 million) were treated with DMSO vehicle, **XJ-4-85** (5 µM) or **XJ-4-97** (5 µM) for 2 h. B10-F10-Luc2 cells in 10 cm plates at ~80% confluence were treated with DMSO, **XJ-4-85** (5 µM) or **XJ-4-97** (5 µM) for 2 h. At the time of collection, cells washed with ice-cold saline solution (Saline: 2F7123 Baxter 0.9% Sodium Chloride Irrigation, USP), lysed with an 80:20 ratio of methanol to water and quickly scraped into an Eppendorf tube, followed by three freeze–thaw cycles between liquid nitrogen and 37 °C. The insoluble material was pelleted in a cooled centrifuge (4 °C), and the metabolite-containing supernatant was transferred to a new tube. The protein concentration of the supernatant was determined, and 10 µg of protein was transferred to a new Eppendorf tube and evaporated until dry using a SpeedVac concentrator (Eppendorf). Metabolites were reconstituted in 100 µL acetonitrile/water 80:20 (vol/vol) for a final concentration of [0.1 mg/mL] of protein, mixed by vortexing rigorously for 1 min and centrifuged at 20,000 x g for 15 min in a refrigerated centrifuge to remove debris. The metabolite-containing supernatant was transferred to an LC-MS vial (with insert) and capped for LC-MS/MS analysis. Chromatographic separation was achieved on a Thermo Scientific (Bremen, Germany) Vanquish Flex liquid chromatography system equipped with a Millipore-Sigma (St. Louis, MO) ZIC-pHILIC column. Metabolite mass spectra were acquired on a Thermo Scientific Orbitrap Exploris 480 mass spectrometer collecting both precursor and product ion spectra. Metabolites were identified with CompoundDiscoverer 3.3 (Thermo Scientific) equipped with an in-house spectral library generated from purified chemical standards, or previously characterized metabolites from biological extracts. Spectra were also searched against the publicly available mzCloud database (Thermo Scientific). Our method compares spectra collect from experimental samples to precursor and product ion spectra from our library or mzCloud. All metabolites were identified with a 5 ppm mass tolerance for precursor ions and a 10 ppm tolerance for product ions. All data were reviewed by an analytical scientist to confirm IDs generated by the software. The normalized areas were used as variables for the multivariate and univariate statistical data analysis. All multivariate analyses and modelling on the normalized data were carried out using MetaboAnalyst v.5.0 (http://www.metaboanalyst.ca). Univariate statistical differences in the metabolites between two groups were analyzed using a two-tailed Student's *t*-test.

**Animal studies**

C57BL/6 mice (male, 18 ± 20 g, 6–8 weeks old) were purchased from Jackson Laboratory. Mice were housed in an animal facility of the University of Texas at Austin under constant environmental conditions (RT, 21 ± 1 °C; relative humidity, 40–70%; a 12 h light–dark cycle). To establish the B16-F10-Luc2 tumor-bearing model, mice were shaved at the injection site, and 200 μL of $3.0 \times 10^5$ B16-F10-Luc2 melanoma cells in serum-free DMEM medium were subcutaneously injected into the flanks of male C57BL/6 mice (8-10 weeks old, average mice weight: 28 g) with a 29 G needle. In all mouse experiments, age-matched and weight-matched mice were randomized to each group to eliminate age and weight differences. All animal experiments were approved by the Animal Resources Center of the University of Texas at Austin. When the tumor volume of B16-F10-Luc2 tumor-bearing mice reached approximately 50 $mm^3$ in size (around one week), the mice were administered intraperitoneally with 100 μL of vehicle (sterile saline : $PEG_{40}$ Castor oil : 100% ethanol solution = 18 : 1 : 1), **XJ-4-85**, **XJ-4-97**, **XJ-4-119** at indicated dose every day. Tumor dimensions were measured daily with a digital caliper, and tumor volume was calculated using the formula (length x $width^2$ x 0.5), where length represents the largest tumor diameter and width represents the perpendicular tumor diameter. For *in vivo* bioluminescent assays, a fresh stock solution of luciferin was prepared at 15 mg/mL in DPBS and sterilized through a 0.2 μm filter. Each mouse was intraperitoneally (i.p.) injected with 100 μL of luciferin solution (15 mg/mL) 10 minutes before imaging. The imaging was then collected on an Xenogen IVIS Spectrum system. During imaging, mice were under isoflurane anesthesia on a bed maintained at physiological temperature. The imaging results were analyzed using Living Image 4.1. Mice weight was recorded daily to monitor potential drug toxicity. Mice were sacrificed when tumors reached 1500 $mm^3$ or upon ulceration/bleeding.

#### 3. CHEMICAL SYNTHESIS

All chemical reagents and solvents were obtained from commercial suppliers such as Sigma Aldrich, Ambeeds, Alfa Aesar, Combi-Blocks, and used as supplied without further purification unless stated otherwise. Analytical thin layer chromatography (TLC) was performed on Merck Silica gel 60 F254 plates (0.25 mm). Flash column chromatography was accomplished with Silica Gel 60 (230- 400 mesh) purchased from Fisher Scientific. Detection was accomplished by using UV-light (254 nm). Nuclear magnetic resonance (NMR) spectra ($^1$H and $^{13}$C) were recorded on a Varian spectrometer at 600 MHz at RT. Chemical shifts were provided in parts per million (ppm) with coupling constants in Hz. $^1$H and $^{13}$C spectra were calibrated in relation to deuterated solvents, namely $CDCl_3$ (7.26 ppm for $^1$H and 77.16 ppm for $^{13}$C) and $CD_3COCD_3$ (2.05 ppm for $^1$H, 29.84 and 206.26 ppm for $^{13}$C). Splitting patterns for apparent multiplets were indicated as s (singlet), d (doublet), t (triplet), q (quartet), m (multiplet), br (broadened) as well as combinations of them. High resolution mass spectrometry was obtained with an Agilent 6545B LC/Q-TOF (Agilent Technologies, Santa Clara, CA, USA). Purity of all final products was greater than 95% as determined by analytical high performance liquid chromatography (HPLC) with a Shimadzu 1100 Series spectrometer detector.

##### General synthetic procedures for preparation of terminal alkynes

**Method A**: The synthetic procedure was performed following previously described methods (*66*). To a solution of secondary amines (1.0 equiv., 10.0 mmol) in DMF (50 mL) was added $K_2CO_3$ (3.0 equiv., 4.15 g, 30.0 mmol) and corresponding *p*-toluenesulfonates (1.2 equiv., 12.0 mmol) sequentially. The reaction mixture was heated to 60 °C and stirred for 6 h. Upon completion, which was indicated by TLC, the reaction solution was concentrated under reduced pressure. And the residue was dissolved with ethyl acetate (200 mL) and washed with brine (50 mL x 3). The combined organic layers were dried over $Na_2SO_4$, filtered and concentrated under reduced pressure. The crude product was purified by silica gel column chromatography to yield alkynes as a yellowish oil.

**Method B**: To a solution of secondary amines (1.0 equiv., 10.0 mmol) in DMF (50 mL) was added $K_2CO_3$ (3.0 equiv., 4.15 g, 30.0 mmol), potassium iodide (2.0 equiv., 3.32 g, 20.0 mmol) and 6-chloro-1-hexyne (1.1 equiv., 1.83 g, 11.0 mmol) sequentially. The reaction mixture was heated to

60 °C and stirred for 6 h. Upon completion, which was indicated by TLC, the reaction solution was concentrated under reduced pressure. And the residue was dissolved with ethyl acetate (200 mL) and washed with brine (50 mL x 3). The combined organic layers were dried over $Na_2SO_4$, filtered and concentrated under reduced pressure. The crude product was purified by silica gel column chromatography to yield alkynes as a yellowish oil.

**Method C**: A solution of 4-pentynoic acid (1.0 equiv., 2.0 g, 20.4 mmol) and 200 µL of DMF (*cat.*) in $CH_2Cl_2$ (20 mL) was cooled (0° C) and oxalyl chloride (1.2 equiv., 2.1 mL, 24.5 mmol) was added dropwise. The reaction mixture was warmed to RT and stirred for 2 h. Excess solvent and oxalyl chloride were removed *in vacuo*, and the dark red residue was redissolved in 20 mL of $CH_2Cl_2$ . This solution was added to a pre-cooled (0° C) solution of RF001 (1.1 equiv., 6.4 g, 22.4 mmol) and DIPEA (2.0 equiv., 7.0 mL, 40.8 mmol). The reaction mixture was warmed to RT and stirred for 2 h. The reaction was poured into a saturated sodium bicarbonate aqueous solution, and the aqueous layer was extracted with $CH_2Cl_2$. The combined organic layers were washed with brine, dried over $Na_2SO_4$ and concentrated under reduced pressure. The crude product was purified by column chromatography to afford the alkynes.

**4-(Bis(4-fluorophenyl)methylene)-1-(pent-4-yn-1-yl)piperidine (XJ-3-7)**

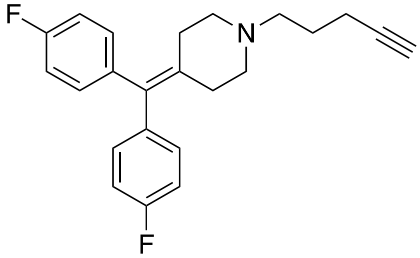


**XJ-3-7**

**Method A**, light yellowish oil, 72% yield, 2.5 g, $R_f$ = 0.20 (hexane/EtOAc=3:1, UV detection on TLC plate). $^1$H NMR (800 MHz, $CDCl_3$) δ 7.09 – 7.02 (m, 4H), 6.99 – 6.94 (m, 4H), 2.56 – 2.43 (m, 6H), 2.37 (t, $J$ = 5.7 Hz, 4H), 2.24 (td, $J$ = 7.1, 2.6 Hz, 2H), 1.94 (t, $J$ = 2.6 Hz, 1H), 1.74 (q, $J$ = 7.4 Hz, 2H). $^{13}$C NMR (201 MHz, $CDCl_3$) δ 161.66, 160.44, 137.72, 137.70, 135.89, 133.31, 130.86, 130.82, 114.54, 114.44, 83.67, 68.02, 56.78, 54.76, 31.15, 25.46, 16.05. $^{19}$F NMR (564 MHz, $CDCl_3$) δ -119.29. HRMS (ESI) calculated for $C_{23}H_{23}F_2N$ [M+H]$^+$: 352.1871, found: 352.1868.

**1-(4-(Bis(4-fluorophenyl)methylene)piperidin-1-yl)pent-4-yn-1-one (XJ-3-19)**

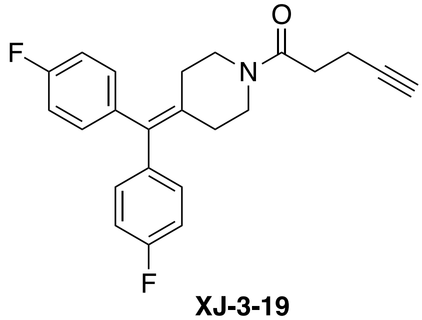

**XJ-3-19**

**Method C**, white solid, 72% yield, 5.9 g, $R_f$ = 0.25 (hexane/EtOAc=3:1, UV detection on TLC plate). $^1$H NMR (800 MHz, $CDCl_3$) δ 7.16 – 7.12 (m, 4H), 7.11 – 7.06 (m, 4H), 3.74 (t, $J$ = 5.9 Hz, 2H), 3.59 – 3.56 (m, 2H), 2.2.71 – 2.66 (m, 2H), 2.65 – 2.63 (m, 2H), 2.48 – 2.46 (m, 2H), 2.43 (t, $J$ = 5.9 Hz, 2H), 2.06 (t, $J$ = 2.6 Hz, 1H). $^{13}$C NMR (201 MHz, $CDCl_3$) δ 168.84, 161.84, 160.61, 137.24, 137.22, 137.05, 137.03, 135.48, 133.61, 130.70, 130.66, 114.79, 114.78, 114.69, 114.67, 83.10, 68.25, 45.95, 42.62, 31.81, 31.53, 30.70, 14.10. $^{19}$F NMR (564 MHz, $CDCl_3$) δ -118.49. HRMS (ESI) calculated for $C_{23}H_{21}F_2NO$ $[M+Na]^+$: 388.1483, found: 388.1489.

**4-(Bis(4-fluorophenyl)methylene)-1-(hept-6-yn-1-yl)piperidine (XJ-3-61)**

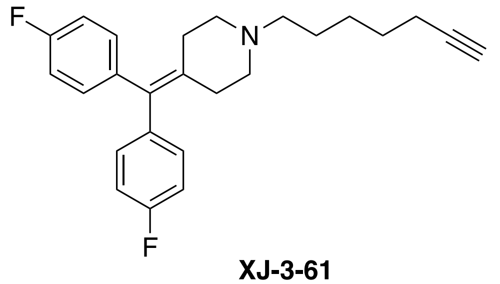

**XJ-3-61**

**Method A**, yellowish oil, 65% yield, 2.5 g, $R_f$ = 0.15 (hexane/EtOAc=4:1, UV detection on TLC plate). $^1$H NMR (800 MHz, Acetone-$d_6$) δ 7.19 – 7.13 (m, 4H), 7.10 – 7.05 (m, 4H), 2.45 (t, $J$ = 5.6 Hz, 4H), 2.34 – 2.29 (m, 7H), 2.17 (td, $J$ = 7.1, 2.7 Hz, 2H), 1.55 – 1.50 (m, 2H), 1.51 – 1.46 (m, 2H), 1.47 – 1.40 (m, 2H). $^{13}$C NMR (200 MHz, Acetone-$d_6$) δ 161.59, 160.38, 138.20, 138.18, 136.63, 132.88, 131.00, 130.96, 114.35, 114.24, 83.59, 68.46, 68.42, 57.46, 54.54, 31.18, 27.90, 26.02, 25.97, 17.36. $^{19}$F NMR (564 MHz, Acetone-$d_6$) δ -117.97. HRMS (ESI) calculated for $C_{25}H_{27}F_2N$ $[M+H]^+$: 380.2184, found: 380.2180.

**4-(Bis(4-fluorophenyl)methylene)-1-(hex-5-yn-1-yl)piperidine (XJ-3-69)**

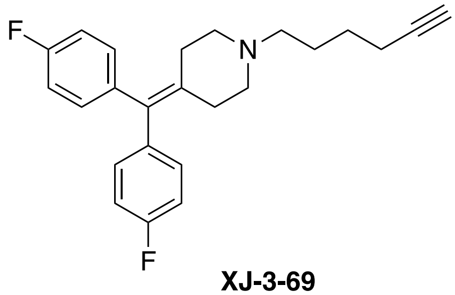

**XJ-3-69**

**Method B**, yellowish oil, 52% yield, 1.9 g, $R_f$ = 0.30 (hexane/EtOAc=3:1, UV detection on TLC plate). $^1$H NMR (800 MHz, Acetone-$d_6$) δ 7.18 – 7.13 (m, 4H), 7.09 – 7.04 (m, 4H), 2.45 (t, $J$ = 5.6 Hz, 4H), 2.35 – 2.28 (m, 7H), 2.20 – 2.17 (m, 2H), 1.60 – 1.57 (m, 2H), 1.56 – 1.52 (m, 2H). $^{13}$C NMR (201 MHz, Acetone-$d_6$) δ 161.59, 160.38, 138.19, 138.17, 136.58, 132.91, 131.00, 130.96, 114.35, 114.25, 83.61, 68.52, 56.90, 54.51, 31.16, 25.87, 25.47, 17.27. $^{19}$F NMR (564 MHz, Acetone-$d_6$) δ -117.97. HRMS (ESI) calculated for $C_{24}H_{25}F_2N$ $[M+H]^+$: 366.2028, found: 366.2045.

**(4-Fluorophenyl)((2*S*,5*R*)-4-(hex-5-yn-1-yl)-2,5-dimethylpiperazin-1-yl)methanone (XJ-4-95)**

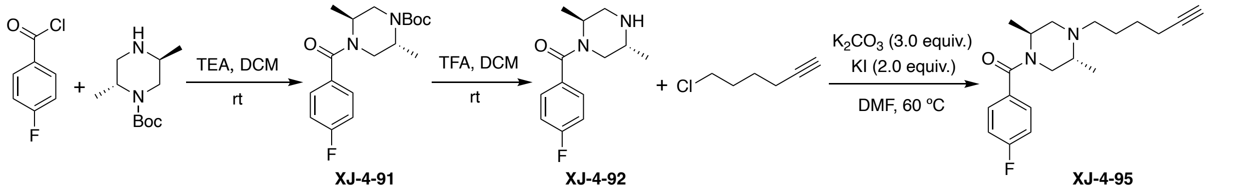


XJ-4-91 XJ-4-92 XJ-4-95

*Synthetic procedures for the intermediates* ***XJ-4-91*** *and* ***XJ-4-92***: To a solution of *tert*-butyl (2*R*,5*S*)-2,5-dimethylpiperazine-1-carboxylate (1.0 equiv., 9.0 g, 42.0 mmol) in dichloromethane (100 mL) was added 4-fluorobenzoyl chloride (1.2 equiv., 8.0 g, 50.4 mmol) and triethylamine (2.0 equiv., 11.7 mL, 84.0 mmol), respectively. The resulting solution was stirred at RT for 1 h. Once completion, the reaction solution was concentrated under reduced pressure. The residue was diluted with ethyl acetate and washed with brine. The organic layers were dried over anhydrous sodium sulfate, filtered and concentrated under reduced pressure. The crude was purified by column chromatography to give **XJ-4-91** (85% yield). To a solution of **XJ-4-91** (12.0 g) in dichloromethane (100 mL) was added 50 mL of trifluoroacetic acid (TFA). The resulting mixture was stirred at RT overnight. Upon completion, the reaction solution was concentrated under reduced pressure. The residue was dissolved with ethyl acetate (200 mL) and washed with sodium hydroxide solution (4 mol/L) and brine, respectively. The organic layers were dried over anhydrous sodium sulfate, filtered and concentrated under reduced pressure. The residue was resuspended in dichloromethane followed by addition of hexane. The white precipitate was filtered and dried to provide **XJ-4-92**.

***tert*-butyl (2R,5S)-4-(4-fluorobenzoyl)-2,5-dimethylpiperazine-1-carboxylate (XJ-4-91)**

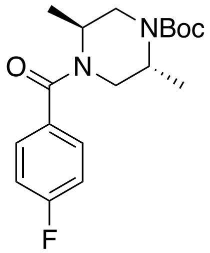

**XJ-4-91**

White powder, 85% yield, 12.0 g, $R_f$ = 0.25 (hexane/EtOAc=5:1, UV detection on TLC plate). $^1$H NMR (500 MHz, Acetone-$d_6$) δ 7.74 – 7.40 (m, 2H), 7.23 (t, $J$ = 8.6 Hz, 2H), 4.91 – 4.40 (m, 1H), 4.27 (s, 1H), 4.02 – 3.45 (m, 2H), 3.39 – 3.09 (m, 2H), 1.46 (s, 9H), 1.34 – 1.02 (m, 6H). $^{13}$C NMR (126 MHz, Acetone-$d_6$) δ 169.50, 163.06 (d, $J$ = 247.0 Hz), 154.64, 154.42, 133.19, 129.47, 129.06, 115.38, 115.21, 79.01, 50.10, 47.29, 46.29, 45.78, 44.53, 43.13, 42.61, 41.80, 41.35, 39.85, 27.70, 14.31. $^{19}$F NMR (471 MHz, Acetone-$d_6$) δ -113.30. HRMS (ESI) calculated for $C_{18}H_{25}FN_2O_3$ [M+H]$^+$: 359.1741, found: 359.1748.

**((2*S*,5*R*)-2,5-dimethylpiperazin-1-yl)(4-fluorophenyl)methanone (XJ-4-92)**

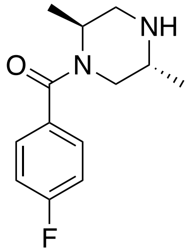

**XJ-4-92**

White powder, 50% yield, 4.2 g, $R_f$ = 0.40 (dichloromethane / methanol = 20 : 1, UV detection on TLC plate). $^1$H NMR (500 MHz, Acetone-$d_6$) δ 7.67 – 7.47 (m, 2H), 7.30 – 7.15 (m, 2H), 4.76 – 4.45 (m, 1H), 3.98 (d, $J$ = 14.6 Hz, 1H), 3.92 – 3.81 (m, 1H), 3.75 (dd, $J$ = 14.5, 3.6 Hz, 1H), 3.59 (dd, $J$ = 13.5, 5.0 Hz, 1H), 3.21 (dd, $J$ = 13.4, 2.0 Hz, 1H), 1.50 (d, $J$ = 7.1 Hz, 3H), 1.47 (d,

$J$ = 6.9 Hz, 3H). $^{13}$C NMR (126 MHz, Acetone-$d_6$) δ 169.85, 163.30 (d, $J$ = 247.4 Hz), 132.21, 129.48, 129.41, 115.52, 115.34, 47.56, 45.33, 41.30, 41.04, 15.03, 12.62. $^{19}$F NMR (471 MHz, Acetone-$d_6$) δ -112.69. HRMS (ESI) calculated for $C_{13}H_{17}FN_2O$ $[M+H]^+$: 237.1398, found: 237.1402.

**(4-Fluorophenyl)((2*S*,5*R*)-4-(hex-5-yn-1-yl)-2,5-dimethylpiperazin-1-yl)methanone (XJ-4-95)**

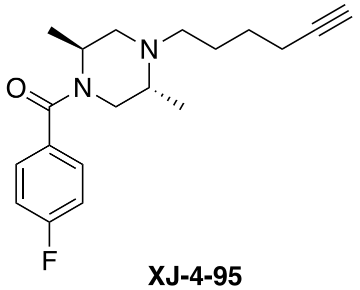


**XJ-4-95**

**Method B**, light yellow oil, 60% yield, 1.9 g, $R_f$ = 0.25 (hexane/EtOAc=3:1, UV detection on TLC plate). $^1$H NMR (500 MHz, Acetone-$d_6$) 7.49 – 7.42 (m, 2H), 7.26 – 7.16 (m, 2H), 4.33 (s, 1H), 3.73 (s, 1H), 3.43 (d, $J$ = 12.9 Hz, 1H), 3.03 (s, 1H), 2.80 (t, $J$ = 1.1 Hz, 1H), 2.72 (dd, $J$ = 11.8, 4.2 Hz, 1H), 2.47 (dt, $J$ = 12.4, 6.9 Hz, 1H), 2.41 – 2.37 (m, 1H), 2.34 (t, $J$ = 2.7 Hz, 1H), 2.25 – 2.19 (m, 1H), 1.68 – 1.50 (m, 4H), 1.34 (d, $J$ = 6.7 Hz, 3H), 0.93 (d, $J$ = 6.6 Hz, 3H). $^{13}$C NMR (200 MHz, Acetone-$d_6$) δ 169.22, 162.89 (d, $J$ = 246.5 Hz),133.74, 133.71, 129.12, 129.05, 115.24, 115.07, 84.14, 68.99, 53.21, 52.36, 48.83, 26.12, 26.05, 17.70, 15.98, 6.32. $^{19}$F NMR (471 MHz, Acetone-$d_6$) δ -113.67. HRMS (ESI) calculated for $C_{19}H_{25}FN_2O$ $[M+Na]^+$: 339.1843, found: 339.1830.

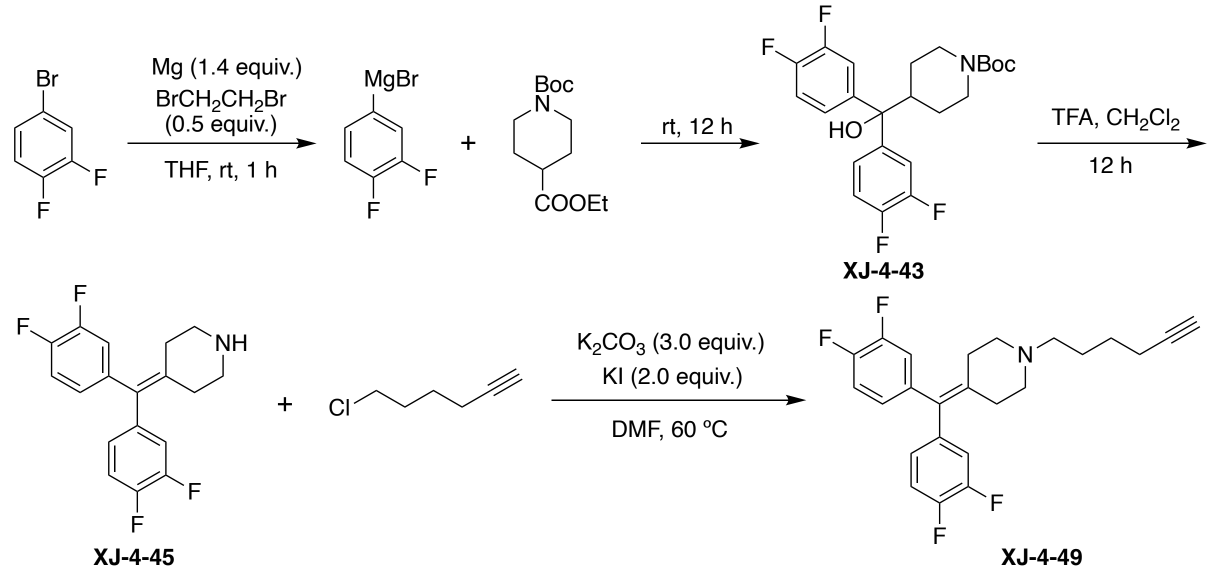


*Synthetic procedures for the intermediates **XJ-4-43** and **XJ-4-45***: The synthetic procedure for Grignard reagent was performed following previously described methods (*67*). A dry flask (500 mL) was equipped with magnesium turnings (3.5 equiv., 2.6 g, 108.8 mmol) and a magnetic stirring bar. The flask was evacuated and heated with a heat gun for 10 min. When the flask was cooled down to RT, it was refilled with argon. THF (200 mL) was then added followed by 1,2-dibromoethane (1.25 equiv., 38.9 mmol, 3.4 mL) and the mixture was heated with a heat gun until ebullition started. After 2 min, the flask was placed into an ice bath and 3,4-difuoro-1-bromobenzene (2.5 equiv., 15.0 g, 77.7 mmol) was added dropwise over 1 h by syringe. 1-Boc-isonipecotic acid ester (1.0 equiv., 8.0 g, 31.1 mmol) was added and the reaction solution was stirred overnight at RT. The reaction was quenched with saturated ammonia chloride solution and extracted with ethyl acetate. The organic layers were combined, dried over anhydrous sodium sulfate, filtered and concentrated under reduced pressure. The residue was resuspended in dichloromethane followed by addition of hexane. The resuspension was cooled down in ice bath and the white precipitate was filtered and dried to provide **XJ-4-43**.

To a solution of **XJ-4-43** in dichloromethane (100 mL) was added 50 mL of trifluoroacetic acid (TFA). The resulting mixture was stirred overnight at RT. Upon completion, the reaction solution was concentrated under reduced pressure. The residue was dissolved with ethyl acetate (200 mL) and washed with sodium hydroxide solution (4 mol/L) and brine, respectively. The organic layers were dried over anhydrous sodium sulfate, filtered and concentrated under reduced pressure. The residue was resuspended in dichloromethane followed by addition of hexane. The white precipitate was filtered and dried to provide **XJ-4-45**.

***tert*-Butyl 4-(bis(3,4-difluorophenyl)(hydroxy)methyl)piperidine-1-carboxylate (XJ-4-43)**

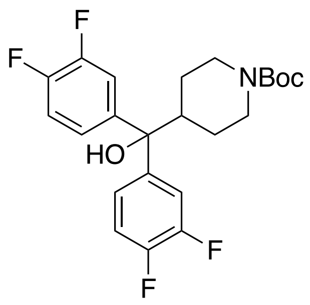

**XJ-4-43**

White powder, 42% yield, 5.7 g, $R_f$ = 0.30 (hexane/EtOAc=4:1, UV detection on TLC plate). $^{1}$H NMR (500 MHz, $CDCl_3$) δ 7.35 – 7.22 (m, 2H), 7.04 – 7.17 (m, 4H), 4.27 – 4.00 (m, 2H), 2.69 (t, *J* = 12.9 Hz, 2H), 2.31 – 2.48 (m, 1H), 2.24 (s, 1H), 1.43 (s, 9H), 1.16 – 1.35 (m, 2H). $^{13}$C NMR (126 MHz, $CDCl_3$) δ 154.61, 151.17 (d, *J* = 12.5 Hz), 150.11 (d, *J* = 12.7 Hz), 149.19 (d, *J* = 12.7 Hz), 148.13 (d, *J* = 12.6 Hz), 142.39 (t, *J* = 4.2 Hz), 121.61 (dd, *J* = 6.1, 3.5 Hz), 117.15, 117.02, 115.32, 115.17, 79.69, 78.51, 44.34, 28.36, 26.24. $^{19}$F NMR (471 MHz, $CDCl_3$) δ -139.96, -143.28. HRMS (ESI) calculated for $C_{23}H_{25}F_4NO_3$ $[M+Na]^+$: 426.1663, found: 462.1676.

**4-(Bis(3,4-difluorophenyl)methylene)piperidine (XJ-4-45)**

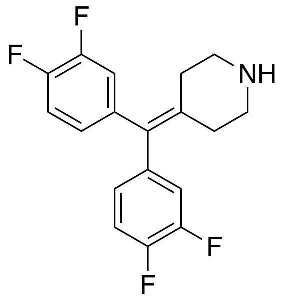

**XJ-4-45**

$^{1}$H NMR (500 MHz, Acetone-*d*$_6$) δ 7.38 – 7.28 (m, 2H), 7.27 – 7.20 (m, 2H), 7.14 – 7.06 (m, 2H), 3.35 (t, *J* = 6.0 Hz, 4H), 2.69 (t, *J* = 6.0 Hz, 4H). $^{13}$C NMR (126 MHz, Methanol-*d*$_4$) δ 151.00 (d, *J* = 12.8 Hz), 150.48 (d, *J* = 12.7 Hz), 149.02 (d, *J* = 13.0 Hz), 148.51 (d, *J* = 12.7 Hz),137.69, 135.71, 131.44, 125.75, 125.72, 125.69, 125.67, 118.01, 117.87, 117.23, 117.09, 44.53, 27.75. $^{19}$F

NMR (471 MHz, Methanol-$d_4$) δ -139.51 (d, $J$ = 20.8 Hz), -141.57 (d, $J$ = 20.8 Hz). HRMS (ESI) calculated for $C_{18}H_{15}F_4N$ $[M+Na]^+$: 322.1213, found: 322.1219.

**4-(Bis(3,4-difluorophenyl)methylene)-1-(hex-5-yn-1-yl)piperidine (XJ-4-49)**

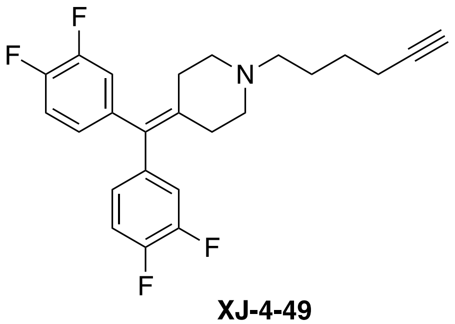

**XJ-4-49**

**Method B**, colorless oil, 65% yield, 2.0 g, $R_f$ = 0.30 (hexane/acetone=4:1, UV detection on TLC plate) . $^1$H NMR (500 MHz, Acetone-$d_6$) δ 7.30 (dt, $J$ = 10.8, 8.5 Hz, 2H), 7.13 (ddd, $J$ = 11.5, 7.8, 2.1 Hz, 2H), 7.06 – 6.99 (m, 2H), 2.82 (d, $J$ = 16.9 Hz, 1H), 2.48 (t, $J$ = 5.6 Hz, 4H), 2.39 – 2.30 (m, 6H), 2.22 (td, $J$ = 6.8, 2.7 Hz, 2H), 1.66 – 1.48 (m, 4H). $^{13}$C NMR (126 MHz, Acetone-$d_6$) δ 150.74 (d, $J$ = 12.7 Hz), 149.97 (d, $J$ = 12.6 Hz), 148.78 (d, $J$ = 12.8 Hz), 148.01 (d, $J$ = 12.6 Hz), 139.39 – 139.04 (m), 139.01, 131.60, 126.33 (dd, $J$ = 6.2, 3.3 Hz), 118.51, 118.37, 117.12, 116.99, 84.06, 68.94, 57.26, 54.72, 31.60, 26.33, 25.93, 17.73. $^{19}$F NMR (471 MHz, Acetone-$d_6$) δ -140.33, -140.38, -142.89, -142.93. HRMS (ESI) calculated for $C_{24}H_{23}F_4N$ $[M+H]^+$: 402.1839, found: 402.1849.

**2.2 Representative preparation of sulfonyl 1,2,3-triazoles by click chemistry**

The synthetic procedure was performed following previously described methods (*68*). To a stirred mixture of alkynes (1.2 equiv., 6 mmol), sulfonyl azides (1.0 equiv., 5 mmol), and copper(I) iodide (CuI, 0.05 mmol) in $CHCl_3$ (30 mL) was slowly added 2,6-lutidine (0.070 mL, 6 mmol) at 0 °C. After stirring the reaction mixture for 12 h at 0 °C, it was diluted with $CH_2Cl_2$ (30 mL) and then quenched with aqueous ammonia chloride solution (50 mL). The mixture was stirred for an additional 30 min and two layers were separated. The aqueous layer was extracted with $CH_2Cl_2$ (50 mL x 3) and the combined organic layers were dried over anhydrous sodium sulfate, filtered, and concentrated in *vacuo*. The crude residue was purified by column chromatography to give the desired products.

**4-(Bis(4-fluorophenyl)methylene)-1-(2-(1-tosyl-1H-1,2,3-triazol-4-yl)ethyl)piperidine (TH207)**

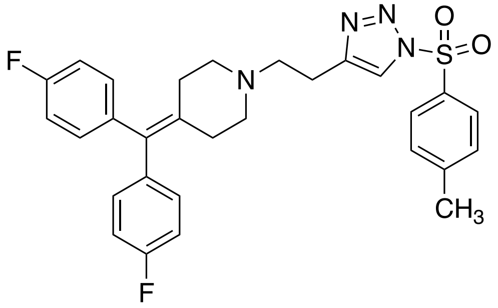

**TH207**

Light brown oil, 63% yield, 1.6 g, R$_f$ = 0.20 (hexane/EtOAc =2:1, UV detection on TLC plate). $^1$H NMR (800 MHz, $CDCl_3$) δ 8.00 7.95 (m, 3H), 7.37 (d, *J* = 8.2 Hz, 2H), 7.10 – 7.02 (m, 4H), 7.00 - 6.95 (m, 4H), 2.93 (t, *J* = 7.5 Hz, 2H), 2.68 (t, *J* = 7.5 Hz, 2H), 2.52 (t, *J* = 5.6 Hz, 4H), 2.44 (s, 3H), 2.37 (t, *J* = 5.7 Hz, 4H). $^{13}$C NMR (201 MHz, $CDCl_3$) δ 162.14, 160.92, 147.15, 146.20, 138.13, 138.11, 135.97, 134.04, 133.27, 131.41, 131.40, 131.38, 131.36, 131.33, 131.27, 131.25, 131.23, 131.21, 131.19, 130.47, 130.37, 130.32, 128.65, 128.62, 128.59, 121.06, 121.03, 121.00, 115.13, 115.11, 115.08, 115.02, 115.01, 114.98, 114.96, 114.95, 114.89, 114.86, 114.84, 56.89, 54.96, 31.73, 23.53, 21.88. $^{19}$F NMR (564 MHz, $CDCl_3$) δ -119.10. HRMS (ESI) calculated for $C_{29}H_{28}F_2N_4O_2S[M+H]^+$: 535.1974, found: 535.1979.

**1-(Bis(4-fluorophenyl)methyl)-4-(2-(1-tosyl-1*H*-1,2,3-triazol-4-yl)ethyl)piperazine (TH208)**

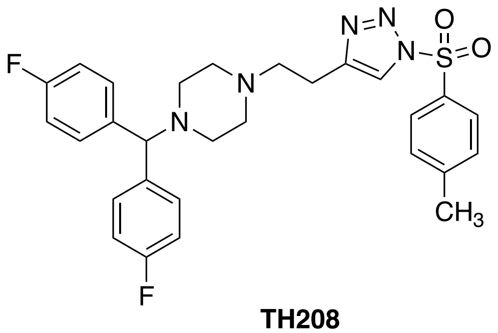

**TH208**

White powder, 56% yield, 1.5 g, R$_f$ = 0.25 (hexane/ EtOAc=2:1, UV detection on TLC plate). $^1$H NMR (800 MHz, $CDCl_3$) δ 8.00 – 7.97 (m, 2H), 7.95 (s, 1H), 7.43 – 7.32 (m, 6H), 7.02 – 6.93 (m, 4H), 4.22 (s, 1H), 2.96 – 2.83 (m, 2H), 2.71 – 2.62 (m, 2H), 2.60 – 2.21 (m, 11H). $^{13}$C NMR

(200 MHz, $CDCl_3$) δ 162.44, 161.22, 147.14, 146.09, 138.22, 133.26, 130.46, 130.38, 130.31, 129.32, 129.29, 129.18, 129.14, 128.66, 128.63, 128.60, 121.04, 120.97, 115.56, 115.45, 115.41, 115.30, 56.90, 53.17, 51.74, 23.21, 21.89, 21.86, 21.82. $^{19}F$ NMR (564 MHz, $CDCl_3$) δ -118.83. HRMS (ESI) calculated for $C_{28}H_{29}F_2N_5O_2S$ $[M+H]^+$: 538.2083, found: 538.2075.

**4-(Bis(4-fluorophenyl)methylene)-1-(2-(1-tosyl-1*H*-1,2,3-triazol-4-yl)ethyl)piperidine (TH220)**

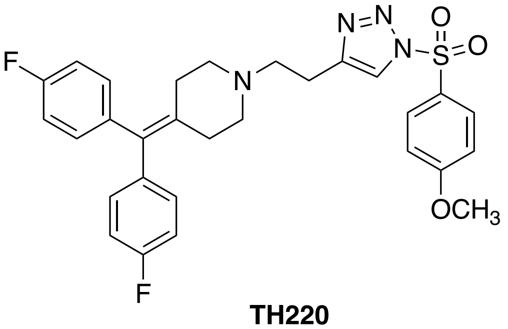


**TH220**

White powder, 69% yield, 1.9 g, $R_f$ = 0.35 (hexane/ EtOAc=1:1, UV detection on TLC plate). $^1H$ NMR (800 MHz, $CDCl_3$) δ 8.08 – 8.01 (m, 2H), 7.97 (s, 1H), 7.09 – 7.04 (m, 4H), 7.04 – 7.01 (m, 2H), 7.00 - 6.96 (m, 4H), 3.88 (s, 3H), 2.98 – 2.85 (m, 2H), 2.74 – 2.63 (m, 2H), 2.57 – 2.48 (m, 4H), 2.42 – 2.10 (m, 4H). $^{13}C$ NMR (200 MHz, $CDCl_3$) δ 165.27, 162.15, 160.93, 146.07, 138.11, 135.98, 134.04, 132.89, 131.5 – 130.89 (m, 4C), 131.40, 131.36, 131.33, 131.27, 131.23, 131.20, 131.17, 131.14, 131.12, 129.40, 120.91, 120.87, 120.84, 115.30 – 114.74 (m, 4C), 57.11 – 56.70 (m, 1C), 55.94 (q, J = 49.0 Hz, 1C), 54.94, 31.64, 23.43. $^{19}F$ NMR (564 MHz, $CDCl_3$) δ -119.12. HRMS (ESI) calculated for $C_{29}H_{28}F_2N_4O_3S$ $[M+H]^+$: 551.1923, found: 551.1919.

**4-(Bis(4-fluorophenyl)methylene)-1-(2-(1-((4-fluorophenyl)sulfonyl)-1*H*-1,2,3-triazol-4-yl)ethyl)piperidine (TH221)**

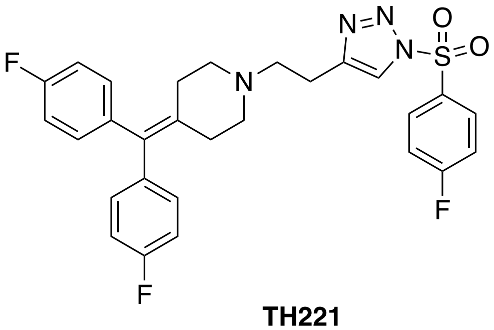

**TH221**

Light yellow oil, 45% yield, 1.2 g, $R_f$ = 0.30 (hexane/EtOAc=2:1, UV detection on TLC plate). $^1$H NMR (800 MHz, $CDCl_3$) δ 8.17 – 8.13 (m, 2H), 7.99 (s, 1H), 7.30 – 7.24 (m, 2H), 7.09 – 7.03 (m, 4H), 7.01 – 6.94 (m, 4H), 2.95 (t, *J* = 7.5 Hz, 2H), 2.69 (t, *J* = 7.5 Hz, 2H), 2.53 (t, *J* = 5.6 Hz, 4H), 2.37 (t, *J* = 5.6 Hz, 4H). $^{13}$C NMR (200 MHz, $CDCl_3$) δ 167.47, 166.18, 162.16, 160.93, 146.34, 138.85, 138.08 (d, *J* = 3.5 Hz), 135.83, 134.13, 132.23 (d, *J* = 3.2 Hz), 131.75 (d, *J* = 10.2 Hz), 131.29 (d, *J* = 7.5 Hz), 121.06, 117.31 (d, *J* = 23.1 Hz), 115.00 (d, *J* = 21.2 Hz),77.21, 77.05, 76.89, 56.81, 54.94, 31.61, 23.38. $^{19}$F NMR (564 MHz, $CDCl_3$) δ -102.34. HRMS (ESI) calculated for $C_{28}H_{25}F_3N_4O_2S$ $[M+H]^+$: 539.1723, found: 539.1716.

**4-(bis(4-fluorophenyl)methylene)-1-(2-(1-(cyclopropylsulfonyl)-1*H*-1,2,3-triazol-4-yl)ethyl) piperidine (TH223)**

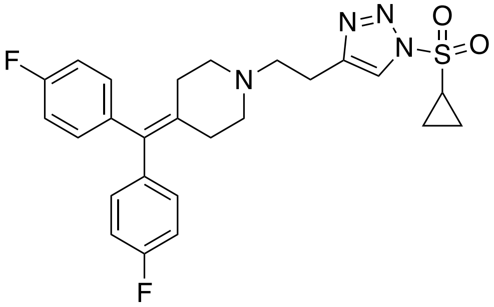

**TH223**

Light brown oil, 70% yield, 1.7 g, $R_f$ = 0.25 (hexane/EtOAc =1:1, UV detection on TLC plate). $^1$H NMR (800 MHz, $CDCl_3$) δ 7.92 (s, 1H), 7.12 – 7.04 (m, 4H), 7.00 – 6.92 (m, 4H), 2.98 (t, *J* = 7.5 Hz, 2H), 2.86 (tt, *J* = 7.9, 4.7 Hz, 1H), 2.71 (t, *J* = 7.5 Hz, 2H), 2.54 (t, *J* = 5.6 Hz, 4H), 2.44 – 2.30 (m, 4H), 1.67 – 1.47 (m, 2H), 1.36 – 1.16 (m, 2H). $^{13}$C NMR (200 MHz, $CDCl_3$) δ 162.12, 160.90, 145.98, 138.13, 138.12, 135.96, 134.04, 131.33, 131.29, 121.42, 115.02, 114.92, 56.90,

54.94, 32.16, 23.38, 7.80. $^{19}$F NMR (564 MHz, CDCl$_3$) δ -119.07. HRMS (ESI) calculated for $C_{25}H_{26}F_2N_4O_2S$ $[M+H]^+$: 485.1717, found: 485.1796.

**4-((4-(2-(4-(Bis(4-fluorophenyl)methylene)piperidin-1-yl)ethyl)-1*H*-1,2,3-triazol-1-yl)sulfonyl)-*N*-propylbenzamide (TH225)**

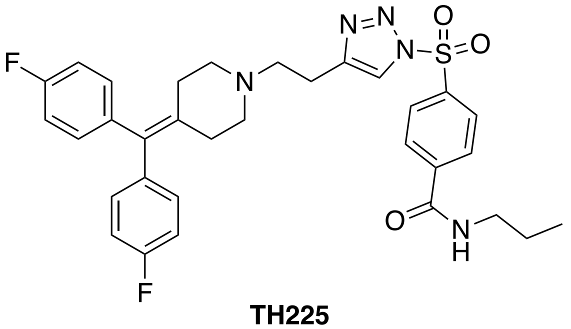


**TH225**

White powder, 80% yield, 2.4 g, $R_f$ = 0.20 (hexane/EtOAc=1:1, UV detection on TLC plate). $^1$H NMR (800 MHz, CDCl$_3$) δ 8.14 – 8.10 (m, 2H), 8.00 (s, 1H), 7.96 – 7.92 (m, 2H), 7.09 – 7.01 (m, 4H), 7.00 – 6.94 (m, 4H), 6.43 (t, *J* = 5.9 Hz, 1H), 3.45 – 3.35 (m, 2H), 2.92 (t, *J* = 7.5 Hz, 2H), 2.67 (t, *J* = 7.5 Hz, 2H), 2.52 (t, *J* = 5.9 Hz, 4H), 2.36 (t, *J* = 5.6 Hz, 4H), 1.62 (h, *J* = 7.4 Hz, 2H), 0.96 (t, *J* = 7.4 Hz, 3H). $^{13}$C NMR (200 MHz, CDCl$_3$) δ 165.28, 162.15, 160.93, 146.53, 141.43, 138.41, 138.08, 135.78, 134.16, 131.47-131.09 (m, 2C), 128.85-128.80 (m, 2C),128.41-128.23 (m, 2C), 121.32, 121.25, 115.15- 114.87 (m, 2C) 56.75, 54.92, 42.11, 31.60, 23.35, 22.86, 22.76, 11.47, 11.34. $^{19}$F NMR (564 MHz, CDCl$_3$) δ -119.07. HRMS (ESI) calculated for $C_{32}H_{33}F_2N_5O_3S$ $[M+H]^+$: 606.2345, found: 606.2329.

**4-(Bis(4-fluorophenyl)methylene)-1-(3-(1-((4-methoxyphenyl)sulfonyl)-1H-1,2,3-triazol-4-yl)propyl)piperidine (XJ-3-9)**

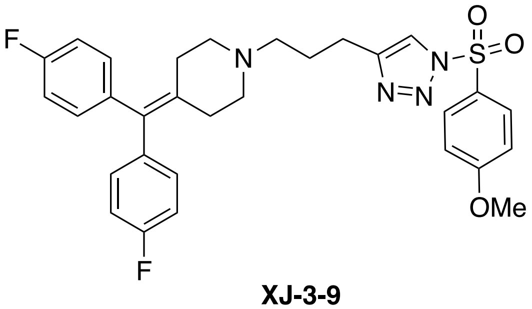


**XJ-3-9**

Light brown oil, 74% yield, 2.1 g, $R_f$ = 0.40 (hexane/acetone=3:1, UV detection on TLC plate) $^1$H NMR (800 MHz, $CDCl_3$) δ 8.10 – 7.94 (m, 2H), 7.85 (s, 1H), 7.05 – 7.02 (m, 4H), 7.02 – 7.00 (m, 2H), 6.98 – 6.94 (m, 4H), 3.87 (s, 3H), 2.74 (t, $J$ = 7.6 Hz, 2H), 2.44 (t, $J$ = 5.7 Hz, 4H), 2.40 – 2.36 (m, 2H), 2.33 (t, $J$ = 5.7 Hz, 4H), 1.86 (p, $J$ = 7.6 Hz, 2H). $^{13}$C NMR (201 MHz, $CDCl_3$) δ 164.78, 161.65, 160.43, 147.21, 137.72, 137.70, 135.85, 133.29, 130.86, 130.82, 130.65, 126.75, 119.82, 114.54, 114.43, 56.99, 55.46, 54.69, 31.20, 25.85, 22.89. $^{19}$F NMR (564 MHz, $CDCl_3$) δ -119.21. HRMS (ESI) calculated for $C_{30}H_{30}F_2N_4O_3S$ $[M+H]^+$: 565.2079, found: 565.2069.

**1-(4-(Bis(4-fluorophenyl)methylene)piperidin-1-yl)-3-(1-((4-methoxyphenyl)sulf-onyl)-1*H*-1,2,3-triazol-4-yl)propan-1-one (XJ-3-23)**

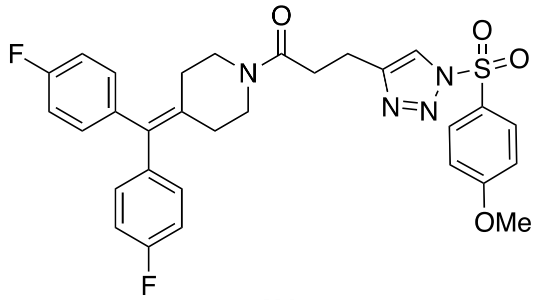


**XJ-3-23**

Light yellow powder, 68% yield, 2.0 g, $R_f$ = 0.10 (hexane/acetone=3:1, UV detection on TLC plate). $^1$H NMR (800 MHz, $CDCl_3$) δ 8.06 – 7.99 (m, 2H), 7.95 (s, 1H), 7.06 – 7.01 (m, 5H), 7.01 – 6.96 (m, 5H), 3.87 (s, 3H), 3.61 (t, $J$ = 5.9 Hz, 2H), 3.43 (t, $J$ = 5.8 Hz, 2H), 3.10 – 3.03 (m, 2H), 2.78 – 2.67 (m, 2H), 2.27 – 2.34 (m, 4H). $^{13}$C NMR (201 MHz, $CDCl_3$) δ 169.22, 161.22 (d, $J$ = 245.9 Hz),160.61, 146.34, 137.20, 137.18, 137.09, 137.07, 135.47, 133.55, 130.70, 130.66, 126.74, 120.86, 114.78, 114.67, 114.54, 55.46, 45.88, 42.59, 31.60, 31.43, 30.68, 20.46. $^{19}$F NMR (564 MHz, $CDCl_3$) δ -118.52. HRMS (ESI) calculated for $C_{30}H_{28}F_2N_4O_4S$ $[M+H]^+$: 579.1872, found: 579.1870.

**4-(Bis(4-fluorophenyl)methylene)-1-(4-(1-((4-methoxyphenyl)sulfonyl)-1H-1,2,3-triazol-4-yl)butyl)piperidine (XJ-3-41)**

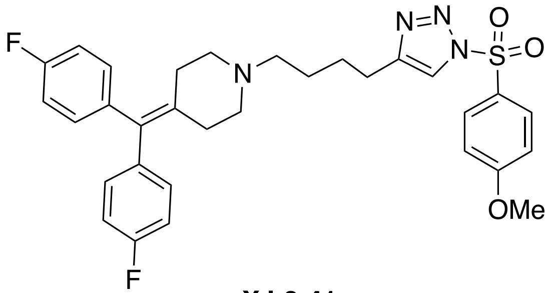

**XJ-3-41**

Light brown oil, 82% yield, 2.4 g, $R_f$ = 0.20 (hexane/acetone=4:1, UV detection on TLC plate). $^1$H NMR (800 MHz, Acetone-$d_6$) δ 8.29 (s, 1H), 8.10 – 8.04 (m, 2H), 7.22 – 7.18 (m, 2H), 7.18 – 7.13 (m, 4H), 7.10 – 7.04 (m, 4H), 3.93 (s, 3H), 2.75 – 2.64 (m, 2H), 2.42 (t, *J* = 5.5 Hz, 4H), 2.33 (t, *J* = 7.2 Hz, 2H), 2.30 (t, *J* = 5.6 Hz, 4H), 1.75 – 1.67 (m, 2H), 1.51 (p, *J* = 7.3 Hz, 2H). $^{13}$C NMR (200 MHz, Acetone-$d_6$) δ 204.84, 164.98, 161.59, 160.38, 147.64, 138.18, 138.16, 136.52, 132.93, 131.02, 130.98, 130.42, 126.96, 120.58, 114.79, 114.39, 114.29, 57.11, 55.24, 54.50, 31.13, 26.19, 25.87, 24.41. $^{19}$F NMR (564 MHz, $CDCl_3$) δ -119.17, -164.90. HRMS (ESI) calculated for $C_{31}H_{32}F_2N_4O_3S$ $[M+H]^+$: 579.2236, found: 579.2248.

**4-(Bis(4-fluorophenyl)methylene)-1-(5-(1-((4-methoxyphenyl)sulfonyl)-1*H*-1,2,3-triazol-4-yl) pentyl)piperidine (XJ-3-65)**

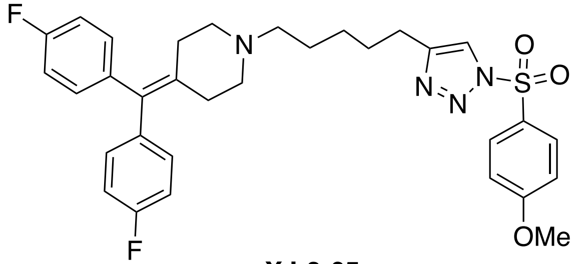

**XJ-3-65**

Light brown oil, 67% yield, 2.0 g, $R_f$ = 0.20 (hexane/acetone=3:1, UV detection on TLC plate). $^1$H NMR (800 MHz, Acetone-$d_6$) δ 8.29 (s, 1H), 8.08 – 8.04 (m, 2H), 7.22 – 7.19 (m, 2H), 7.18 – 7.13 (m, 4H), 7.10 – 7.04 (m, 4H), 3.93 (s, 3H), 2.72 – 2.66 (m, 2H), 2.46 – 2.39 (m, 4H), 2.32 – 2.25 (m, 6H), 1.71 – 1.63 (m, 2H), 1.53 – 1.44 (m, 2H), 1.40 – 1.32 (m, 2H). $^{13}$C NMR (200 MHz, Acetone-$d_6$) δ 204.91, 164.98, 161.60, 160.38, 147.64, 138.17, 138.16, 136.52, 132.95, 131.04, 131.00, 130.44, 126.96, 120.58, 114.81, 114.41, 114.31, 57.37, 55.27, 54.52, 31.13, 28.17, 26.24,

26.07, 24.52. $^{19}$F NMR (564 MHz, CDCl$_3$) δ -119.21, -164.90. HRMS (ESI) calculated for $C_{32}H_{34}F_2N_4O_3S$ $[M+H]^+$: 593.2392, found: 593.2384.

**4-Butyl-1-((4-methoxyphenyl)sulfonyl)-1*H*-1,2,3-triazole (XJ-4-5)**

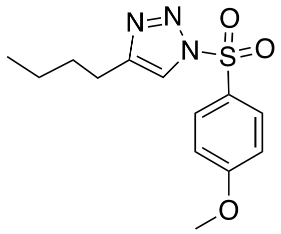


**XJ-4-5**

White powder, 85% yield, 1.3 g, $R_f$ = 0.50 (hexane/EtOAc =4:1, UV detection on TLC plate). $^1$H NMR (800 MHz, Acetone-$d_6$) δ 8.37 (s, 1H), 8.18 – 8.10 (m, 2H), 7.32 – 7.23 (m, 2H), 4.03 (s, 3H), 2.80 – 2.73 (m, 2H), 1.75 – 1.66 (m, 2H), 1.48 – 1.29 (m, 2H), 1.01 – 0.93 (m, 2H). $^{13}$C NMR (201 MHz, Acetone-$d_6$) δ 164.97, 147.73, 130.45, 126.91, 120.58, 114.81, 55.32, 30.47, 24.29, 21.52, 12.81. $^{19}$F NMR (564 MHz, CDCl$_3$) δ -119.21, -164.90. HRMS (ESI) calculated for $C_{13}H_{17}N_3O_3S$ $[M+H]^+$: 296.1063, found: 296.1057.

**4-(Bis(4-fluorophenyl)methylene)-1-(4-(1-(propylsulfonyl)-1H-1,2,3-triazol-4-yl)butyl)piperidine (XJ-4-27)**

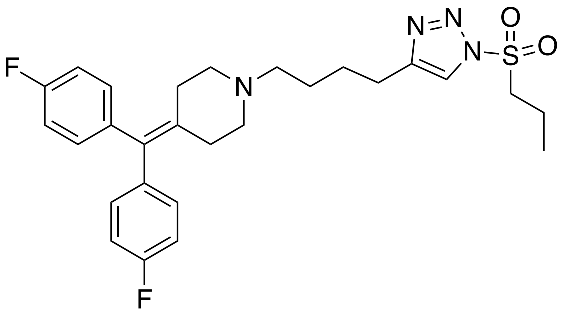


**XJ-4-27**

Light brown oil, 70% yield, 1.8 g, $R_f$ = 0.40 (hexane/acetone=3:1, UV detection on TLC plate). $^1$H NMR (800 MHz, Acetone-$d_6$) δ 8.21 (s, 1H), 7.24 – 7.12 (m, 4H), 7.12 – 6.90 (m, 4H), 3.82 – 3.67 (m, 2H), 2.79 (t, *J* = 7.6 Hz, 2H), 2.44 (t, *J* = 5.7 Hz, 4H), 2.36 (t, *J* = 7.2 Hz, 2H), 2.33 – 2.26 (m, 4H), 1.80 – 1.67 (m, 4H), 1.55 (p, *J* = 7.4 Hz, 2H), 1.01 (t, *J* = 7.5 Hz, 3H). $^{13}$C NMR (201 MHz, Acetone-$d_6$) δ 161.59, 160.38, 147.37, 138.18, 138.17, 136.57, 132.91, 131.00, 130.96,

121.31, 114.36, 114.25, 57.12, 55.69, 54.52, 31.16, 26.24, 25.84, 24.35, 16.29, 11.14. $^{19}$F NMR (564 MHz, Acetone-$d_6$) δ -117.96. HRMS (ESI) calculated for $C_{27}H_{32}F_2N_4O_2S$ $[M+H]^+$: 515.2287, found: 515.2298.

**1-(4-(1-(Benzo[*d*][1,3]dioxol-5-ylsulfonyl)-1*H*-1,2,3-triazol-4-yl)butyl)-4-(bis(4-fluorophenyl) methylene)piperidine (XJ-4-71)**

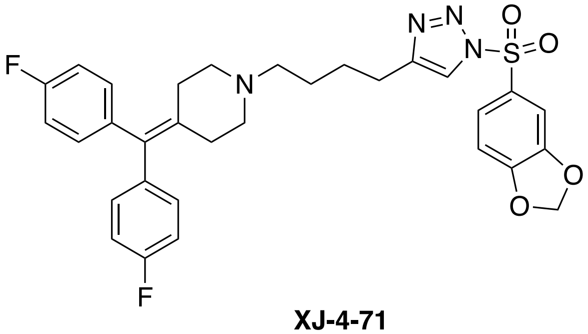

**XJ-4-71**

Light yellow oil, 78% yield, 2.3 g, $R_f$ = 0.30 (hexane/acetone=3:1, UV detection on TLC plate). $^1$H NMR (800 MHz, Acetone-$d_6$) δ 8.30 (s, 1H), 7.75 – 7.72 (m, 1H), 7.51 – 7.45 (m, 1H), 7.18 – 7.43 (m, 4H), 7.12 – 7.09 (m, 1H), 7.09 – 7.05 (m, 4H), 6.23 (s, 2H), 2.84 – 2.59 (m, 2H), 2.59 – 2.38 (m, 4H), 2.34 – 2.31 (m, 2H), 2.32 – 2.28 (m, 4H), 1.73 – 1.67 (m, 2H), 1.54 – 1.48 (m, 2H). $^{13}$C NMR (200 MHz, Acetone-$d_6$) δ 161.60, 160.38, 153.78, 148.57, 147.70, 138.18, 138.16, 136.51, 132.93, 131.02, 131.00, 130.98, 130.96, 128.49, 124.87, 120.72, 114.36, 114.25, 108.33, 106.95, 103.15, 57.08, 54.49, 31.11, 26.15, 25.84, 24.37. $^{19}$F NMR (564 MHz, Acetone-$d_6$) δ -117.92. HRMS (ESI) calculated for $C_{31}H_{30}F_2N_4O_4S$ $[M+H]^+$: 593.2029, found: 593.2015.

**1-(4-(1-(Benzo[*d*][1,3]dioxol-5-ylsulfonyl)-1*H*-1,2,3-triazol-4-yl)butyl)-4-(bis(3,4-difluor-op-henyl)methylene)piperidine (XJ-4-85)**

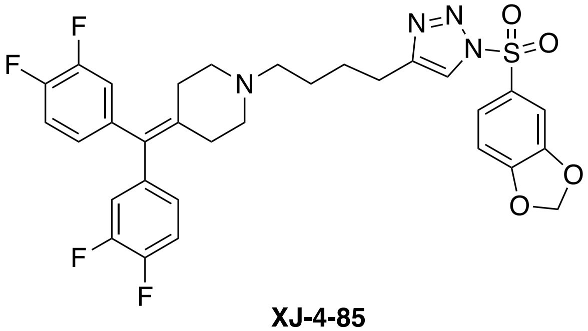

**XJ-4-85**

Light yellow oil, 76% yield, 2.4 g, $R_f$ = 0.25 (hexane/acetone=3:1, UV detection on TLC plate). $^1$H NMR (800 MHz, Acetone-$d_6$) δ 8.30 (s, 1H), 7.74 (dd, $J$ = 8.4, 2.0 Hz, 1H), 7.47 (d, $J$ = 2.0 Hz, 1H), 7.28 (dt, $J$ = 10.7, 8.4 Hz, 2H), 7.12 – 7.07 (m, 3H), 7.02 – 6.96 (m, 2H), 6.24 (s, 2H), 2.73 (t, $J$ = 7.6 Hz, 2H), 2.43 (t, $J$ = 5.6 Hz, 4H), 2.33 (t, $J$ = 7.2 Hz, 2H), 2.31 – 2.28 (m, 4H), 1.71 (p, $J$ = 7.6 Hz, 2H), 1.51 (p, $J$ = 7.3 Hz, 2H). $^{13}$C NMR (126 MHz, Acetone-$d_6$) δ 154.22, 150.75 (d, $J$ = 12.7 Hz), 149.98 (d, $J$ = 12.7 Hz), 149.01, 148.79 (d, $J$ = 12.9 Hz), 148.17, 148.02 (d, $J$ = 12.6 Hz), 139.21 (dd, $J$ = 5.7, 4.0 Hz), 139.17, 139.01, 131.60, 129.00, 126.32 (dd, $J$ = 6.2, 3.3 Hz), 125.34, 121.14, 118.52, 118.39, 117.12, 116.99, 108.79, 107.47, 103.60, 57.47, 54.74, 31.60, 26.64, 26.36, 24.88. $^{19}$F NMR (471 MHz, Acetone-$d_6$) δ -140.32, -140.37, -142.88, -142.92. HRMS (ESI) calculated for $C_{31}H_{28}F_4N_4O_4S$ $[M+H]^+$: 629.1840, found: 629.1817.

**((2*S*,5*R*)-4-(4-(1-(Benzo[*d*][1,3]dioxol-5-ylsulfonyl)-1*H*-1,2,3-triazol-4-yl)butyl)-2,5-dime-th-ylpiperazin-1-yl)(4-fluorophenyl)methanone (XJ-4-97)**

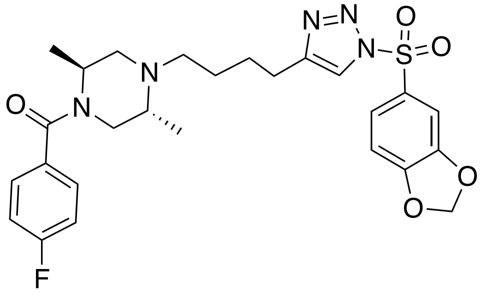

**XJ-4-97**

White solid, 62% yield, 1.7 g, $R_f$ = 0.20 (hexane/acetone=3:1, UV detection on TLC plate). $^1$H NMR (800 MHz, Acetone-$d_6$) δ 8.31 (s, 1H), 7.75 (dt, $J$ = 8.3, 1.9 Hz, 1H), 7.47 (t, $J$ = 1.8 Hz, 1H), 7.45 – 7.38 (m, 2H), 7.22 – 7.16 (m, 2H), 7.14 – 7.10 (m, 1H), 6.25 (s, 2H), 3.38 (s, 1H), 2.95 (s, 1H), 2.74 (t, $J$ = 7.6 Hz, 2H), 2.68 (dd, $J$ = 12.0, 4.1 Hz, 2H), 2.47 – 2.40 (m, 2H), 2.38 – 2.28 (m, 2H), 1.82 – 1.63 (m, 2H), 1.48 (p, $J$ = 7.3 Hz, 2H), 1.28 (d, $J$ = 6.8, 3H), 0.89 (d, $J$ = 6.6 Hz, 3H). $^{13}$C NMR (201 MHz, Acetone-$d_6$) δ 168.75, 162.41 (d, $J$ = 246.5 Hz), 153.81, 148.59, 147.73, 133.22, 128.63, 128.59, 124.88, 120.72, 114.75, 114.65, 108.35, 106.94, 103.19, 52.85, 51.87, 48.30, 28.79, 28.69, 28.60, 28.50, 28.40, 28.31, 28.21, 25.94, 25.84, 24.26, 15.50, 5.82. $^{19}$F NMR (564 MHz, acetone-$d_6$) δ -113.75. HRMS (ESI) calculated for $C_{26}H_{30}FN_5O_5S$ $[M+H]^+$: 544.2024, found: 544.2002.

**1-(4-(1*H*-1,2,3-Triazol-4-yl)butyl)-4-(bis(3,4-difluorophenyl)methylene)piperidine (XJ-4-119)**

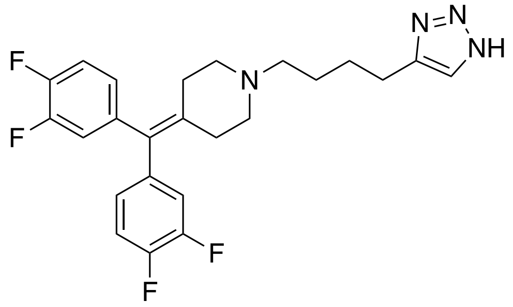

**XJ-4-119**

Colorless oil, 57% yield, 0.4 g, $R_f$ = 0.10 (hexane/acetone=3:1, UV detection on TLC plate). $^{1}H$ NMR (500 MHz, Acetone-$d_6$) δ 7.55 (s, 1H), 7.35 – 7.23 (m, 2H), 7.18 – 7.09 (m, 2H), 7.05 – 6.96 (m, 2H), 2.84 (brs, 1H), 2.74 (t, *J* = 7.5 Hz, 2H), 2.48 (t, *J* = 5.5 Hz, 4H), 2.38 (t, *J* = 7.2 Hz, 2H), 2.33 (t, *J* = 5.6 Hz, 4H), 1.77 – 1.66 (m, 2H), 1.62 – 1.49 (m, 2H). $^{13}C$ NMR (126 MHz, Acetone-$d_6$) δ 150.73 (d, *J* = 12.9 Hz), 149.97 (d, *J* = 12.7 Hz), 148.77 (d, *J* = 12.8 Hz), 148.01 (d, *J* = 12.7 Hz), 139.20 (t, *J* = 5.0 Hz), 147.96, 139.24, 139.20, 139.16, 138.90, 131.84, 131.63, 130.16, 126.33 (dd, *J* = 6.2, 3.3 Hz), 118.51, 118.38, 117.14, 117.00, 57.56, 54.71, 31.52, 27.03, 26.40, 24.52. $^{19}F$ NMR (471 MHz, Acetone-$d_6$) δ -140.30, -140.35, -142.85, -142.90. HRMS (ESI) calculated for $C_{26}H_{30}FN_5O_5S$ $[M+H]^+$: 445.2010, found: 445.2004.

### 4. NMR AND HRMS SPECTRA

#### $^{1}$H NMR (800 MHz, $CDCl_3$) spectrum of XJ-3-7

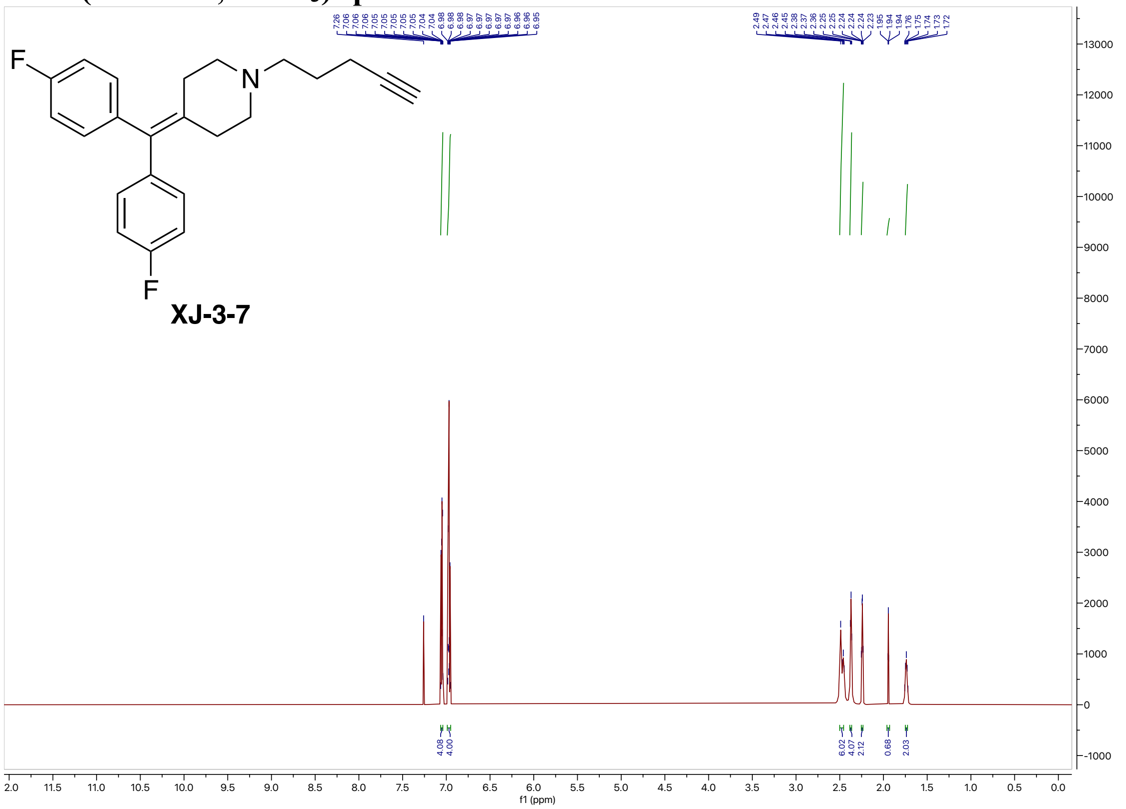


#### $^{13}$C NMR (200 MHz, $CDCl_3$) spectrum of XJ-3-7

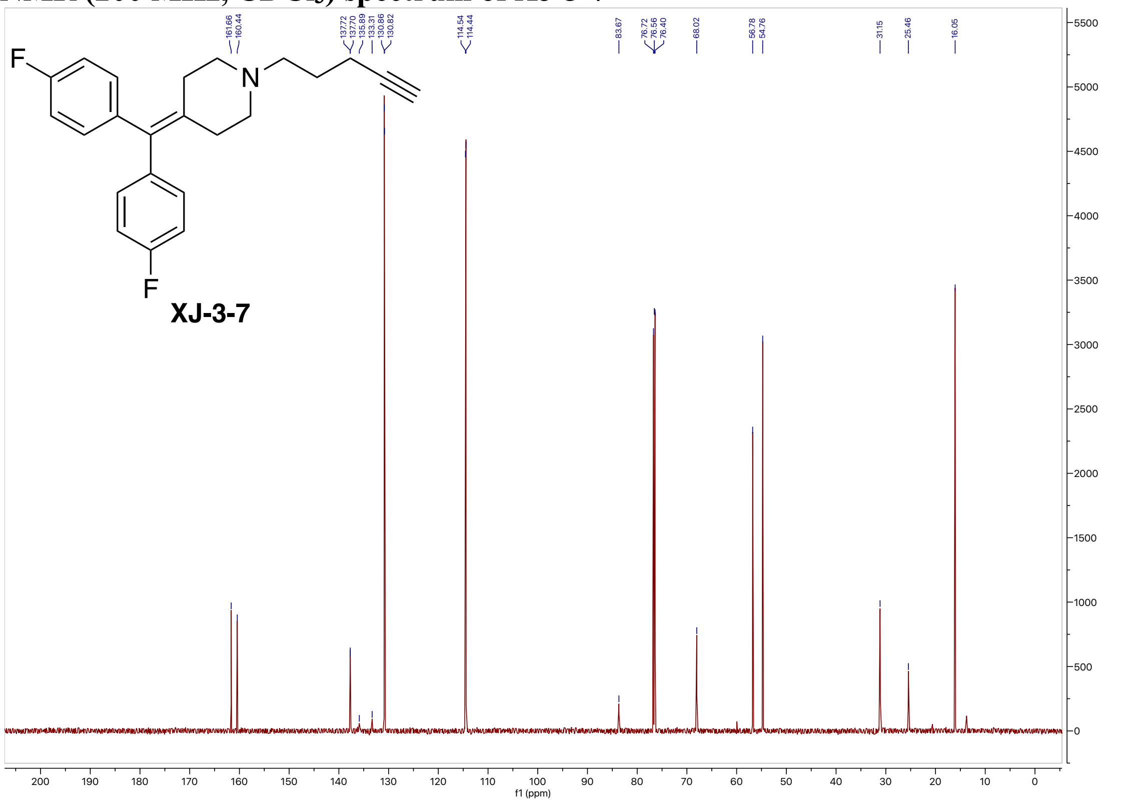

#### $^{19}$F NMR (564 MHz, $CDCl_3$) spectrum of XJ-3-7

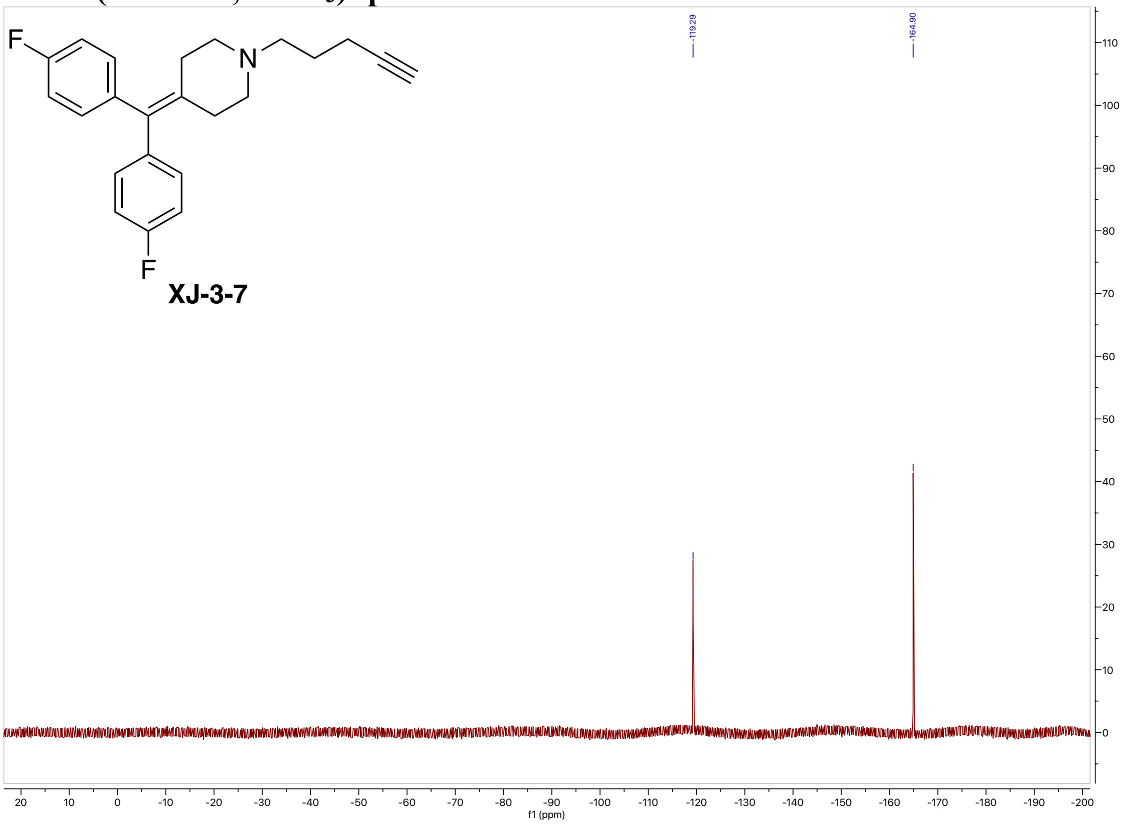


#### High-resolution mass spectrometry of XJ-3-7

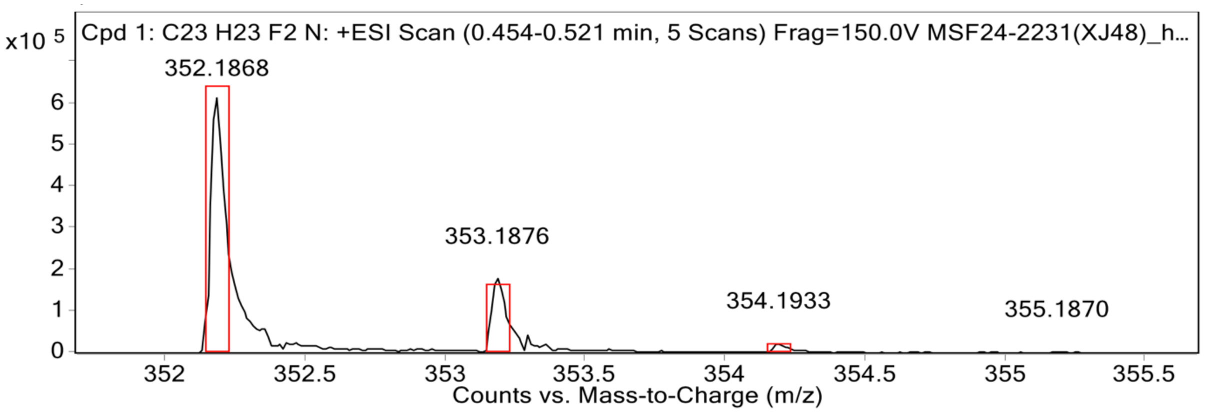


**MS Spectrum Peak List**

| Obs. m/z | Calc. m/z | Charge | Abundance | Formula | Ion Species | Tgt Mass Error (ppm) |
|---|---|---|---|---|---|---|
| 352.1868 | 352.1871 | 1 | 615600 | C23H23F2N | (M+H)+ | 0.88 |
| 353.1876 | 353.1904 | 1 | 179289 | C23H23F2N | (M+H)+ | 8.01 |
| 354.1933 | 354.1937 | 1 | 23615 | C23H23F2N | (M+H)+ | 1.22 |
| 355.1870 | 355.1970 | 1 | 3385 | C23H23F2N | (M+H)+ | 28.19 |
| 356.1794 | 356.2003 | 1 | 940 | C23H23F2N | (M+H)+ | 58.8 |

#### [1]H NMR (800 MHz, CDCl3) spectrum of XJ-3-19

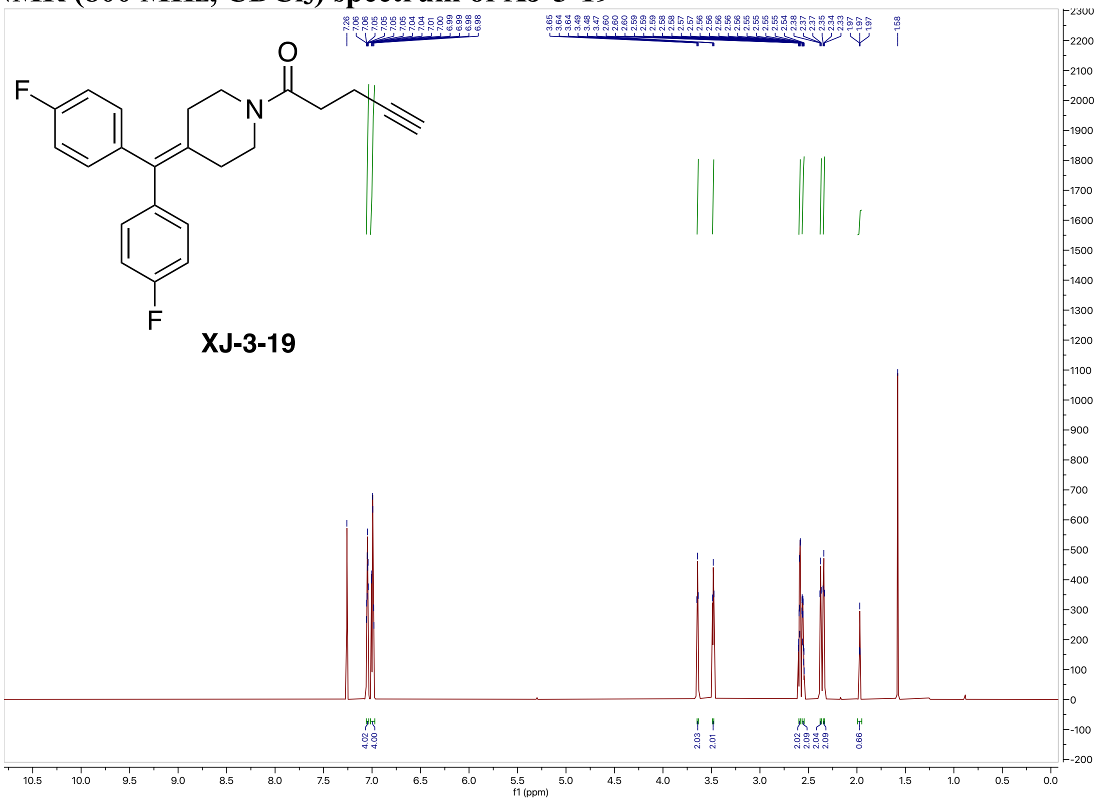

#### [13]C NMR (200 MHz, CDCl3) spectrum of XJ-3-19

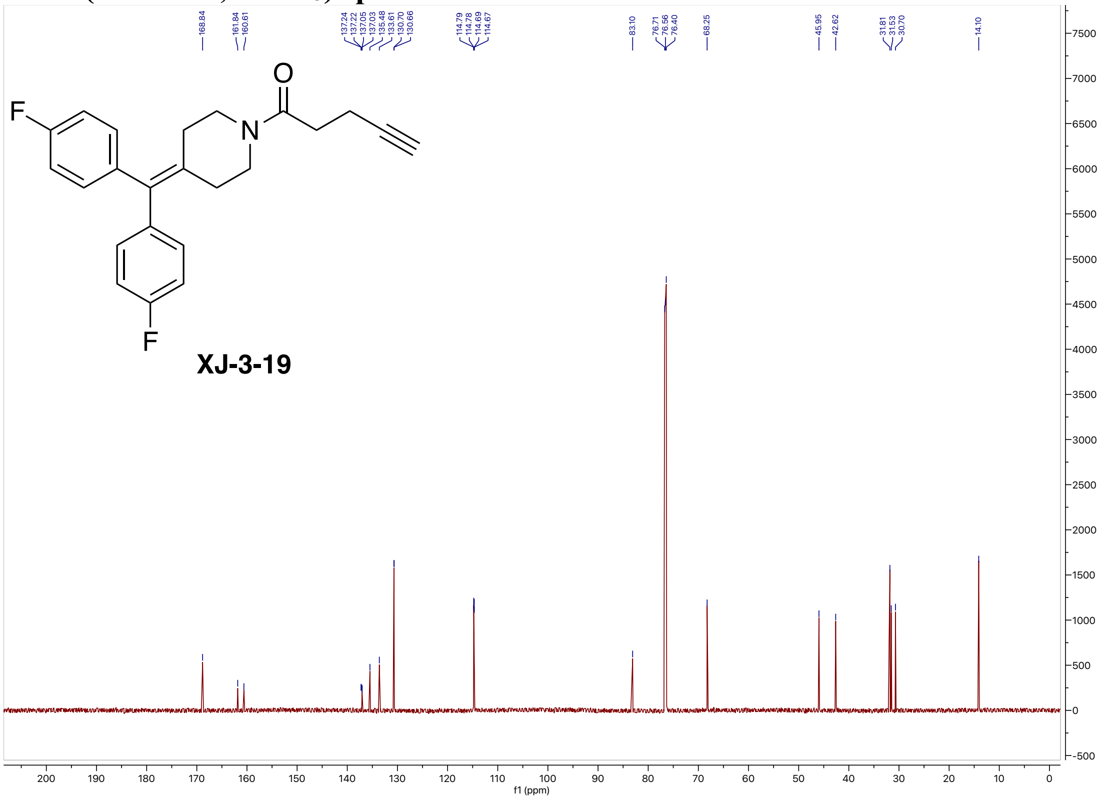

#### $^{19}$F NMR (564 MHz, $CDCl_3$) spectrum of XJ-3-19

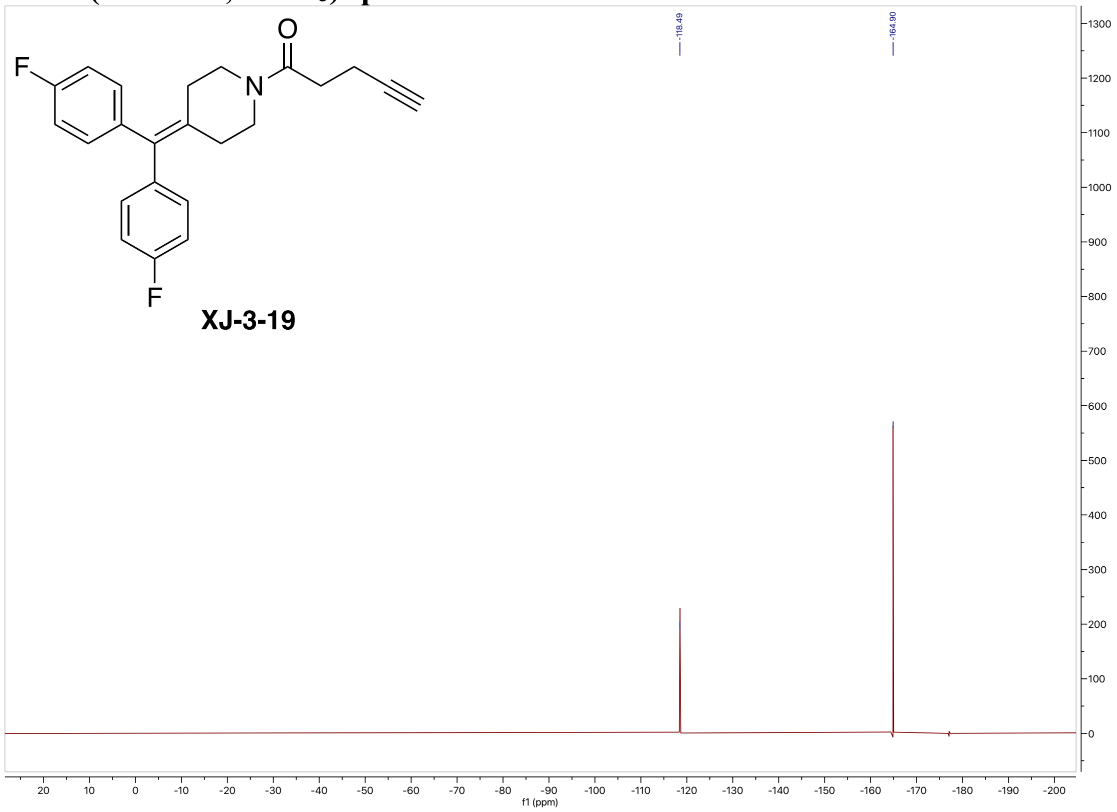


#### High-resolution mass spectrometry of XJ-3-19

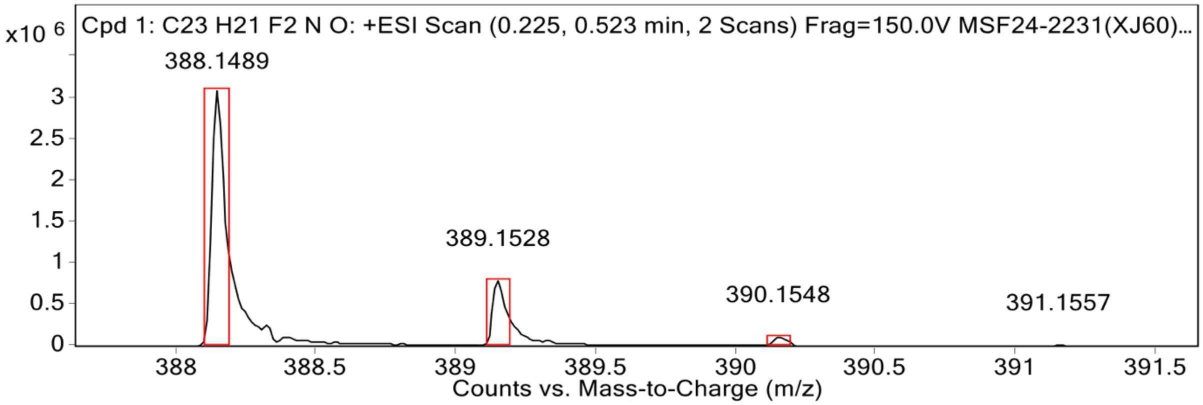


**MS Spectrum Peak List**

| Obs. m/z | Calc. m/z | Charge | Abundance | Formula | Ion Species | Tgt Mass Error (ppm) |
|---|---|---|---|---|---|---|
| 388.1489 | 388.1483 | 1 | 3092621 | C23H21F2NO | (M+Na)+ | -1.34 |
| 389.1528 | 389.1516 | 1 | 789793 | C23H21F2NO | (M+Na)+ | -3.08 |
| 390.1548 | 390.1548 | 1 | 109992 | C23H21F2NO | (M+Na)+ | -0.04 |
| 391.1557 | 391.1578 | 1 | 12086 | C23H21F2NO | (M+Na)+ | 5.44 |

#### [1]H NMR (800 MHz, Acetone-$d_6$) spectrum of XJ-3-61

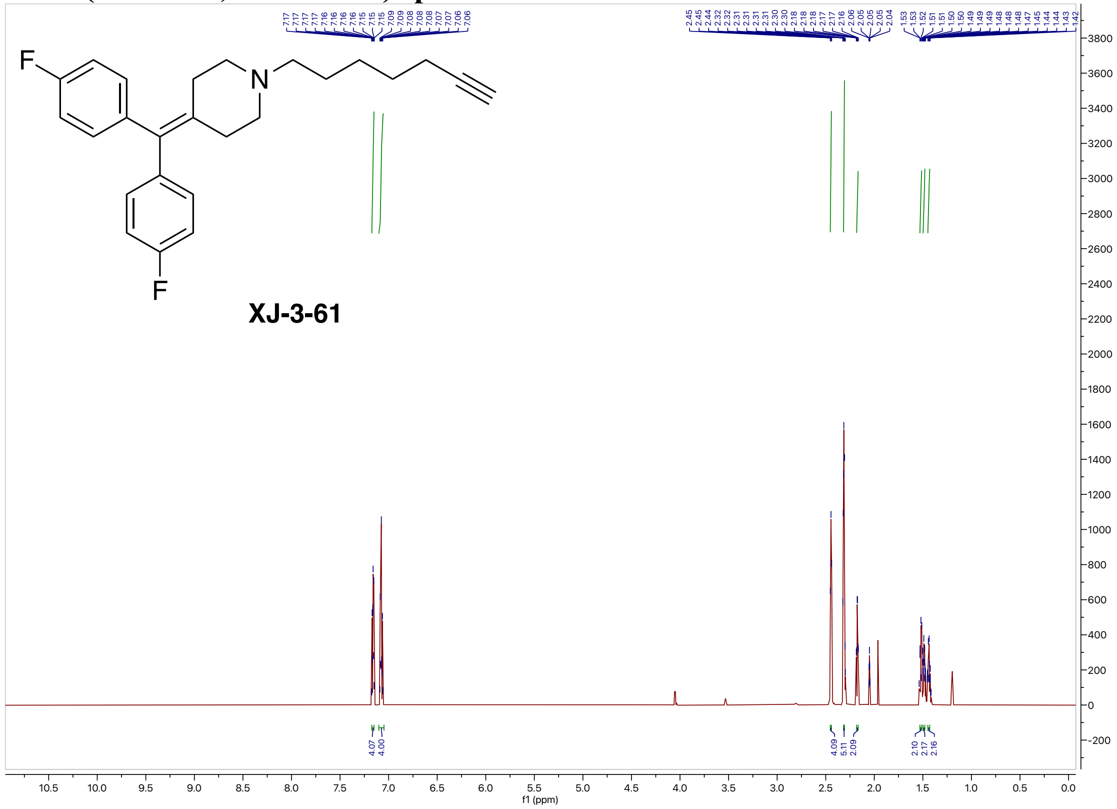

#### [13]C NMR (200 MHz, Acetone-$d_6$) spectrum of XJ-3-61

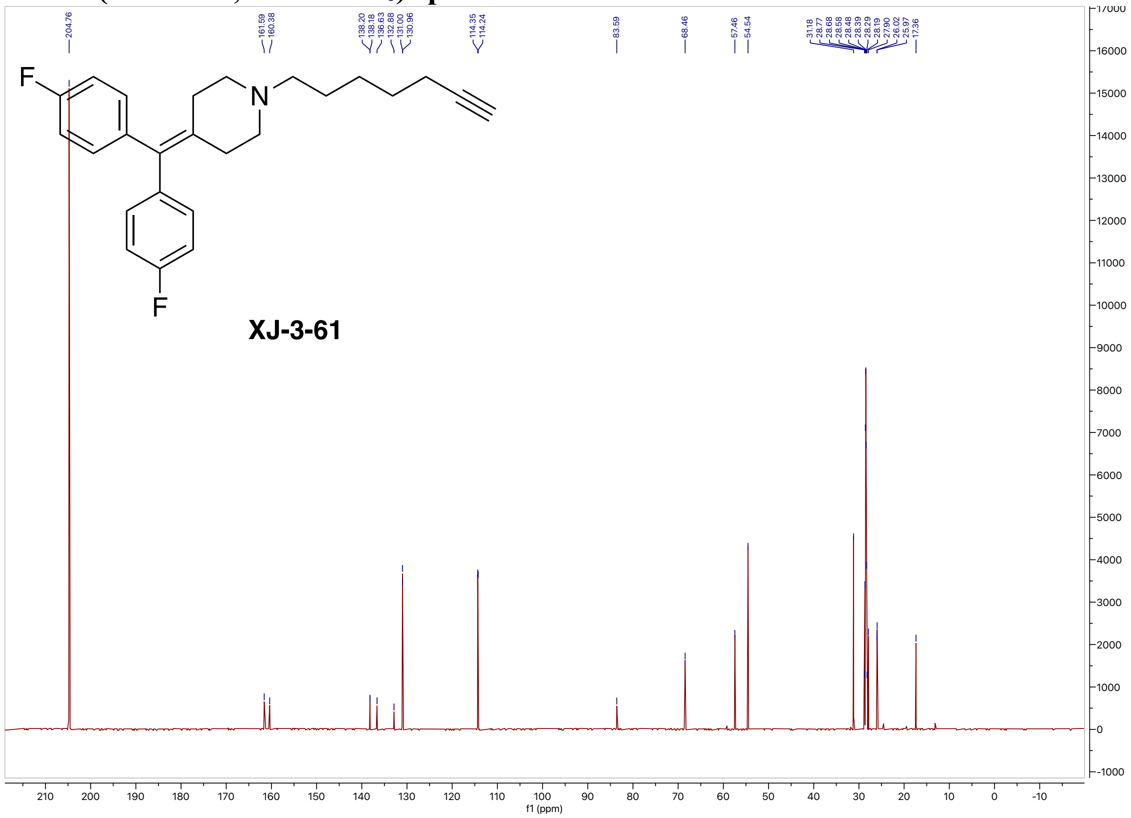

#### $^{19}$F NMR (564 MHz, Acetone-$d_6$) spectrum of XJ-3-61

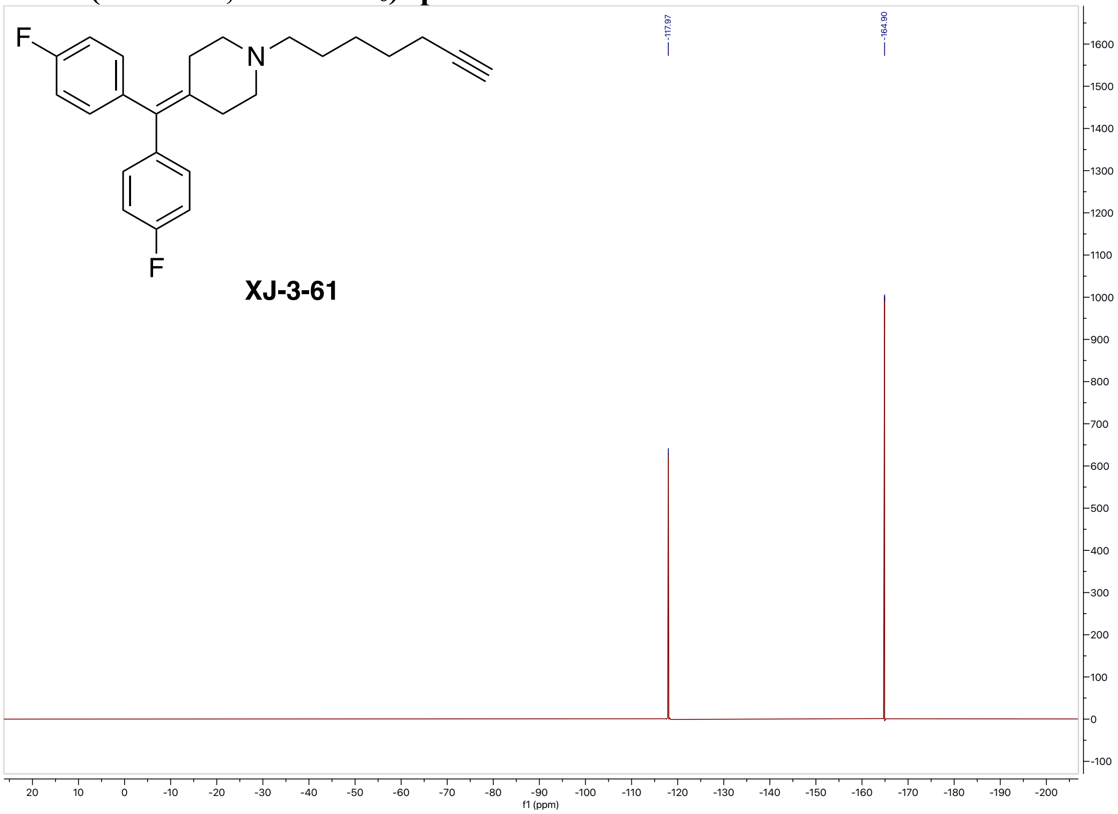


#### High-resolution mass spectrometry of XJ-3-61

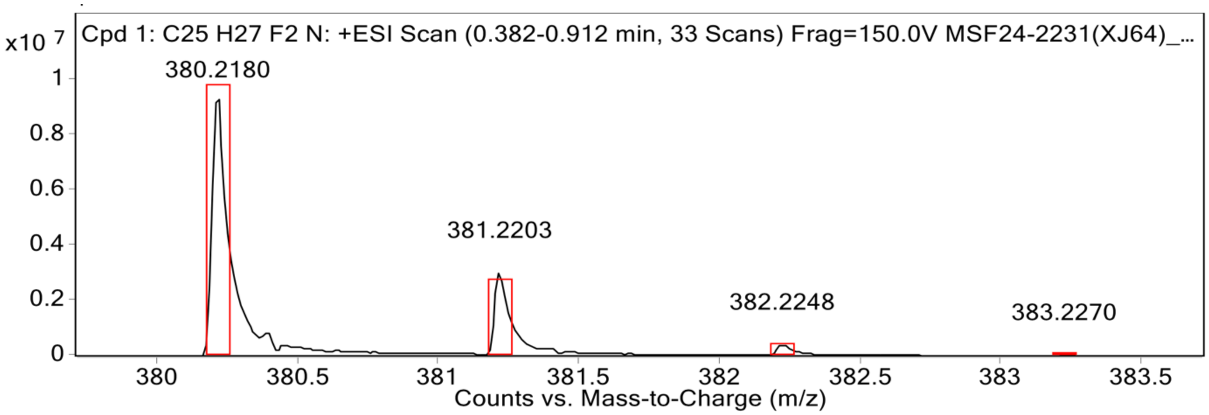


**MS Spectrum Peak List**

| Obs. m/z | Calc. m/z | Charge | Abundance | Formula | Ion Species | Tgt Mass Error (ppm) |
|---|---|---|---|---|---|---|
| 380.2180 | 380.2184 | 1 | 9465681 | C25H27F2N | (M+H)+ | 1.17 |
| 381.2203 | 381.2217 | 1 | 3017255 | C25H27F2N | (M+H)+ | 3.72 |
| 382.2248 | 382.2250 | 1 | 410629 | C25H27F2N | (M+H)+ | 0.66 |
| 383.2270 | 383.2283 | 1 | 33586 | C25H27F2N | (M+H)+ | 3.61 |

#### $^{1}$H NMR (800 MHz, Acetone-$d_6$) spectrum of XJ-3-69

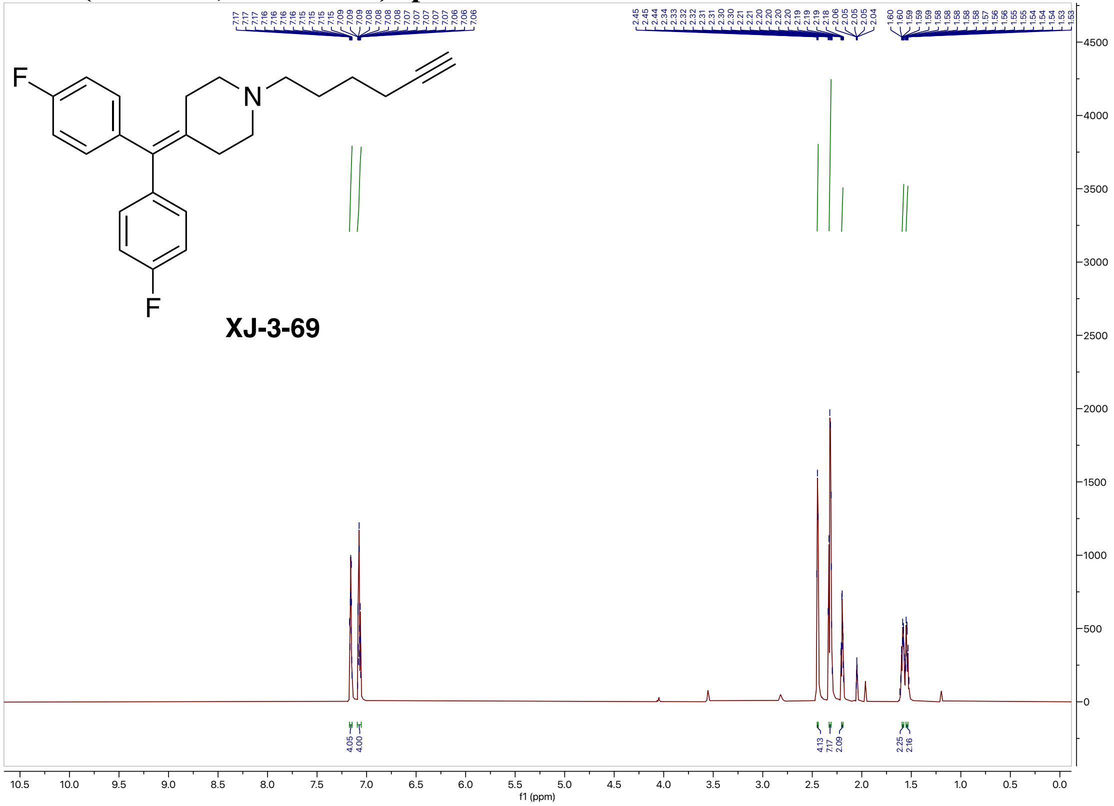

#### $^{13}$C NMR (200 MHz, Acetone-$d_6$) spectrum of XJ-3-69

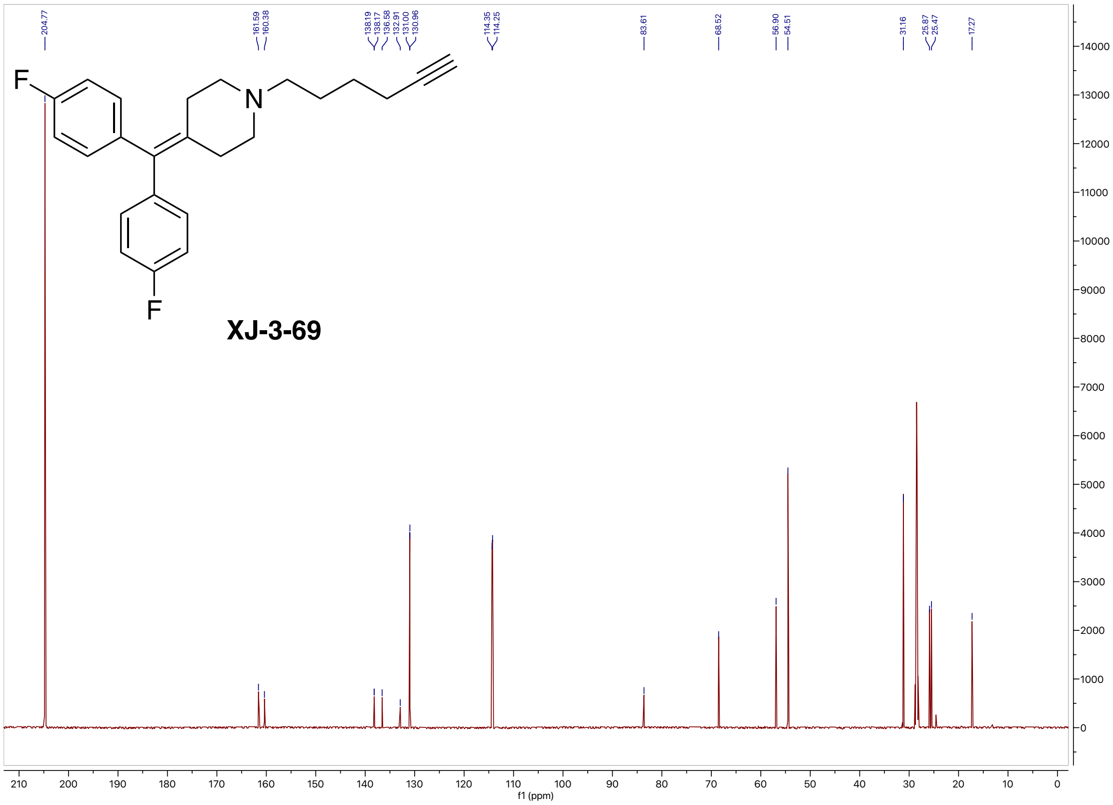

#### $^{19}$F NMR (564 MHz, Acetone-$d_6$) spectrum of XJ-3-69

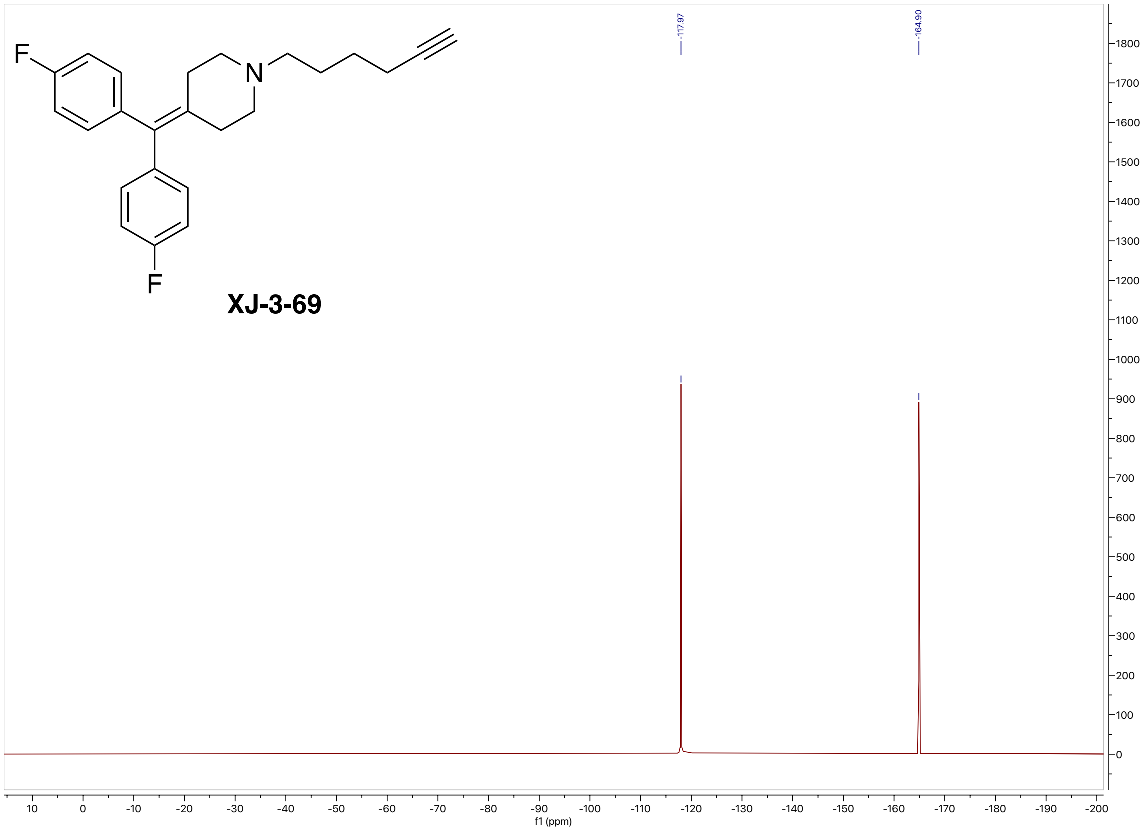


#### High-resolution mass spectrometry of XJ-3-69

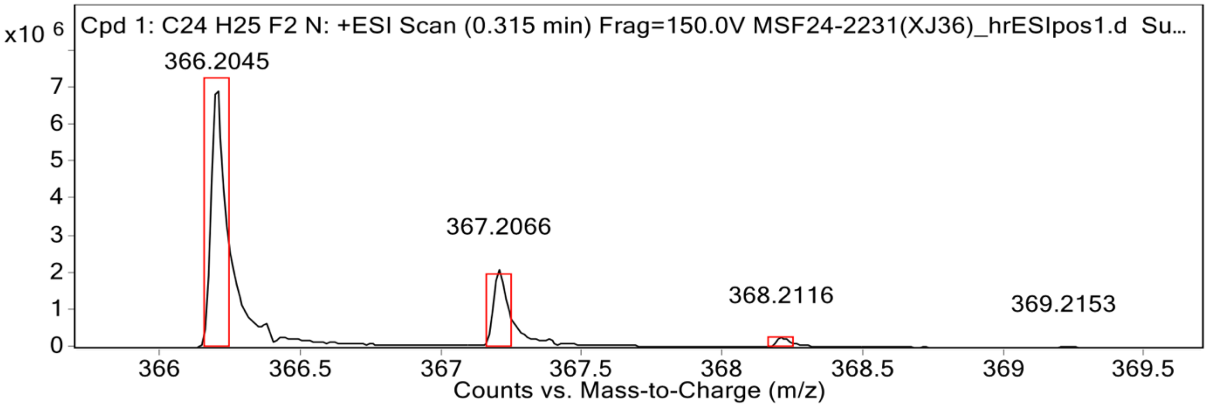


**MS Spectrum Peak List**

| Obs. m/z | Calc. m/z | Charge | Abundance | Formula | Ion Species | Tgt Mass Error (ppm) |
|---|---|---|---|---|---|---|
| 366.2045 | 366.2028 | 1 | 7048315 | C24H25F2N | (M+H)+ | -4.64 |
| 367.2066 | 367.2061 | 1 | 2114487 | C24H25F2N | (M+H)+ | -1.36 |
| 368.2116 | 368.2094 | 1 | 260065 | C24H25F2N | (M+H)+ | -5.98 |
| 369.2153 | 369.2127 | 1 | 26172 | C24H25F2N | (M+H)+ | -7.1 |

#### $^{1}$H NMR (500 MHz, Acetone-$d_6$) spectrum of XJ-4-91

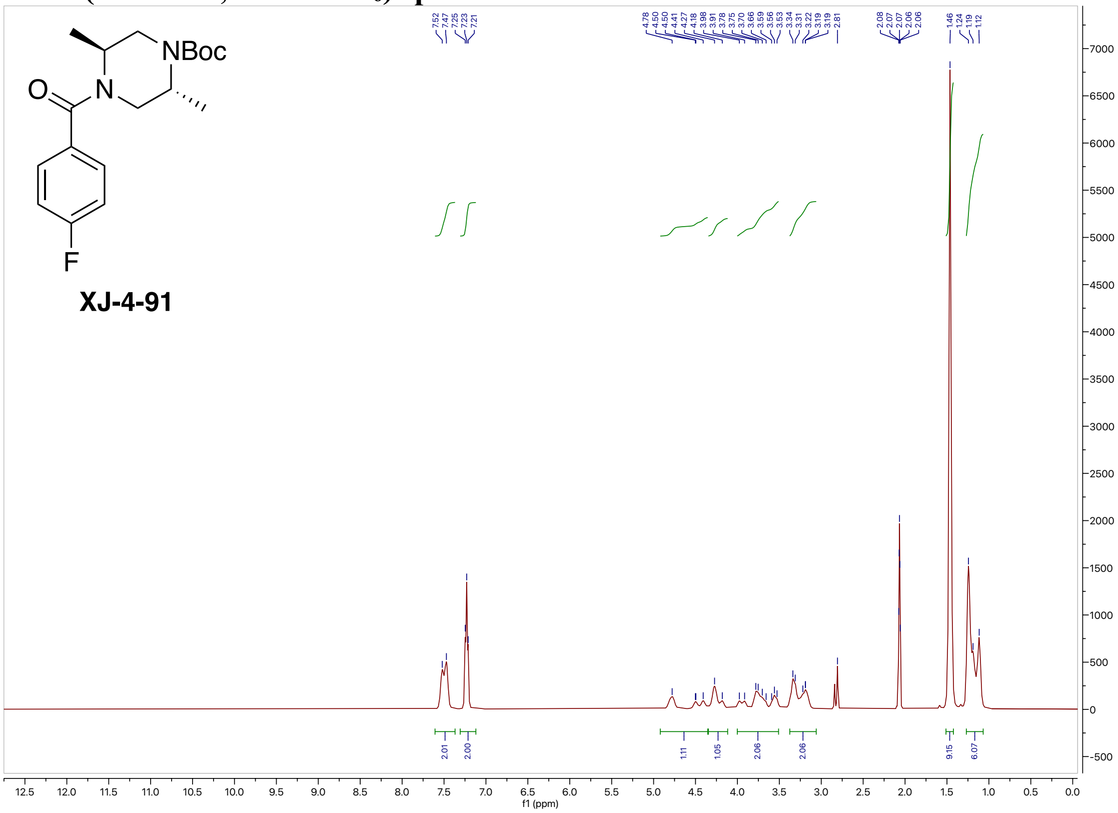

#### $^{13}$C NMR (200 MHz, Acetone-$d_6$) spectrum of XJ-4-91

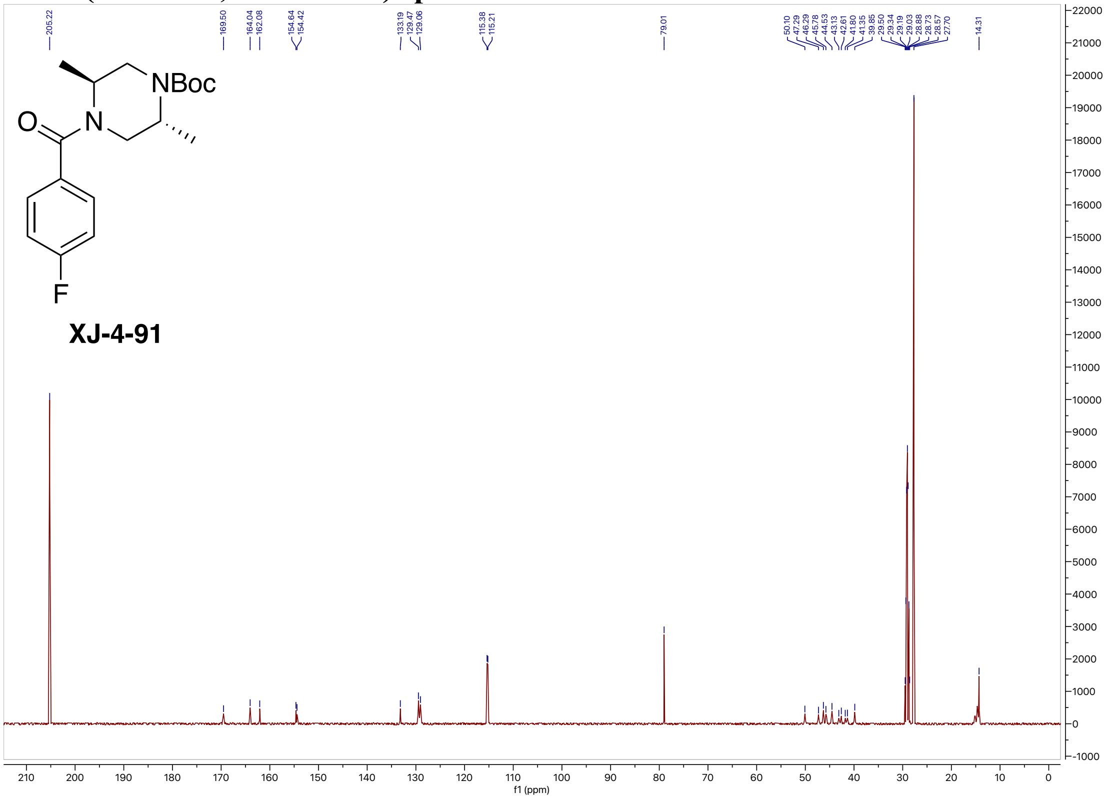

#### $^{19}$F NMR (471 MHz, Acetone-$d_6$) spectrum of XJ-4-91

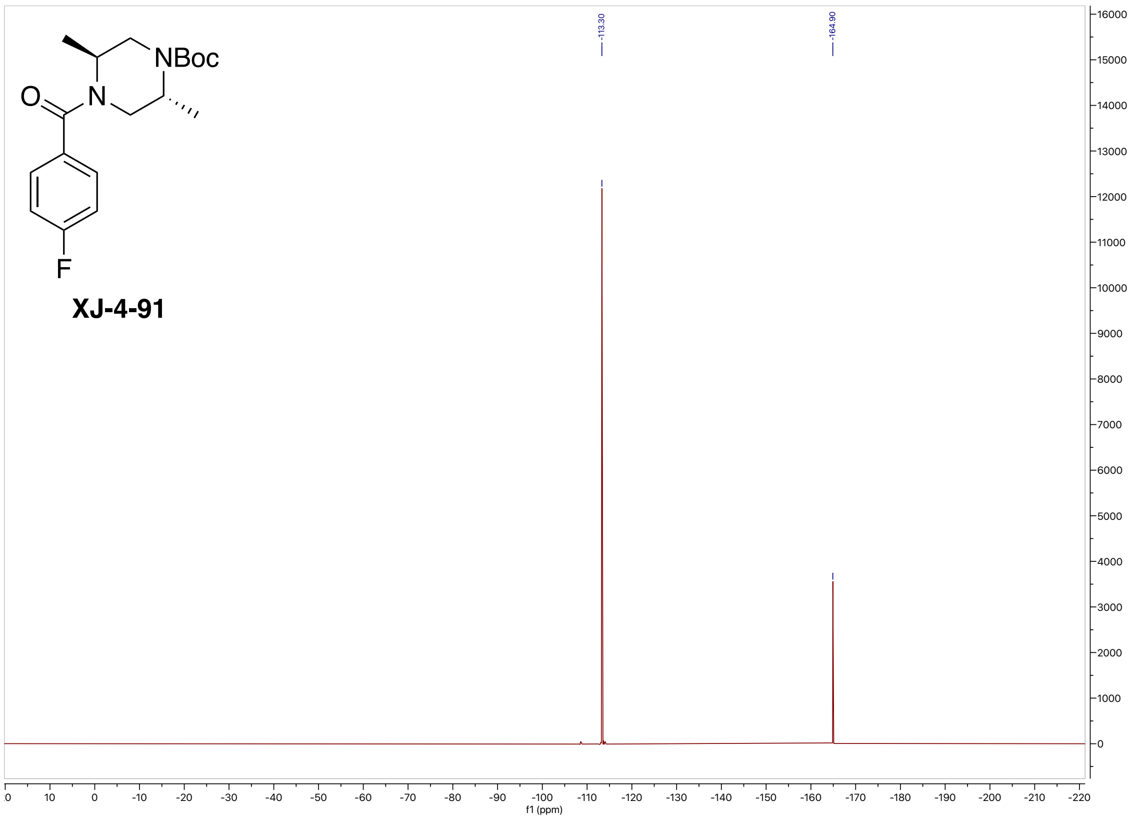


#### High-resolution mass spectrometry of XJ-4-91

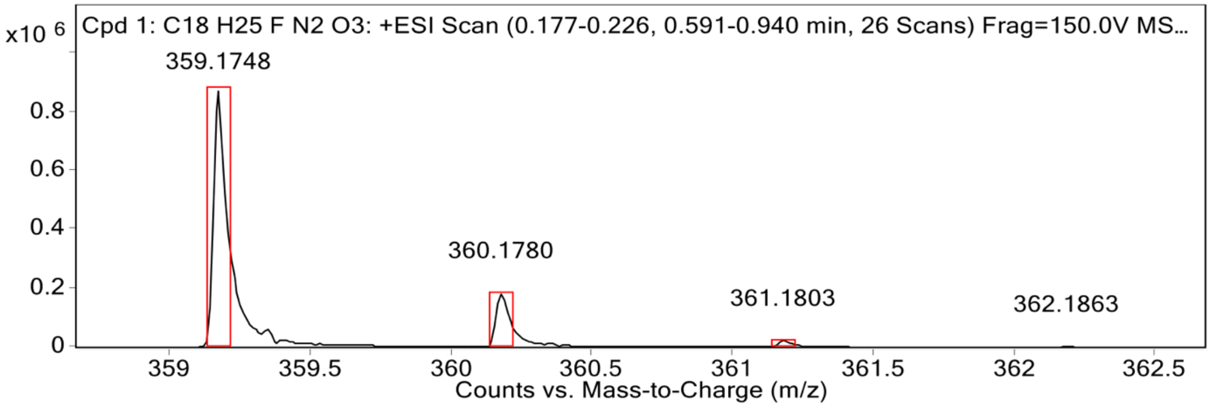


**MS Spectrum Peak List**

| Obs. m/z | Calc. m/z | Charge | Abundance | Formula | Ion Species | Tgt Mass Error (ppm) |
|---|---|---|---|---|---|---|
| 359.1748 | 359.1741 | 1 | 876723 | C18H25FN2O3 | (M+Na)+ | -1.79 |
| 360.1780 | 360.1773 | 1 | 184901 | C18H25FN2O3 | (M+Na)+ | -1.76 |
| 361.1803 | 361.1800 | 1 | 22931 | C18H25FN2O3 | (M+Na)+ | -0.98 |
| 362.1863 | 362.1826 | 1 | 2573 | C18H25FN2O3 | (M+Na)+ | -10.11 |
| 418.2017 | | | 7927745 | | | |

#### $^{1}$H NMR (500 MHz, Acetone-$d_6$) spectrum of XJ-4-92

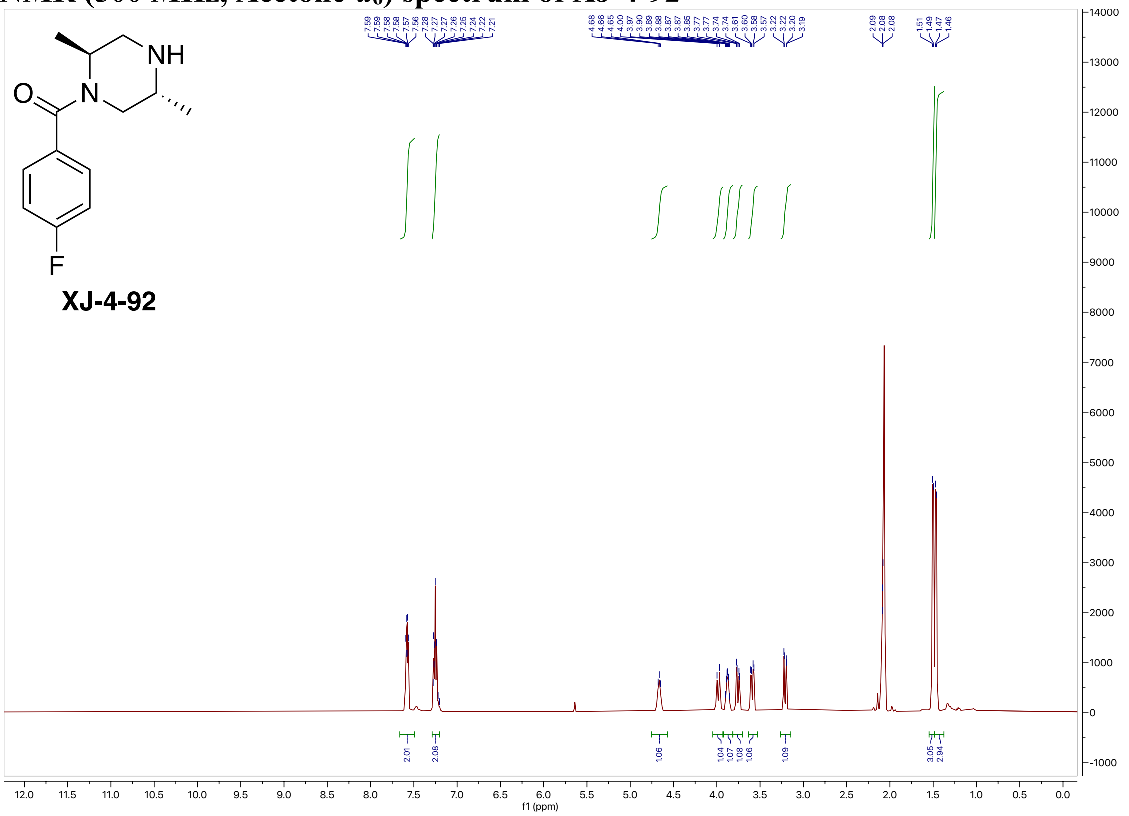


#### $^{13}$C NMR (200 MHz, Acetone-$d_6$) spectrum of XJ-4-92

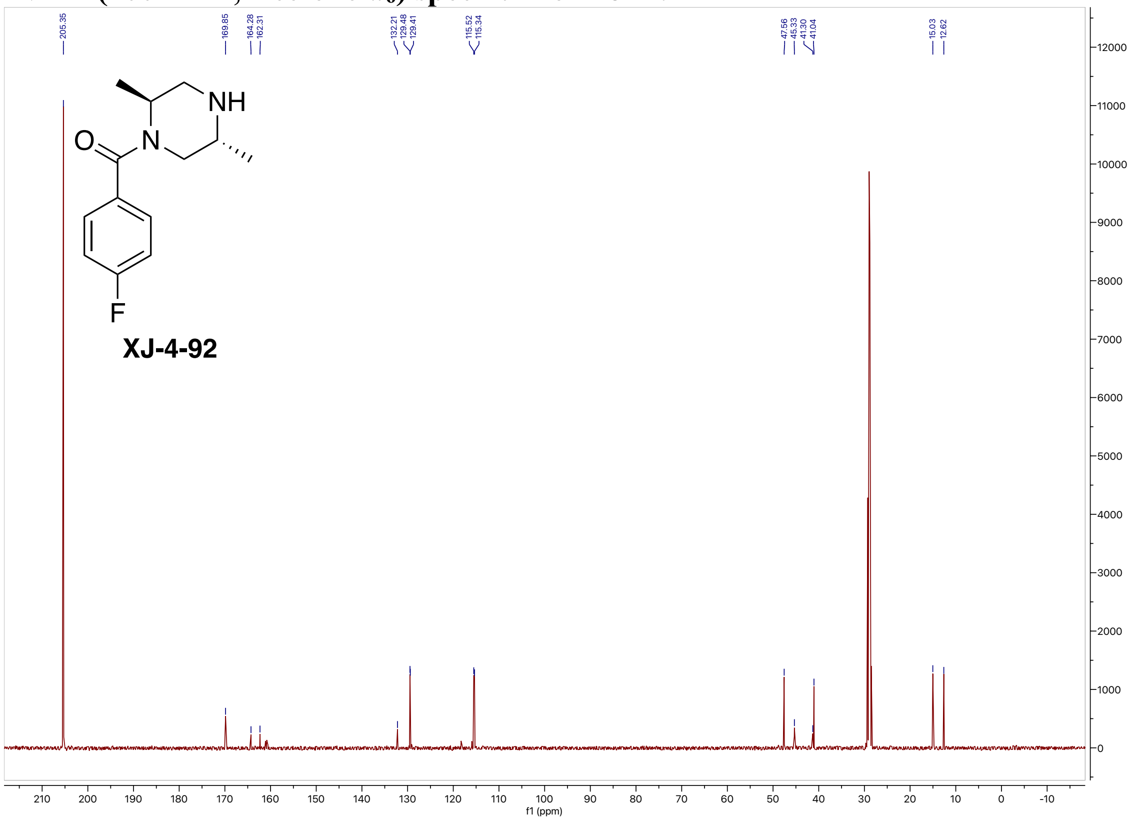

#### $^{19}$F NMR (471 MHz, Acetone-$d_6$) spectrum of XJ-4-92

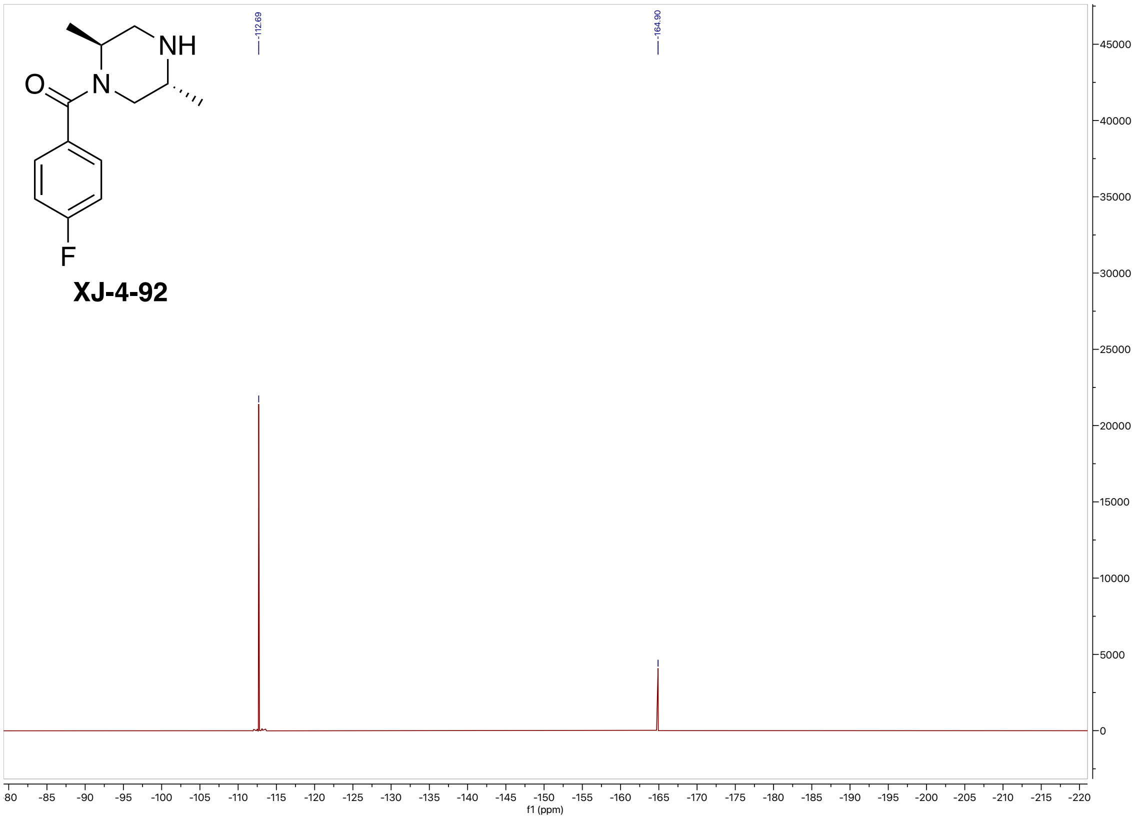


#### High-resolution mass spectrometry of XJ-4-92

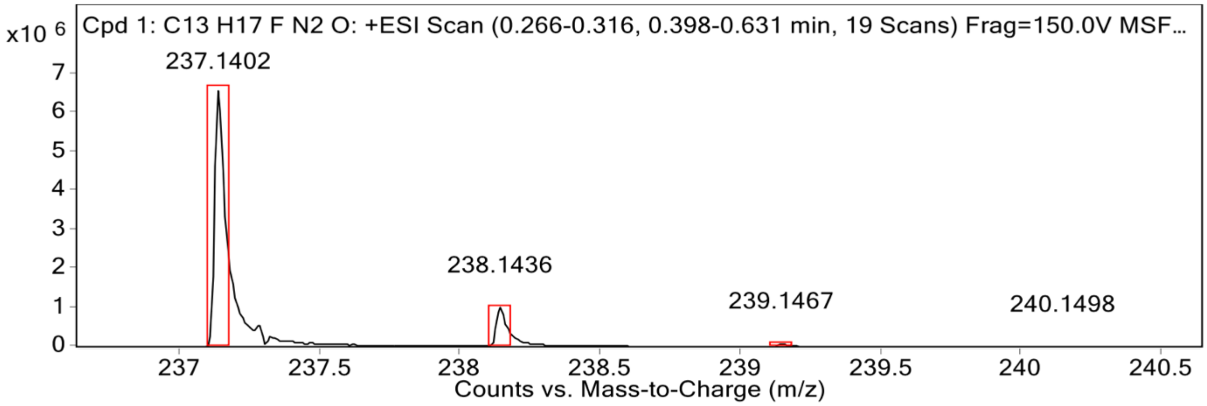


**MS Spectrum Peak List**

| Obs. m/z | Calc. m/z | Charge | Abundance | Formula | Ion Species | Tgt Mass Error (ppm) |
|---|---|---|---|---|---|---|
| 237.1402 | 237.1398 | 1 | 6674957 | C13H17FN2O | (M+H)+ | -1.78 |
| 238.1436 | 238.1429 | 1 | 1010224 | C13H17FN2O | (M+H)+ | -3.11 |
| 239.1467 | 239.1456 | 1 | 86052 | C13H17FN2O | (M+H)+ | -4.37 |
| 240.1498 | 240.1482 | 1 | 5026 | C13H17FN2O | (M+H)+ | -6.81 |

#### $^{1}$H NMR (500 MHz, Acetone-$d_6$) spectrum of XJ-4-95

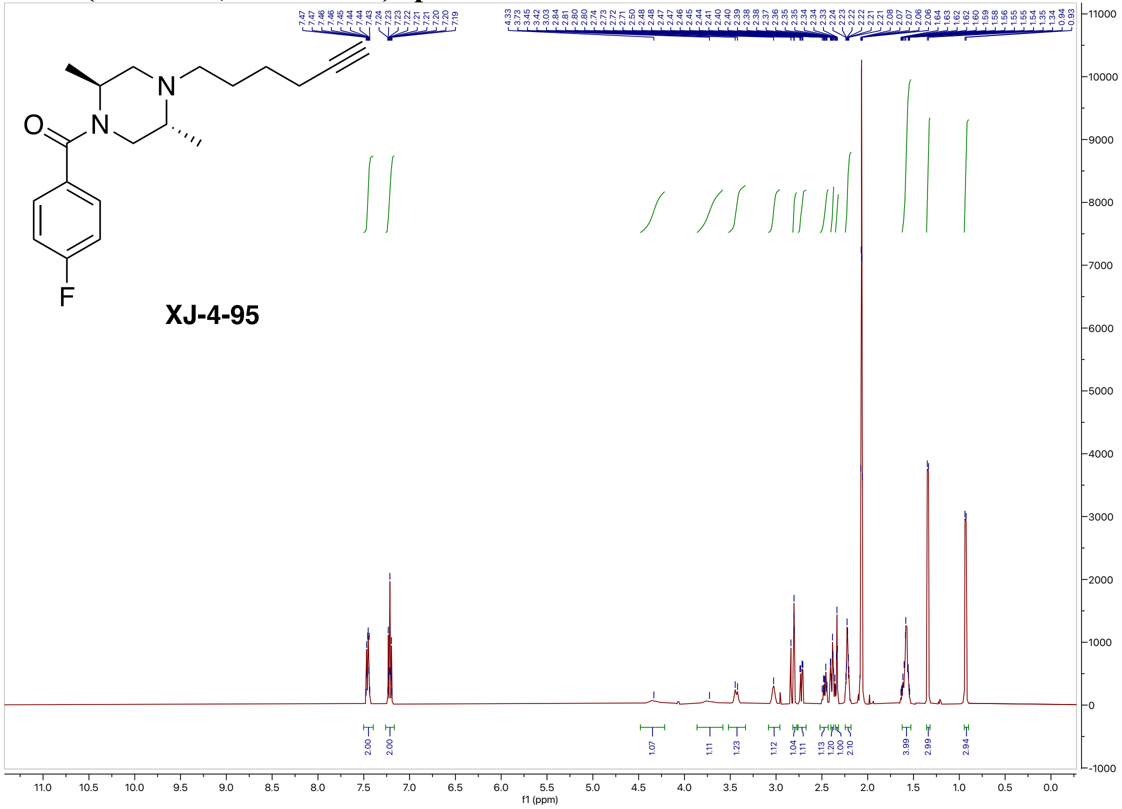


#### $^{13}$C NMR (200 MHz, Acetone-$d_6$) spectrum of XJ-4-95

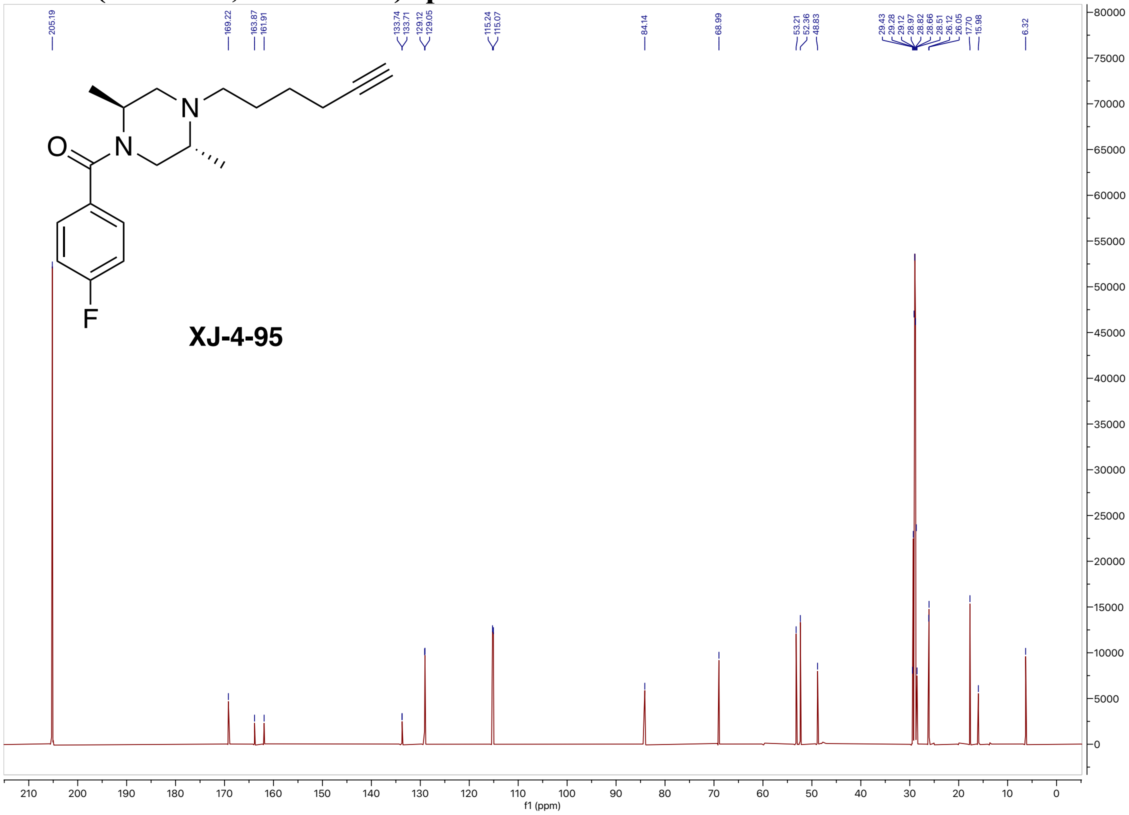

#### $^{19}$F NMR (471 MHz, Acetone-$d_6$) spectrum of XJ-4-95

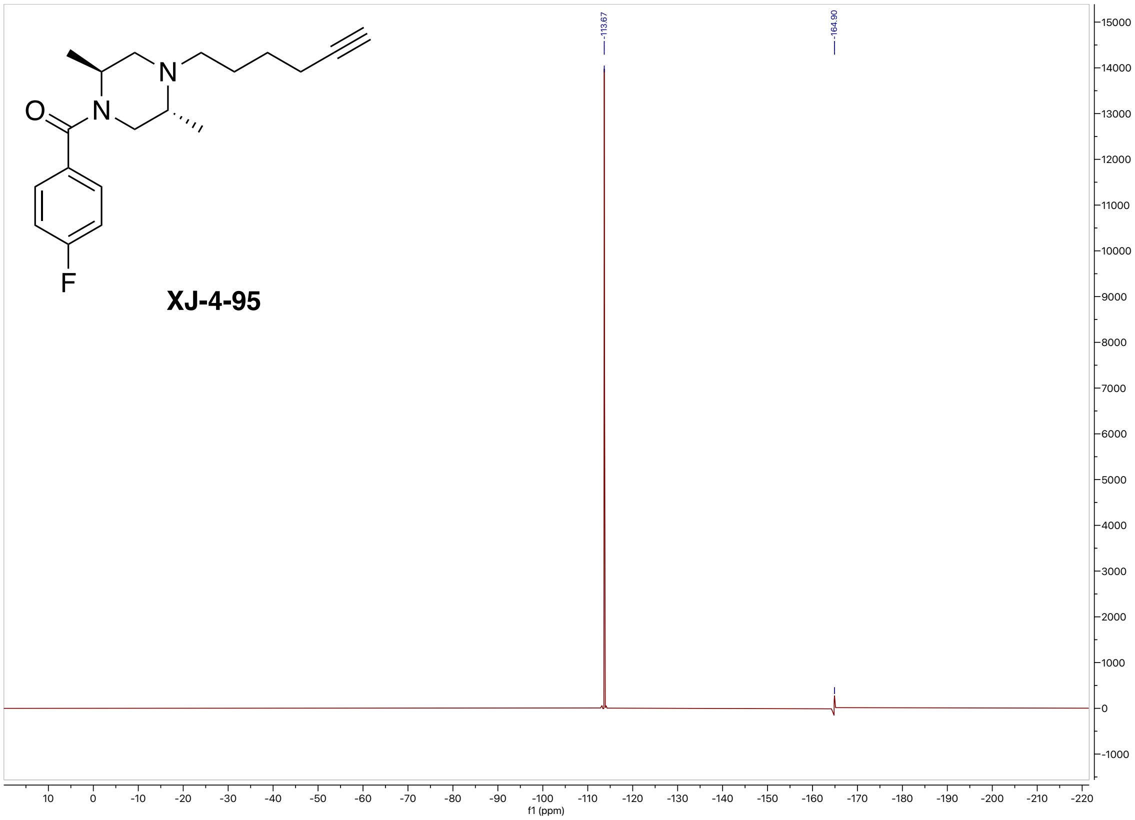


#### High-resolution mass spectrometry of XJ-4-95

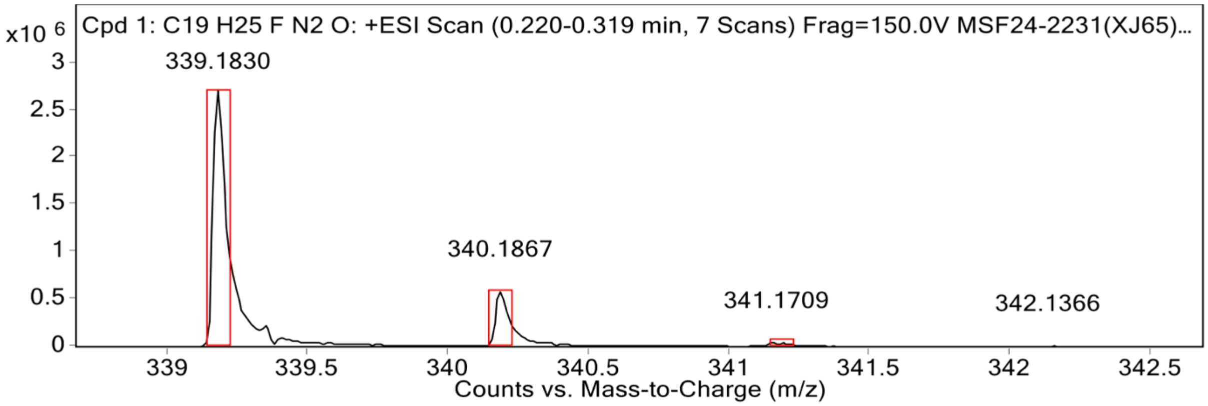


**MS Spectrum Peak List**

| Obs. m/z | Calc. m/z | Charge | Abundance | Formula | Ion Species | Tgt Mass Error (ppm) |
|---|---|---|---|---|---|---|
| 339.1830 | 339.1843 | 1 | 2706115 | C19H25FN2O | (M+Na)+ | 3.87 |
| 340.1867 | 340.1875 | 1 | 584158 | C19H25FN2O | (M+Na)+ | 2.22 |
| 341.1709 | 341.1905 | 1 | 40580 | C19H25FN2O | (M+Na)+ | 57.31 |
| 342.1366 | 342.1933 | 1 | 9354 | C19H25FN2O | (M+Na)+ | 165.87 |
| 343.1334 | 343.1961 | 1 | 8122 | C19H25FN2O | (M+Na)+ | 182.87 |

**$^{1}$H NMR (500 MHz, Acetone-$d_6$) spectrum of XJ-4-43**

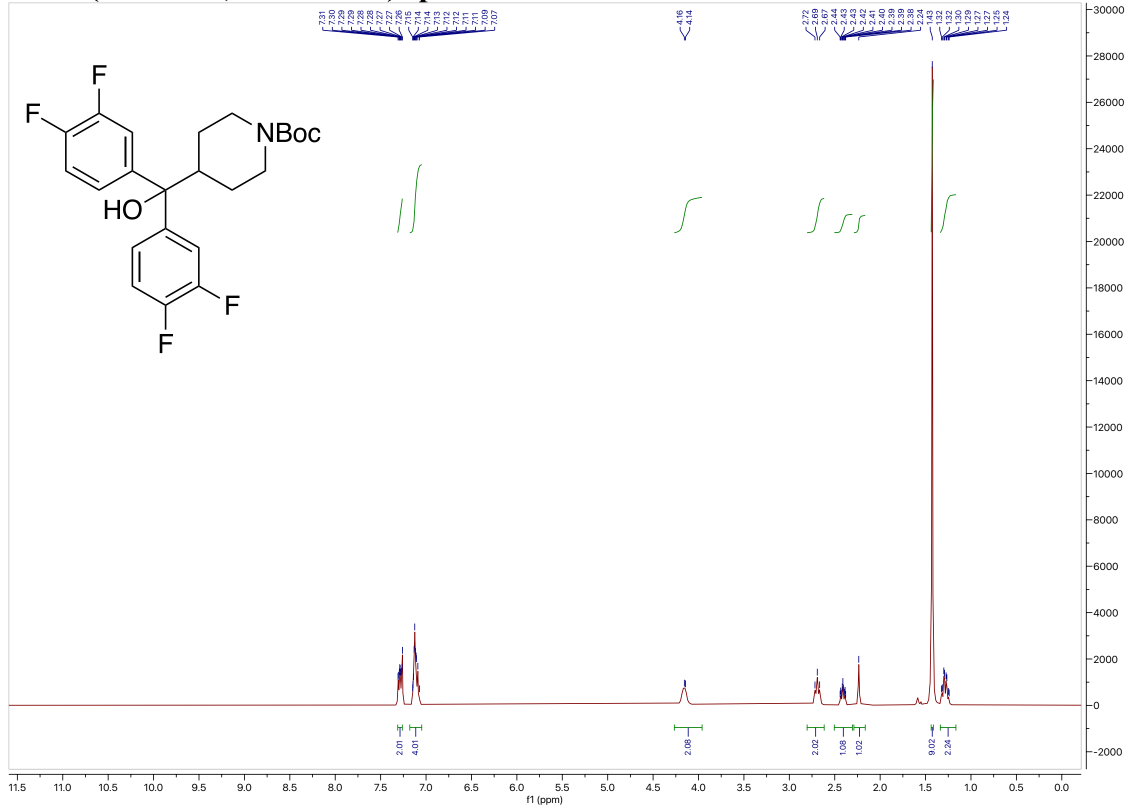

**$^{13}$C NMR (200 MHz, Acetone-$d_6$) spectrum of XJ-4-43**

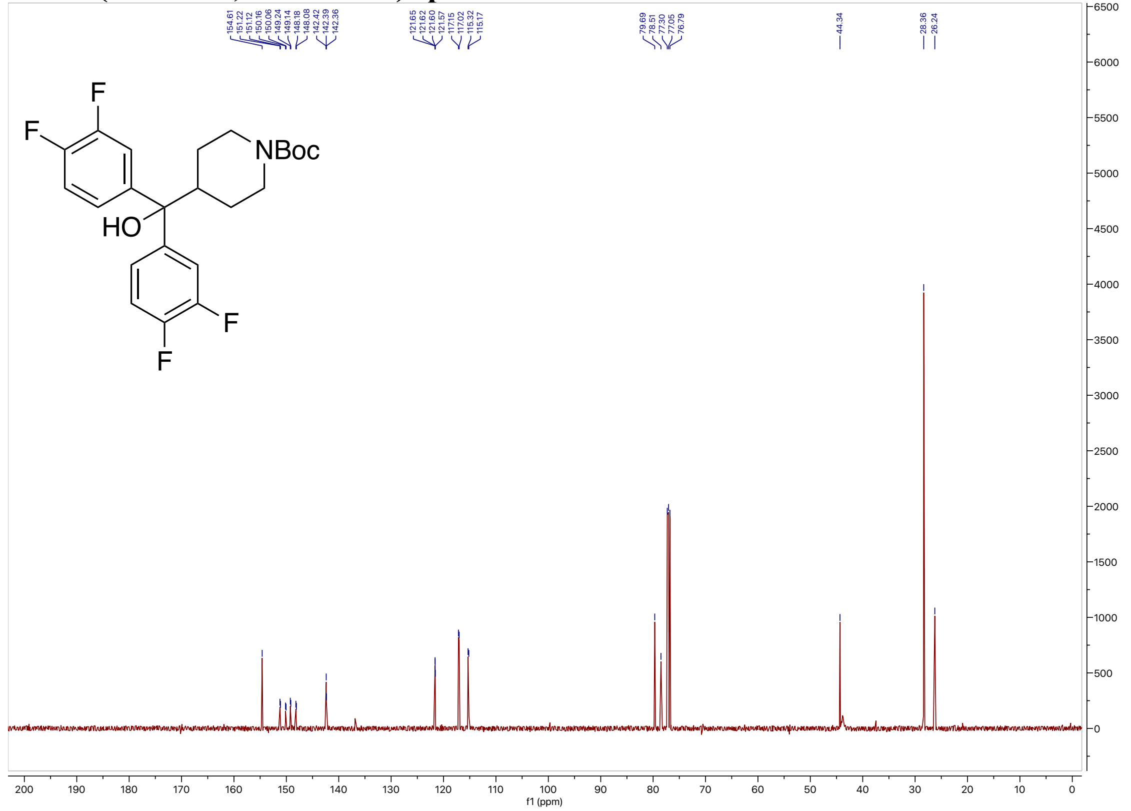

#### $^{19}$F NMR (471 MHz, Acetone-$d_6$) spectrum of XJ-4-43

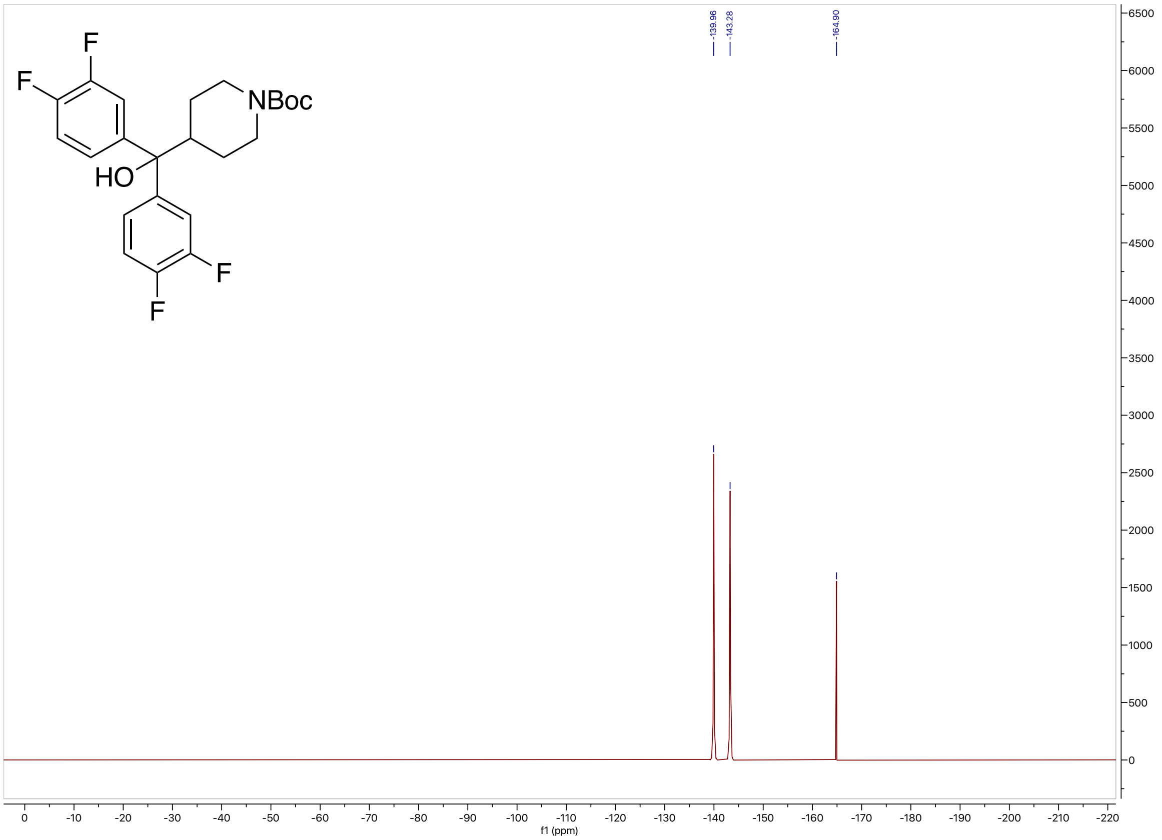


#### High-resolution mass spectrometry of XJ-4-43

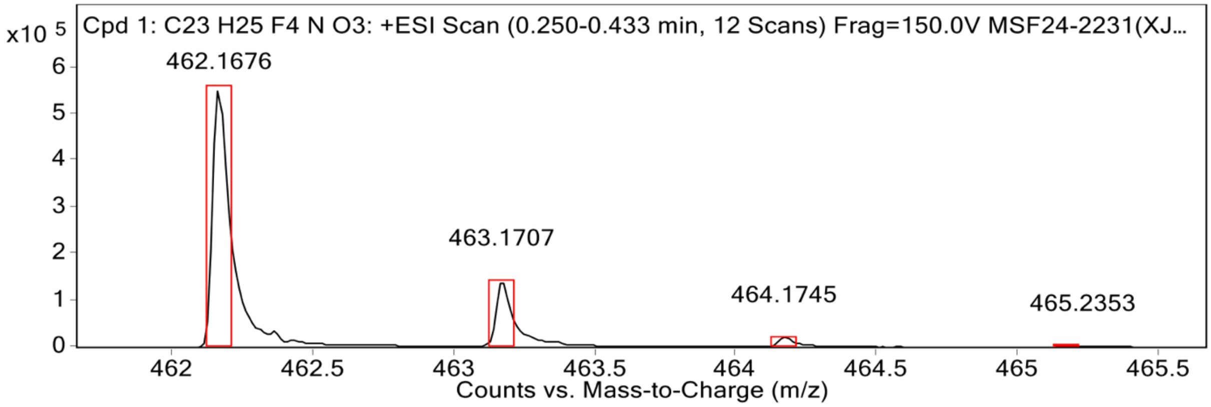


**MS Spectrum Peak List**

| Obs. m/z | Calc. m/z | Charge | Abundance | Formula | Ion Species | Tgt Mass Error (ppm) |
|---|---|---|---|---|---|---|
| 462.1676 | 462.1663 | 1 | 553367 | C23H25F4NO3 | (M+Na)+ | -2.97 |
| 463.1707 | 463.1696 | 1 | 143807 | C23H25F4NO3 | (M+Na)+ | -2.4 |
| 464.1745 | 464.1725 | 1 | 22014 | C23H25F4NO3 | (M+Na)+ | -4.27 |
| 465.2353 | 465.1753 | 1 | 3464 | C23H25F4NO3 | (M+Na)+ | -129.06 |
| 466.2093 | 466.1780 | 1 | 1912 | C23H25F4NO3 | (M+Na)+ | -67.14 |

#### $^{1}$H NMR (500 MHz, Acetone-$d_6$) spectrum of XJ-4-45

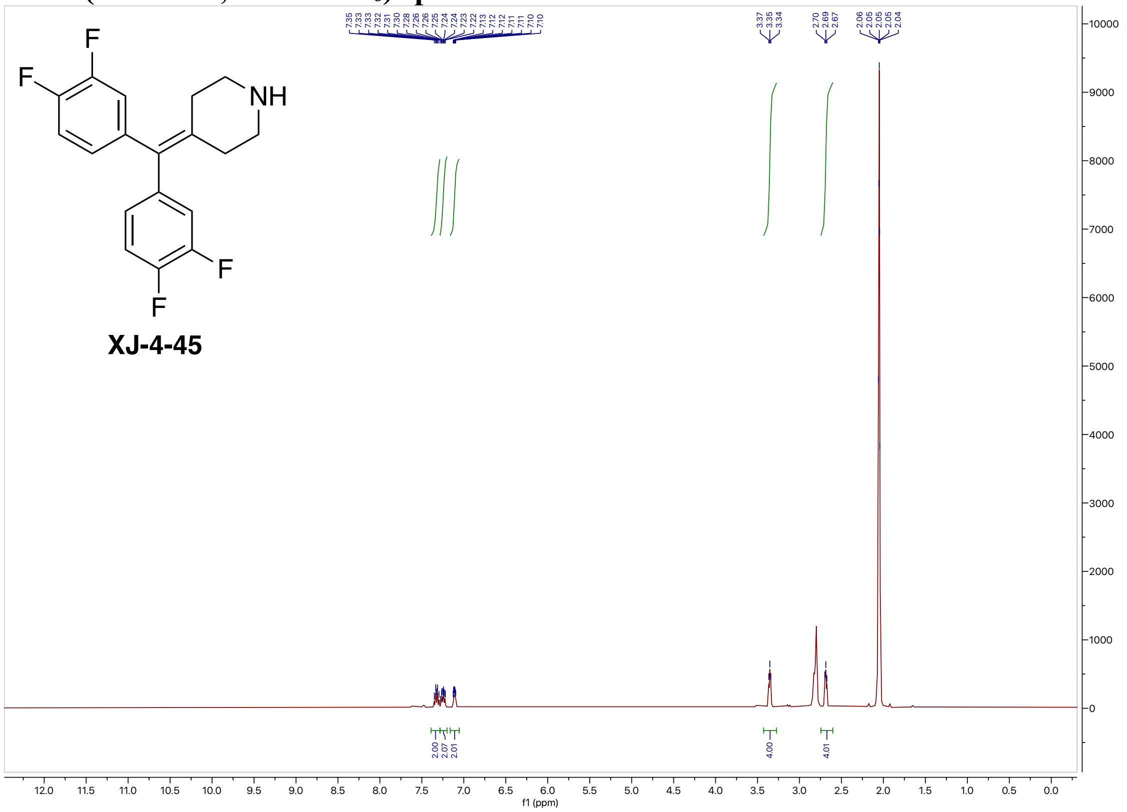


#### $^{13}$C NMR (200 MHz, Acetone-$d_6$) spectrum of XJ-4-45

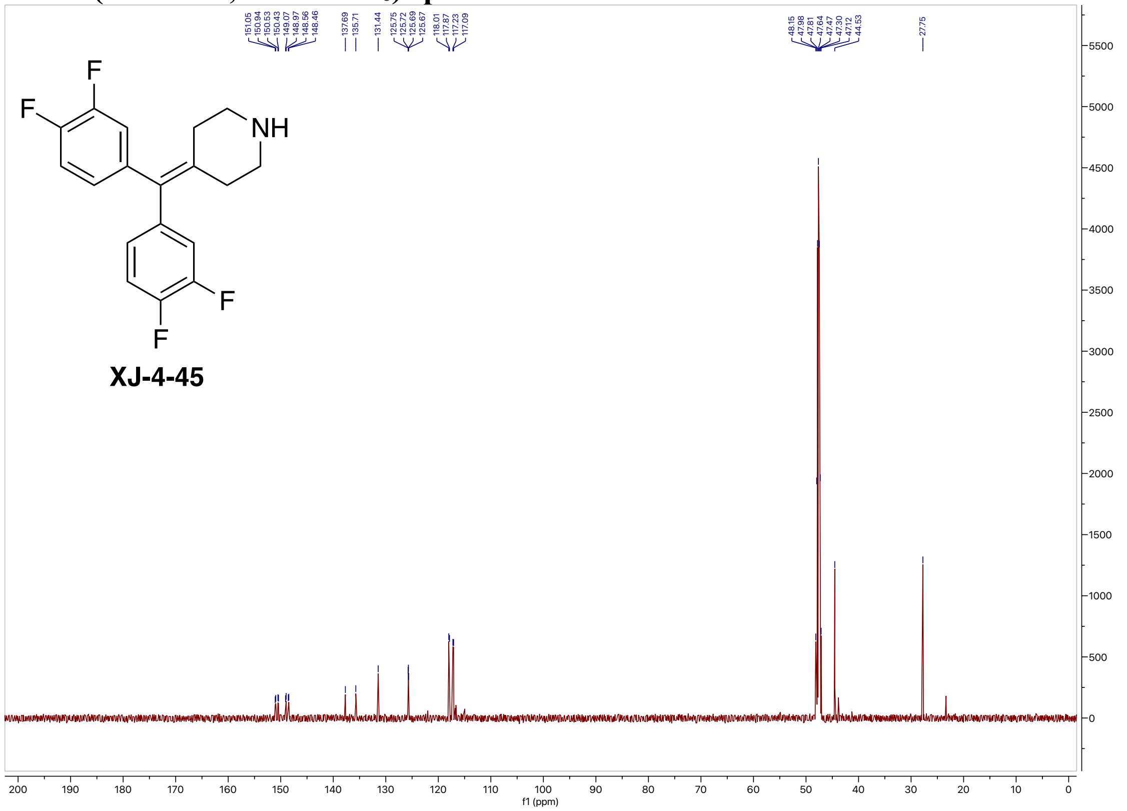

#### $^{19}$F NMR (471 MHz, Acetone-$d_6$) spectrum of XJ-4-45

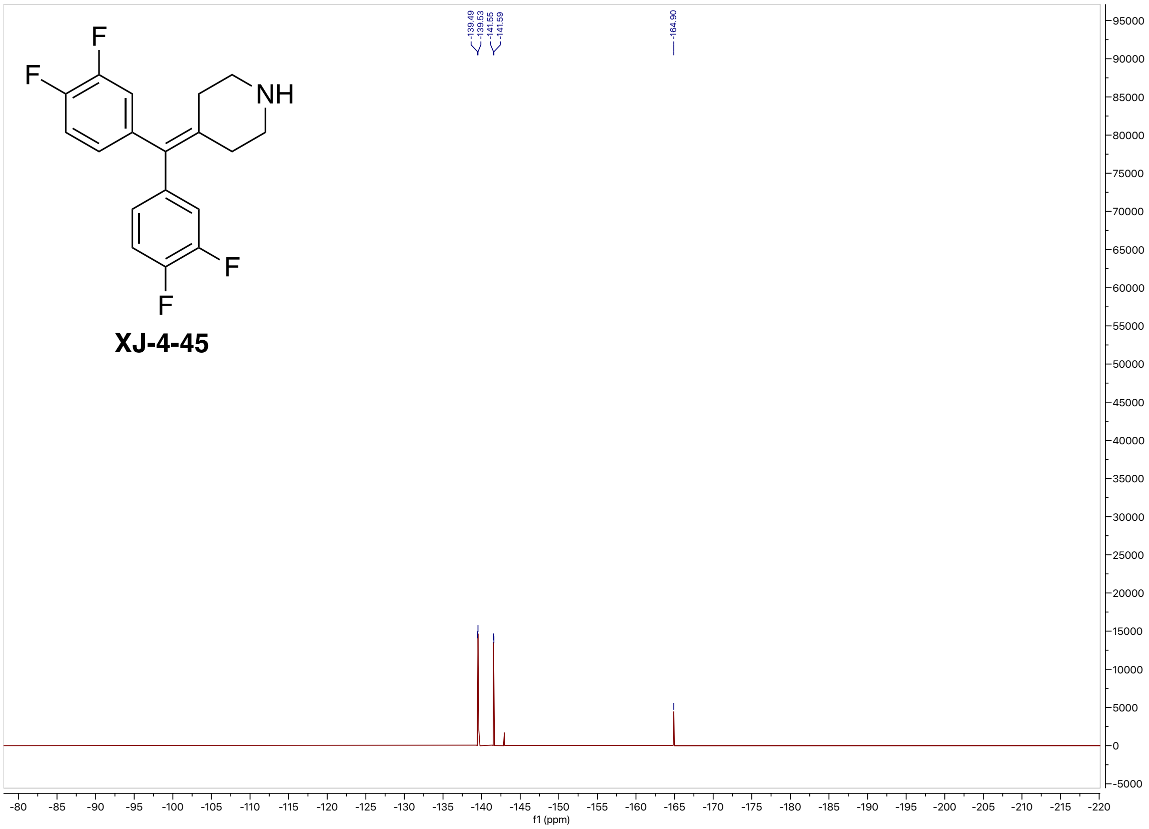


#### High-resolution mass spectrometry of XJ-4-45

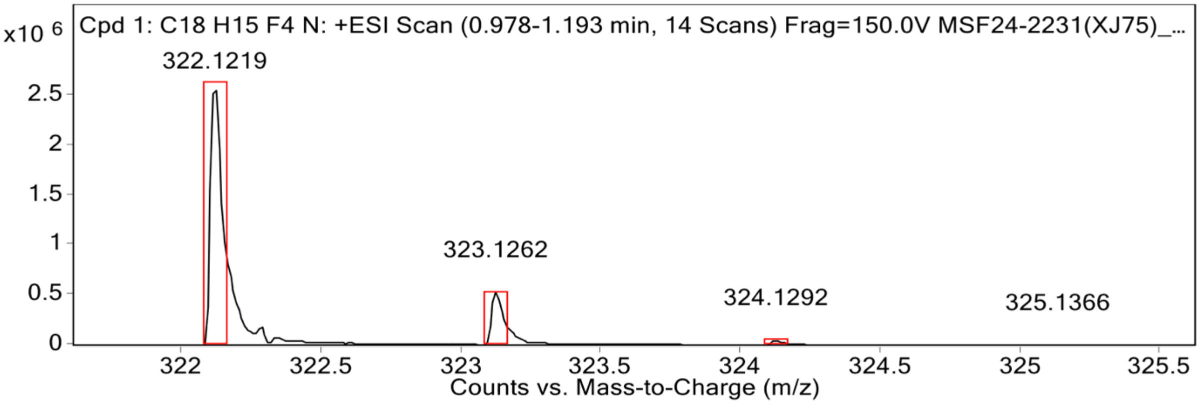


**MS Spectrum Peak List**

| Obs. m/z | Calc. m/z | Charge | Abundance | Formula | Ion Species | Tgt Mass Error (ppm) |
|---|---|---|---|---|---|---|
| 322.1219 | 322.1213 | 1 | 2609764 | C18H15F4N | (M+H)+ | -1.61 |
| 323.1262 | 323.1246 | 1 | 531722 | C18H15F4N | (M+H)+ | -4.82 |
| 324.1292 | 324.1279 | 1 | 52932 | C18H15F4N | (M+H)+ | -4.04 |
| 325.1366 | 325.1311 | 1 | 2710 | C18H15F4N | (M+H)+ | -16.79 |

#### $^{1}$H NMR (500 MHz, Acetone-$d_6$) spectrum of XJ-4-49

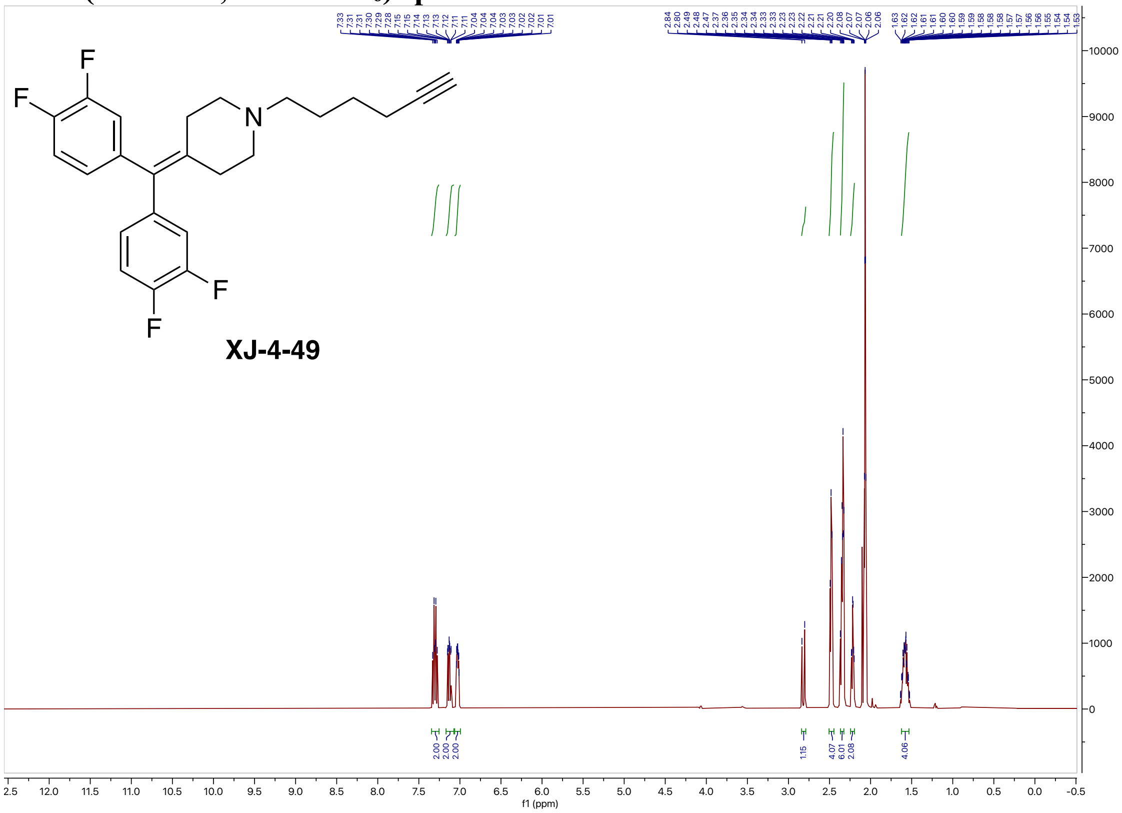

#### $^{13}$C NMR (200 MHz, Acetone-$d_6$) spectrum of XJ-4-49

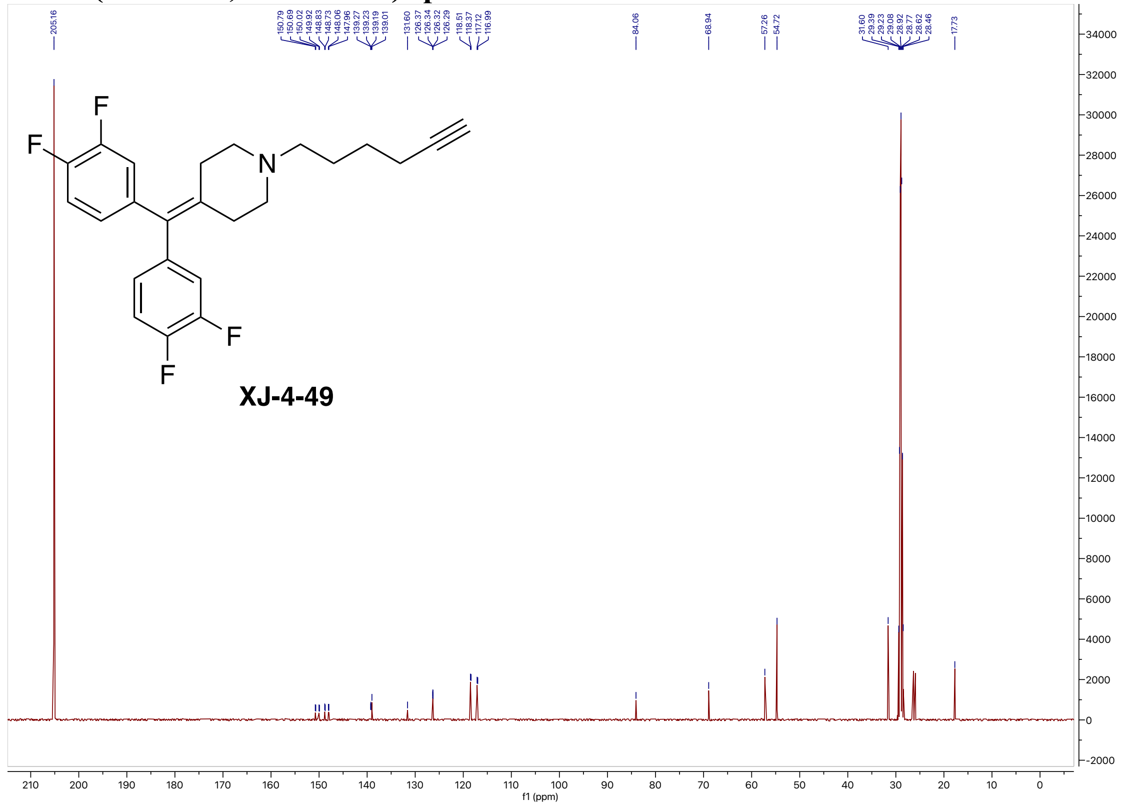

#### [19]F NMR (471 MHz, Acetone-$d_6$) spectrum of XJ-4-49

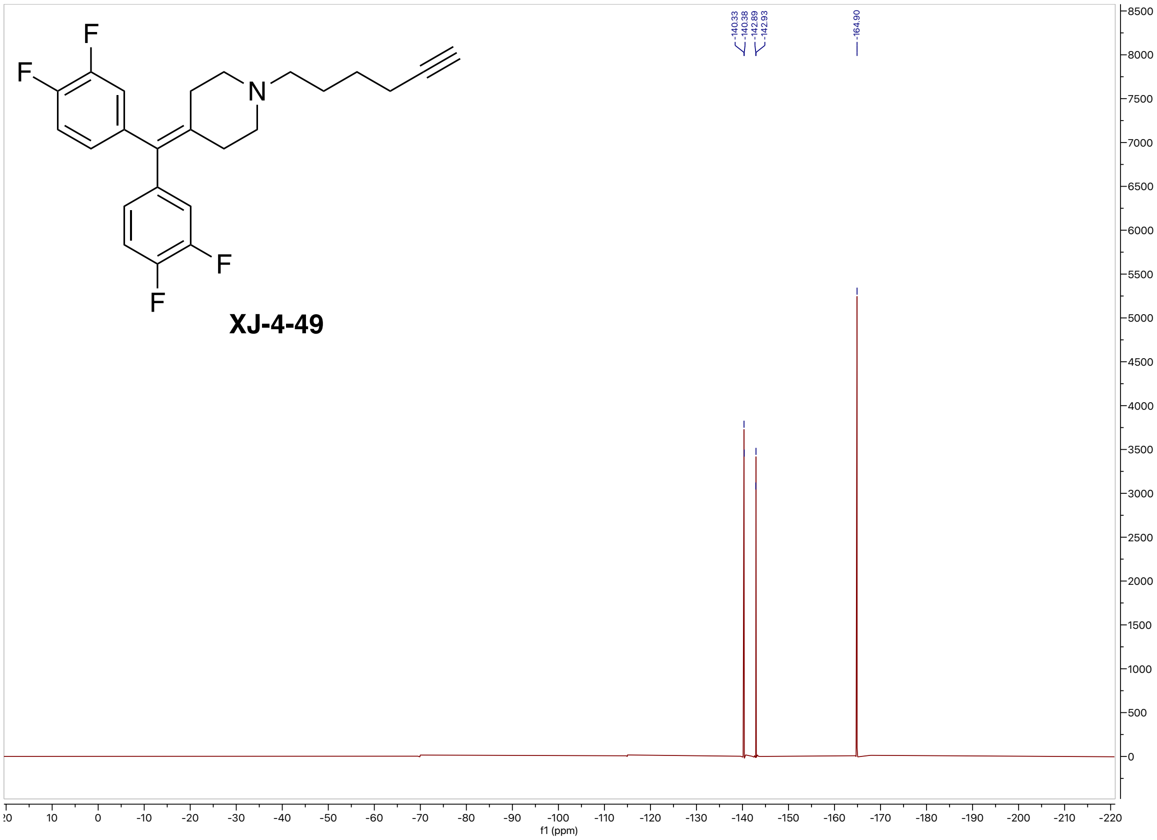


#### High-resolution mass spectrometry of XJ-4-49

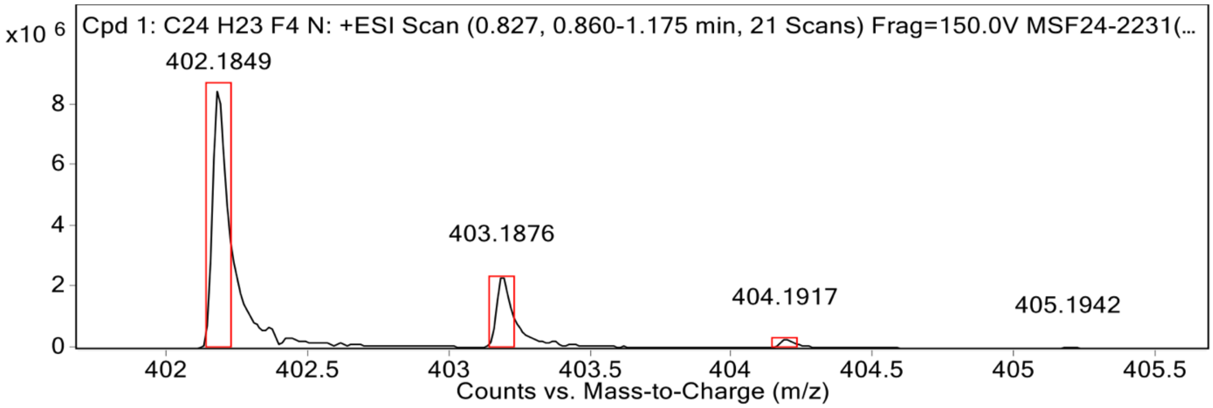


**MS Spectrum Peak List**

| Obs. m/z | Calc. m/z | Charge | Abundance | Formula | Ion Species | Tgt Mass Error (ppm) |
|---|---|---|---|---|---|---|
| 402.1849 | 402.1839 | 1 | 8608482 | C24H23F4N | (M+H)+ | -2.49 |
| 403.1876 | 403.1872 | 1 | 2370501 | C24H23F4N | (M+H)+ | -1 |
| 404.1917 | 404.1905 | 1 | 293250 | C24H23F4N | (M+H)+ | -2.82 |
| 405.1942 | 405.1938 | 1 | 22931 | C24H23F4N | (M+H)+ | -0.89 |

#### [1]H NMR (800 MHz, CDCl3) spectrum of TH207

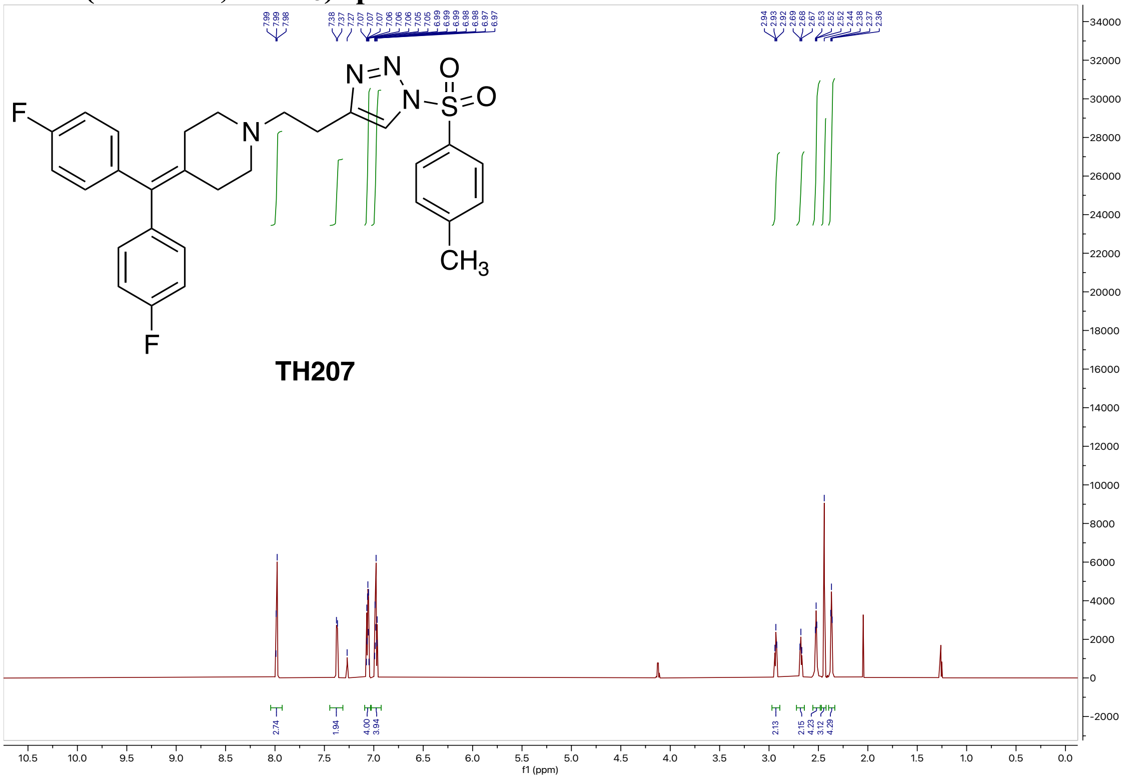

#### [13]C NMR (201 MHz, CDCl3) spectrum of TH207

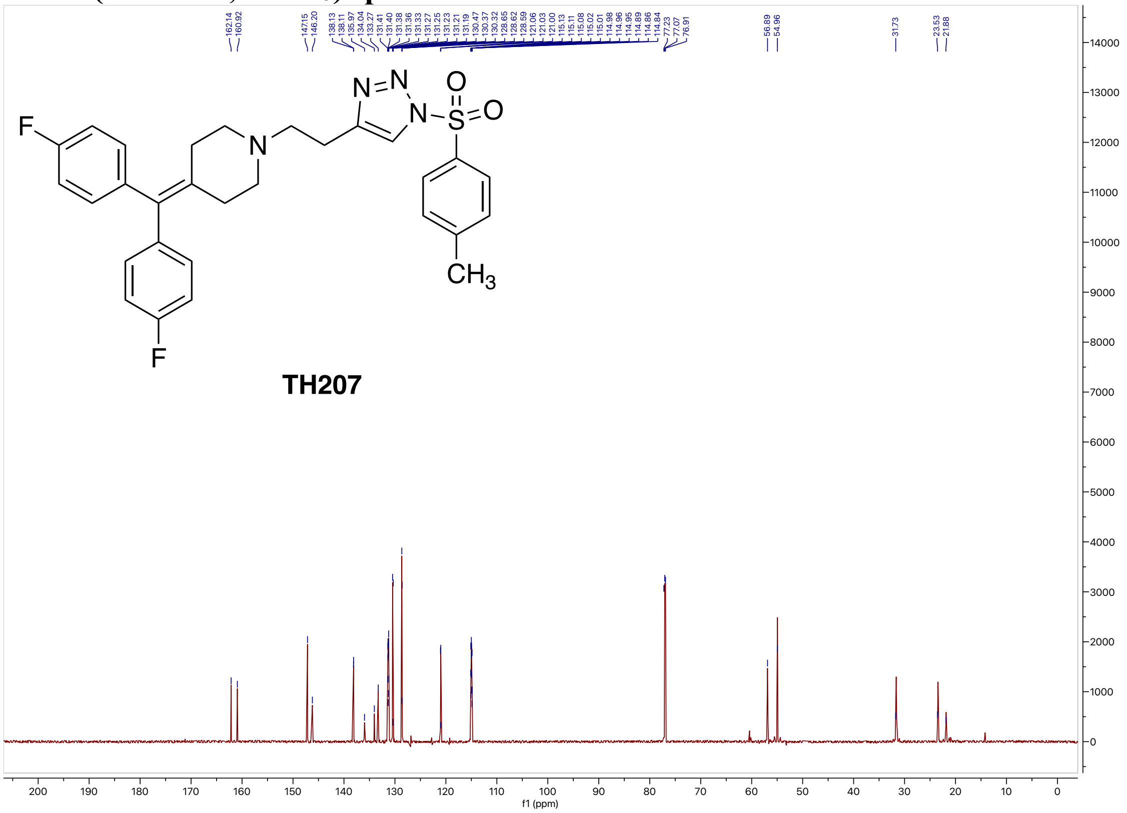

#### $^{19}F$ NMR (564 MHz, $CDCl_3$) spectrum of TH207

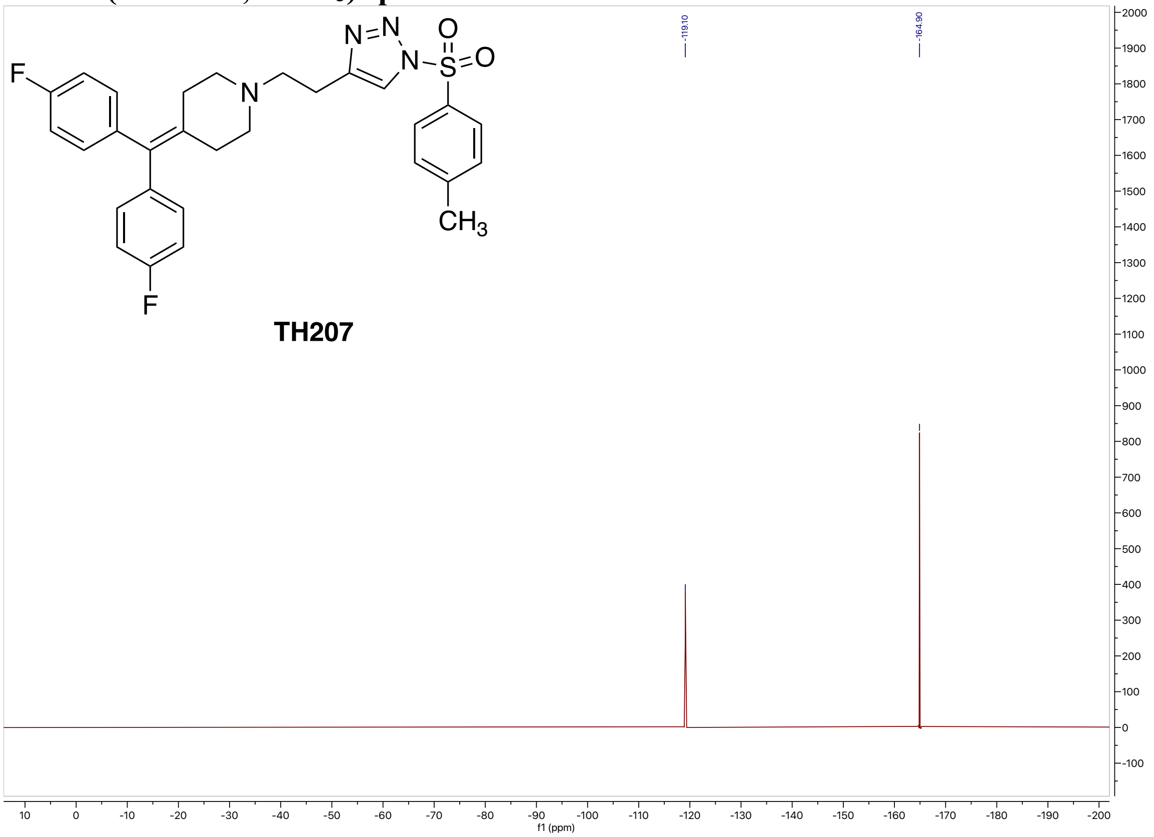


#### High-resolution mass spectrometry of TH207

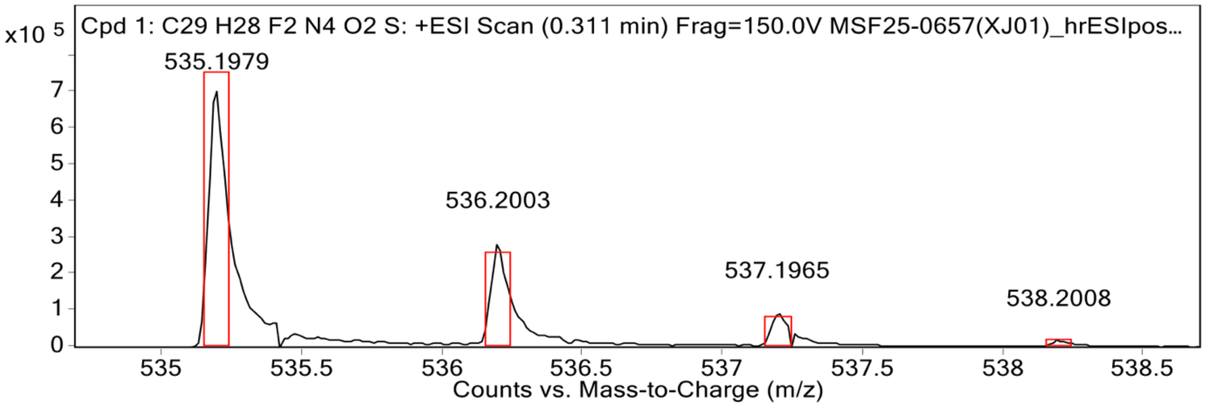


**MS Spectrum Peak List**

| Obs. m/z | Calc. m/z | Charge | Abundance | Formula | Ion Species | Tgt Mass Error (ppm) |
|---|---|---|---|---|---|---|
| 535.1979 | 535.1974 | 1 | 707520 | C29H28F2N4O2S | (M+H)+ | -0.93 |
| 536.2003 | 536.2004 | 1 | 286207 | C29H28F2N4O2S | (M+H)+ | 0.15 |
| 537.1965 | 537.1990 | 1 | 93887 | C29H28F2N4O2S | (M+H)+ | 4.53 |
| 538.2008 | 538.1995 | 1 | 17384 | C29H28F2N4O2S | (M+H)+ | -2.37 |

#### HPLC spectrum of TH207

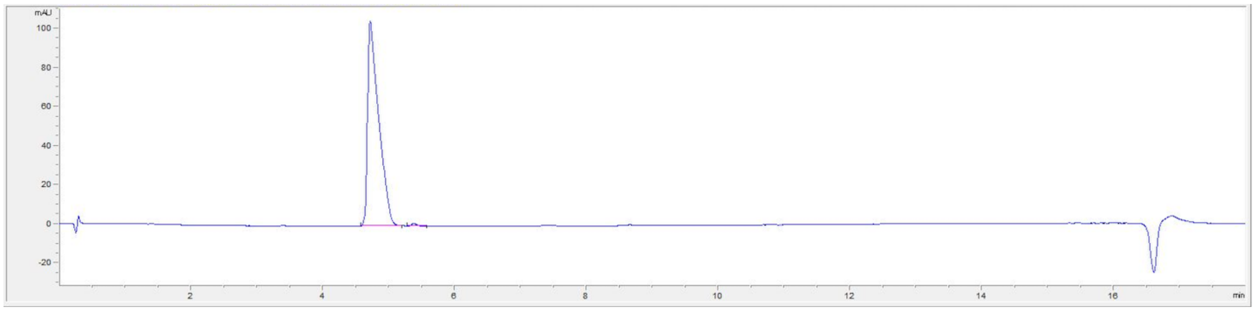

| # | Time | Area | Height | Width | Area% | Symmetry |
|---|---|---|---|---|---|---|
| 1 | 4.716 | 1224.7 | 105.3 | 0.1597 | 99.326 | 0.265 |
| 2 | 5.384 | 8.3 | 1.5 | 0.0846 | 0.674 | 0.817 |

#### $^1$H NMR (800 MHz, $CDCl_3$) spectrum of TH208

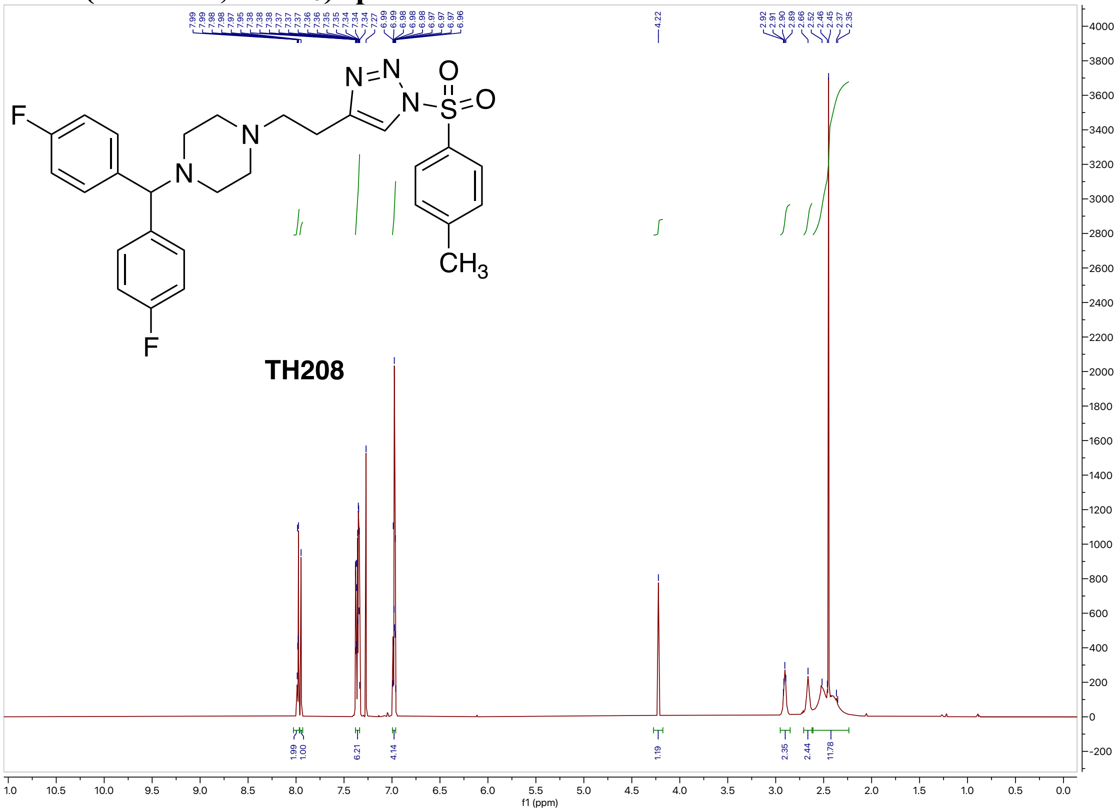

$^{13}$C NMR (200 MHz, $CDCl_3$) spectrum of TH208

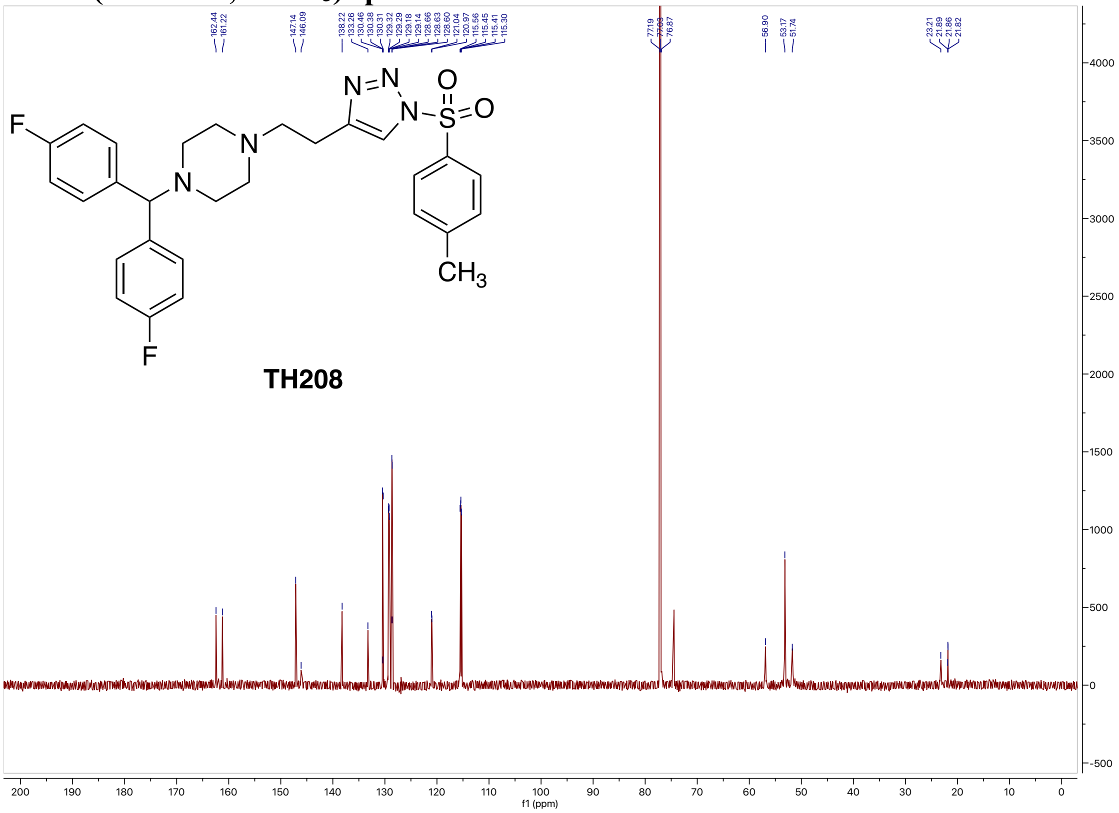


$^{19}$F NMR (564 MHz, $CDCl_3$) spectrum of TH208

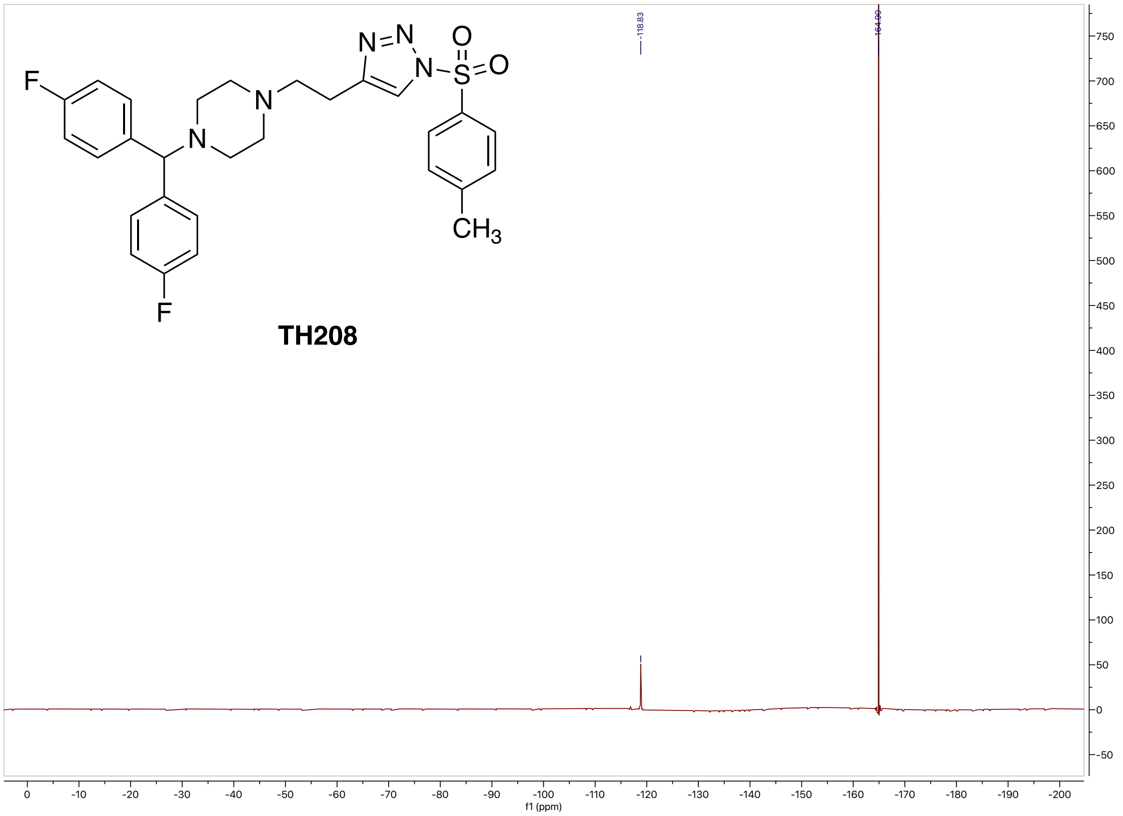

#### High-resolution mass spectrometry of TH208

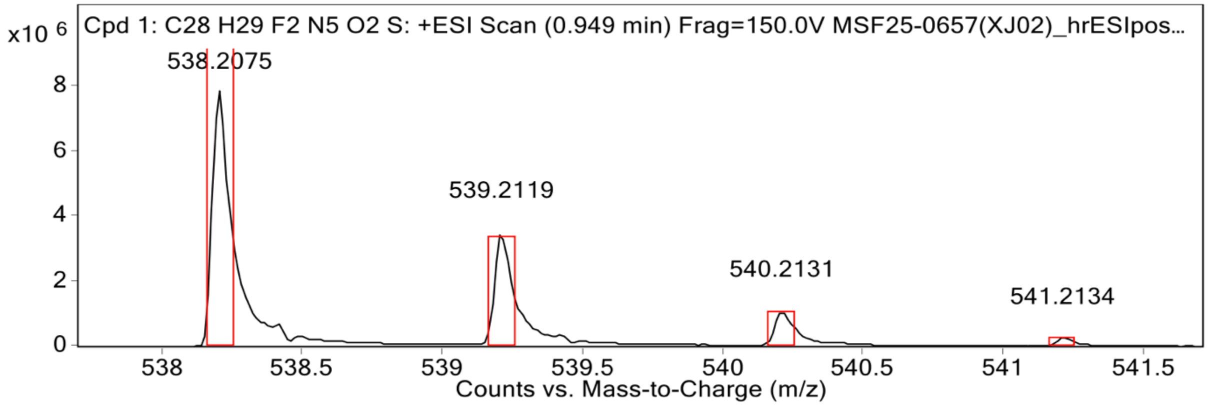


**MS Spectrum Peak List**

| Obs. m/z | Calc. m/z | Charge | Abundance | Formula | Ion Species | Tgt Mass Error (ppm) |
|---|---|---|---|---|---|---|
| 538.2075 | 538.2083 | 1 | 7881577 | C28H29F2N5O2S | (M+H)+ | 1.5 |
| 539.2119 | 539.2112 | 1 | 3511696 | C28H29F2N5O2S | (M+H)+ | -1.18 |
| 540.2131 | 540.2097 | 1 | 1079373 | C28H29F2N5O2S | (M+H)+ | -6.32 |
| 542.2276 | 542.2117 | 1 | 1279686 | C28H29F2N5O2S | (M+H)+ | -29.37 |
| 543.2308 | 543.2134 | 1 | 440881 | C28H29F2N5O2S | (M+H)+ | -32.07 |

#### HPLC spectrum of TH208

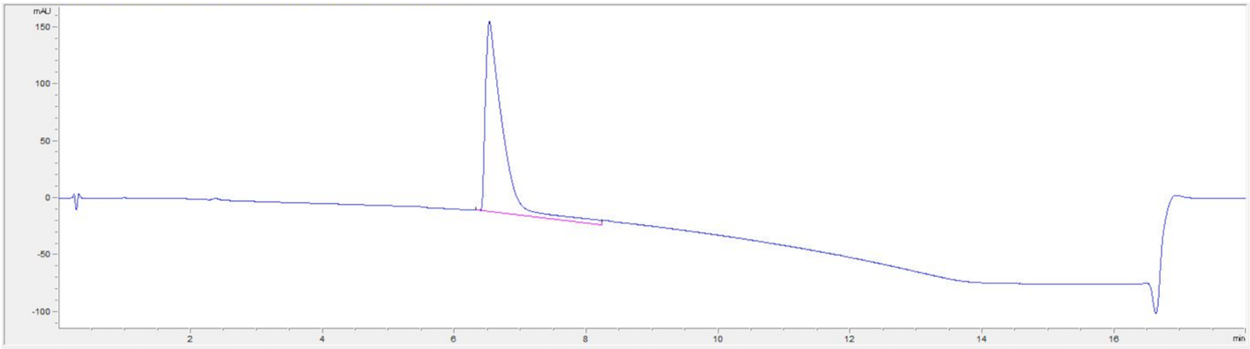

| # | Time | Area | Height | Width | Area% | Symmetry |
|---|---|---|---|---|---|---|
| 1 | 6.532 | 3060.2 | 167.8 | 0.2508 | 100.000 | 0.271 |

#### $^{1}$H NMR (800 MHz, $CDCl_3$) spectrum of TH220

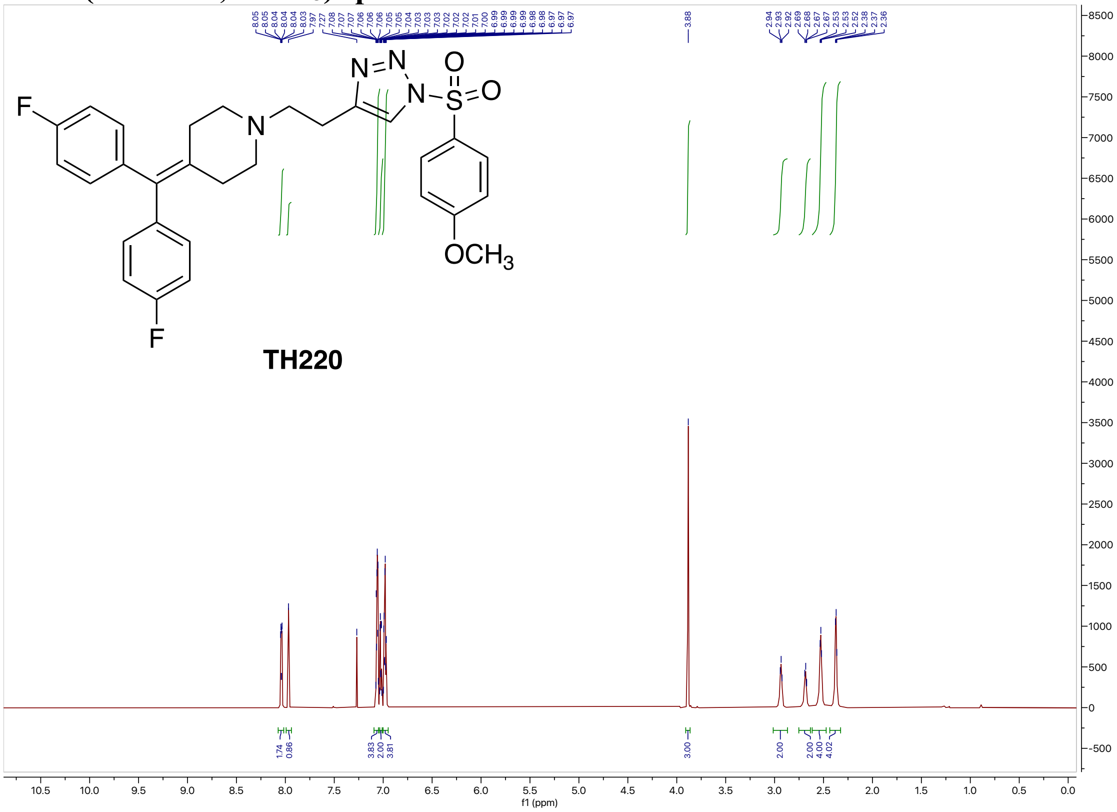

#### $^{13}$C NMR (200 MHz, $CDCl_3$) spectrum of TH220

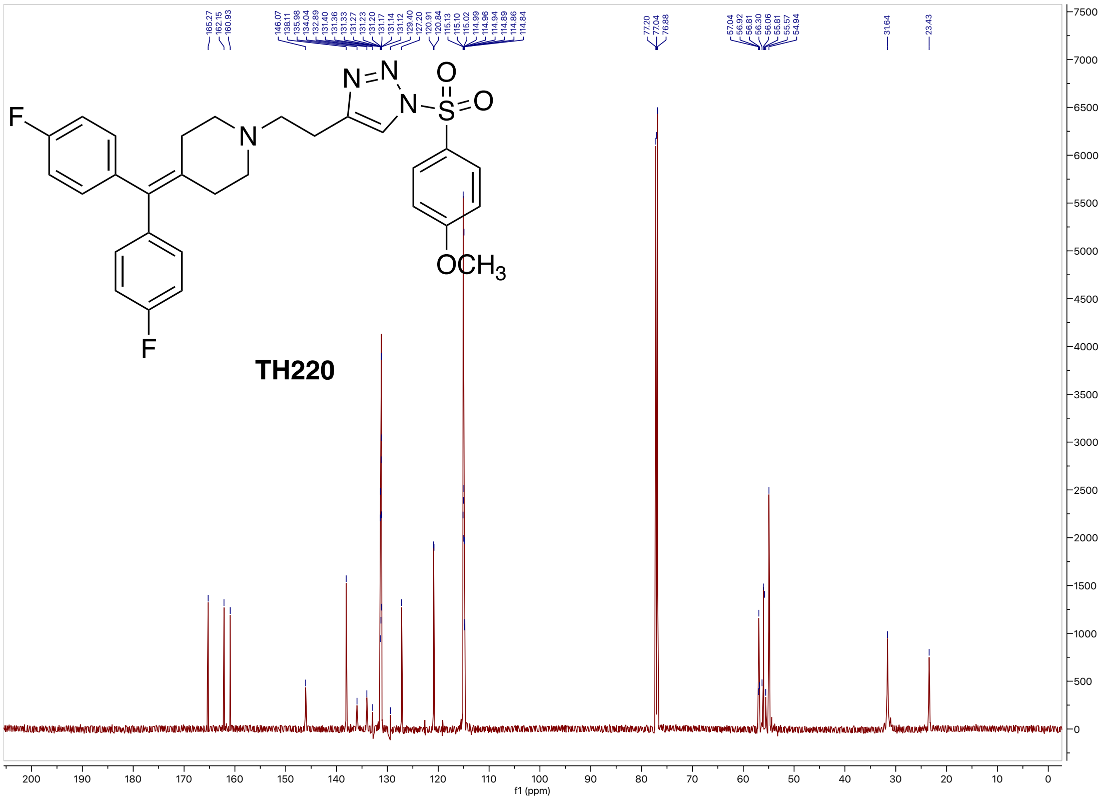

#### $^{19}$F NMR (564 MHz, $CDCl_3$) spectrum of TH220

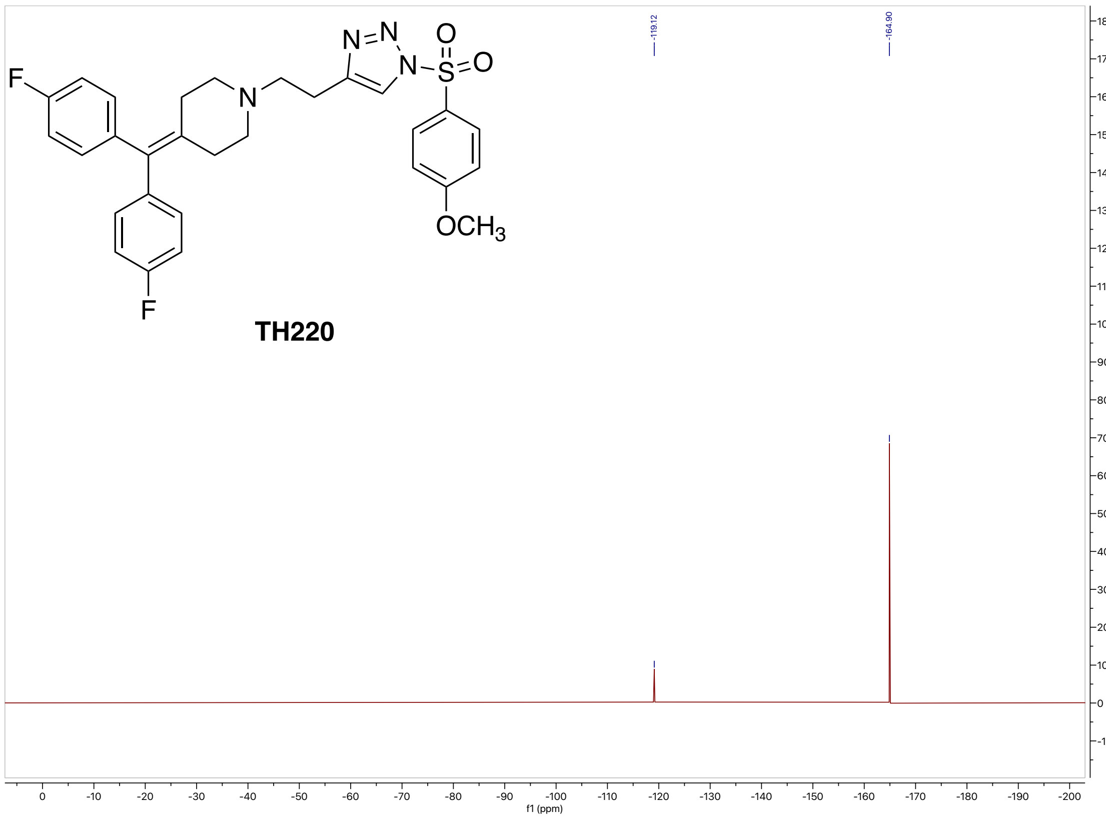

#### High-resolution mass spectrometry of TH220

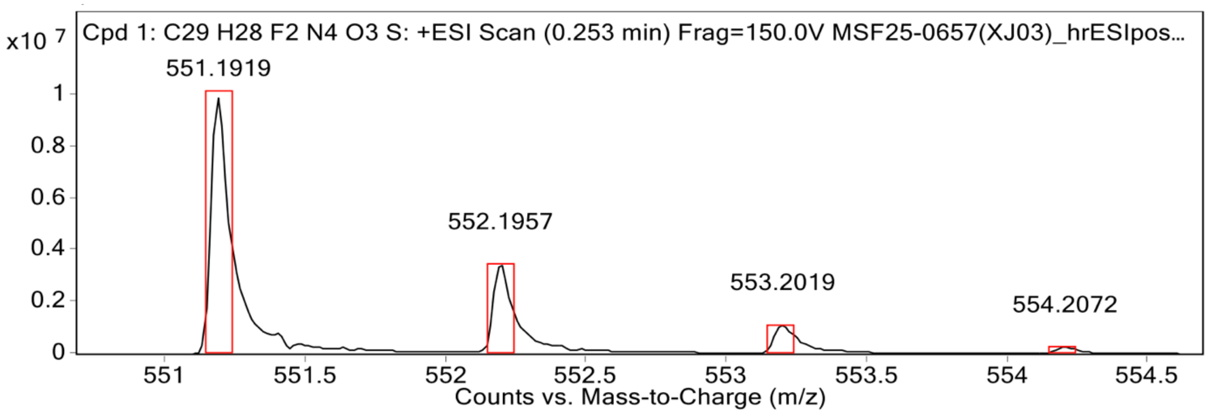


**MS Spectrum Peak List**

| Obs. m/z | Calc. m/z | Charge | Abundance | Formula | Ion Species | Tgt Mass Error (ppm) |
|---|---|---|---|---|---|---|
| 551.1919 | 551.1923 | 1 | 9906547 | C29H28F2N4O3S | (M+H)+ | 0.63 |
| 552.1957 | 552.1953 | 1 | 3481037 | C29H28F2N4O3S | (M+H)+ | -0.77 |
| 553.2019 | 553.1940 | 1 | 1134935 | C29H28F2N4O3S | (M+H)+ | -14.29 |
| 554.2072 | 554.1946 | 1 | 275003 | C29H28F2N4O3S | (M+H)+ | -22.64 |
| 557.1806 | 557.1997 | 1 | 66472 | C29H28F2N4O3S | (M+H)+ | 34.24 |

#### HPLC spectrum of TH220

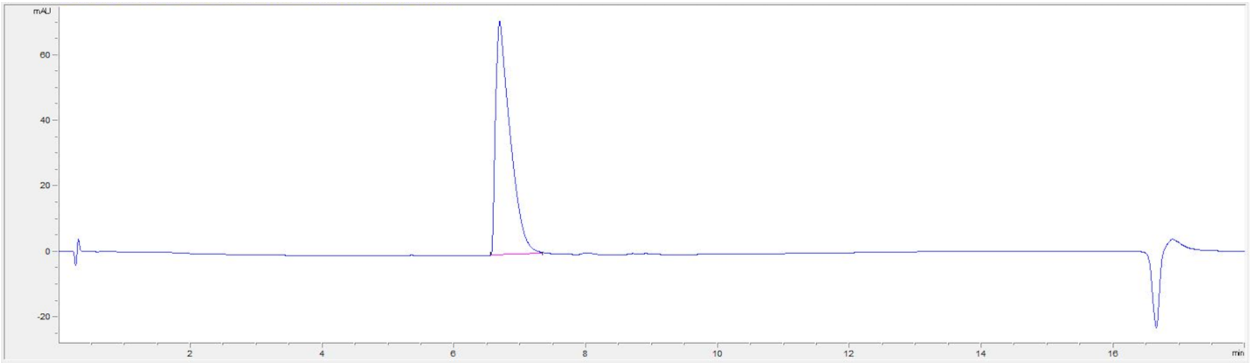

| # | Time | Area | Height | Width | Area% | Symmetry |
|---|---|---|---|---|---|---|
| 1 | 6.532 | 3060.2 | 167.8 | 0.2508 | 100.000 | 0.271 |

#### [1]H NMR (800 MHz, $CDCl_3$) spectrum of TH221

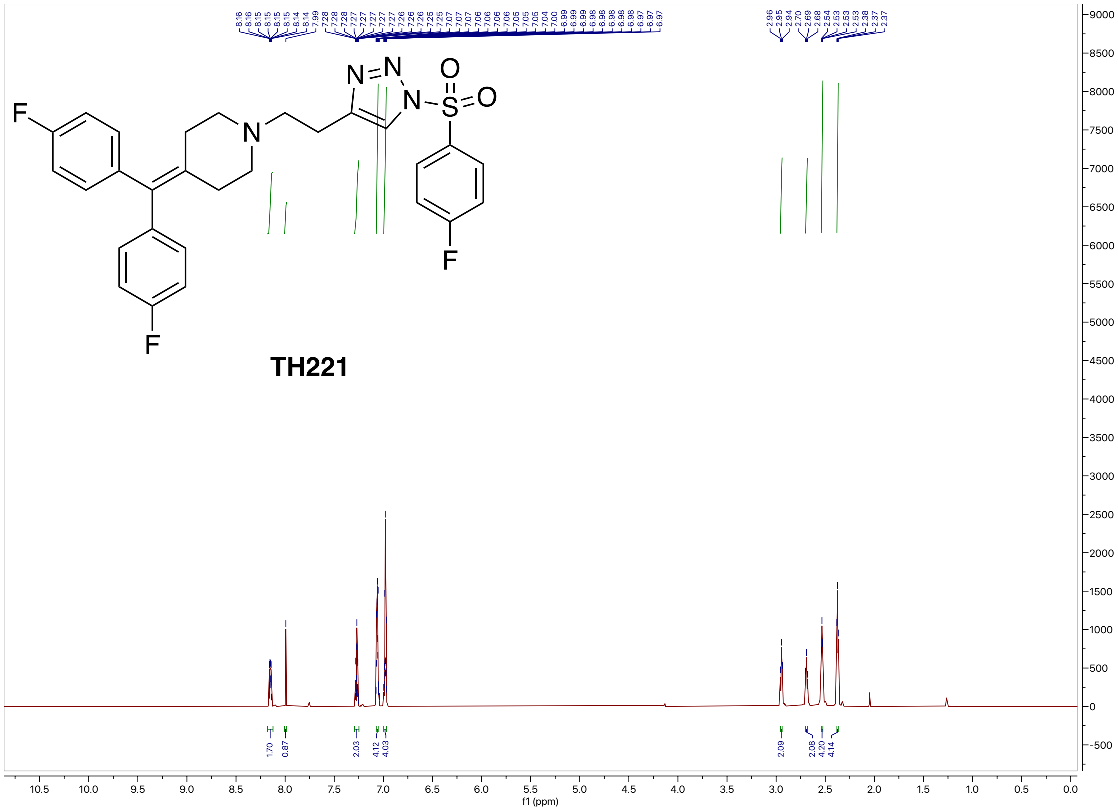

[13]C NMR (200 MHz, CDCl3) spectrum of TH221

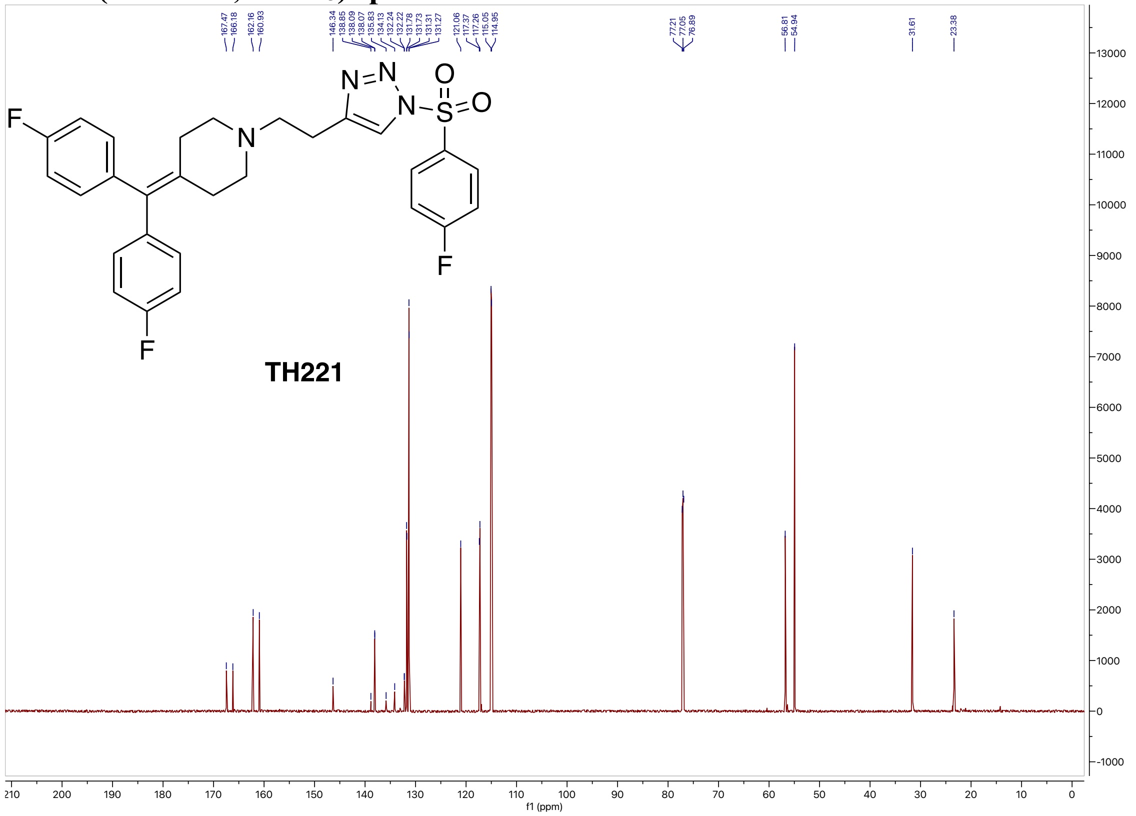

[19]F NMR (564 MHz, CDCl3) spectrum of TH221

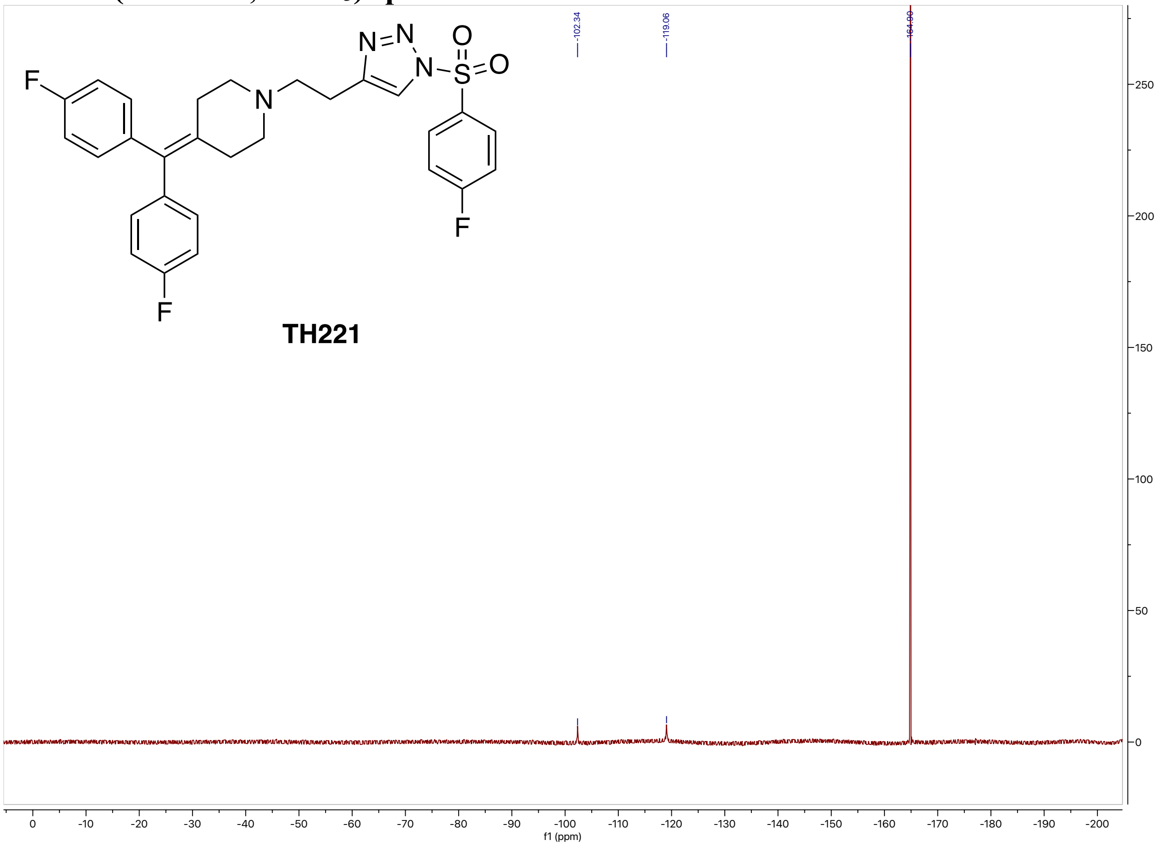

#### High-resolution mass spectrometry of TH221

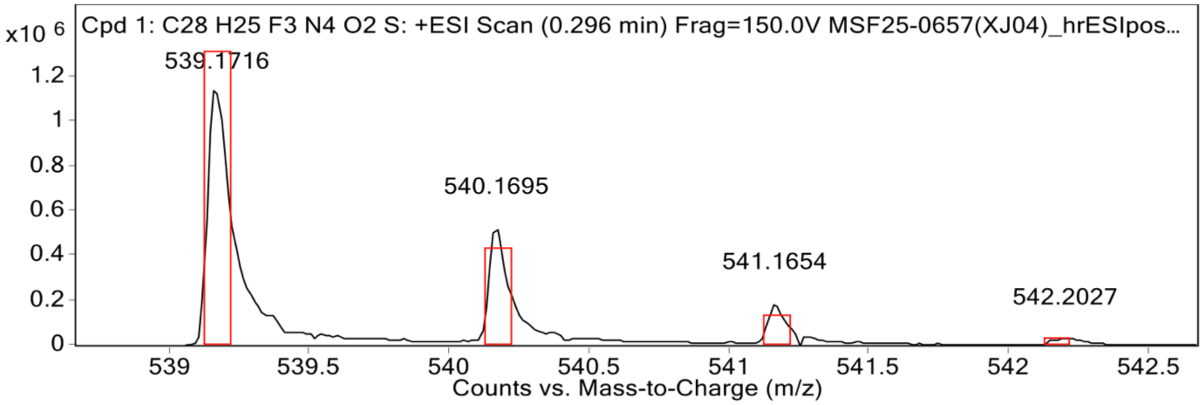


**MS Spectrum Peak List**

| Obs. m/z | Calc. m/z | Charge | Abundance | Formula | Ion Species | Tgt Mass Error (ppm) |
|---|---|---|---|---|---|---|
| 539.1716 | 539.1723 | 1 | 1157265 | C28H25F3N4O2S | (M+H)+ | 1.4 |
| 540.1695 | 540.1753 | 1 | 523671 | C28H25F3N4O2S | (M+H)+ | 10.7 |
| 541.1654 | 541.1737 | 1 | 187254 | C28H25F3N4O2S | (M+H)+ | 15.47 |
| 542.2027 | 542.1743 | 1 | 28962 | C28H25F3N4O2S | (M+H)+ | -52.38 |
| 543.1950 | 543.1758 | 1 | 8410 | C28H25F3N4O2S | (M+H)+ | -35.39 |

#### HPLC spectrum of TH221

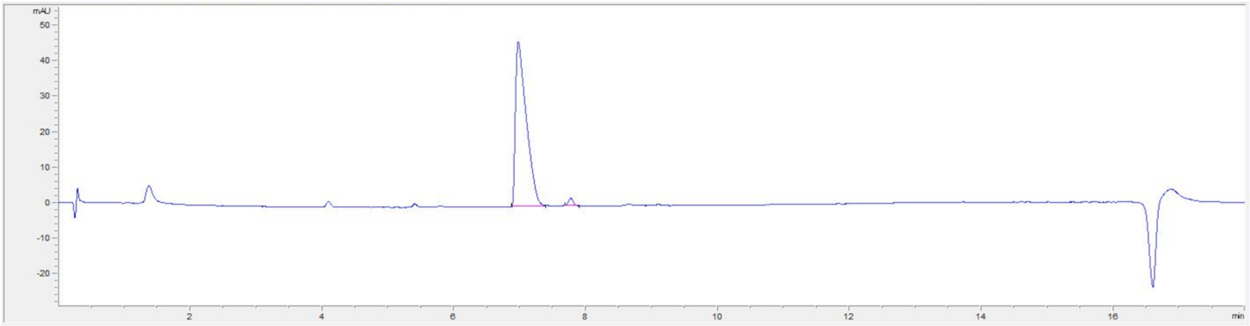


| # | Time | Area | Height | Width | Area% | Symmetry |
|---|---|---|---|---|---|---|
| 1 | 6.977 | 530.5 | 46.6 | 0.1607 | 97.990 | 0.336 |
| 2 | 7.775 | 10.9 | 2.2 | 0.0755 | 2.010 | 0.851 |

#### $^{1}$H NMR (800 MHz, $CDCl_3$) spectrum of TH223

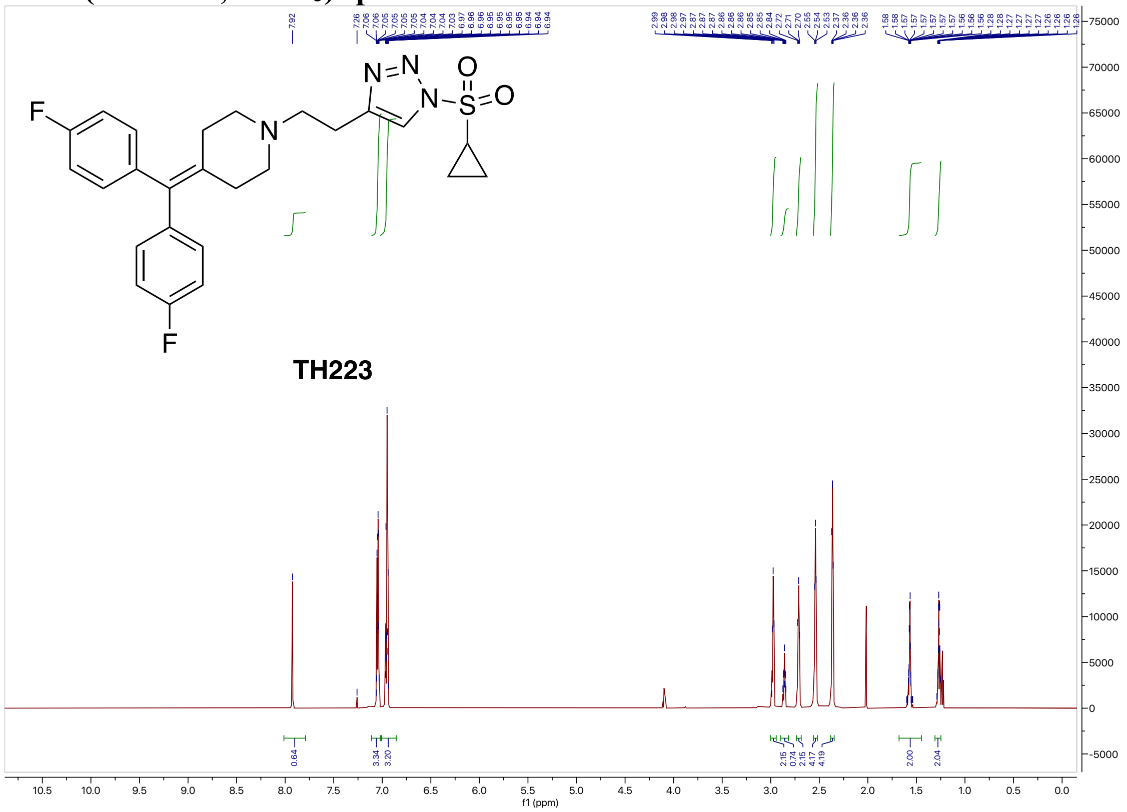


#### $^{13}$C NMR (200 MHz, $CDCl_3$) spectrum of TH223

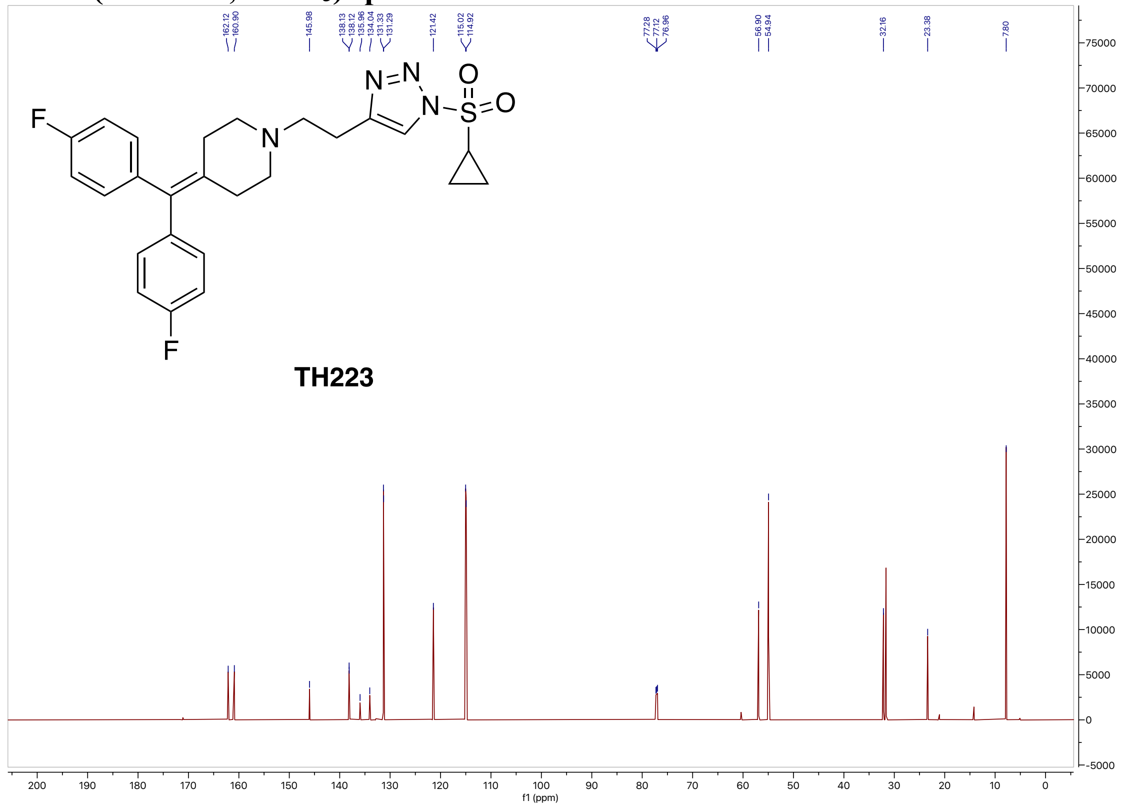

#### $^{19}$F NMR (564 MHz, $CDCl_3$) spectrum of TH223

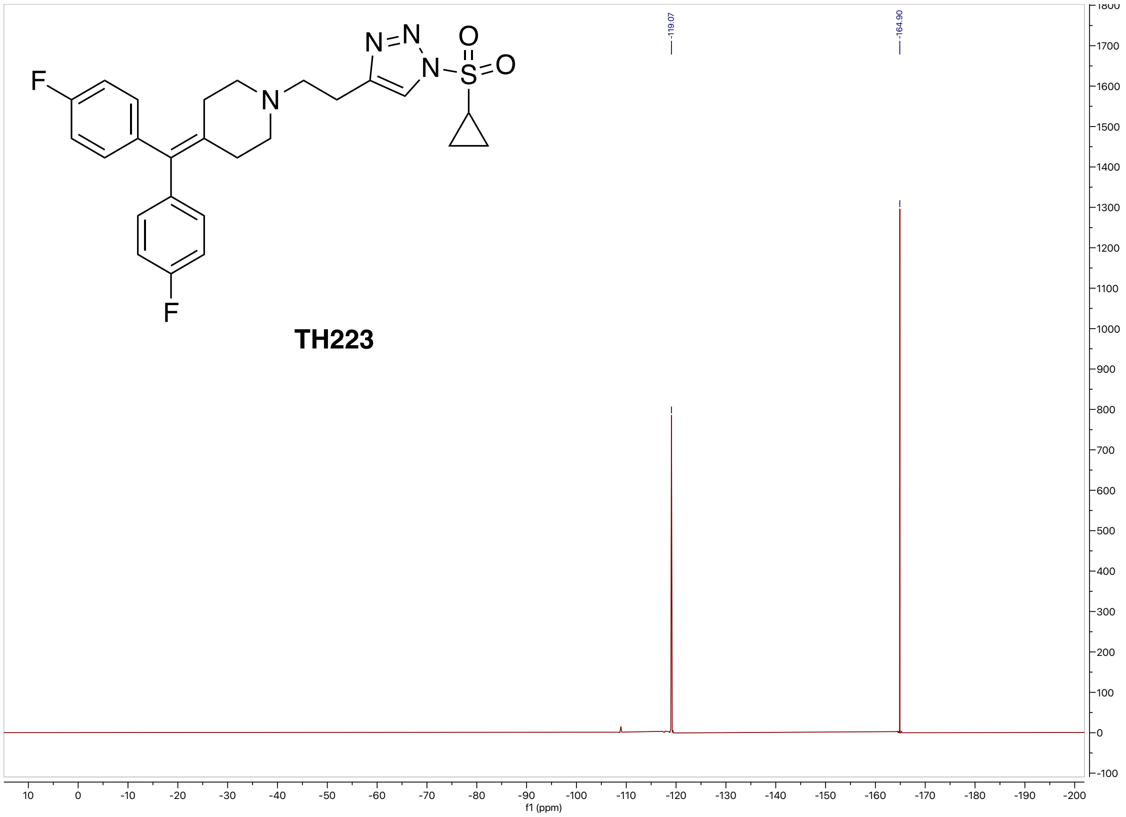


#### High-resolution mass spectrometry of TH223

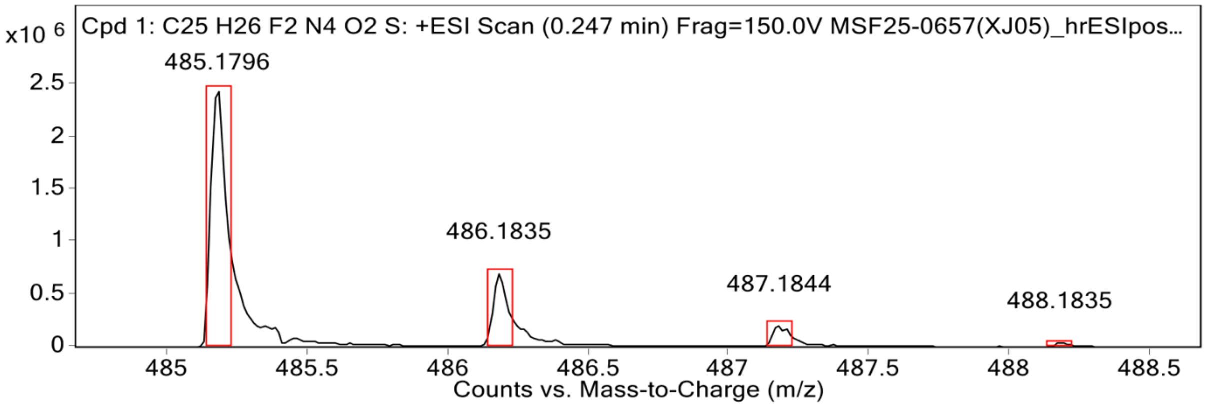


**MS Spectrum Peak List**

| Obs. m/z | Calc. m/z | Charge | Abundance | Formula | Ion Species | Tgt Mass Error (ppm) |
|---|---|---|---|---|---|---|
| 485.1796 | 485.1817 | 1 | 2474359 | C25H26F2N4O2S | (M+H)+ | 4.46 |
| 486.1835 | 486.1847 | 1 | 697307 | C25H26F2N4O2S | (M+H)+ | 2.52 |
| 487.1844 | 487.1826 | 1 | 197976 | C25H26F2N4O2S | (M+H)+ | -3.56 |
| 488.1835 | 488.1833 | 1 | 38292 | C25H26F2N4O2S | (M+H)+ | -0.26 |
| 489.1326 | 489.1848 | 1 | 3502 | C25H26F2N4O2S | (M+H)+ | 106.85 |

**HPLC spectrum of TH223**

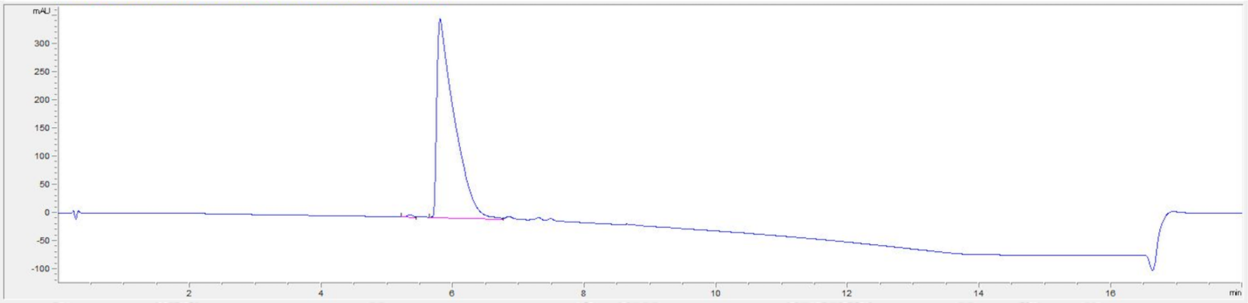


| # | Time | Area | Height | Width | Area% | Symmetry |
|---|---|---|---|---|---|---|
| 1 | 5.349 | 41.8 | 5.5 | 0.1269 | 0.628 | 0.656 |
| 2 | 5.802 | 6607 | 353.7 | 0.2479 | 99.372 | 0.206 |

**[1]H NMR (800 MHz, $CDCl_3$) spectrum of TH225**

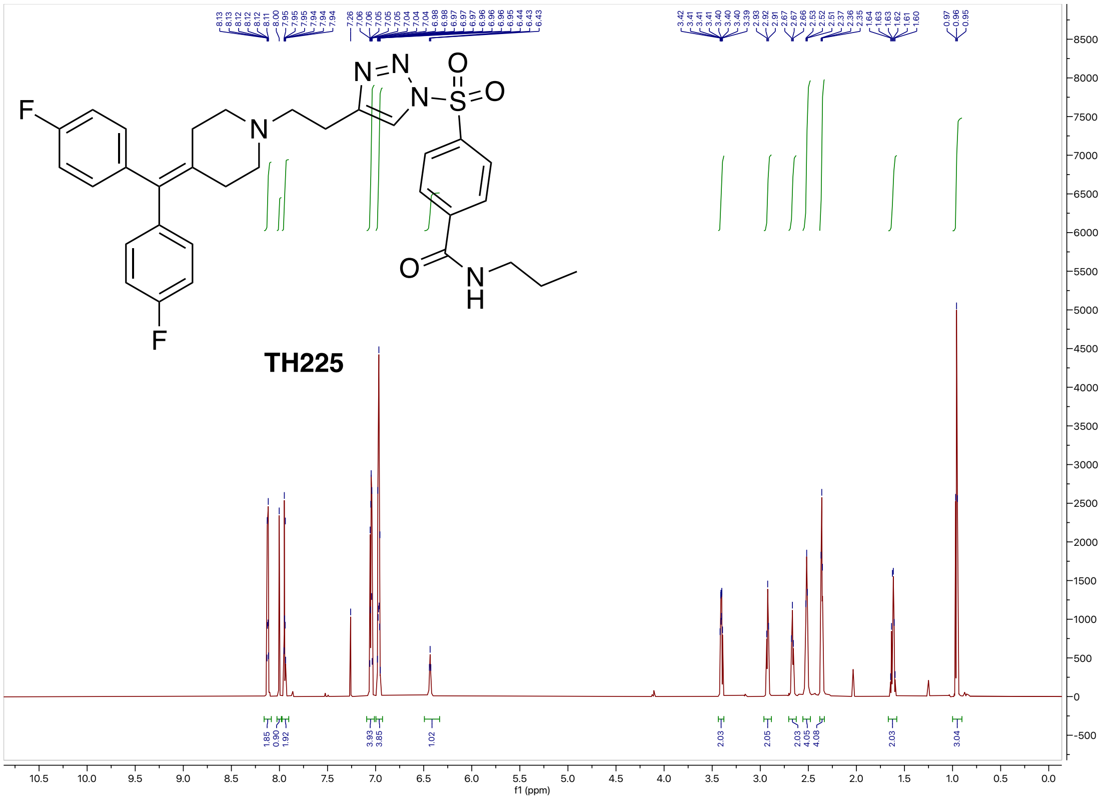

**$^{13}$C NMR (200 MHz, $CDCl_3$) spectrum of TH225**

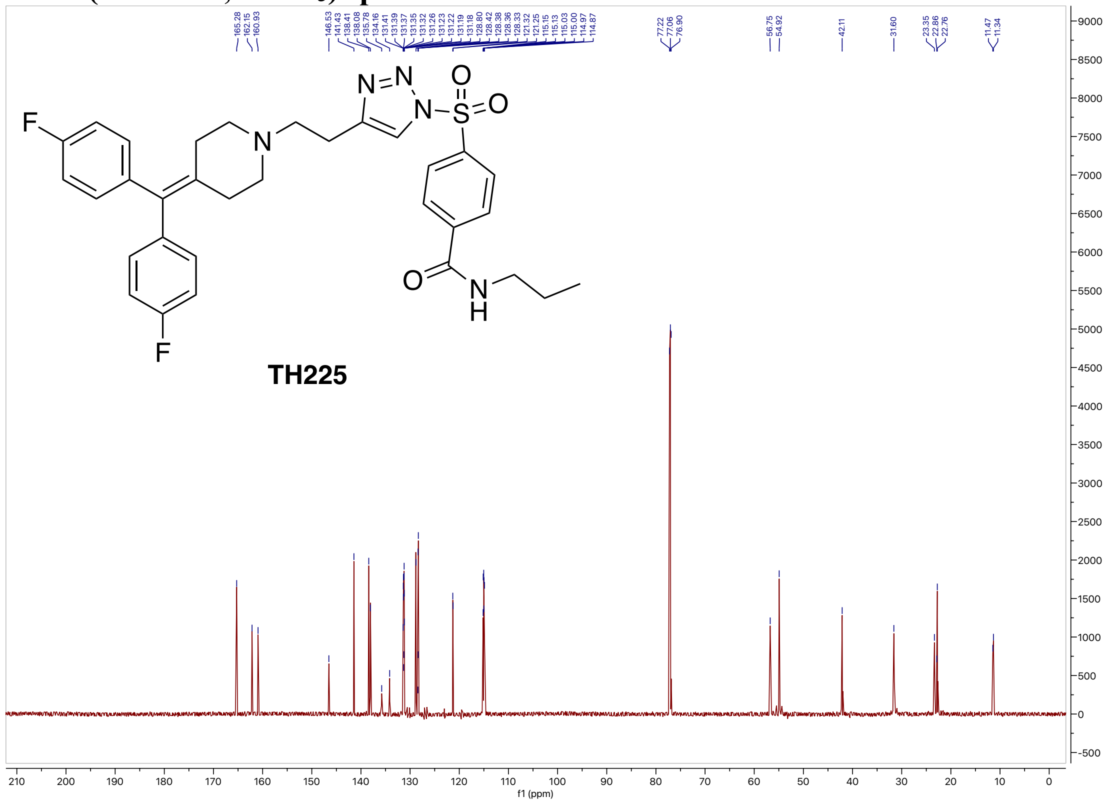


**$^{19}$F NMR (564 MHz, $CDCl_3$) spectrum of TH225**

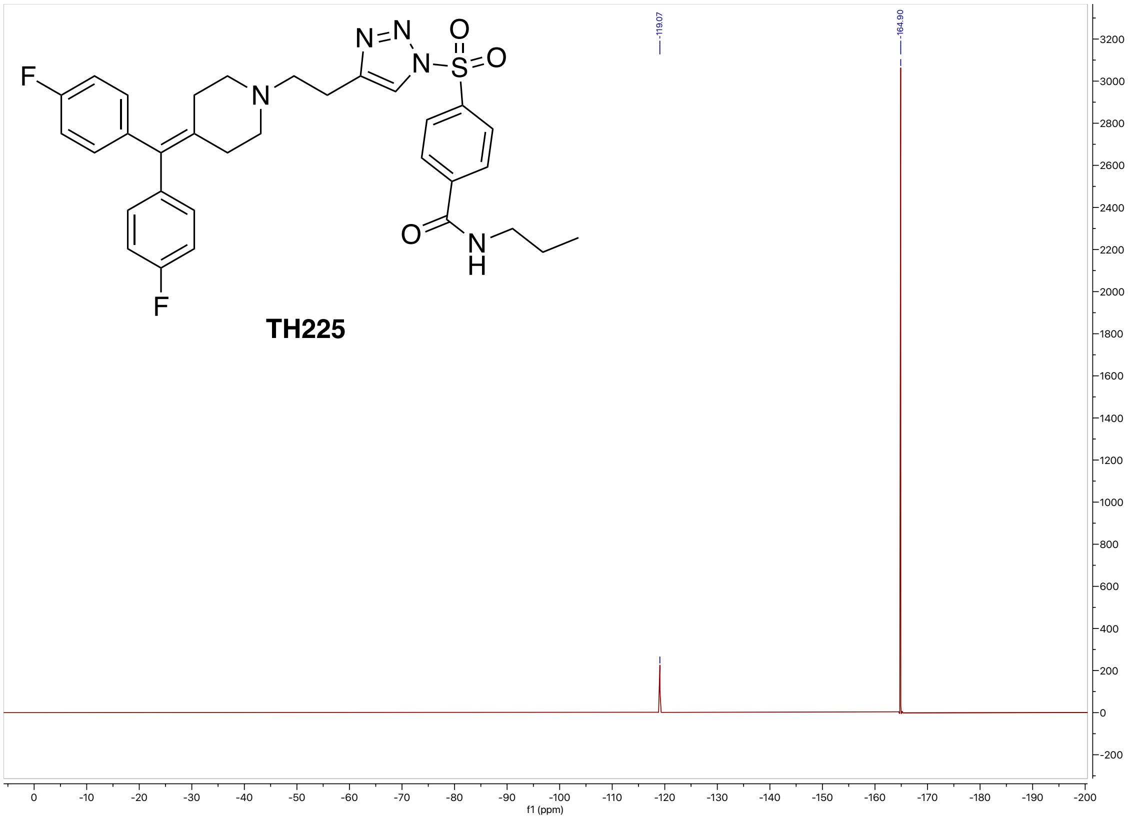

**High-resolution mass spectrometry of TH225**

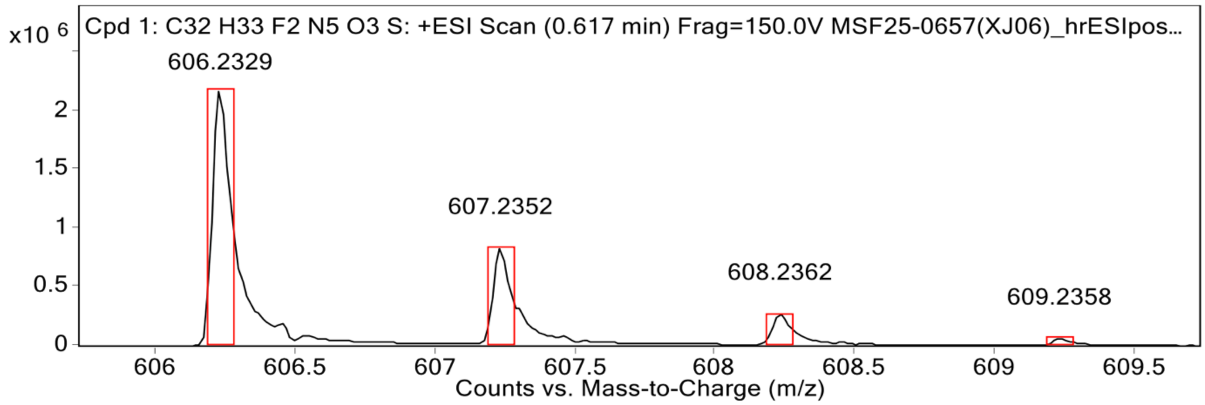


MS Spectrum Peak List

| Obs. m/z | Calc. m/z | Charge | Abundance | Formula | Ion Species | Tgt Mass Error (ppm) |
|---|---|---|---|---|---|---|
| 606.2329 | 606.2345 | 1 | 2172541 | C32H33F2N5O3S | (M+H)+ | 2.61 |
| 607.2352 | 607.2375 | 1 | 828311 | C32H33F2N5O3S | (M+H)+ | 3.78 |
| 608.2362 | 608.2366 | 1 | 269153 | C32H33F2N5O3S | (M+H)+ | 0.6 |
| 609.2358 | 609.2372 | 1 | 58211 | C32H33F2N5O3S | (M+H)+ | 2.25 |
| 610.2378 | 610.2385 | 1 | 7725 | C32H33F2N5O3S | (M+H)+ | 1.24 |

**HPLC spectrum of TH225**

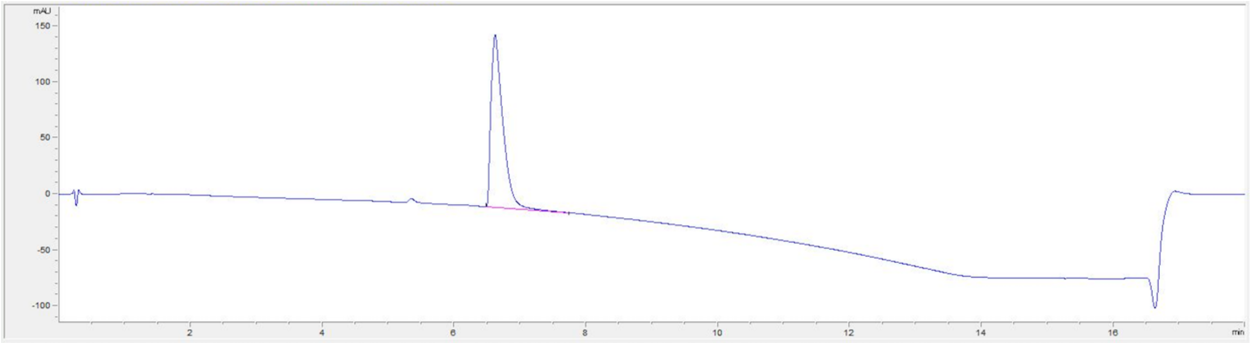


| # | Time | Area | Height | Width | Area% | Symmetry |
|---|---|---|---|---|---|---|
| 1 | 6.624 | 1921.7 | 154.4 | 0.1852 | 100.000 | 0.485 |

#### [1]H NMR (800 MHz, $CDCl_3$) spectrum of XJ-3-9

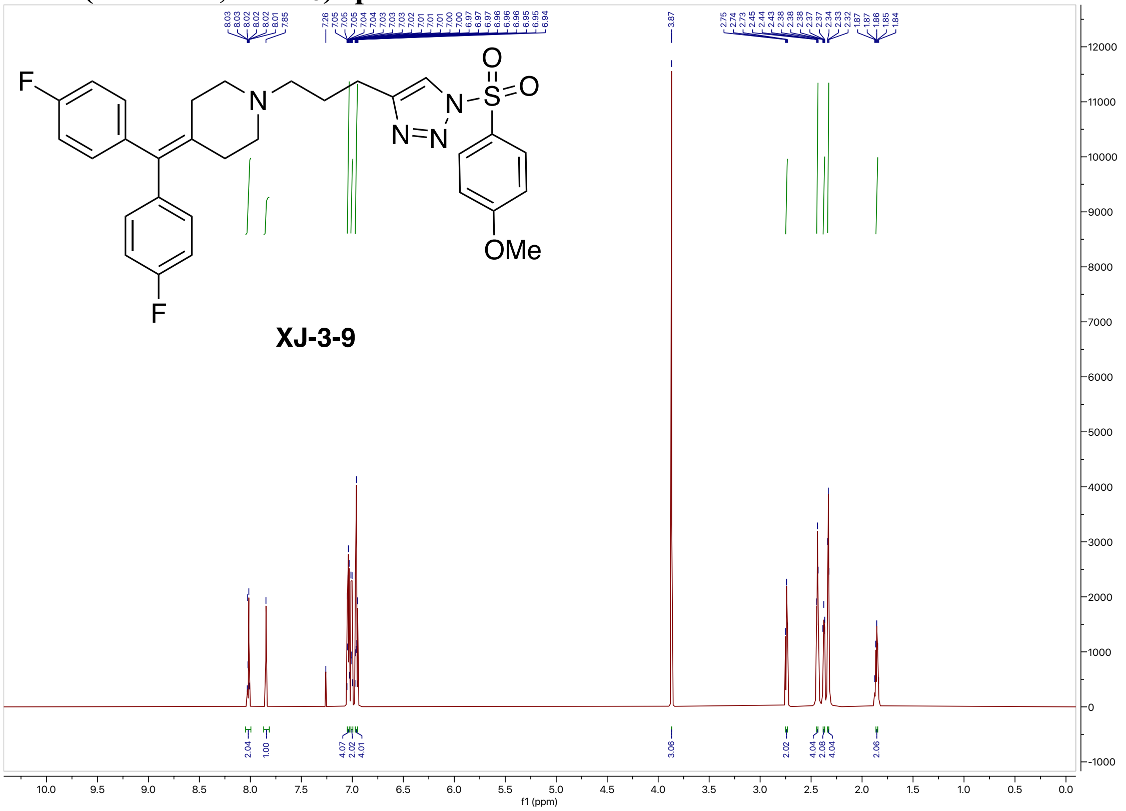

#### [13]C NMR (200 MHz, $CDCl_3$) spectrum of XJ-3-9

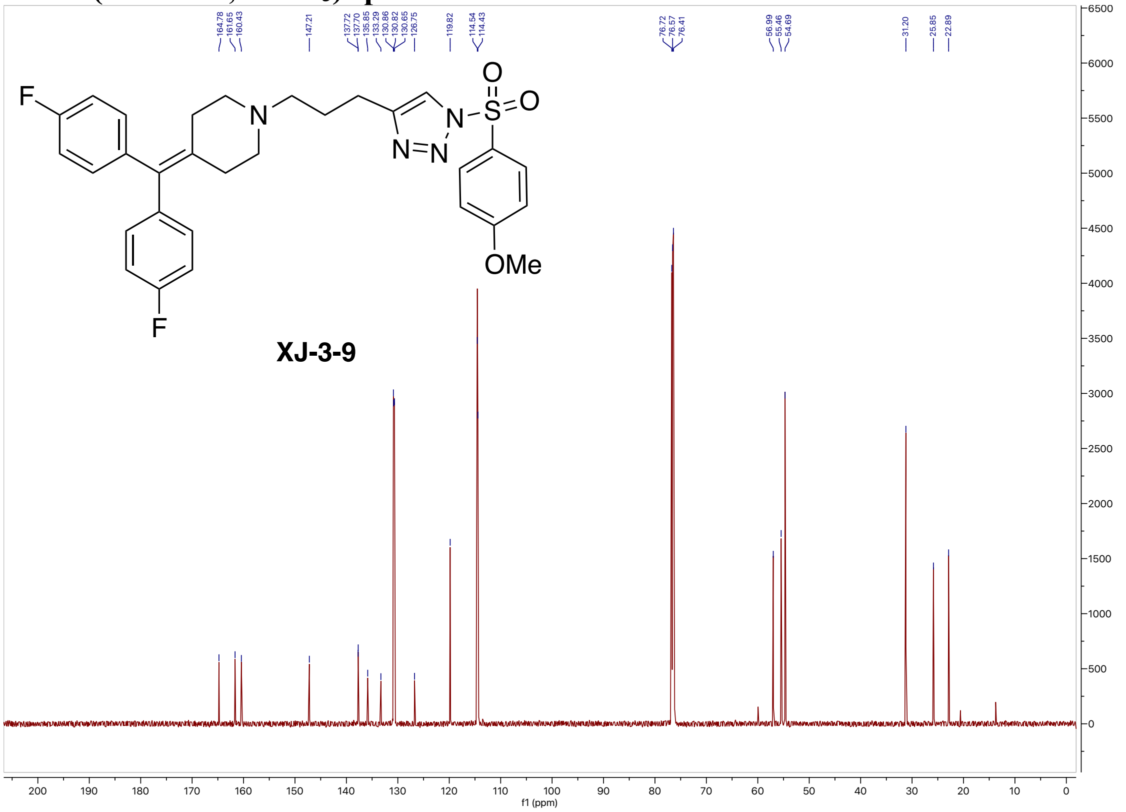

#### $^{19}F$ NMR (564 MHz, $CDCl_3$) spectrum of XJ-3-9

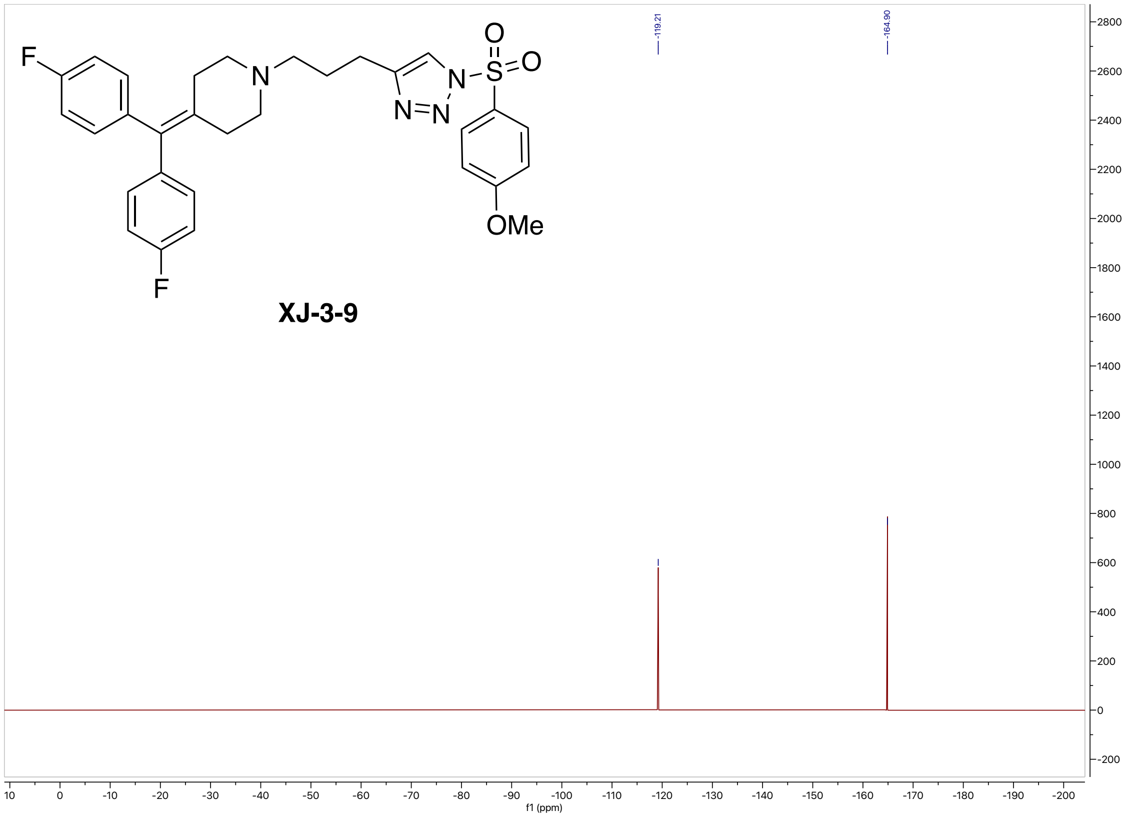

#### High-resolution mass spectrometry of XJ-3-9

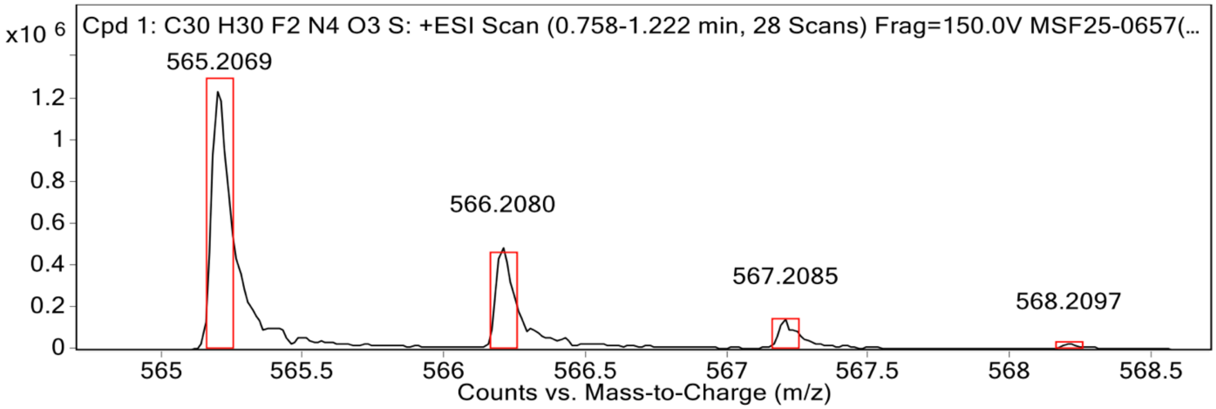


**MS Spectrum Peak List**

| Obs. m/z | Calc. m/z | Charge | Abundance | Formula | Ion Species | Tgt Mass Error (ppm) |
|---|---|---|---|---|---|---|
| 565.2069 | 565.2079 | 1 | 1258939 | C30H30F2N4O3S | (M+H)+ | 1.93 |
| 566.2080 | 566.2110 | 1 | 490108 | C30H30F2N4O3S | (M+H)+ | 5.25 |
| 567.2085 | 567.2098 | 1 | 146029 | C30H30F2N4O3S | (M+H)+ | 2.32 |
| 568.2097 | 568.2104 | 1 | 25616 | C30H30F2N4O3S | (M+H)+ | 1.29 |
| 569.2113 | 569.2118 | 1 | 5787 | C30H30F2N4O3S | (M+H)+ | 0.88 |

#### HPLC spectrum of XJ-3-9

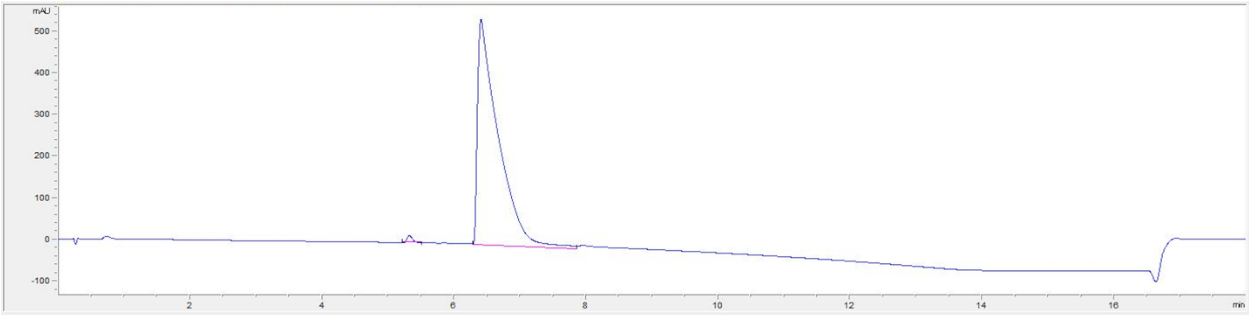

| # | Time | Area | Height | Width | Area% | Symmetry |
|---|---|---|---|---|---|---|
| 1 | 5.324 | 94.6 | 16.5 | 0.091 | 0.796 | 0.731 |
| 2 | 6.407 | 11795.7 | 542.2 | 0.2858 | 99.204 | 0.187 |

#### $^1$H NMR (800 MHz, $CDCl_3$) spectrum of XJ-3-23

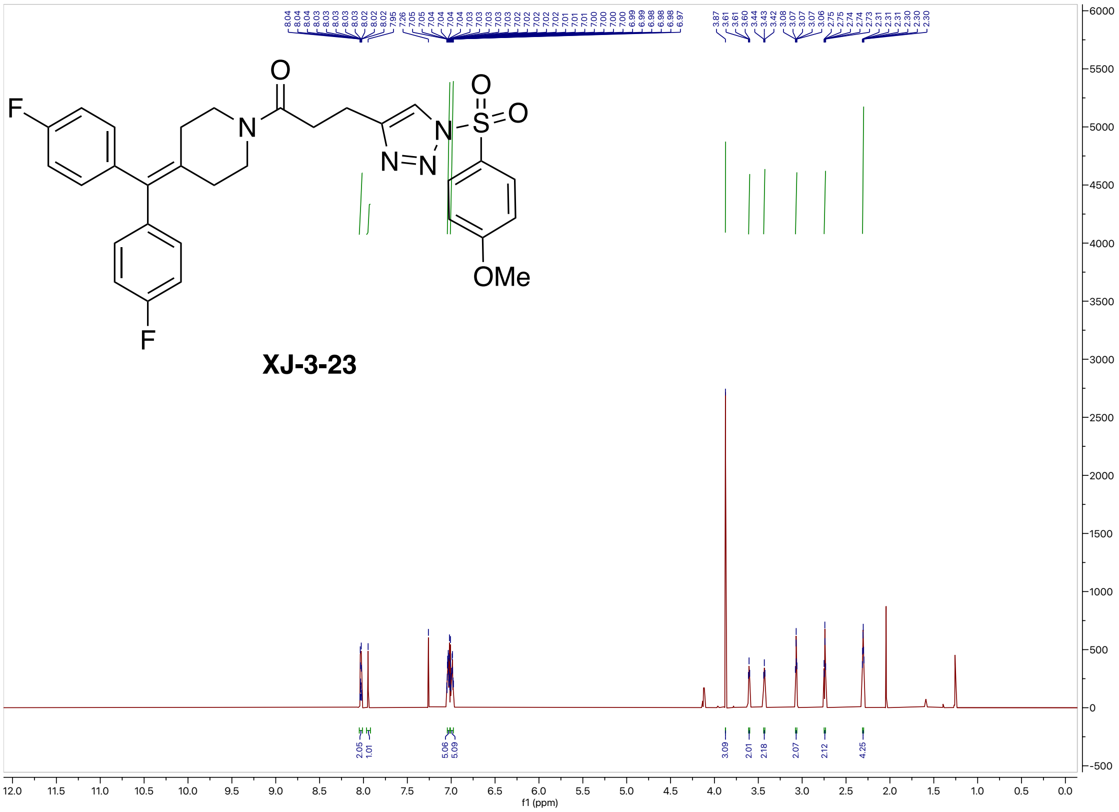

#### $^{13}$C NMR (200 MHz, $CDCl_3$) spectrum of XJ-3-23

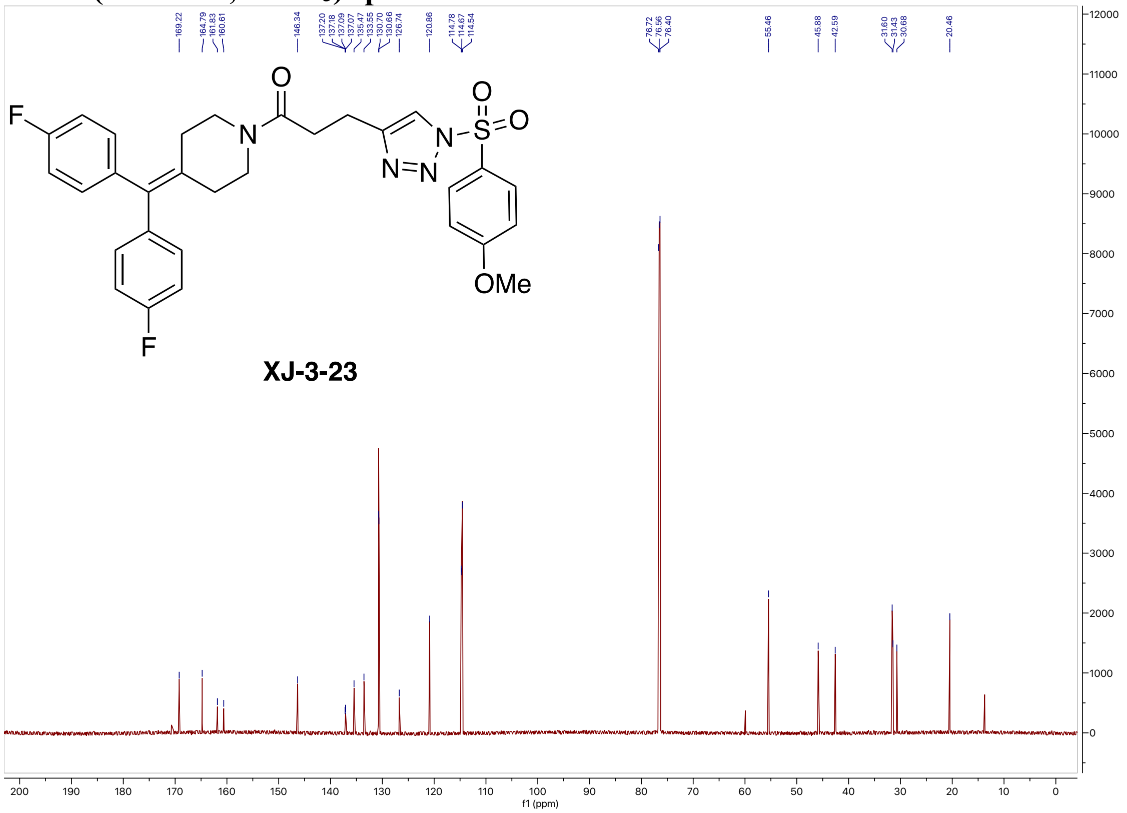


#### $^{19}$F NMR (564 MHz, $CDCl_3$) spectrum of XJ-3-23

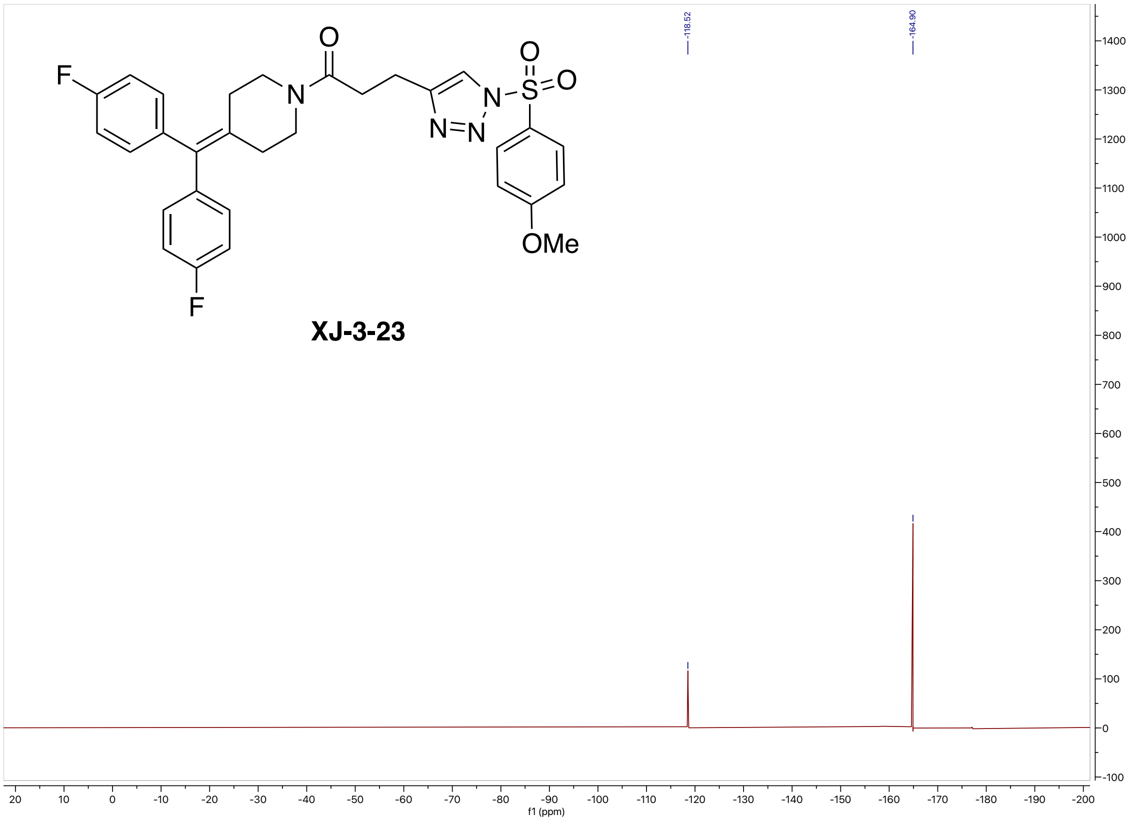

#### High-resolution mass spectrometry of XJ-3-23

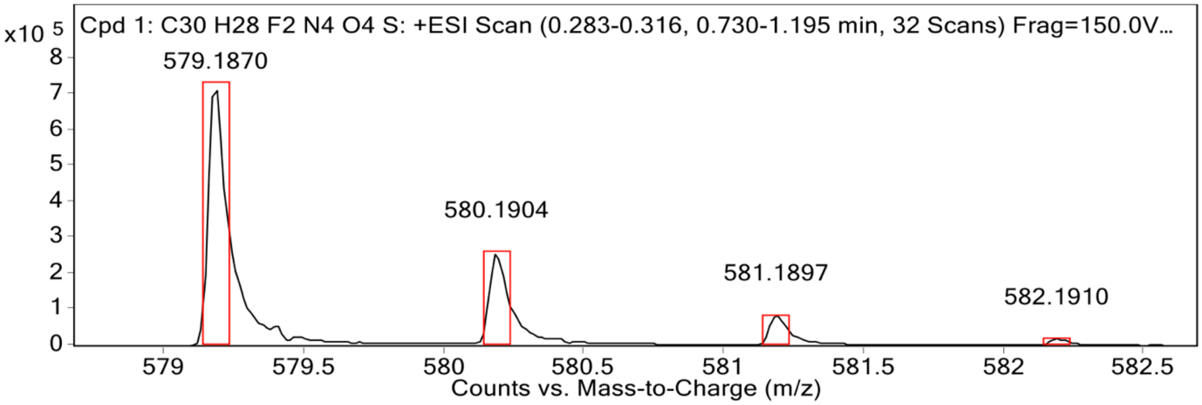


**MS Spectrum Peak List**

| Obs. m/z | Calc. m/z | Charge | Abundance | Formula | Ion Species | Tgt Mass Error (ppm) |
|---|---|---|---|---|---|---|
| 409.1834 | | | 1485649 | | | |
| 579.1870 | 579.1872 | 1 | 720136 | C30H28F2N4O4S | (M+H)+ | 0.34 |
| 580.1904 | 580.1902 | 1 | 260039 | C30H28F2N4O4S | (M+H)+ | -0.21 |
| 581.1897 | 581.1891 | 1 | 84986 | C30H28F2N4O4S | (M+H)+ | -1.01 |
| 582.1910 | 582.1898 | 1 | 18531 | C30H28F2N4O4S | (M+H)+ | -2.12 |

#### HPLC spectrum of XJ-3-23

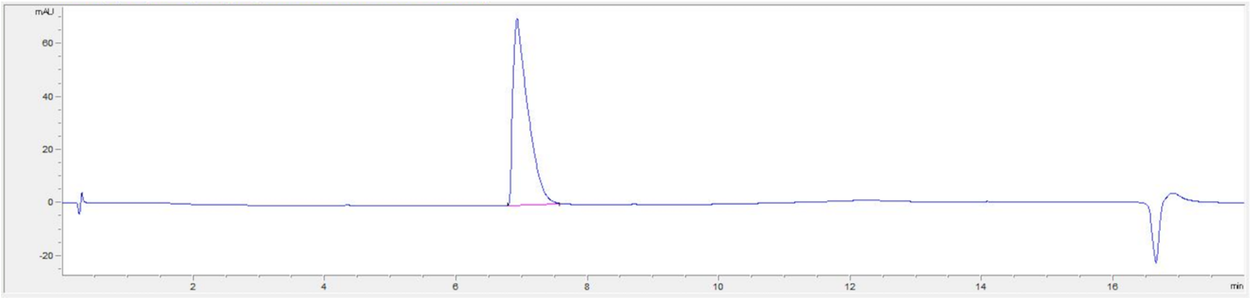


| # | Time | Area | Height | Width | Area% | Symmetry |
|---|---|---|---|---|---|---|
| 1 | 6.924 | 1117.8 | 70.6 | 0.2228 | 100.000 | 0.338 |

**$^1$H NMR (800 MHz, $CDCl_3$) spectrum of XJ-3-41**

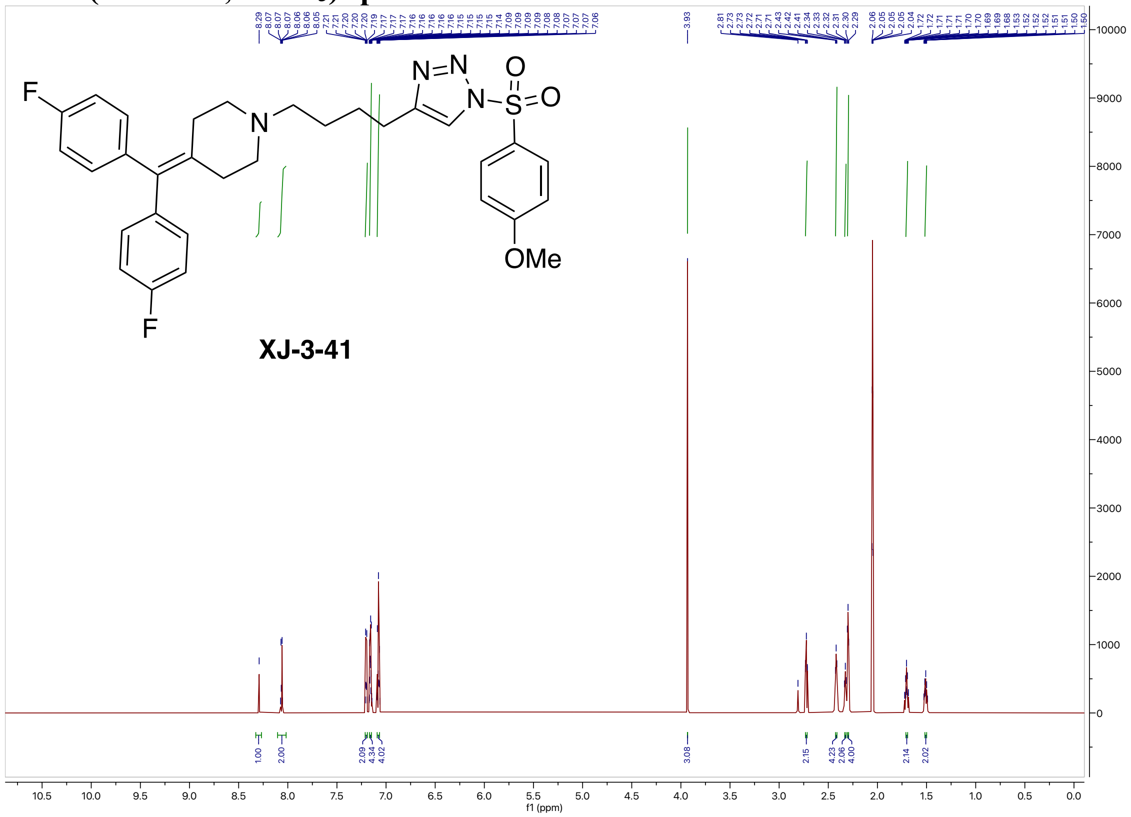

**$^{13}$C NMR (200 MHz, $CDCl_3$) spectrum of XJ-3-41**

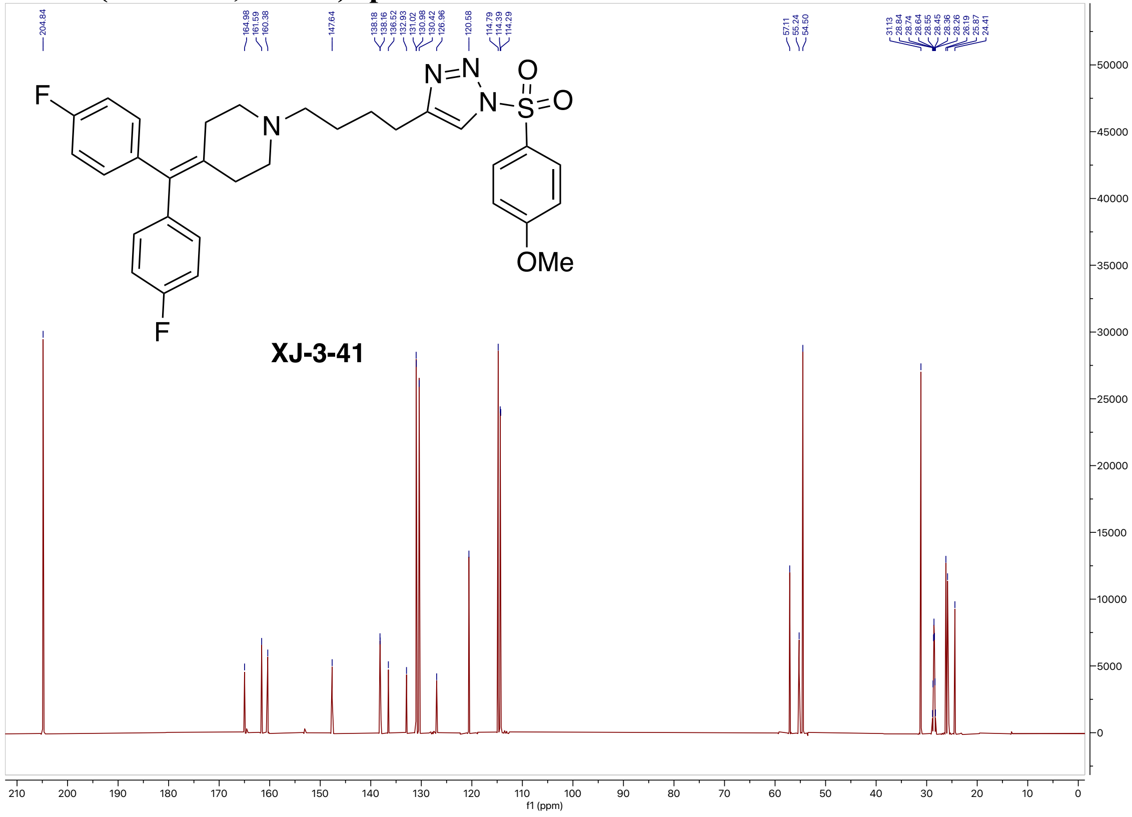

#### [19]F NMR (564 MHz, $CDCl_3$) spectrum of XJ-3-41

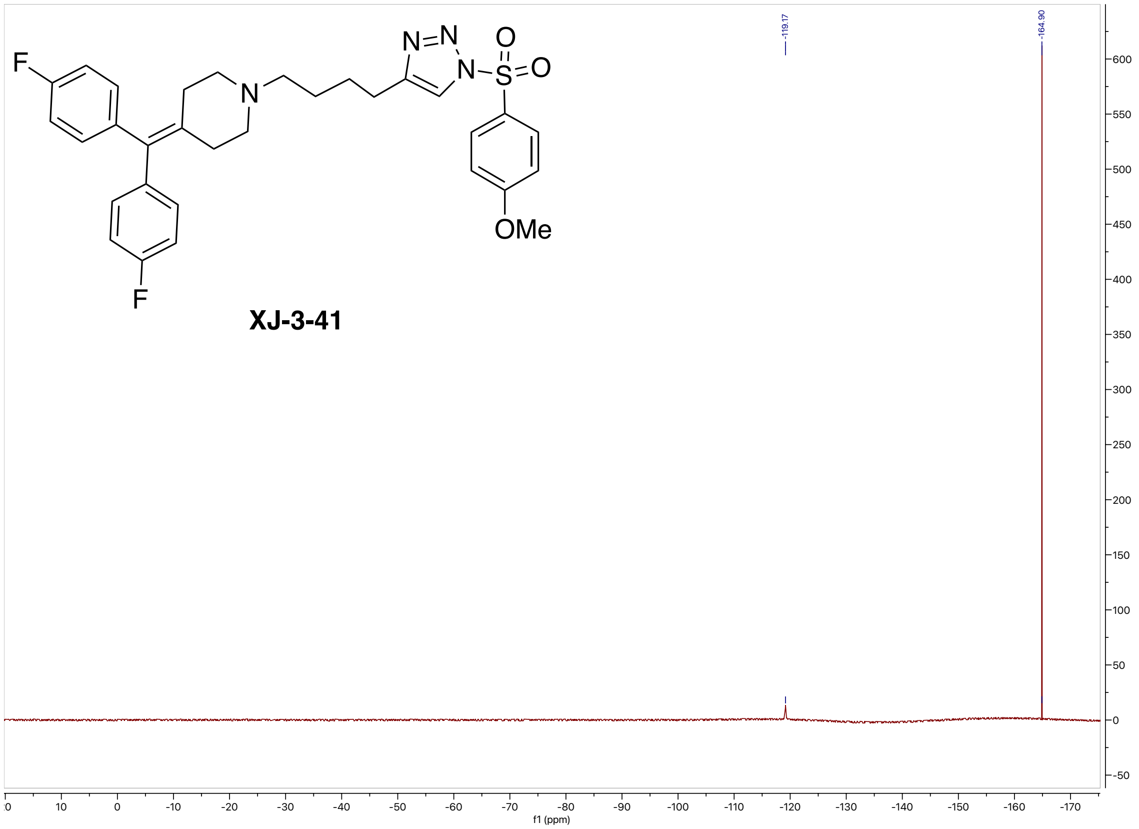


#### High-resolution mass spectrometry of XJ-3-41

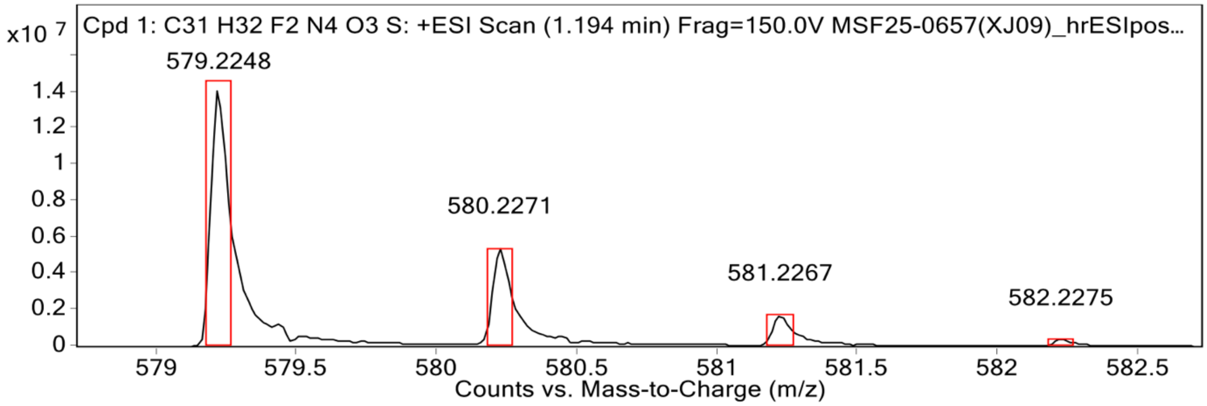


**MS Spectrum Peak List**

| Obs. m/z | Calc. m/z | Charge | Abundance | Formula | Ion Species | Tgt Mass Error (ppm) |
|---|---|---|---|---|---|---|
| 579.2248 | 579.2236 | 1 | 14186934 | C31H32F2N4O3S | (M+H)+ | -2.14 |
| 580.2271 | 580.2266 | 1 | 5409072 | C31H32F2N4O3S | (M+H)+ | -0.77 |
| 581.2267 | 581.2256 | 1 | 1719537 | C31H32F2N4O3S | (M+H)+ | -1.96 |
| 582.2275 | 582.2262 | 1 | 361836 | C31H32F2N4O3S | (M+H)+ | -2.35 |
| 583.2364 | 583.2276 | 1 | 102686 | C31H32F2N4O3S | (M+H)+ | -15.12 |

**HPLC spectrum of XJ-3-41**

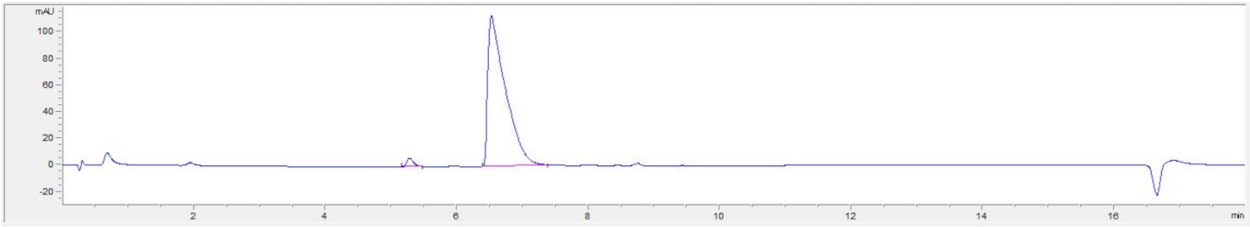

| # | Time | Area | Height | Width | Area% | Symmetry |
|---|---|---|---|---|---|---|
| 1 | 5.282 | 44.1 | 6.5 | 0.103 | 2.054 | 0.599 |
| 2 | 6.528 | 2104.5 | 113.3 | 0.2508 | 97.946 | 0.236 |

**[1]H NMR (800 MHz, Acetone-$d_6$) spectrum of XJ-3-65**

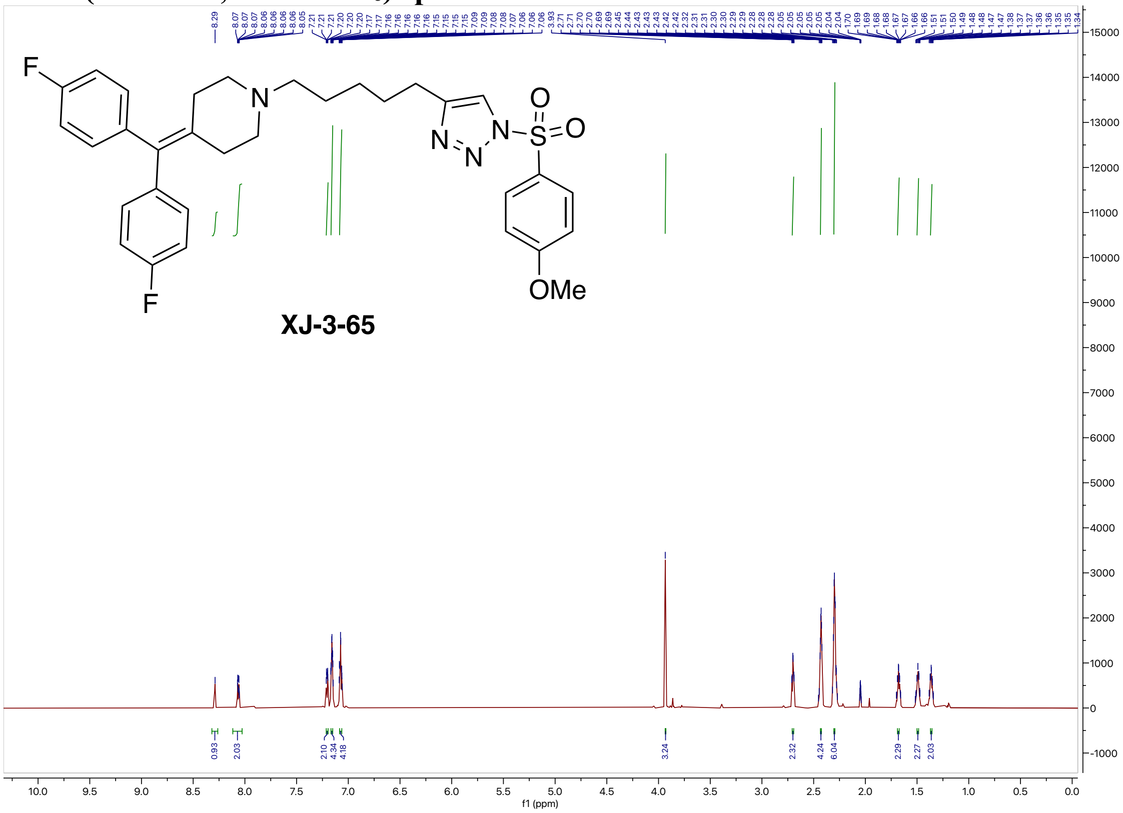

#### $^{13}$C NMR (200 MHz, Acetone-$d_6$) spectrum of XJ-3-65

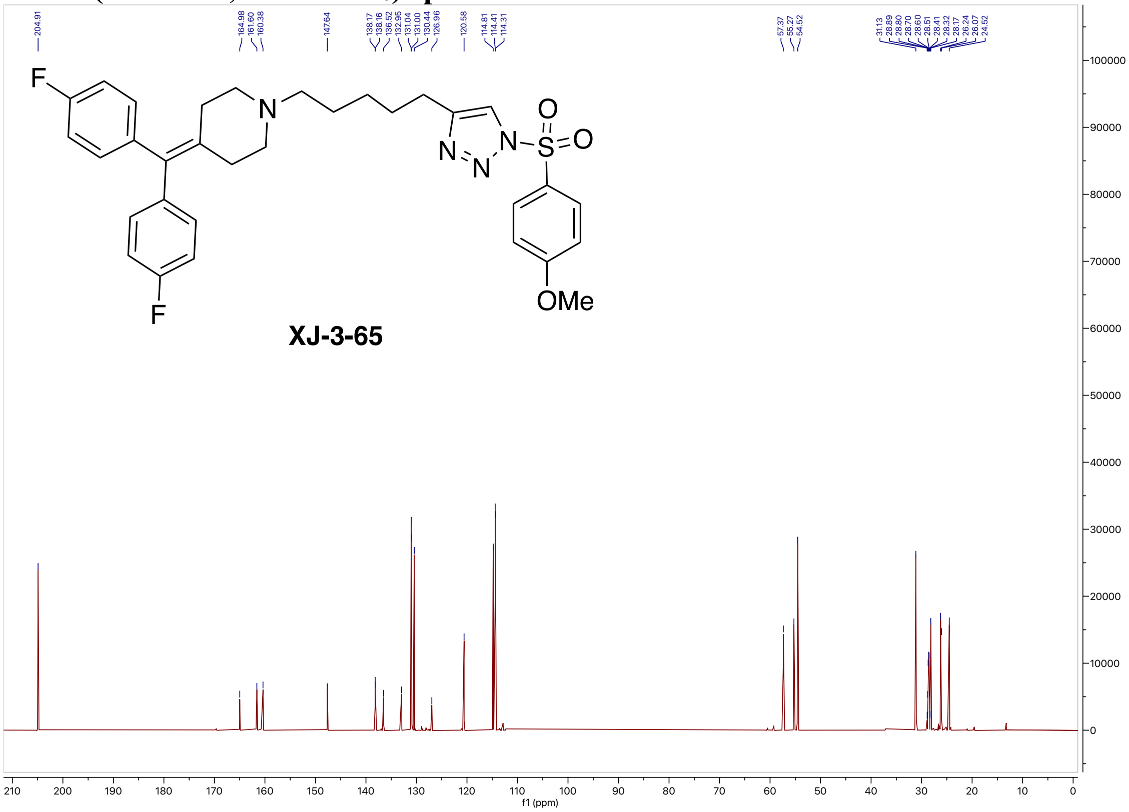

#### $^{19}$F NMR (564 MHz, Acetone-$d_6$) spectrum of XJ-3-65

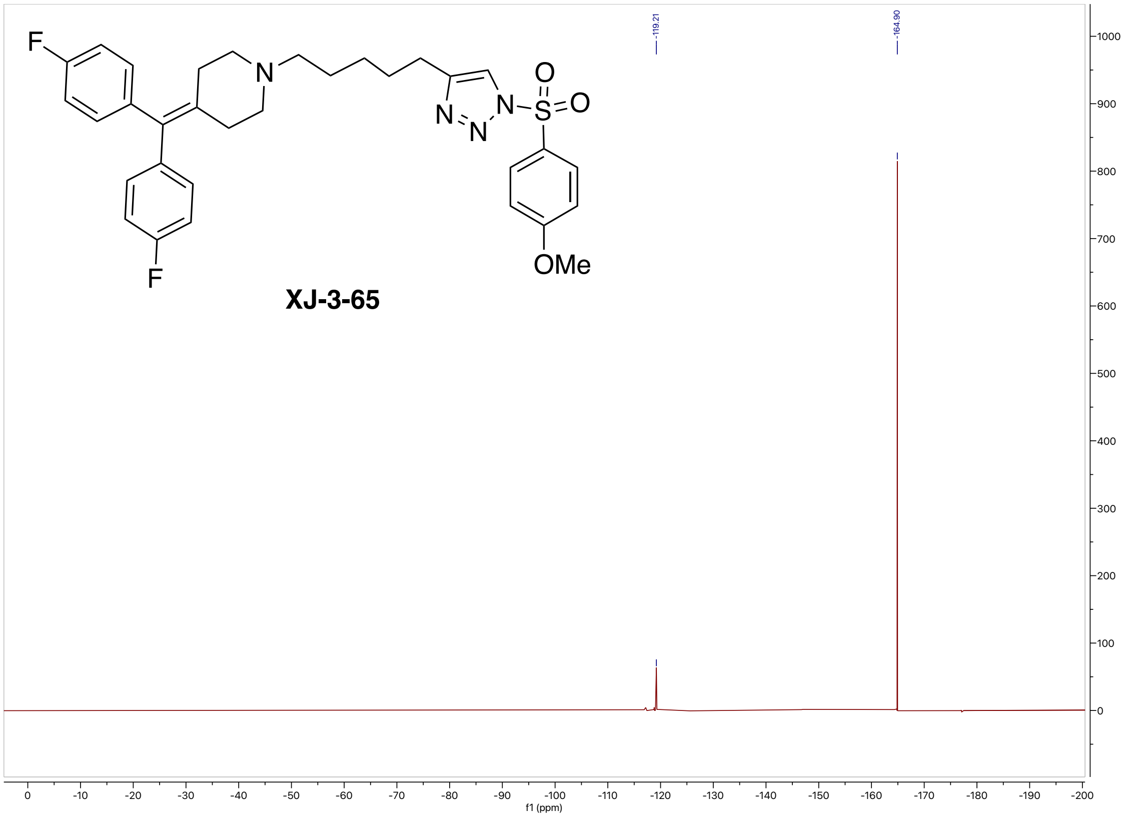

#### High-resolution mass spectrometry of XJ-3-65

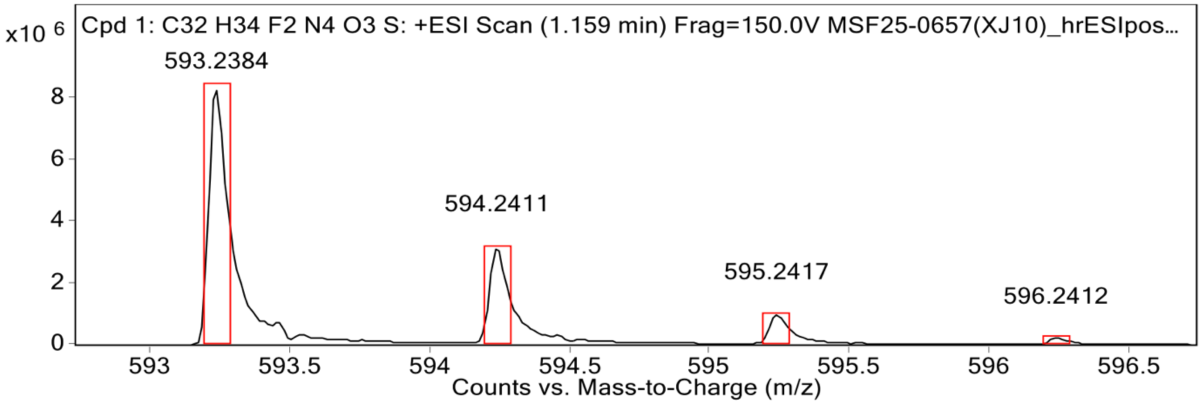


MS Spectrum Peak List

| Obs. m/z | Calc. m/z | Charge | Abundance | Formula | Ion Species | Tgt Mass Error (ppm) |
|---|---|---|---|---|---|---|
| 593.2384 | 593.2392 | 1 | 8346610 | C32H34F2N4O3S | (M+H)+ | 1.36 |
| 594.2411 | 594.2423 | 1 | 3215849 | C32H34F2N4O3S | (M+H)+ | 2.07 |
| 595.2417 | 595.2414 | 1 | 1010595 | C32H34F2N4O3S | (M+H)+ | -0.56 |
| 596.2412 | 596.2420 | 1 | 220327 | C32H34F2N4O3S | (M+H)+ | 1.3 |
| 607.2539 | | | 9153810 | | | |

#### HPLC spectrum of XJ-3-65

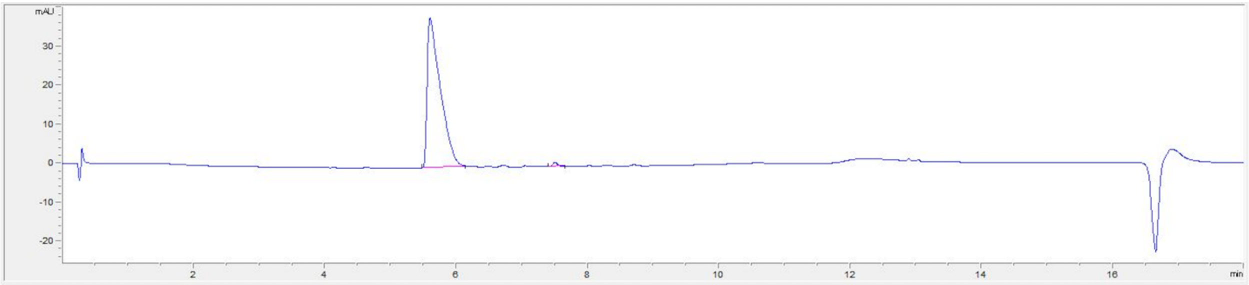


| # | Time | Area | Height | Width | Area% | Symmetry |
|---|---|---|---|---|---|---|
| 1 | 5.604 | 538.3 | 38.3 | 0.1939 | 98.956 | 0.28 |
| 2 | 7.506 | 5.7 | 1.1 | 0.0809 | 1.044 | 0.803 |

#### $^{1}$H NMR (800 MHz, Acetone-$d_6$) spectrum of XJ-4-5

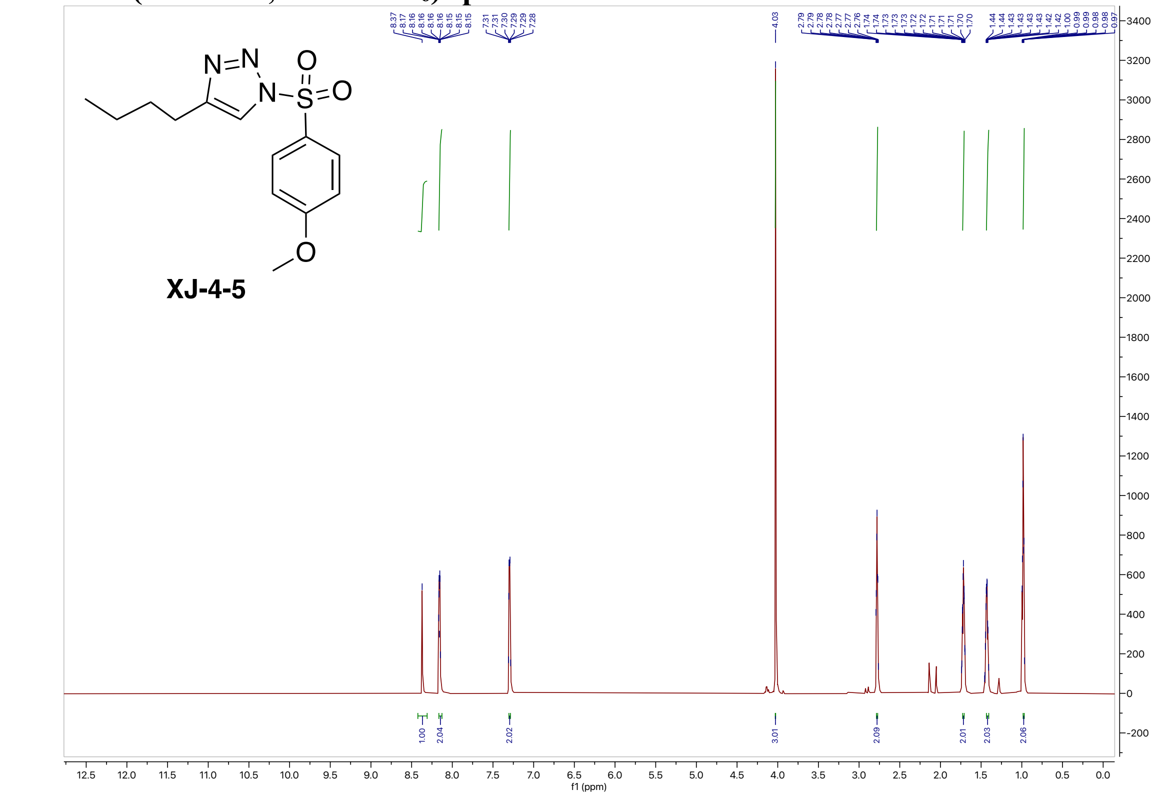


#### $^{13}$C NMR (200 MHz, Acetone-$d_6$) spectrum of XJ-4-5

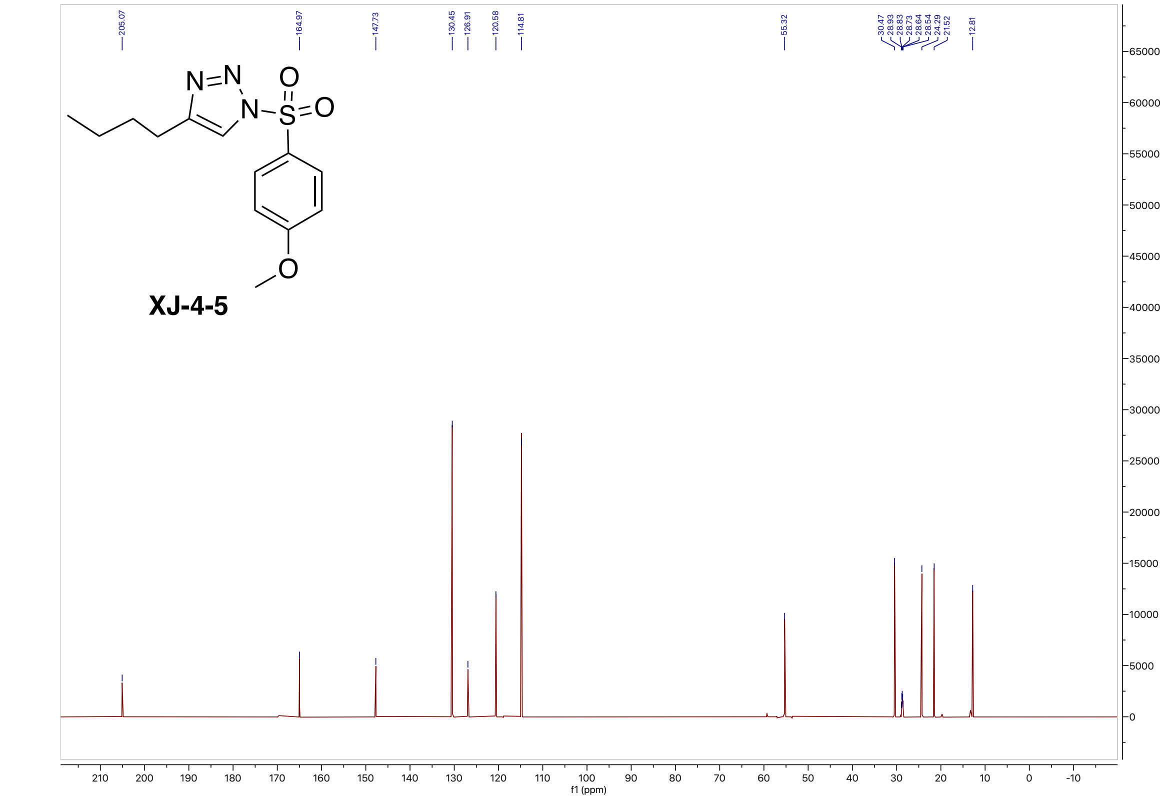

#### High-resolution mass spectrometry of XJ-4-5

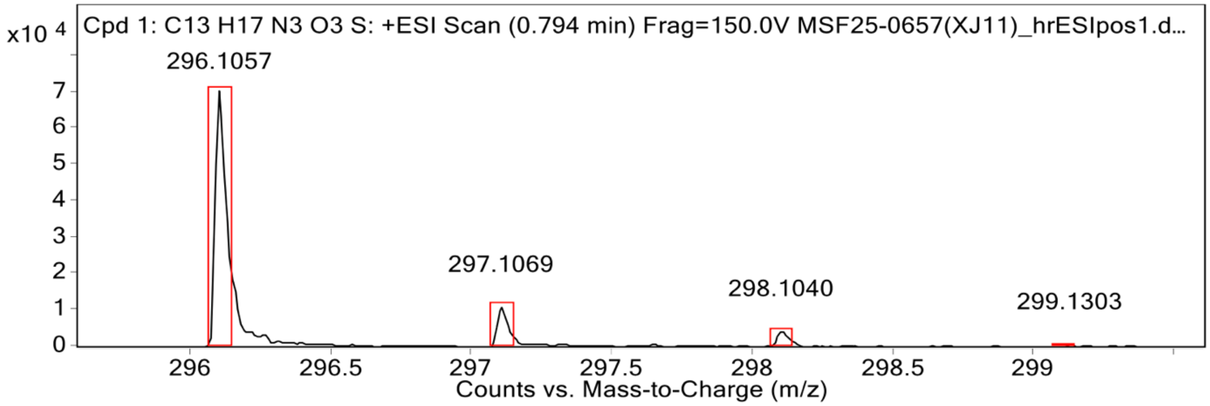


**MS Spectrum Peak List**

| Obs. m/z | Calc. m/z | Charge | Abundance | Formula | Ion Species | Tgt Mass Error (ppm) |
|---|---|---|---|---|---|---|
| 296.1057 | 296.1063 | 1 | 71186 | C13H17N3O3S | (M+H)+ | 2.28 |
| 297.1069 | 297.1091 | 1 | 11029 | C13H17N3O3S | (M+H)+ | 7.46 |
| 298.1040 | 298.1049 | 1 | 4371 | C13H17N3O3S | (M+H)+ | 3.05 |
| 299.1303 | 299.1067 | 1 | 764 | C13H17N3O3S | (M+H)+ | -78.91 |
| 607.2530 | | | 641157 | | | |

#### HPLC spectrum of XJ-4-5

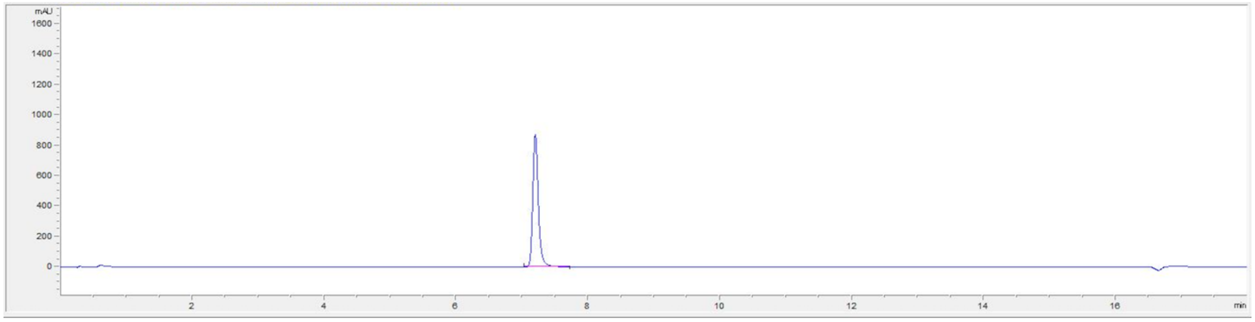


| # | Time | Area | Height | Width | Area% | Symmetry |
|---|---|---|---|---|---|---|
| 1 | 7.199 | 5117.9 | 874.4 | 0.0883 | 100.000 | 0.751 |

#### $^{1}H$ NMR (800 MHz, Acetone-$d_6$) spectrum of XJ-4-27

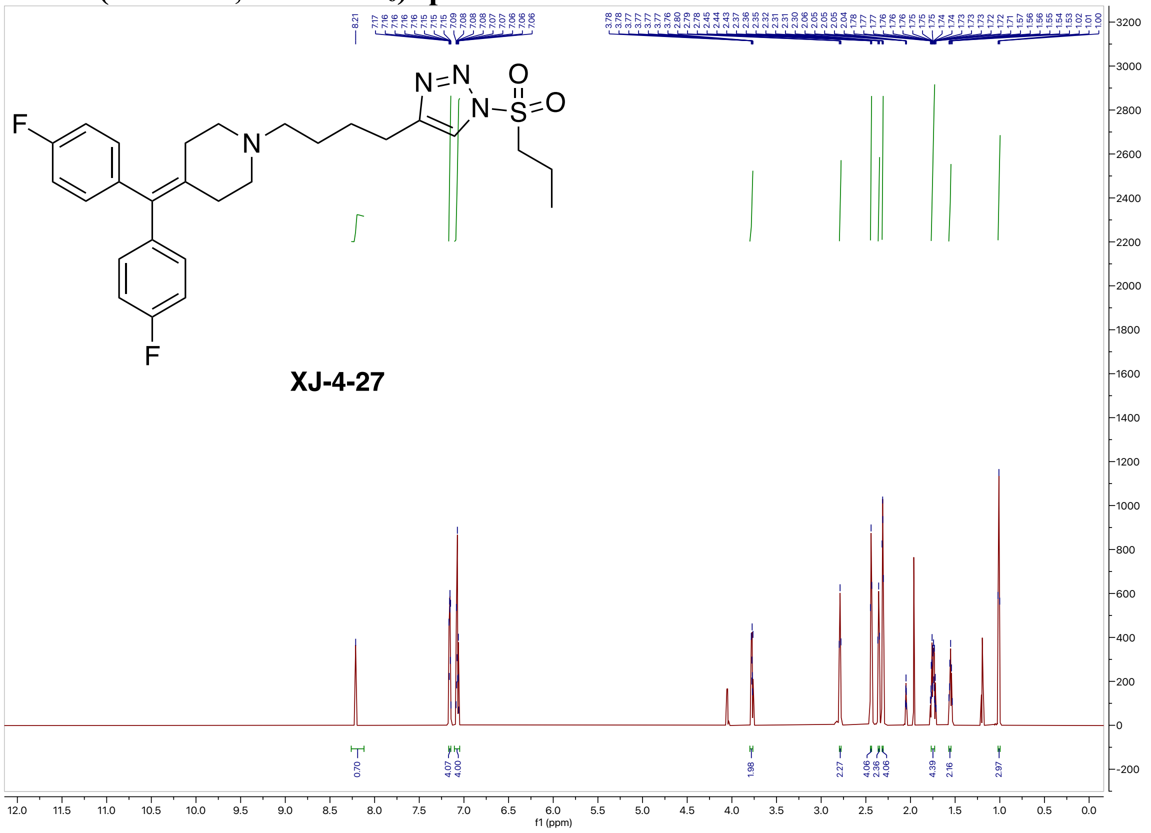


#### $^{13}C$ NMR (200 MHz, Acetone-$d_6$) spectrum of XJ-4-27

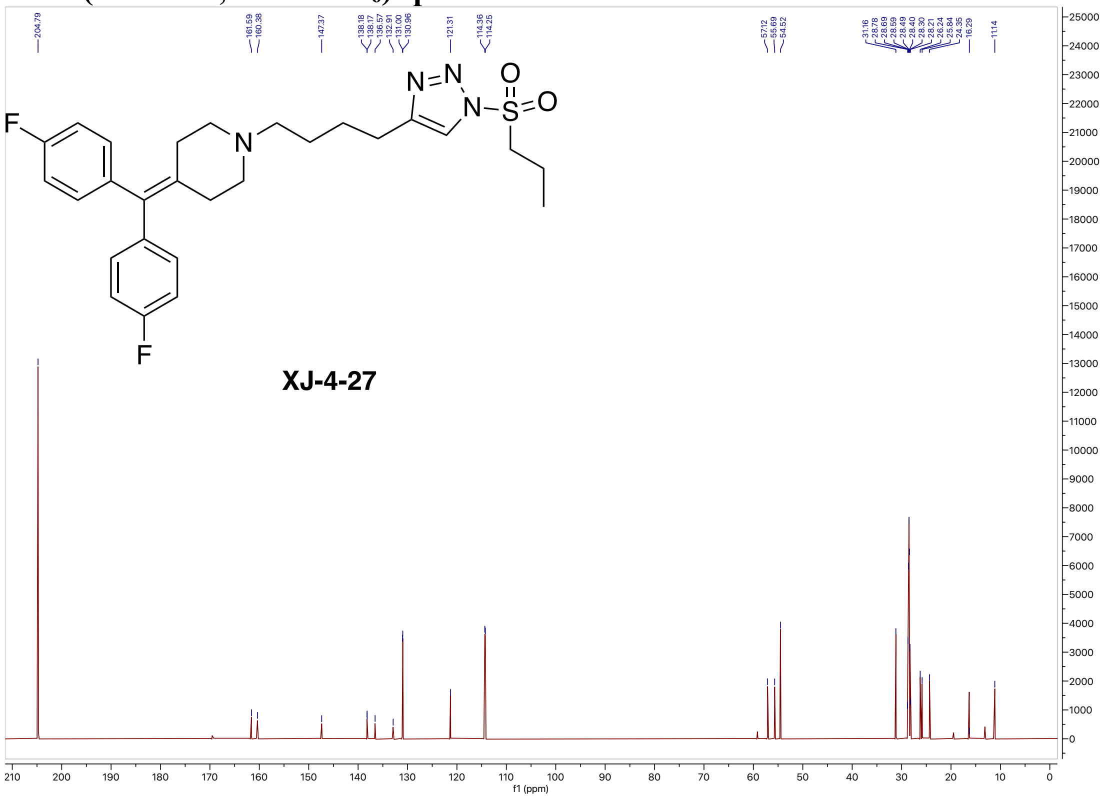

#### $^{19}$F NMR (564 MHz, Acetone-$d_6$) spectrum of XJ-4-27

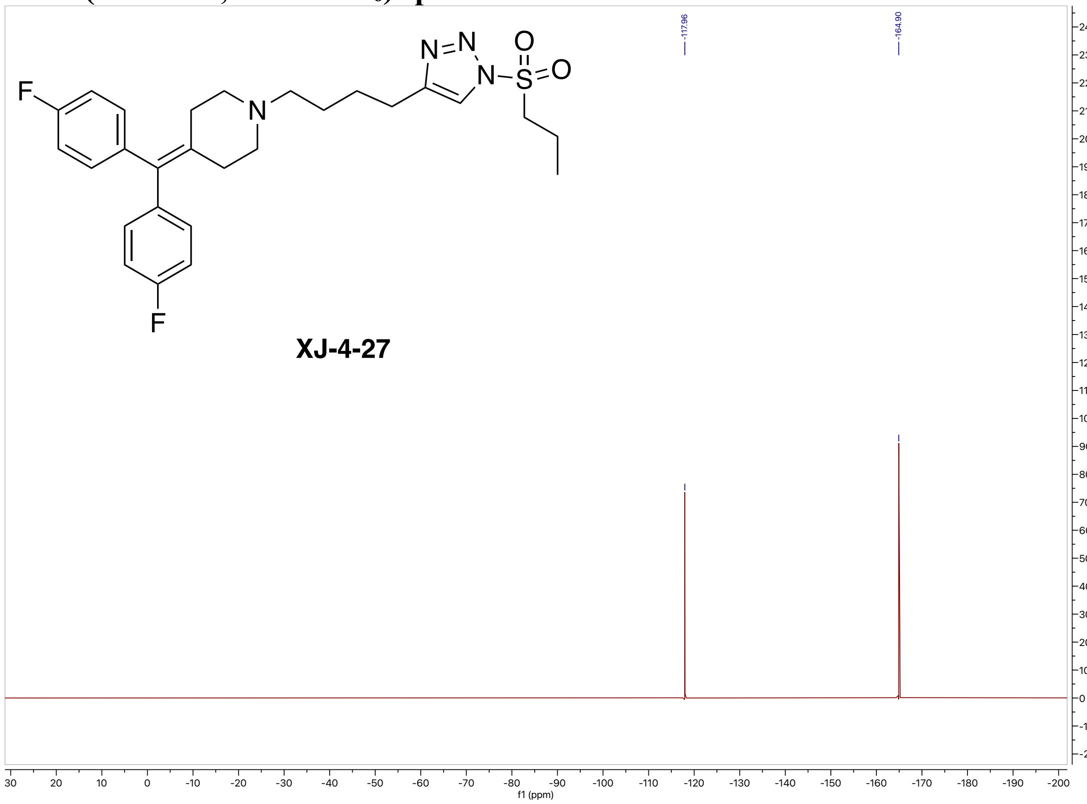


#### High-resolution mass spectrometry of XJ-4-27

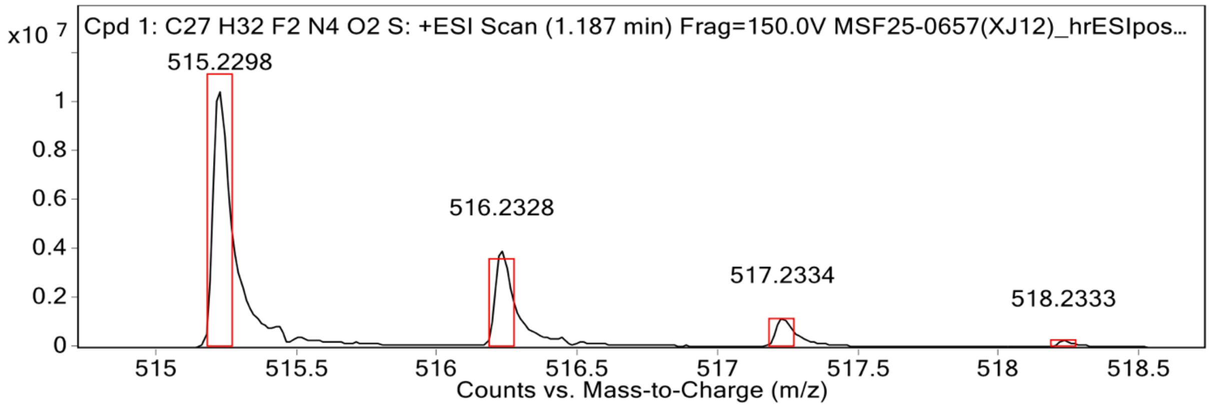


**MS Spectrum Peak List**

| Obs. m/z | Calc. m/z | Charge | Abundance | Formula | Ion Species | Tgt Mass Error (ppm) |
|---|---|---|---|---|---|---|
| 515.2298 | 515.2287 | 1 | 10591421 | C27H32F2N4O2S | (M+H)+ | -2.26 |
| 516.2328 | 516.2317 | 1 | 3986373 | C27H32F2N4O2S | (M+H)+ | -2.13 |
| 517.2334 | 517.2300 | 1 | 1218145 | C27H32F2N4O2S | (M+H)+ | -6.67 |
| 518.2333 | 518.2306 | 1 | 258297 | C27H32F2N4O2S | (M+H)+ | -5.2 |
| 519.2347 | 519.2321 | 1 | 42466 | C27H32F2N4O2S | (M+H)+ | -5.12 |

**HPLC spectrum of XJ-4-27**

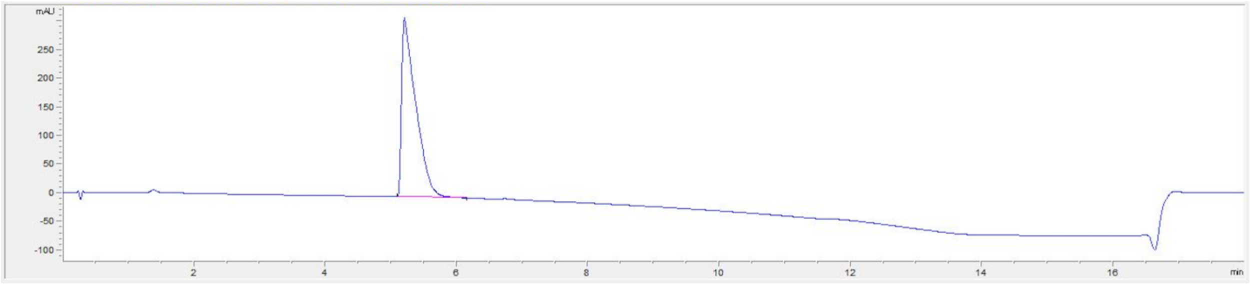

| # | Time | Area | Height | Width | Area% | Symmetry |
|---|---|---|---|---|---|---|
| 1 | 5.205 | 4682.8 | 312.7 | 0.2027 | 100.000 | 0.231 |

**$^{1}$H NMR (800 MHz, Acetone-$d_6$) spectrum of XJ-4-71**

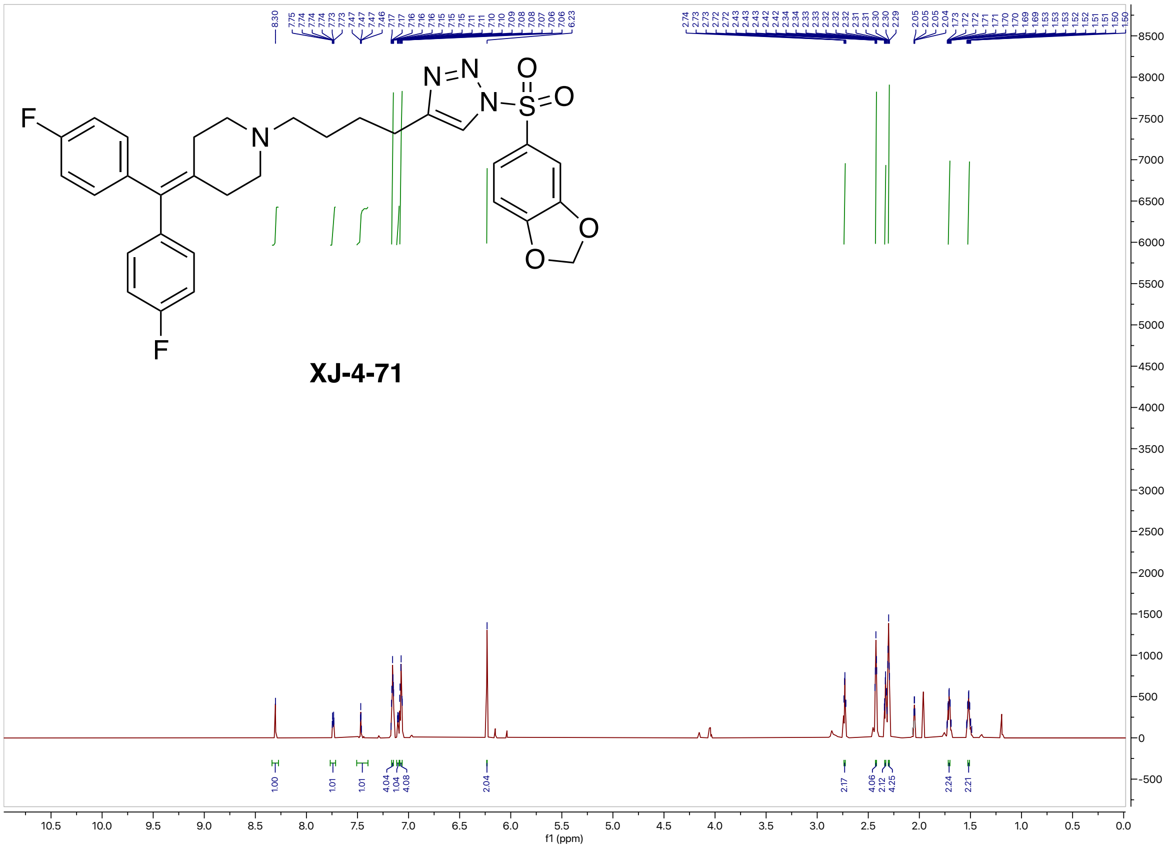

#### $^{13}$C NMR (200 MHz, Acetone-$d_6$) spectrum of XJ-4-71

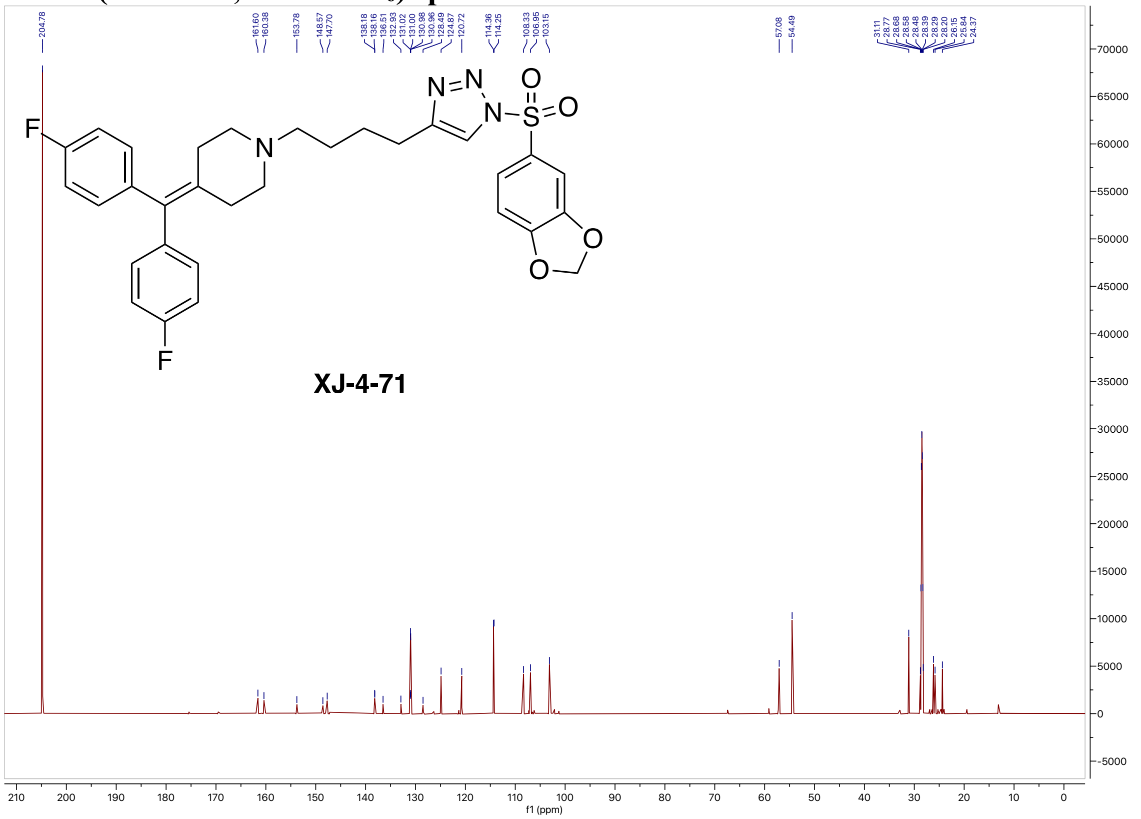

#### $^{19}$F NMR (564 MHz, Acetone-$d_6$) spectrum of XJ-4-71

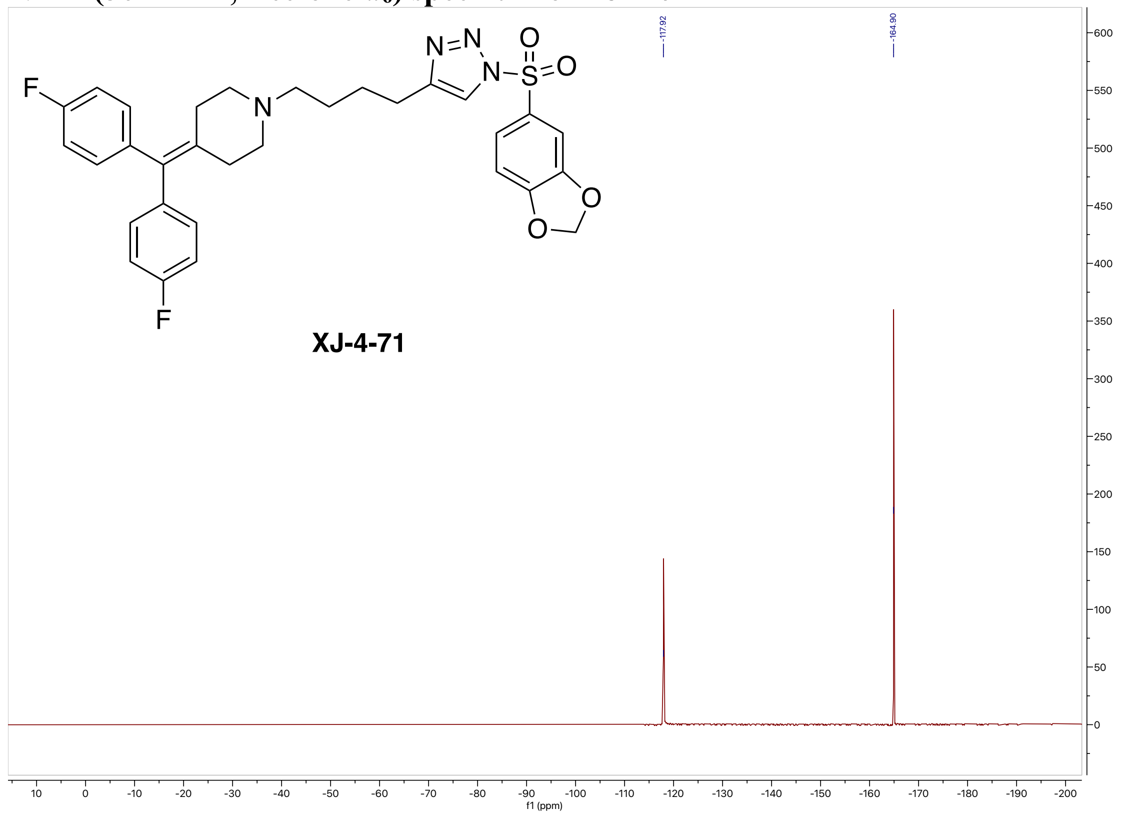

**High-resolution mass spectrometry of XJ-4-71**

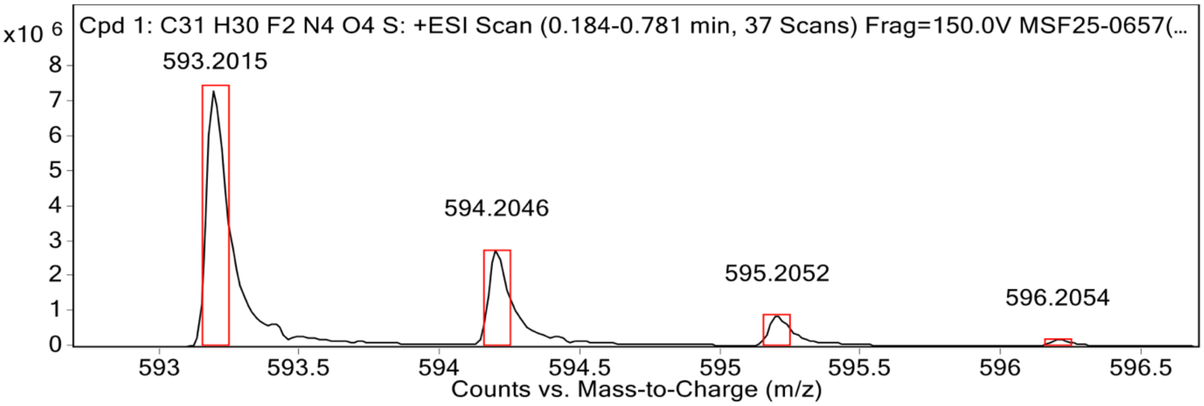


**MS Spectrum Peak List**

| Obs. m/z | Calc. m/z | Charge | Abundance | Formula | Ion Species | Tgt Mass Error (ppm) |
|---|---|---|---|---|---|---|
| 593.2015 | 593.2029 | 1 | 7377400 | C31H30F2N4O4S | (M+H)+ | 2.32 |
| 594.2046 | 594.2059 | 1 | 2747123 | C31H30F2N4O4S | (M+H)+ | 2.19 |
| 595.2052 | 595.2049 | 1 | 884225 | C31H30F2N4O4S | (M+H)+ | -0.59 |
| 596.2054 | 596.2056 | 1 | 185947 | C31H30F2N4O4S | (M+H)+ | 0.31 |
| 597.2189 | 597.2070 | 1 | 50594 | C31H30F2N4O4S | (M+H)+ | -19.96 |

**HPLC spectrum of XJ-4-71**

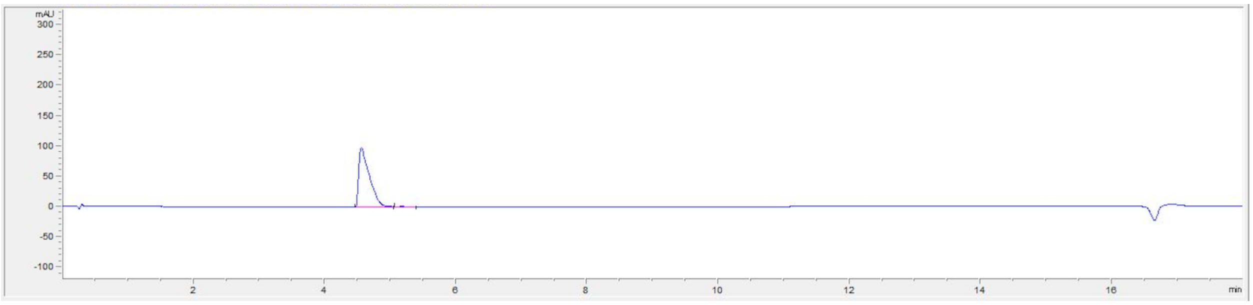

| # | Time | Area | Height | Width | Area% | Symmetry |
|---|---|---|---|---|---|---|
| 1 | 4.559 | 1112.3 | 98.7 | 0.1575 | 98.936 | 0.29 |
| 2 | 5.18 | 12 | 1.1 | 0.1437 | 1.064 | 0.855 |

#### $^{1}$H NMR (800 MHz, Acetone-$d_6$) spectrum of XJ-4-85

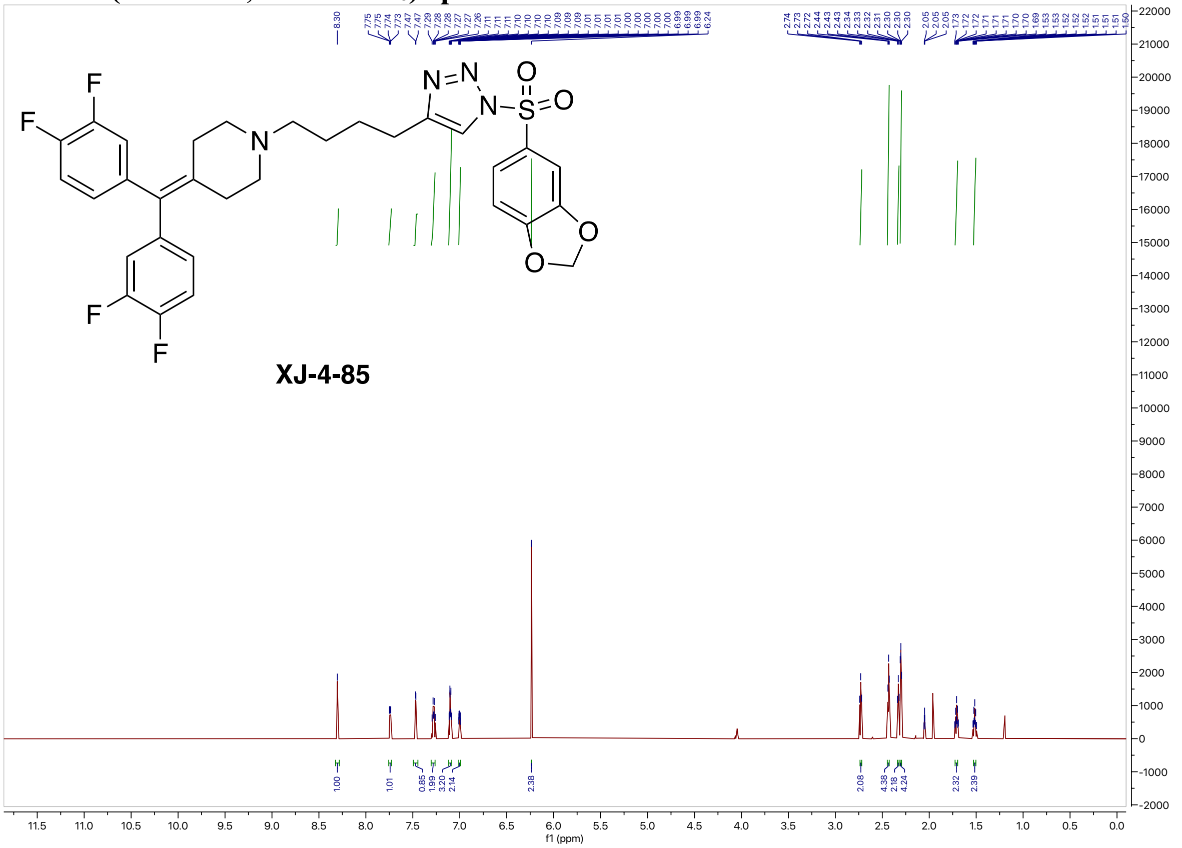


#### $^{13}$C NMR (126 MHz, Acetone-$d_6$) spectrum of XJ-4-85

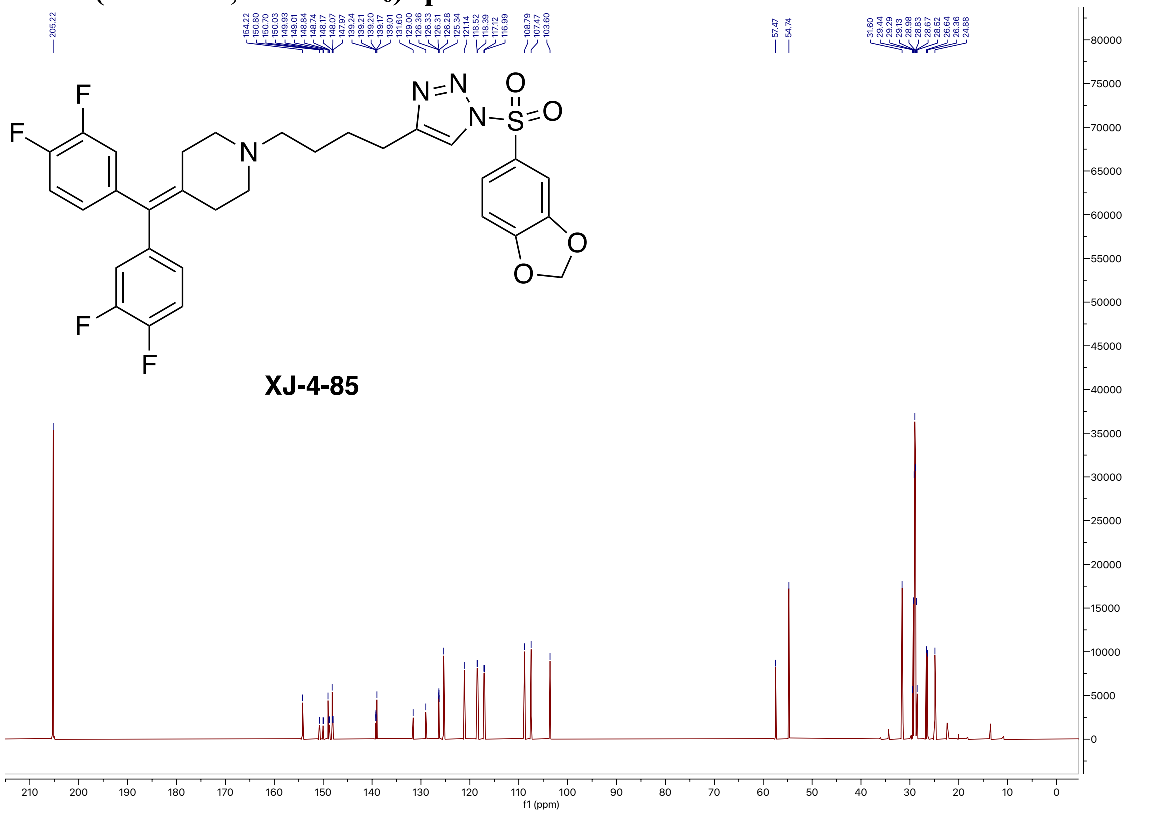

#### $^{19}$F NMR (471 MHz, Acetone-$d_6$) spectrum of XJ-4-85

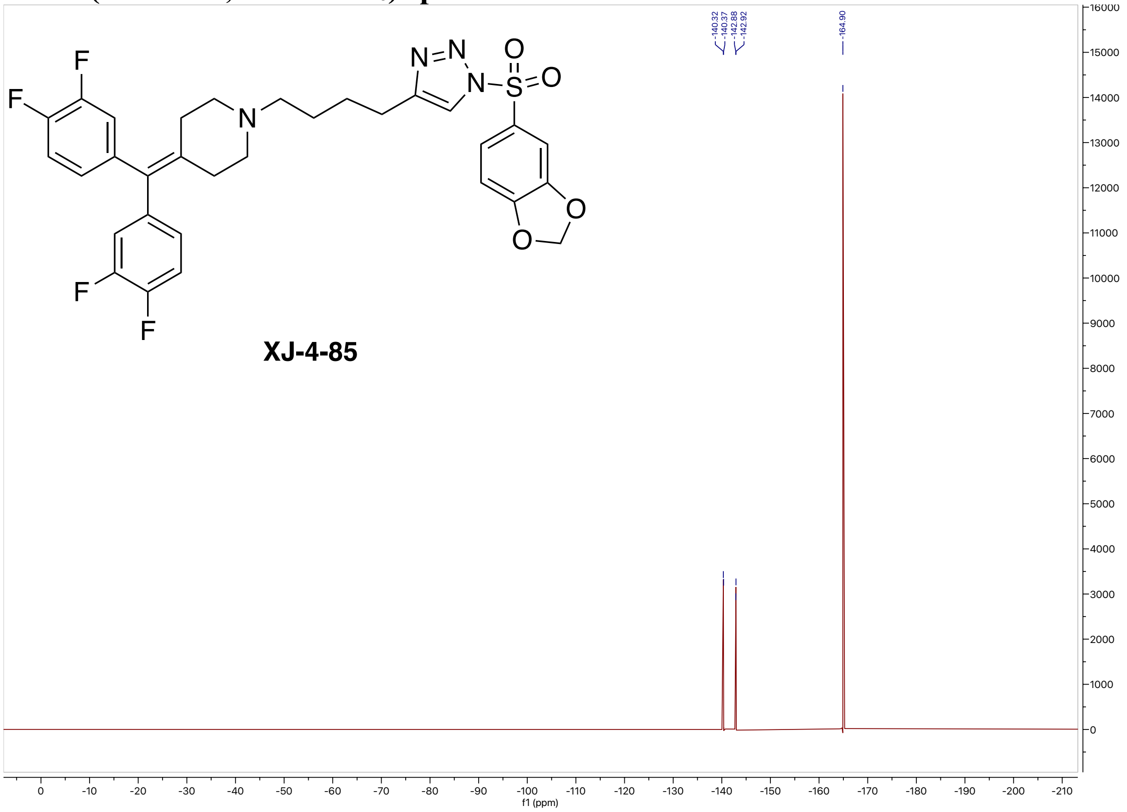


#### High-resolution mass spectrometry of XJ-4-85

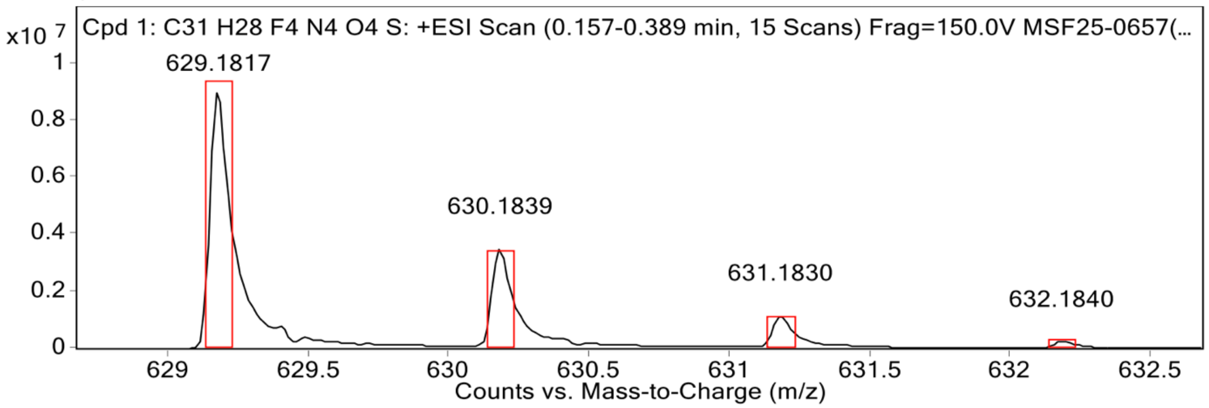


**MS Spectrum Peak List**

| Obs. m/z | Calc. m/z | Charge | Abundance | Formula | Ion Species | Tgt Mass Error (ppm) |
|---|---|---|---|---|---|---|
| 629.1817 | 629.1840 | 1 | 9123326 | C31H28F4N4O4S | (M+H)+ | 3.64 |
| 630.1839 | 630.1871 | 1 | 3505858 | C31H28F4N4O4S | (M+H)+ | 4.99 |
| 631.1830 | 631.1860 | 1 | 1169833 | C31H28F4N4O4S | (M+H)+ | 4.73 |
| 632.1840 | 632.1867 | 1 | 250715 | C31H28F4N4O4S | (M+H)+ | 4.34 |
| 633.1859 | 633.1881 | 1 | 50432 | C31H28F4N4O4S | (M+H)+ | 3.58 |

#### HPLC spectrum of XJ-4-85

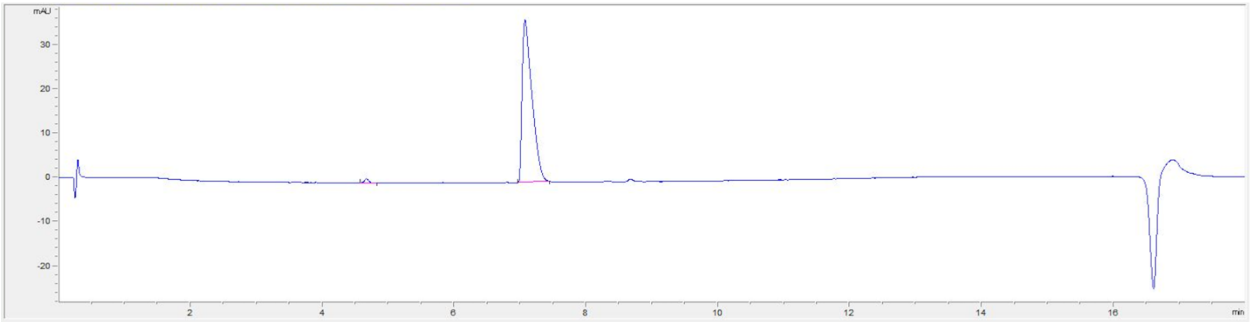

| # | Time | Area | Height | Width | Area% | Symmetry |
|---|---|---|---|---|---|---|
| 1 | 4.671 | 6 | 1 | 0.0959 | 1.551 | 0.784 |
| 2 | 7.072 | 381.7 | 36.8 | 0.1491 | 98.449 | 0.384 |

#### $^{1}$H NMR (800 MHz, Acetone-$d_6$) spectrum of XJ-4-97

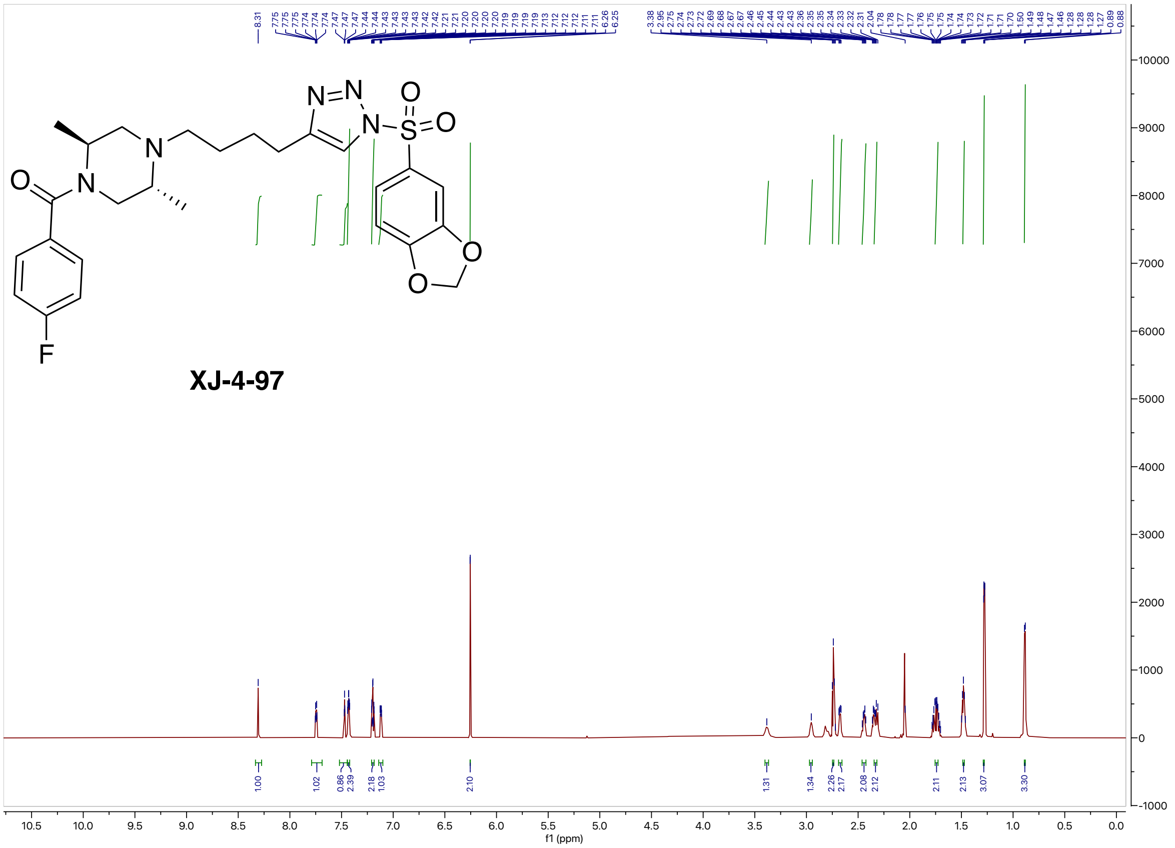

#### $^{13}C$ NMR (201 MHz, Acetone-$d_6$) spectrum of XJ-4-97

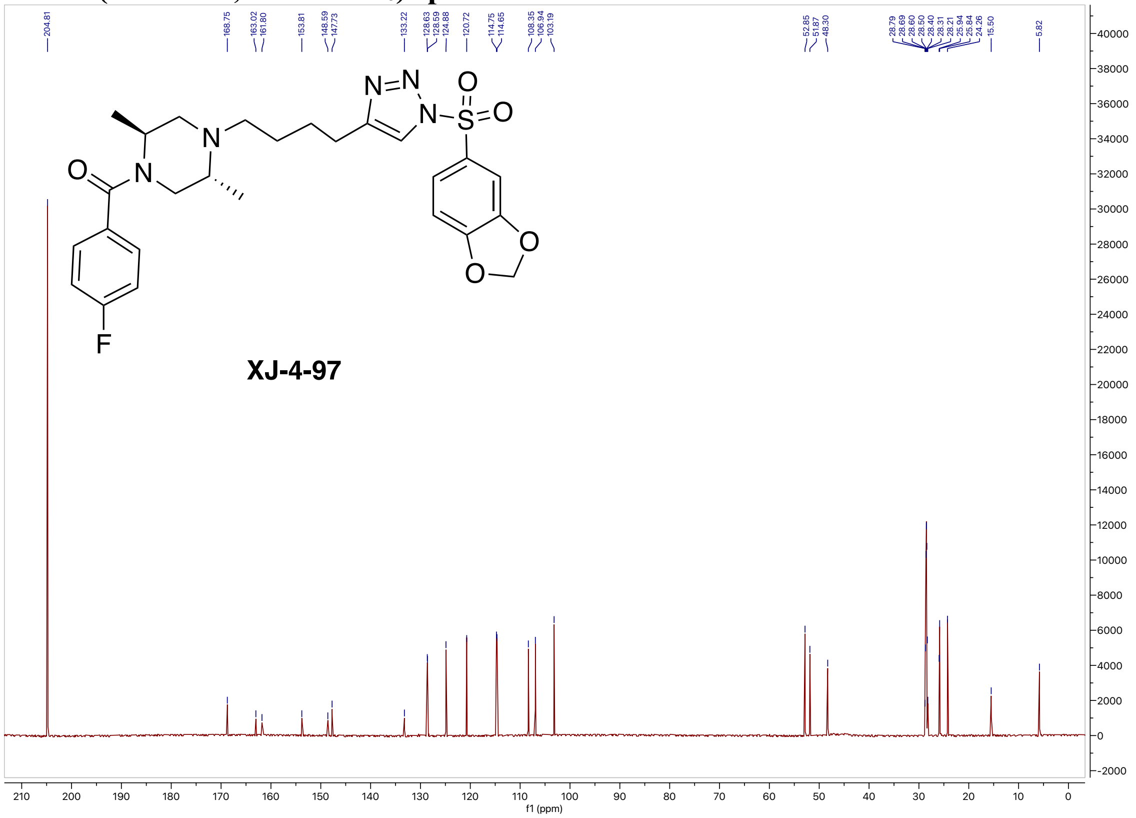

#### $^{19}F$ NMR (471 MHz, Acetone-$d_6$) spectrum of XJ-4-97

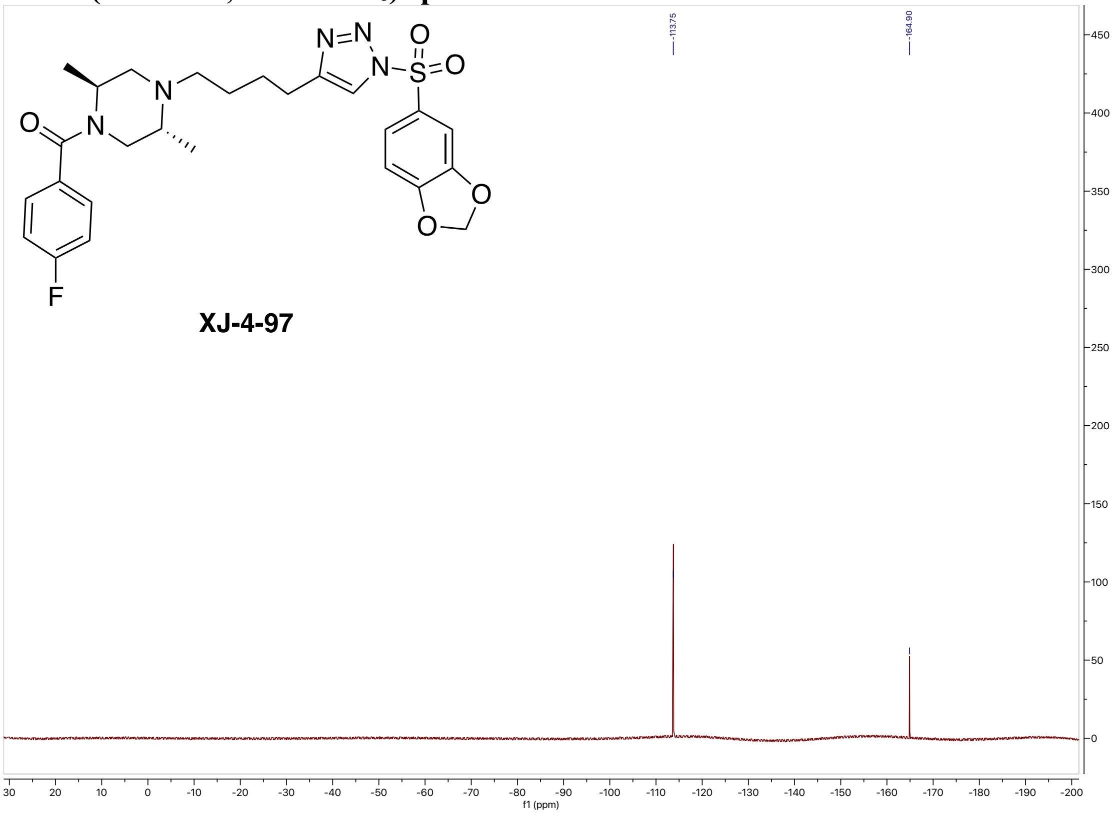

**High-resolution mass spectrometry of XJ-4-97**

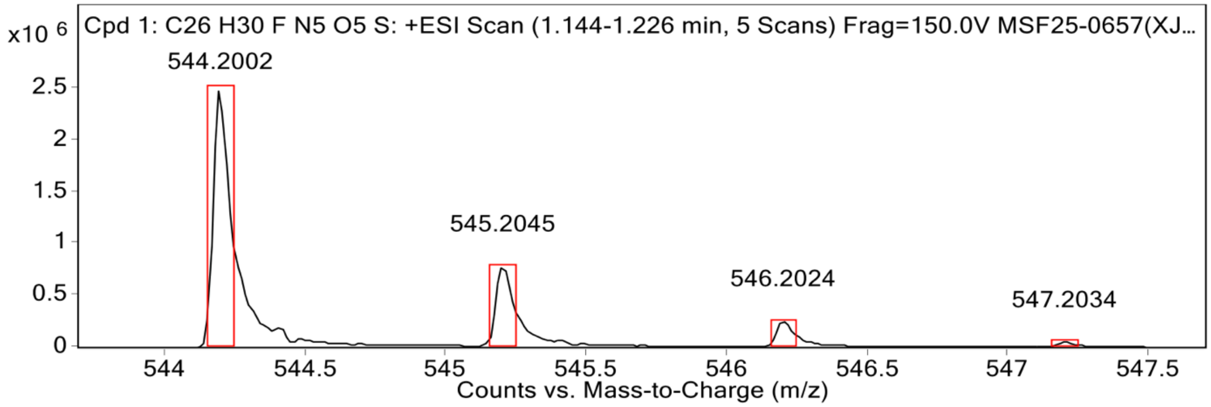


**MS Spectrum Peak List**

| Obs. m/z | Calc. m/z | Charge | Abundance | Formula | Ion Species | Tgt Mass Error (ppm) |
|---|---|---|---|---|---|---|
| 544.2002 | 544.2024 | 1 | 2492194 | C26H30FN5O5S | (M+H)+ | 4.1 |
| 545.2045 | 545.2054 | 1 | 782368 | C26H30FN5O5S | (M+H)+ | 1.62 |
| 546.2024 | 546.2037 | 1 | 254228 | C26H30FN5O5S | (M+H)+ | 2.37 |
| 547.2034 | 547.2046 | 1 | 51287 | C26H30FN5O5S | (M+H)+ | 2.33 |
| 566.1822 | | | 4702052 | | | |

**HPLC spectrum of XJ-4-97**

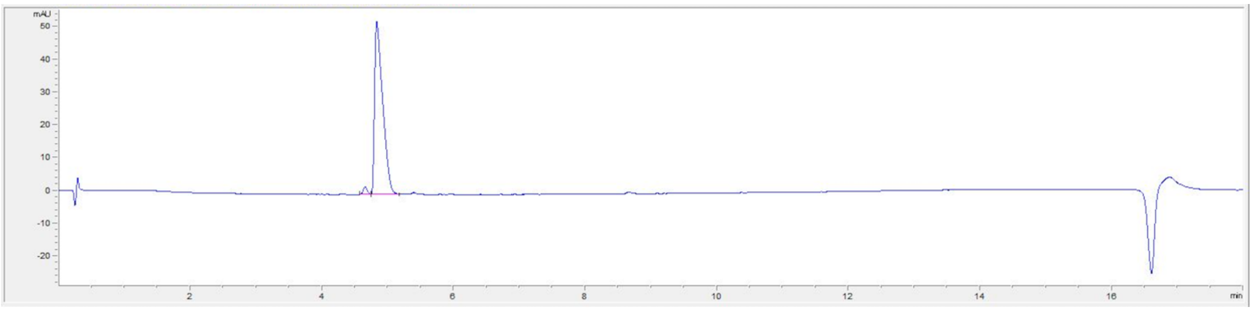


| # | Time | Area | Height | Width | Area% | Symmetry |
|---|---|---|---|---|---|---|
| 1 | 4.659 | 10.1 | 2.3 | 0.0696 | 2.201 | 0.847 |
| 2 | 4.834 | 448.8 | 52.8 | 0.1211 | 97.799 | 0.342 |

#### $^{1}$H NMR (500 MHz, Acetone-$d_6$) spectrum of XJ-4-119

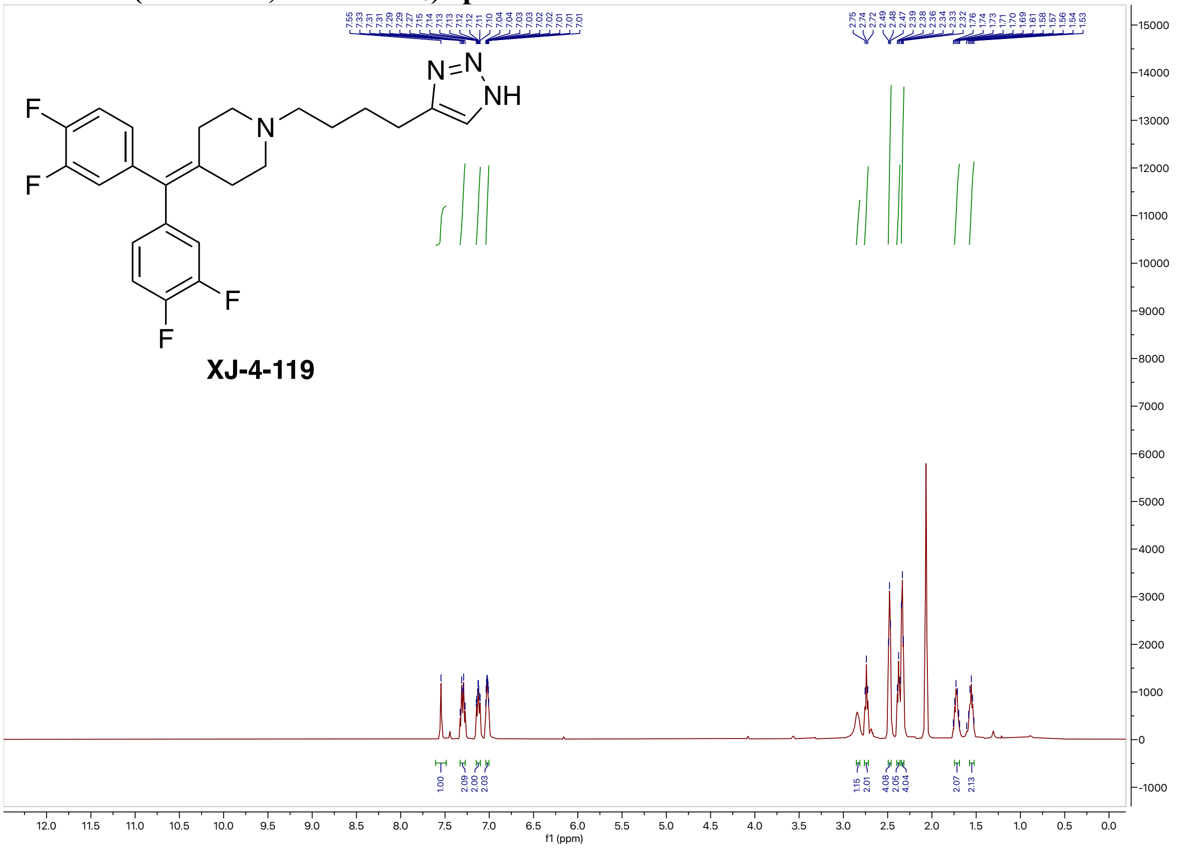


#### $^{13}$C NMR (126 MHz, Acetone-$d_6$) spectrum of XJ-4-119

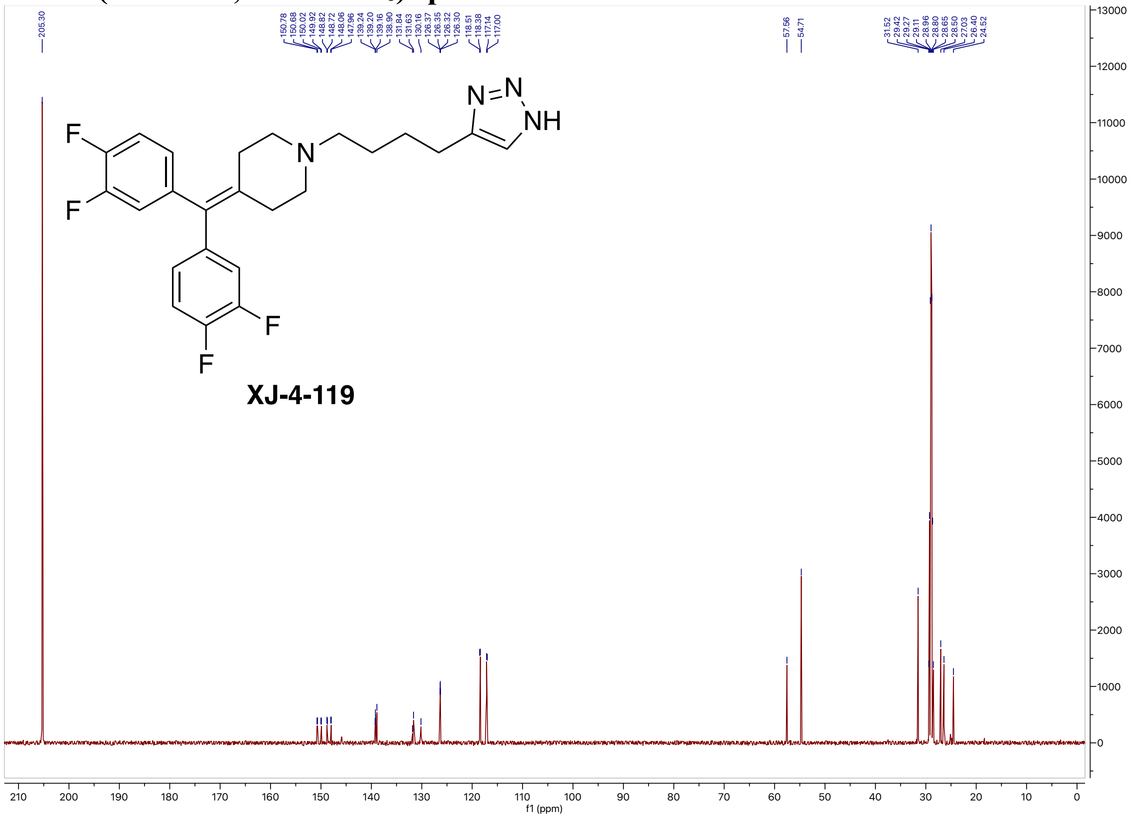

#### $^{19}$F NMR (471 MHz, Acetone-$d_6$) spectrum of XJ-4-119

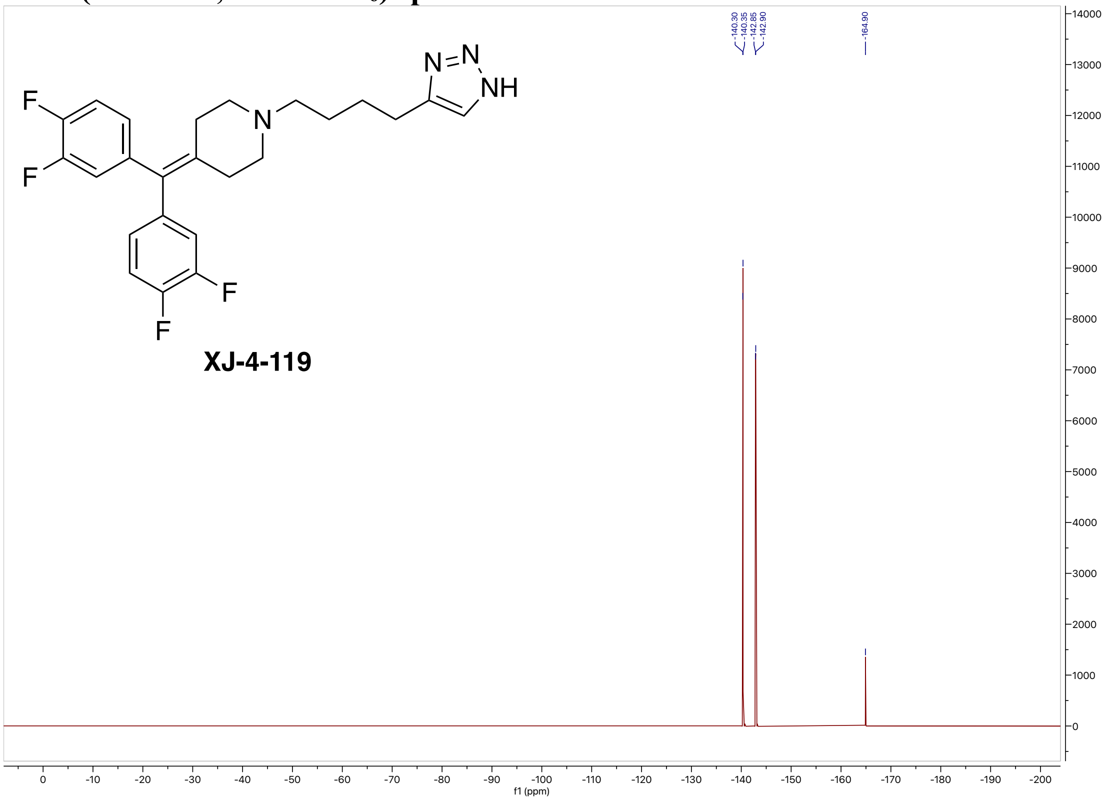


#### High-resolution mass spectrometry of XJ-4-119

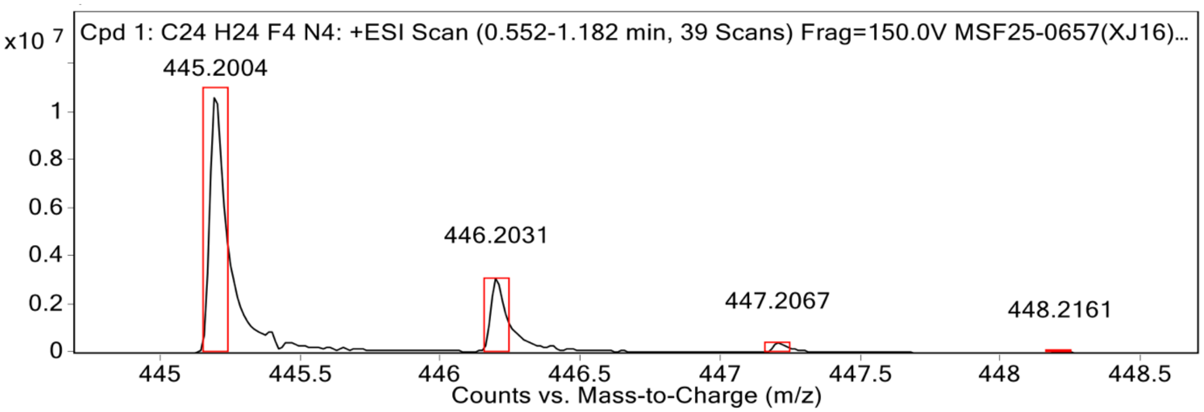


**MS Spectrum Peak List**

| Obs. m/z | Calc. m/z | Charge | Abundance | Formula | Ion Species | Tgt Mass Error (ppm) |
|---|---|---|---|---|---|---|
| 445.2004 | 445.2010 | 1 | 10907299 | C24H24F4N4 | (M+H)+ | 1.29 |
| 446.2031 | 446.2040 | 1 | 3127257 | C24H24F4N4 | (M+H)+ | 2.21 |
| 447.2067 | 447.2071 | 1 | 400560 | C24H24F4N4 | (M+H)+ | 0.81 |
| 448.2161 | 448.2101 | 1 | 32815 | C24H24F4N4 | (M+H)+ | -13.32 |

**HPLC spectrum of XJ-4-119**

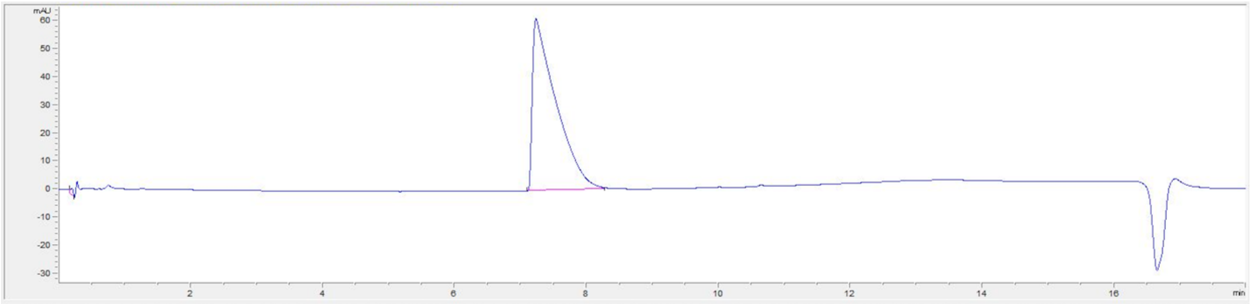


| # | Time | Area | Height | Width | Area% | Symmetry |
|---|---|---|---|---|---|---|
| 1 | 0.205 | 6.8 | 2.4 | 0.0434 | 0.445 | 1.454 |
| 2 | 7.235 | 1523.1 | 61.4 | 0.3316 | 99.555 | 0.165 |
